## Supplementary material for "Analysis of meiosis in *Pristionchus pacificus* reveals plasticity in homolog pairing and synapsis in the nematode lineage": Table S6 X0 genotypes

| chr | MB | f2.0 | f2.0 | f2.0 | f2.0 | f2.0 | f2.0 | f2.0 | f2.0 | f2.0 | f2.0 | f2.0 | f2.0 | f2.0 | f2.0 | f2.0 | f2.0 | f2.0 | f2.0 | f2.0 | f2.0 | f2.0 | f2.0 | f2.0 | f2.0 | f2.0 | f2.0 | f2.0 | f2.0 | f2.0 | f2.0 | f2.0 | f2.0 | f2.0 | f2.0 | f2.0 | f2.0 | f2.0 | f2.0 | f2.0 | f2.0 | f2.0 | f2.0 | f2.0 | f2.0 | f2.0 | f2.0 | f2.0 | f2.0 | f2.0 | f2.0 | f2.0 | f2.0 | f2.0 | f2.0 | f2.0 | f2.0 | f2.0 | f2.0 | f2.0 | f2.0 | f2.0 | f2.0 | f2.0 | f2.0 | f2.0 | f2.0 | f2.0 | f2.0 | f2.0 | f2.0 | f2.0 | f2.0 | f2.0 | f2.0 | f2.0 | f2.0 | f2.0 | f2.0 | f2.0 | f2.0 | f2.0 | f2.0 | f2.0 | f2.0 | f2.0 | f2.0 | f2.0 | f2.0 | f2.0 | f2.0 | f2.0 | f2.093 |
|---|---|---|---|---|---|---|---|---|---|---|---|---|---|---|---|---|---|---|---|---|---|---|---|---|---|---|---|---|---|---|---|---|---|---|---|---|---|---|---|---|---|---|---|---|---|---|---|---|---|---|---|---|---|---|---|---|---|---|---|---|---|---|---|---|---|---|---|---|---|---|---|---|---|---|---|---|---|---|---|---|---|---|---|---|---|---|---|---|---|---|---|---|---|
| Chrl | 0 | A | W | W | NA | W | W | W | NA | W | A | A | W | W | W | W | W | NA | W | W | NA | W | A | A | A | A | NA | A | W | NA | A | W | W | W | W | A | A | A | W | A | W | W | A | NA | A | A | A | W | A | W | W | A | A | NA | A | A | A | A | A | W | A | A | A | W | W | W | NA | W | A | W | A | A | A | W | W | W | A | W | W | A | A | W | A | A | NA | A | W | NA | A | W | A | A | W |
| Chrl | 0.1 | A | W | W | W | W | W | W | W | W | A | A | W | W | W | W | W | W | W | W | W | W | A | A | A | A | A | A | W | W | A | W | W | W | W | A | A | A | W | A | W | W | A | W | A | A | A | W | A | W | W | A | A | NA | A | A | A | A | A | W | A | A | A | W | W | W | W | W | A | W | A | A | A | W | W | W | A | W | W | A | A | W | A | A | W | A | W | A | A | W | A | A | W |
| Chrl | 0.2 | A | W | W | NA | W | W | W | W | W | A | A | W | W | W | W | W | W | W | W | W | W | A | A | A | A | A | A | W | W | A | W | W | W | W | A | A | A | W | A | W | W | A | W | A | A | A | W | A | W | NA | A | A | W | A | A | A | A | A | W | A | A | A | W | W | W | W | W | A | W | A | A | A | W | W | W | A | W | W | A | A | W | A | A | W | A | W | A | A | W | A | A | W |
| Chrl | 0.3 | A | W | W | W | W | W | W | W | W | A | A | W | W | W | W | W | W | W | W | W | W | A | A | A | NA | A | A | W | W | A | W | W | W | W | A | A | A | W | A | W | W | A | W | A | A | A | W | A | W | W | A | A | W | A | A | A | A | A | W | A | A | A | W | W | W | W | W | A | W | A | A | A | W | W | W | A | W | W | A | A | W | A | A | W | A | W | A | A | W | A | A | W |
| Chrl | 0.4 | A | W | W | W | W | W | W | W | W | A | A | W | W | W | W | W | NA | W | W | W | W | A | A | A | A | A | A | W | W | A | W | W | W | W | A | A | A | W | A | W | W | A | W | A | A | A | W | A | W | W | A | A | W | A | A | A | A | A | W | A | A | A | W | W | W | NA | W | A | W | A | A | A | W | W | W | A | W | W | A | A | W | A | A | W | A | W | A | A | W | A | A | W |
| Chrl | 0.5 | A | W | W | W | W | W | W | W | W | A | A | W | W | W | W | W | W | W | W | W | W | A | A | A | A | A | A | W | W | A | W | W | W | W | A | A | A | W | A | W | W | A | W | A | A | A | W | A | W | NA | A | A | W | A | A | A | A | A | W | A | A | A | W | W | W | W | W | A | W | A | A | A | W | W | W | A | W | W | A | A | W | A | A | W | A | W | A | A | W | A | A | W |
| Chrl | 0.6 | A | W | W | W | W | W | W | W | W | A | A | W | W | W | W | W | W | W | W | W | W | A | A | A | A | A | A | W | W | A | W | W | W | W | A | A | A | W | A | W | W | A | W | A | A | A | W | A | W | NA | A | A | W | A | A | A | A | A | W | A | A | A | W | W | W | W | W | A | W | A | A | A | W | W | W | A | W | W | A | A | W | A | A | W | A | W | A | A | W | A | A | W |
| Chrl | 0.7 | A | W | W | W | W | W | W | W | W | A | A | NA | W | W | W | W | W | W | W | W | W | A | A | A | A | A | A | W | W | A | W | W | W | W | A | A | A | W | A | W | W | A | W | A | A | A | W | A | W | W | A | A | W | A | A | A | A | A | W | A | A | A | W | W | W | W | W | A | W | A | A | A | W | W | W | A | W | W | A | A | W | A | A | W | A | W | A | A | W | A | A | W |
| Chrl | 0.8 | A | W | W | W | W | W | W | W | W | A | A | A | W | W | W | W | W | W | W | W | W | A | A | A | A | A | A | W | W | A | W | W | W | W | A | A | A | W | A | W | W | A | W | A | A | A | W | A | W | W | A | A | W | A | A | A | A | A | W | A | A | A | W | W | W | W | W | A | W | A | A | A | W | W | W | A | W | W | A | A | W | A | A | W | A | W | A | A | W | A | A | W |
| Chrl | 0.9 | A | W | W | W | W | W | W | W | W | A | A | A | W | W | W | W | W | W | W | W | W | A | A | A | A | A | A | W | W | A | W | W | W | W | A | A | A | W | A | W | W | A | W | A | A | A | W | A | W | W | A | A | W | A | A | A | A | A | W | A | A | A | W | W | W | W | W | A | W | A | A | A | W | W | W | A | W | W | A | A | W | A | A | W | A | W | A | A | W | A | A | W |
| Chrl | 1 | A | W | W | W | W | W | W | W | W | A | A | A | W | W | W | W | W | W | W | W | W | A | A | A | A | A | A | W | W | A | W | W | W | W | A | A | A | W | A | W | W | A | W | A | A | A | W | A | W | NA | A | A | W | A | A | A | A | A | W | A | A | A | W | W | W | W | W | A | W | A | A | A | W | W | W | A | W | W | A | A | W | A | A | W | A | W | A | A | W | A | A | W |
| Chrl | 1.1 | A | W | W | W | W | W | W | W | W | A | A | A | W | W | W | W | W | W | W | W | W | A | A | A | NA | A | A | W | W | A | W | W | W | W | A | A | A | W | A | W | W | A | W | A | A | A | W | A | W | W | A | A | W | A | A | A | A | A | W | A | A | A | W | W | W | W | W | A | W | A | A | A | W | W | W | A | NA | W | A | A | W | A | A | W | A | W | A | A | W | A | A | W |
| Chrl | 1.3 | A | W | A | W | W | W | W | W | W | A | A | A | W | NA | W | W | W | W | W | W | W | A | A | A | A | A | A | W | W | A | W | W | W | W | A | A | A | W | A | W | W | A | W | A | W | A | W | A | W | W | A | A | W | A | A | A | A | A | W | A | A | A | W | NA | W | W | NA | A | W | A | A | A | W | W | W | A | W | W | A | A | W | A | A | W | A | W | A | A | W | A | A | W |
| Chrl | 1.5 | A | W | A | W | W | W | W | W | W | A | A | A | W | A | W | W | W | W | W | W | W | A | A | A | A | A | A | W | W | A | W | W | W | W | A | A | A | W | A | W | W | A | W | A | W | A | W | A | W | W | A | A | W | A | A | A | A | A | W | A | A | W | W | W | W | W | W | A | W | A | A | A | W | W | W | A | W | W | A | A | W | A | W | W | A | W | A | A | W | A | A | W |
| Chrl | 1.6 | A | W | A | W | W | W | W | W | W | A | A | A | W | A | W | W | W | W | W | W | W | A | A | A | A | A | A | W | W | A | W | W | W | W | A | A | A | W | A | W | W | A | W | A | W | A | W | A | W | W | A | A | W | A | A | A | A | A | W | A | A | W | W | W | W | W | W | A | W | A | A | A | W | W | W | A | W | W | A | A | W | A | W | W | A | W | A | A | W | A | A | W |
| Chrl | 1.7 | A | W | A | NA | W | W | W | W | W | A | A | A | W | A | NA | W | NA | W | W | NA | W | A | A | A | A | A | A | W | W | A | W | W | W | W | A | A | A | W | A | W | W | A | W | A | NA | A | W | A | W | NA | A | A | W | A | A | A | A | A | W | A | A | W | W | W | W | W | W | A | W | A | A | A | W | W | W | A | W | W | A | A | W | A | W | W | A | W | A | A | W | A | A | W |
| Chrl | 1.8 | A | W | A | W | W | W | W | W | W | A | A | A | W | A | W | W | W | W | W | NA | W | A | A | A | A | A | A | NA | W | A | W | W | W | W | A | A | A | W | A | W | W | A | W | A | W | A | W | A | W | NA | A | A | W | A | A | A | A | A | W | A | A | W | W | W | W | NA | W | A | W | A | A | A | W | W | W | A | W | W | A | A | W | A | W | W | A | W | NA | A | W | A | A | W |
| Chrl | 1.9 | A | W | A | W | W | W | W | W | W | A | A | A | W | A | W | W | W | W | W | W | W | A | A | A | A | A | A | W | W | A | W | W | W | W | A | A | A | W | A | W | W | A | W | A | W | A | W | A | W | W | A | A | W | A | A | A | A | A | W | A | A | W | W | W | W | W | W | A | W | A | A | A | W | W | W | A | W | W | A | A | W | A | W | W | A | W | A | A | W | A | A | W |
| Chrl | 2 | A | W | A | W | W | W | W | W | W | A | A | A | W | A | W | W | W | W | W | W | W | A | A | A | A | A | A | W | W | A | W | W | W | W | A | A | A | W | A | W | W | A | W | A | W | A | W | A | W | W | A | A | W | A | A | A | A | A | W | A | A | W | W | W | W | W | W | A | W | A | A | A | W | W | W | A | W | W | A | A | W | A | W | W | A | W | A | A | W | A | A | W |
| Chrl | 2.1 | A | W | A | W | W | W | W | W | W | A | A | A | W | A | W | W | W | W | W | W | W | A | A | A | A | A | A | W | W | A | W | W | W | W | A | A | A | W | A | W | W | A | W | A | W | A | W | A | W | W | A | A | W | A | NA | A | A | A | W | A | A | W | W | W | W | W | W | A | W | A | A | A | W | W | W | A | W | W | A | A | W | A | W | W | A | W | A | A | W | A | A | W |
| Chrl | 2.2 | A | W | A | W | W | W | W | W | W | A | A | A | W | A | W | W | NA | W | W | W | W | A | A | A | A | A | A | W | W | A | W | W | W | NA | A | A | A | W | A | W | W | A | NA | A | W | A | W | A | W | W | A | A | W | A | A | A | A | A | W | A | A | W | W | W | W | W | W | A | W | A | A | A | W | W | W | A | W | W | A | A | W | A | W | W | A | W | A | A | W | A | A | W |
| Chrl | 2.3 | A | W | A | W | W | W | W | W | W | A | A | A | W | A | W | W | W | W | W | NA | W | A | A | A | A | A | A | W | W | A | W | W | W | W | A | A | A | W | A | W | W | A | A | A | W | A | W | A | W | W | A | A | W | A | A | A | A | A | W | A | A | W | W | W | W | W | W | A | W | A | A | A | W | W | W | A | W | W | A | A | W | A | W | W | A | W | A | A | W | A | A | W |
| Chrl | 2.4 | A | W | A | W | W | W | W | W | W | A | A | A | W | A | W | W | W | W | W | W | NA | A | A | A | A | A | A | W | W | A | W | W | W | W | A | A | A | W | A | W | W | A | A | A | W | A | W | A | W | NA | A | A | W | A | A | A | A | A | W | A | A | W | W | W | W | W | W | A | W | A | A | A | W | W | W | A | W | W | A | A | W | A | W | W | A | W | A | A | W | A | A | W |
| Chrl | 2.5 | A | W | A | W | W | W | W | W | W | A | A | A | W | A | W | W | W | W | W | W | W | A | A | A | A | A | A | W | W | A | W | W | W | W | A | A | A | W | A | W | W | A | A | A | W | A | W | A | W | W | A | A | W | A | A | A | A | A | W | A | A | W | W | W | W | W | W | A | W | A | A | A | W | W | W | A | W | W | A | A | W | A | W | W | A | W | A | A | W | A | A | W |
| Chrl | 2.6 | A | W | A | W | W | W | W | W | W | NA | A | NA | W | NA | W | W | W | W | W | W | W | A | A | A | A | A | NA | W | NA | A | W | W | W | W | A | A | NA | W | NA | W | W | A | A | A | W | A | W | A | W | W | A | NA | W | A | A | NA | NA | A | W | A | A | W | W | W | W | W | W | A | W | A | NA | A | W | W | W | NA | W | W | A | A | W | A | W | W | A | W | A | A | W | A | A | W |
| Chrl | 2.7 | A | W | A | W | W | W | W | W | W | A | A | A | W | A | W | W | NA | W | W | W | W | A | A | A | A | A | A | W | W | A | W | W | W | W | A | A | A | W | A | W | W | A | A | A | W | A | W | A | W | W | A | A | W | A | A | A | A | A | W | A | A | W | W | W | W | W | NA | A | W | A | A | A | W | W | W | A | W | W | A | A | W | A | W | W | A | W | A | A | W | A | NA | W |
| Chrl | 2.8 | A | W | A | W | W | W | W | W | W | A | A | A | W | A | NA | NA | A | W | W | W | W | A | A | A | A | A | A | W | W | A | W | W | W | W | A | A | A | W | A | W | NA | A | A | A | W | A | W | A | W | W | A | A | W | A | A | A | A | A | W | A | A | W | W | W | W | W | W | A | W | A | A | A | W | W | W | A | W | W | A | A | W | A | W | W | A | W | A | A | W | A | W | W |
| Chrl | 2.9 | A | W | A | W | W | W | W | W | W | A | A | A | W | A | A | W | A | W | W | A | W | A | A | A | A | A | A | W | W | A | W | W | W | W | A | A | A | W | A | W | A | A | A | A | W | A | W | A | W | W | A | A | W | A | A | A | A | A | W | A | A | W | W | W | W | W | W | A | W | A | A | A | W | W | W | A | W | W | A | A | W | A | W | W | A | W | A | A | W | A | W | W |
| Chrl | 3 | A | W | A | W | W | W | W | W | W | A | A | A | W | A | A | W | A | W | W | A | W | A | A | A | A | A | A | W | W | A | W | W | W | W | A | A | A | W | A | W | A | A | A | A | W | A | W | A | W | W | A | A | W | A | A | A | A | A | W | A | A | W | W | W | W | W | W | A | W | A | A | A | W | W | W | A | W | W | A | A | W | A | W | W | A | W | A | A | W | A | W | W |
| Chrl | 3.1 | A | W | A | W | W | W | W | W | W | A | A | NA | W | A | A | W | A | W | W | A | W | A | A | A | A | A | A | W | W | A | W | W | W | W | A | A | A | W | A | W | A | A | A | A | W | A | W | A | W | W | A | A | W | A | A | A | A | A | W | A | A | W | W | W | W | W | W | A | W | A | A | A | W | W | W | A | W | W | A | A | W | A | W | W | A | W | A | A | W | A | W | W |
| Chrl | 3.2 | A | W | A | W | W | W | W | W | W | A | A | A | W | NA | A | W | NA | W | W | A | W | A | A | A | A | A | A | W | W | A | W | W | W | W | A | A | A | W | A | W | A | A | A | A | W | A | W | A | W | W | A | A | W | A | A | A | A | A | W | A | NA | W | W | W | W | W | W | A | W | A | NA | A | W | W | W | A | W | W | A | A | W | A | W | W | A | W | A | A | W | A | W | W |
| Chrl | 3.3 | A | W | A | W | W | W | W | W | NA | A | A | A | W | A | NA | W | NA | W | W | A | W | A | A | A | A | A | A | W | W | A | W | W | W | W | A | A | NA | W | A | W | A | A | A | A | W | A | W | A | W | NA | A | A | W | A | A | A | A | A | W | A | A | W | W | W | NA | W | W | A | W | A | A | A | W | W | W | NA | W | W | A | A | W | A | W | W | A | W | A | A | W | A | W | W |
| Chrl | 3.4 | A | W | A | W | W | W | W | W | A | A | A | A | W | A | A | W | A | W | W | A | W | A | A | A | A | A | A | W | W | A | W | W | W | W | A | A | W | W | A | W | A | A | A | A | W | A | W | A | W | W | A | A | W | A | A | A | A | A | W | A | A | W | W | W | W | W | W | A | W | A | A | A | W | W | W | A | W | W | A | A | W | A | W | W | A | W | A | A | W | A | W | W |
| Chrl | 3.5 | A | W | A | W | W | W | W | W | A | A | A | A | W | A | A | W | A | W | W | A | W | A | A | A | A | A | A | W | W | A | W | W | W | W | A | A | W | W | A | W | A | A | A | A | W | A | W | A | W | W | A | A | W | A | A | A | A | A | W | A | A | W | W | W | W | W | W | A | W | A | A | A | W | W | W | A | W | W | A | A | W | A | W | W | A | W | A | A | W | A | W | W |
| Chrl | 3.6 | A | W | A | W | W | NA | W | W | A | A | A | A | W | A | A | W | A | W | W | A | W | A | A | A | NA | A | A | W | W | A | W | W | W | W | A | A | W | W | A | W | A | A | NA | A | W | A | W | A | W | W | A | A | NA | A | A | A | A | A | W | A | A | W | W | W | W | W | W | A | W | A | A | A | W | W | W | A | W | W | A | A | W | A | W | W | A | W | A | A | W | A | W | W |
| Chrl | 3.7 | A | W | A | W | W | A | W | W | A | A | A | A | W | A | A | W | A | W | W | A | W | A | A | A | A | A | A | W | W | A | W | W | W | W | A | A | W | W | A | W | A | A | A | A | W | A | W | A | W | W | A | A | W | A | NA | A | A | A | W | A | A | W | W | W | W | W | W | A | W | A | A | A | W | W | W | A | W | W | A | A | W | A | W | W | A | W | A | A | W | A | W | W |
| Chrl | 3.8 | A | W | A | W | W | A | W | W | A | A | A | A | W | A | A | W | A | W | W | A | W | A | A | A | A | A | A | W | W | A | W | W | W | W | A | A | W | W | A | W | A | A | A | A | W | A | W | A | W | NA | A | A | W | A | A | A | A | A | W | A | A | W | W | W | W | W | W | A | W | A | A | A | W | W | W | A | W | W | A | A | W | A | W | W | A | W | A | A | W | A | W | W |
| Chrl | 3.9 | A | W | A | W | W | A | W | W | A | A | A | A | W | A | A | W | A | W | W | A | W | A | A | A | A | A | A | W | W | A | W | W | W | W | A | A | W | W | A | W | A | A | A | A | W | A | W | A | W | W | A | A | W | A | A | A | A | A | W | A | A | W | W | W | W | W | W | A | W | A | A | A | W | W | W | A | W | W | A | A | W | A | W | W | A | W | A | A | W | A | W | W |
| Chrl | 4 | A | W | A | NA | W | A | NA | NA | A | A | A | A | W | A | NA | W | A | W | W | A | W | A | A | A | A | A | A | W | W | A | NA | NA | W | W | A | A | W | NA | A | W | A | A | A | A | NA | A | W | A | W | W | A | NA | W | A | NA | A | A | A | W | A | NA | W | W | NA | NA | W | NA | NA | W | A | A | A | W | W | W | A | W | W | A | A | W | A | NA | W | A | W | NA | A | NA | A | W | W |
| Chrl | 4.1 | A | W | A | W | W | A | W | W | A | A | A | A | W | A | A | W | A | W | W | A | W | A | A | A | A | A | A | W | W | A | W | A | W | W | A | A | W | W | A | W | A | A | A | A | W | A | W | A | W | W | A | A | W | A | A | A | A | A | W | A | A | W | W | W | W | W | W | A | W | A | A | A | W | W | W | A | W | W | A | A | W | A | W | W | A | W | W | A | W | A | W | W |
| Chrl | 4.2 | A | W | A | W | W | NA | W | W | A | A | A | A | W | A | A | W | A | W | W | A | NA | A | A | A | A | A | A | W | W | A | W | A | W | W | A | A | W | W | A | W | A | A | A | A | W | A | W | A | W | W | A | NA | W | A | A | A | A | A | W | A | A | W | W | W | W | W | W | A | W | A | A | A | W | W | W | A | W | W | A | A | W | A | W | W | A | W | W | A | W | A | W | W |
| Chrl | 4.3 | A | W | A | W | W | A | W | W | A | A | A | A | W | A | A | W | A | W | W | A | W | A | A | A | A | A | A | W | W | A | W | A | W | NA | A | A | W | W | A | W | A | A | A | A | W | A | W | A | W | W | A | A | W | A | A | A | A | A | W | A | A | W | W | W | W | W | W | A | W | A | A | A | W | W | W | A | W | W | A | A | W | A | W | W | A | W | NA | A | W | A | W | W |
| Chrl | 4.4 | A | W | A | W | W | A | W | W | A | A | A | A | W | A | A | W | A | W | W | A | W | A | A | A | A | A | A | W | W | A | W | A | W | W | A | A | W | W | A | W | A | A | A | A | W | A | W | A | W | W | A | A | W | A | A | A | A | A | W | A | A | W | W | W | NA | W | W | A | W | A | A | A | W | W | W | A | W | W | A | A | W | A | W | W | A | W | W | A | W | A | W | W |
| Chrl | 4.5 | A | W | A | NA | W | A | W | W | A | A | A | A | W | A | A | W | A | W | W | NA | W | A | A | A | A | A | A | W | W | A | W | A | W | W | A | A | W | W | A | W | A | A | A | A | W | A | W | A | W | W | A | A | W | A | A | A | A | A | W | A | A | W | W | W | NA | W | NA | A | W | A | A | A | W | W | W | A | NA | W | A | A | W | A | W | W | A | W | NA | A | W | A | W | W |
| Chrl | 4.6 | A | W | A | W | W | A | W | W | A | A | A | A | W | A | A | W | A | W | W | A | W | A | NA | A | A | A | A | W | W | A | W | A | W | W | A | A | W | W | A | W | A | A | A | A | W | A | W | A | W | W | A | A | W | A | A | A | A | A | W | A | A | W | W | W | W | W | W | A | W | A | A | A | W | W | W | A | W | W | A | A | W | A | W | W | A | W | W | A | W | A | W | W |
| Chrl | 4.7 | A | W | A | W | W | A | W | W | A | A | A | A | W | A | A | W | A | W | W | A | W | A | A | A | A | A | A | NA | W | A | W | A | W | W | A | A | W | W | A | W | A | A | A | A | W | A | W | A | W | W | A | A | W | A | A | A | A | A | W | A | A | W | W | W | NA | NA | NA | A | W | NA | A | A | W | W | W | A | W | W | A | A | W | A | W | NA | A | W | NA | A | W | A | W | W |
| Chrl | 4.8 | A | W | A | NA | W | A | W | W | A | A | A | A | W | A | A | W | A | W | W | A | W | A | NA | A | A | A | A | NA | W | A | W | A | W | NA | A | A | W | W | A | W | A | W | A | A | W | A | W | A | W | W | A | A | W | A | A | A | A | A | W | A | A | W | W | W | NA | W | NA | A | W | W | A | A | W | W | W | A | W | W | A | A | W | A | W | W | A | W | NA | A | W | A | W | W |
| Chrl | 4.9 | A | W | A | W | W | A | W | W | A | A | A | A | W | A | A | W | A | W | W | A | W | A | A | A | A | A | A | W | W | A | W | A | W | W | A | A | W | W | A | W | A | W | A | A | W | A | W | A | W | W | A | A | W | A | A | A | A | A | W | A | A | W | W | W | A | NA | NA | A | W | W | A | A | W | W | W | A | W | W | A | A | W | A | W | W | A | W | W | A | W | A | W | W |
| Chrl | 5 | A | W | A | W | W | A | W | W | A | A | A | A | W | A | A | W | A | W | W | A | W | A | A | A | A | A | A | W | W | A | W | A | W | W | A | A | W | W | A | W | A | W | A | A | W | A | W | A | W | W | A | A | W | A | A | A | A | A | W | A | A | W | W | W | A | W | W | A | W | W | A | A | W | W | W | A | W | W | A | A | W | A | W | W | A | W | W | A | W | A | W | W |
| Chrl | 5.1 | A | W | A | W | W | A | W | W | A | A | A | A | W | A | A | W | A | W | W | A | W | A | NA | A | A | A | A | W | W | A | W | A | W | W | A | A | W | W | A | W | A | W | A | A | W | A | W | A | W | W | A | A | W | A | A | A | A | A | W | A | A | W | W | W | A | NA | W | A | W | W | A | A | W | W | W | A | W | W | A | A | W | A | W | NA | A | W | W | A | W | A | W | W |
| Chrl | 5.2 | A | W | A | W | W | A | W | W | A | A | A | A | W | A | A | W | A | W | W | A | W | A | W | A | A | A | A | W | W | A | W | A | W | W | A | A | W | W | A | W | A | W | A | A | W | A | W | A | W | W | A | A | W | A | A | A | A | A | W | A | A | W | W | W | A | W | W | A | W | W | A | A | W | W | W | A | W | W | A | A | W | A | W | W | A | W | W | A | W | A | W | W |
| Chrl | 5.3 | A | W | A | W | W | A | W | W | A | A | A | A | W | A | A | W | A | W | W | A | W | A | NA | A | A | A | A | W | W | A | W | A | W | W | A | A | W | W | A | W | A | W | A | A | W | A | W | A | W | W | A | A | W | A | A | A | A | A | W | A | A | W | W | W | A | W | W | A | W | W | A | A | W | W | W | A | W | W | A | A | W | A | W | W | A | W | W | A | W | A | W | W |
| Chrl | 5.4 | A | W | A | W | W | A | W | W | A | A | A | A | W | A | A | W | A | W | W | A | W | A | W | A | A | A | A | W | W | A | W | A | W | W | A | A | W | W | A | W | A | W | A | A | W | A | W | A | W | W | A | A | W | A | A | A | A | A | W | A | A | W | W | W | A | W | W | A | W | W | A | A | W | W | W | A | W | W | A | A | W | A | W | W | A | W | W | A | W | A | W | W |
| Chrl | 5.5 | A | W | A | W | W | A | W | W | A | A | A | A | W | A | A | W | A | W | W | A | W | A | W | A | A | A | A | W | W | A | W | A | W | W | A | A | W | W | A | W | A | W | A | A | W | A | W | A | W | W | A | A | W | A | A | A | A | A | W | A | A | W | W | W | NA | W | W | A | W | W | NA | A | W | W | W | A | W | W | A | A | W | A | W | W | A | W | W | A | W | A | W | W |
| Chrl | 5.6 | A | W | A | W | W | A | W | W | A | A | A | A | W | A | A | W | A | W | W | NA | W | A | W | A | A | A | A | W | W | A | W | A | W | W | A | A | W | W | A | W | A | W | A | A | NA | A | W | A | W | W | A | A | W | A | A | A | A | A | W | A | A | W | W | W | A | W | W | A | W | W | A | A | W | W | W | A | W | W | A | A | W | A | W | W | A | W | NA | A | W | A | W | W |
| Chrl | 5.7 | A | W | A | W | W | A | W | W | A | A | A | A | W | A | A | W | A | W | W | NA | W | A | W | A | A | A | A | W | W | A | W | A | W | W | A | A | W | W | A | W | A | W | A | A | W | A | W | A | W | W | A | A | W | A | A | A | A | A | W | A | A | W | W | W | A | W | W | A | W | W | A | A | W | W | W | A | W | W | A | A | W | A | W | W | A | W | W | A | W | A | W | W |
| Chrl | 5.8 | A | W | A | W | W | A | W | W | A | A | A | A | W | A | A | W | A | W | W | A | W | A | W | A | A | A | A | W | W | A | W | A | W | W | A | A | W | W | A | W | A | W | A | A | W | A | W | A | W | W | A | A | W | A | A | A | A | A | W | A | A | W | W | W | A | W | W | A | W | W | A | A | W | W | W | A | W | W | A | A | W | A | W | W | A | W | W | A | W | A | W | W |
| Chrl | 5.9 | A | W | A | W | W | A | W | W | A | A | A | A | W | A | A | W | A | W | W | NA | W | A | NA | A | A | A | A | NA | W | A | W | A | W | NA | A | A | W | W | A | W | A | W | A | A | W | A | W | A | W | W | A | A | W | A | NA | A | A | A | W | A | A | W | W | W | NA | W | NA | A | W | W | A | A | W | W | W | A | W | W | A | A | W | A | W | W | A | W | W | A | W | A | W | NA |
| Chrl | 6 | A | W | A | W | W | A | W | W | A | A | A | A | W | A | A | W | A | W | W | A | W | A | W | A | A | A | A | W | W | A | W | A | W | NA | A | A | W | W | A | W | A | W | A | A | W | A | W | A | W | W | A | A | W | A | A | A | A | A | W | A | A | W | W | W | A | W | W | A | W | W | A | A | W | W | W | A | W | W | A | A | W | A | W | W | A | W | NA | A | W | A | W | W |
| Chrl | 6.1 | A | W | A | W | W | A | W | W | A | A | A | A | W | A | A | W | A | W | W | A | W | A | W | A | A | A | A | W | W | A | W | A | W | NA | A | A | W | W | A | W | A | W | A | A | W | A | W | A | W | W | A | A | W | A | A | A | A | A | W | A | A | W | W | W | NA | NA | NA | A | W | W | A | A | W | W | W | A | W | W | A | A | W | A | W | W | A | W | W | A | W | A | W | W |
| Chrl | 6.2 | A | W | A | W | W | A | W | NA | A | A | A | A | W | A | A | W | A | W | W | A | W | A | W | A | A | A | A | NA | W | A | W | A | W | W | A | A | W | W | A | W | A | W | A | A | W | A | W | A | W | W | A | A | W | A | A | A | A | A | W | A | A | W | W | W | A | W | W | A | W | W | A | A | W | W | W | A | W | W | A | A | W | A | W | W | A | W | NA | A | W | A | W | W |
| Chrl | 6.3 | A | W | A | W | W | A | W | W | A | A | A | A | W | A | A | W | A | W | W | A | W | A | W | A | A | A | A | W | W | A | W | A | W | NA | A | A | W | W | A | W | A | W | A | A | W | A | W | A | W | W | A | A | W | A | A | A | A | A | W | A | A | W | W | W | A | W | W | A | W | W | A | A | W | W | W | A | W | W | A | A | W | A | W | W | A | W | W | A | W | A | W | W |
| Chrl | 6.4 | A | W | A | W | W | A | W | W | A | A | A | A | W | A | A | W | A | W | W | A | W | A | W | A | A | A | A | NA | W | A | W | A | W | W | A | A | W | W | A | W | A | W | A | A | W | A | W | A | W | W | A | A | W | A | A | A | A | A | W | A | A | W | W | W | A | W | NA | A | W | W | A | A | W | W | W | A | NA | W | A | A | W | A | W | W | A | W | W | A | W | A | W | NA |
| Chrl | 6.5 | A | W | A | W | W | A | W | W | NA | A | NA | A | W | A | A | W | NA | W | W | NA | W | A | W | A | NA | NA | NA | W | W | A | W | A | W | W | A | A | W | W | A | W | A | W | A | A | W | NA | W | A | W | W | A | NA | W | A | NA | A | A | A | W | A | A | W | W | W | NA | W | W | A | W | W | NA | A | W | W | W | A | W | W | A | A | W | A | W | W | A | W | W | NA | W | A | W | W |
| Chrl | 6.7 | A | W | A | W | W | A | W | W | A | A | A | A | W | A | A | W | A | W | W | A | W | A | W | A | A | A | A | W | W | A | W | A | W | W | A | A | W | W | A | W | A | W | A | A | W | A | W | A | W | W | A | A | W | A | A | A | A | A | W | A | A | W | W | W | A | W | W | A | W | W | A | NA | W | W | W | A | W | W | A | A | W | A | W | W | A | W | W | A | W | A | W | W |
| Chrl | 6.8 | A | W | A | W | W | A | W | W | A | A | A | A | W | A | A | W | A | W | W | A | W | A | W | A | A | A | A | W | W | A | W | A | W | NA | A | A | W | W | A | W | A | W | A | A | W | A | W | A | W | W | A | A | W | A | A | A | A | A | W | A | A | W | W | W | A | NA | NA | A | W | W | A | A | W | W | W | A | W | W | A | A | W | A | W | W | A | W | W | A | W | A | W | W |
| Chrl | 6.9 | A | W | A | W | W | A | W | W | A | A | A | A | W | A | A | W | A | W | W | A | W | A | W | A | A | A | A | W | W | A | W | A | W | W | A | A | W | W | A | W | A | NA | A | A | W | A | W | A | W | NA | A | A | W | A | A | A | A | A | W | A | A | W | W | W | A | W | W | A | W | W | A | A | W | W | W | A | W | W | A | A | W | A | W | W | A | W | W | A | W | A | W | W |
| Chrl | 7 | A | W | A | W | W | A | W | NA | A | A | A | A | W | A | NA | W | NA | W | W | A | W | A | W | A | A | A | A | NA | W | A | NA | A | W | NA | NA | A | W | NA | A | W | A | W | A | A | W | A | W | A | W | W | A | A | W | A | NA | A | A | A | W | A | A | W | W | NA | NA | W | W | A | W | W | NA | A | W | W | W | NA | NA | W | A | A | W | A | NA | W | A | W | NA | NA | NA | NA | W | NA |
| Chrl | 7.1 | A | W | A | W | W | A | W | NA | A | A | A | A | W | A | A | W | A | W | W | A | W | A | W | A | A | A | A | W | W | A | NA | A | W | NA | A | A | W | W | A | W | A | W | A | A | W | A | W | A | W | W | A | A | W | A | A | A | A | A | W | A | A | W | W | W | NA | NA | W | A | W | W | A | A | W | W | W | A | W | W | A | A | W | A | NA | W | A | W | W | A | NA | A | W | NA |
| Chrl | 7.2 | A | W | A | W | W | A | W | W | A | A | A | A | W | A | A | W | A | W | W | A | W | A | NA | A | A | A | A | W | W | A | W | A | W | NA | A | A | W | W | A | W | A | W | A | A | W | A | W | A | W | W | A | A | W | A | A | A | A | A | W | A | A | W | W | W | A | W | W | A | W | W | A | A | W | W | W | A | W | W | A | A | W | A | W | W | A | W | W | A | W | A | W | W |
| Chrl | 7.3 | A | W | A | W | W | A | W | W | A | A | A | A | W | A | A | W | A | W | W | A | W | A | W | A | A | A | A | W | W | A | W | A | W | NA | A | A | W | W | A | W | A | W | A | A | W | A | W | A | W | W | A | A | W | A | A | A | A | A | W | A | A | W | W | W | A | W | W | A | W | W | A | A | W | W | W | A | W | W | A | A | W | A | W | W | A | W | NA | A | W | A | W | W |
| Chrl | 7.4 | A | W | A | W | W | A | W | W | A | A | A | A | W | A | NA | W | A | W | W | A | W | A | W | A | A | A | A | W | W | A | W | A | W | W | A | A | W | W | A | W | A | W | A | A | W | A | W | A | W | W | A | A | W | A | A | A | A | A | W | A | A | W | W | W | A | W | W | A | W | W | A | A | W | W | W | A | W | W | A | A | W | A | W | W | A | W | W | A | W | A | W | W |
| Chrl | 7.5 | A | W | A | W | W | A | W | W | A | A | A | A | W | A | A | W | A | W | W | A | W | A | W | A | A | A | A | NA | W | A | W | A | W | W | A | A | W | W | A | W | A | W | A | A | W | A | W | A | W | W | A | A | W | A | A | A | A | A | W | A | A | W | W | W | A | NA | NA | A | W | W | A | A | W | W | W | A | W | W | A | A | W | A | W | W | A | W | W | A | W | A | W | W |
| Chrl | 7.6 | A | W | A | W | W | A | W | W | A | A | A | A | W | A | A | W | NA | W | W | A | W | A | W | A | A | A | A | W | W | A | W | NA | W | W | A | A | W | W | A | W | NA | W | A | A | W | A | W | A | W | W | A | A | W | A | NA | NA | A | A | W | A | NA | W | W | W | A | W | W | A | W | W | A | NA | W | W | W | A | W | W | A | NA | W | A | W | W | NA | W | W | A | W | A | W | W |
| Chrl | 7.7 | A | W | A | W | W | A | W | W | A | A | A | A | W | A | A | W | A | W | W | A | W | A | W | A | A | A | A | W | W | A | W | A | W | W | A | A | W | W | A | W | A | W | A | A | W | A | W | A | W | W | A | A | W | A | A | A | A | A | W | A | A | W | W | W | A | W | W | A | W | W | A | A | W | W | W | A | W | W | A | A | W | A | W | W | A | W | W | A | W | A | W | W |
| Chrl | 7.8 | A | W | A | W | W | A | W | W | A | A | A | A | W | A | A | W | A | W | W | A | W | A | W | A | A | A | A | NA | W | A | W | A | W | NA | A | A | W | W | A | W | A | W | A | A | W | A | W | A | W | W | A | A | W | A | A | A | A | A | W | A | A | W | W | W | NA | W | W | A | W | W | A | A | W | W | W | A | W | W | A | A | W | A | W | W | A | W | W | A | W | A | W | W |
| Chrl | 7.9 | A | W | A | W | W | A | W | W | A | A | A | A | W | A | A | W | A | W | W | A | W | A | W | A | A | A | A | NA | W | A | W | A | W | NA | A | A | W | W | A | W | A | W | A | A | W | A | W | A | W | W | A | A | W | A | A | A | A | A | W | A | A | W | W | W | A | W | NA | A | W | W | A | A | W | W | W | A | W | W | A | A | W | A | W | W | A | W | W | A | W | A | W | W |
| Chrl | 8 | A | W | A | W | W | A | W | W | A | A | A | A | W | A | A | W | A | W | W | A | W | A | W | A | A | A | A | W | W | A | W | A | W | NA | A | A | W | W | A | W | A | W | A | A | W | A | W | A | W | NA | A | A | W | A | A | A | A | A | W | A | A | W | W | W | A | W | W | A | W | W | A | A | W | W | W | A | W | W | A | A | W | A | W | W | A | W | NA | A | W | A | W | W |
| Chrl | 8.1 | A | W | A | W | W | A | W | W | A | A | A | A | W | A | A | W | A | W | W | A | W | A | NA | A | A | A | A | W | W | A | W | A | W | NA | A | A | W | W | A | W | A | W | A | A | W | A | W | A | W | W | A | A | W | A | A | A | A | A | W | A | A | W | W | W | A | W | W | A | W | W | A | A | W | W | W | A | W | W | A | A | W | A | W | W | A | W | W | A | W | A | W | W |
| Chrl | 8.2 | A | W | A | W | W | A | W | NA | A | A | A | A | W | A | A | W | A | W | W | A | W | A | W | A | A | A | A | W | W | A | W | A | W | W | A | A | W | W | A | W | A | W | A | A | W | A | W | A | W | NA | A | A | NA | A | A | A | A | A | W | A | A | W | W | W | A | W | W | A | W | W | A | A | W | W | W | A | W | W | A | A | W | A | W | W | A | W | W | A | W | A | W | W |
| Chrl | 8.3 | A | W | A | W | W | A | W | W | A | A | A | A | W | A | A | W | A | W | W | A | W | A | W | A | A | A | A | W | W | A | W | A | W | W | A | A | W | W | A | W | A | W | A | A | W | A | W | A | W | A | A | A | A | A | A | A | A | A | W | A | A | W | W | W | A | W | W | A | W | W | A | A | W | W | W | A | W | W | A | A | W | A | W | W | A | W | W | A | W | A | W | W |
| Chrl | 8.4 | A | W | A | W | W | A | W | W | A | A | A | A | W | A | A | W | A | W | W | A | W | A | W | A | A | A | A | W | W | A | W | A | W | W | A | A | W | W | A | W | A | W | A | A | W | A | W | A | W | A | A | A | A | A | A | A | A | A | W | A | A | W | W | W | A | W | W | A | W | W | A | A | W | W | W | A | W | W | A | A | W | A | W | W | A | W | W | A | W | A | W | W |
| Chrl | 8.6 | A | W | A | W | W | A | W | W | A | A | A | A | W | A | A | W | A | W | W | A | W | A | W | A | A | A | A | W | W | A | W | A | W | W | A | A | W | W | A | W | A | W | A | A | W | A | W | A | W | A | A | A | A | A | A | A | A | A | W | A | A | W | W | W | A | W | W | A | W | W | A | A | W | W | W | A | W | W | A | A | W | A | W | W | A | W | W | A | W | A | W | W |
| Chrl | 8.7 | A | W | A | W | W | A | NA | W | A | A | A | A | W | A | A | W | A | W | W | A | W | A | W | A | A | A | A | NA | W | A | NA | A | W | W | A | A | W | W | A | W | A | W | A | A | W | A | W | A | W | A | A | A | A | A | A | A | A | A | W | A | A | W | W | W | A | NA | NA | A | W | W | A | A | W | W | W | A | W | W | A | A | W | A | W | W | A | W | W | A | W | A | W | NA |
| Chrl | 8.8 | A | W | A | W | W | A | A | W | A | A | A | A | W | A | A | W | A | W | W | A | W | A | W | A | A | A | A | NA | W | A | W | A | W | W | A | A | W | W | A | W | A | W | A | A | W | A | W | A | W | A | A | A | A | A | A | A | A | A | W | A | A | W | W | W | A | W | NA | A | W | W | A | A | W | W | W | A | W | W | A | A | W | A | W | W | A | W | W | A | W | A | W | W |
| Chrl | 8.9 | A | W | A | W | W | A | A | W | A | A | A | A | W | A | A | W | A | W | W | A | W | A | W | A | A | A | A | NA | W | A | NA | A | W | NA | A | A | W | NA | A | W | A | W | A | A | W | A | W | A | W | A | A | A | A | A | A | A | A | A | W | A | A | W | W | NA | A | NA | NA | A | W | W | A | A | W | W | W | A | NA | W | A | A | W | A | NA | W | A | W | W | A | W | A | W | NA |
| Chrl | 9 | A | W | A | W | W | A | A | W | A | A | A | A | W | A | A | W | A | W | W | A | W | A | W | A | A | A | A | W | W | A | W | A | W | W | A | A | W | W | A | W | A | W | A | A | W | A | W | A | W | A | A | A | A | A | A | A | A | A | W | A | A | W | W | W | A | W | W | A | W | W | A | A | W | W | W | A | W | W | A | A | W | A | W | W | A | W | W | A | W | A | W | W |

| | | | | | | | | | | | | | | | | | | | | | | | | | | | | | | | | | | | | | | | | | | | | | | | | | | | | | | | | | | | | | | | | | | | | | | | | | | | | | | | | | | | | | | | | | | | | | |
|---|---|---|---|---|---|---|---|---|---|---|---|---|---|---|---|---|---|---|---|---|---|---|---|---|---|---|---|---|---|---|---|---|---|---|---|---|---|---|---|---|---|---|---|---|---|---|---|---|---|---|---|---|---|---|---|---|---|---|---|---|---|---|---|---|---|---|---|---|---|---|---|---|---|---|---|---|---|---|---|---|---|---|---|---|---|---|---|---|---|---|---|---|---|
| Chrl | 9.1 | A | W | A | W | W | A | A | W | A | A | A | A | W | A | A | W | A | W | W | A | W | A | NA | A | A | A | A | W | W | A | W | A | W | W | A | A | W | W | A | W | A | W | A | A | W | A | W | A | W | A | A | A | A | A | A | A | A | A | W | A | A | W | W | W | A | W | W | A | W | W | A | A | W | W | W | A | W | W | A | A | W | A | W | W | A | W | W | A | W | A | W | W |
| Chrl | 9.2 | A | NA | A | W | W | A | A | W | A | A | A | A | W | A | A | W | A | W | W | A | W | A | W | A | A | A | A | W | W | A | W | A | W | NA | A | A | W | W | A | W | A | W | A | A | W | A | W | A | W | A | A | A | A | A | A | A | A | A | W | A | A | W | W | W | A | W | W | A | W | W | A | A | W | W | W | A | W | W | A | A | W | A | W | W | A | W | W | A | W | A | W | W |
| Chrl | 9.3 | A | A | A | W | W | A | A | W | A | A | A | A | W | A | A | W | A | W | W | A | W | A | W | A | A | A | A | W | W | A | W | A | W | W | A | A | W | W | A | W | A | W | A | A | W | A | W | A | W | A | A | A | A | A | A | A | A | A | W | A | A | W | W | W | A | W | W | A | W | W | A | A | W | W | W | A | W | W | A | A | W | A | W | W | A | W | W | A | W | A | W | W |
| Chrl | 9.4 | A | A | A | W | W | A | A | W | A | A | A | A | W | A | A | W | A | W | W | NA | W | A | W | A | A | A | A | W | W | A | W | A | W | W | A | A | W | W | A | W | A | W | A | A | W | A | W | A | W | A | A | A | A | A | NA | A | A | A | W | A | A | W | W | W | NA | W | W | A | W | W | A | A | W | W | W | NA | W | W | A | A | W | A | W | W | A | W | W | A | W | A | W | W |
| Chrl | 9.5 | A | A | A | W | W | A | A | W | A | A | A | A | W | NA | A | W | A | W | W | A | W | A | W | A | A | A | A | W | W | A | W | A | W | NA | A | A | W | W | A | W | A | W | A | A | W | A | W | A | W | A | A | A | A | A | A | A | A | A | W | A | A | W | W | W | A | W | W | A | W | W | A | A | W | W | W | A | W | W | A | A | W | A | W | W | A | W | W | A | W | A | W | W |
| Chrl | 9.6 | A | A | A | W | W | A | A | W | A | A | A | A | W | A | A | W | A | W | W | A | W | A | W | A | A | A | A | NA | W | A | W | A | W | W | A | A | W | W | A | W | A | W | A | A | W | A | W | A | W | A | A | A | A | A | A | A | A | A | W | A | A | W | W | W | A | W | W | A | W | W | A | A | W | W | W | A | W | W | A | A | W | A | W | W | A | W | W | A | W | A | W | W |
| Chrl | 9.7 | A | A | A | W | W | A | A | W | A | A | A | A | W | A | A | W | A | W | W | A | W | A | W | A | A | A | A | W | W | A | W | A | W | W | A | A | W | W | A | W | A | W | A | A | W | A | W | A | W | A | A | A | A | A | A | A | A | A | W | A | A | W | W | W | A | W | W | A | W | W | A | A | W | W | W | A | W | W | A | A | W | A | W | W | A | W | W | A | W | A | W | W |
| Chrl | 9.8 | A | A | A | W | W | A | A | W | A | A | A | A | W | A | A | W | A | W | W | A | W | A | W | A | A | A | A | W | W | A | W | A | W | W | A | A | W | W | A | W | A | W | A | A | W | A | W | A | W | A | A | A | A | A | A | A | A | A | W | A | A | W | W | W | A | W | W | A | W | W | A | A | W | W | W | A | W | W | A | A | W | A | W | W | A | W | W | A | W | A | W | W |
| Chrl | 9.9 | A | A | A | W | W | A | A | W | A | A | A | A | W | NA | A | W | A | W | W | A | W | A | W | A | A | A | A | W | W | A | W | A | W | NA | A | A | W | W | A | W | A | W | A | A | W | A | W | A | W | A | A | A | A | A | A | A | A | A | W | A | A | W | W | W | A | W | W | A | W | W | NA | A | W | W | W | A | W | W | A | A | W | A | W | W | A | W | W | A | W | A | W | W |
| Chrl | 10 | A | A | A | W | W | A | A | W | A | A | A | A | W | A | A | W | A | W | W | A | W | A | W | A | A | A | A | W | W | A | W | A | W | W | A | A | W | W | A | W | A | W | A | A | W | A | W | A | W | NA | A | A | A | A | A | A | A | A | W | A | A | W | W | W | A | W | W | A | W | W | A | A | W | W | W | A | W | W | A | A | W | A | W | W | A | W | W | A | W | A | W | W |
| Chrl | 10.1 | A | A | A | W | W | A | A | W | A | A | A | A | W | A | A | W | A | W | W | A | W | A | W | A | A | A | A | W | W | A | W | A | W | W | A | A | W | W | A | W | A | W | A | A | W | A | W | A | W | NA | A | A | A | A | A | A | A | A | W | A | A | W | W | W | NA | W | W | A | W | W | A | A | W | W | W | A | W | W | A | A | W | A | W | W | A | W | W | A | W | A | W | W |
| Chrl | 10.2 | A | A | A | W | W | A | A | W | A | A | A | A | W | A | NA | W | A | W | W | A | W | A | W | A | A | A | A | W | W | A | W | A | W | W | A | A | W | W | A | W | A | W | A | A | W | A | W | A | W | A | A | A | A | A | A | NA | A | A | W | NA | NA | W | W | W | NA | NA | NA | A | W | W | A | A | W | W | W | NA | NA | W | A | A | W | A | NA | W | A | W | W | NA | NA | NA | W | W |
| Chrl | 10.3 | A | A | A | W | W | A | A | W | A | A | A | A | W | A | NA | W | A | W | W | NA | W | A | W | A | NA | A | A | W | W | A | W | A | W | W | NA | A | W | W | A | W | A | W | A | A | W | A | W | A | W | A | A | A | A | NA | A | A | A | A | W | A | A | W | W | W | NA | W | NA | A | W | W | A | A | W | W | W | NA | NA | W | A | A | W | A | W | W | A | W | W | NA | W | A | W | W |
| Chrl | 10.4 | A | A | A | W | W | A | A | W | A | A | A | A | W | A | A | W | A | W | W | A | W | A | NA | A | A | A | A | W | W | A | W | A | W | W | A | A | W | W | A | W | A | W | A | A | W | A | W | A | W | A | A | A | A | A | A | A | A | A | W | A | A | W | W | W | A | W | NA | A | W | W | A | A | W | W | W | A | W | W | A | A | W | A | W | W | A | W | W | A | W | A | W | W |
| Chrl | 10.5 | A | A | A | W | W | A | A | W | A | A | A | A | W | A | A | W | A | W | W | A | W | A | W | A | A | A | A | W | W | A | W | A | W | NA | A | A | W | W | A | W | A | W | A | A | W | A | W | A | W | A | A | A | A | A | A | A | A | A | W | A | A | W | W | W | A | W | W | A | W | W | A | A | W | W | W | A | W | W | A | A | W | A | W | W | A | W | NA | A | W | A | W | W |
| Chrl | 10.6 | A | A | A | W | W | A | A | W | A | A | A | A | W | A | A | W | A | W | W | A | W | A | W | A | A | A | A | W | W | A | W | A | W | NA | A | A | W | W | A | W | A | W | A | A | W | A | W | A | W | A | A | A | A | A | A | A | A | A | W | A | A | W | W | W | A | W | NA | A | W | W | A | A | W | W | W | A | W | W | A | A | W | A | W | W | A | W | W | A | W | A | W | W |
| Chrl | 10.7 | A | A | A | W | W | A | A | W | A | A | A | A | W | A | A | W | A | W | W | A | W | A | W | A | A | A | A | W | W | A | W | A | W | W | A | A | W | W | A | W | A | W | A | A | W | A | W | A | W | A | A | A | A | A | A | A | A | A | W | A | A | W | W | W | A | W | W | A | W | W | A | A | W | W | W | A | W | W | A | A | W | A | W | W | A | W | NA | A | W | A | W | W |
| Chrl | 10.8 | A | A | A | W | W | A | A | W | A | A | A | A | W | A | A | W | A | W | W | A | W | A | W | A | A | A | A | W | W | A | W | A | W | W | A | A | W | W | A | W | A | W | A | A | W | A | W | A | W | A | A | A | A | A | A | A | A | A | W | A | A | W | W | W | A | NA | W | A | W | W | A | A | W | W | W | A | W | W | A | A | W | A | W | W | A | W | W | A | W | A | W | W |
| Chrl | 10.9 | A | A | A | W | W | A | A | W | A | A | A | A | W | A | A | W | A | W | W | A | W | A | W | A | A | A | A | W | W | A | W | A | W | W | A | A | W | W | A | W | A | W | A | A | W | A | W | A | W | A | A | A | A | A | A | A | A | A | W | A | A | W | W | W | A | W | W | A | W | W | A | A | W | W | W | A | W | W | A | A | W | A | W | W | A | W | W | A | W | A | W | W |
| Chrl | 11 | A | A | A | W | W | A | A | W | A | A | A | A | W | A | A | W | A | W | W | A | W | A | NA | A | A | A | A | NA | W | A | W | A | W | NA | A | A | W | W | NA | W | A | W | A | A | W | A | W | A | W | A | A | A | A | A | A | A | A | A | W | A | A | W | W | W | NA | W | NA | A | W | W | A | A | W | W | W | NA | NA | W | A | A | W | A | NA | W | A | W | NA | A | NA | A | W | W |
| Chrl | 11.1 | A | A | A | W | W | A | A | W | A | A | A | A | W | A | A | W | A | W | W | A | W | A | W | A | A | A | A | NA | W | A | W | A | W | NA | A | A | W | W | A | W | A | W | A | A | W | A | W | A | W | A | A | A | A | A | A | A | A | A | W | A | A | W | W | W | A | W | W | A | W | W | A | A | W | W | W | A | W | W | A | A | W | A | W | W | A | W | W | A | W | A | W | W |
| Chrl | 11.2 | A | A | A | W | W | A | A | W | A | A | A | A | W | A | A | W | A | W | W | A | W | A | NA | A | A | A | A | W | W | A | W | A | W | NA | A | A | W | W | A | W | A | W | A | A | W | A | W | A | W | A | A | A | A | A | A | A | A | A | W | A | A | W | W | W | A | W | W | A | W | W | A | A | W | W | W | A | W | W | A | A | W | A | W | W | A | W | W | A | W | A | W | W |
| Chrl | 11.3 | A | A | A | W | W | A | A | W | A | A | A | A | W | A | A | W | A | W | W | A | W | A | W | A | A | A | A | NA | W | A | W | A | W | W | A | A | W | W | A | W | A | W | A | A | W | A | W | A | W | A | A | A | A | A | A | A | A | A | W | A | A | W | W | W | A | W | W | A | W | W | A | A | W | W | W | A | W | W | A | A | W | A | W | W | A | W | W | A | W | A | W | W |
| Chrl | 11.4 | A | A | A | W | W | A | A | W | A | A | A | A | W | A | A | W | A | W | W | A | W | A | W | A | A | A | A | W | W | A | W | A | W | NA | A | A | W | W | A | W | A | W | A | A | W | A | W | A | W | A | A | A | A | A | A | A | A | A | W | A | A | W | W | W | A | W | W | A | W | W | A | A | W | W | W | A | W | W | A | A | W | A | W | W | A | W | W | A | W | A | W | W |
| Chrl | 11.5 | A | A | A | W | W | A | A | W | A | A | A | A | W | A | A | W | A | W | W | A | W | A | W | A | A | A | A | NA | W | A | W | A | W | NA | A | A | W | W | A | W | A | W | A | A | W | A | W | A | W | A | A | A | A | A | A | A | A | A | W | A | A | W | W | W | A | W | W | A | W | W | A | A | W | W | W | A | W | W | A | A | W | A | W | W | A | W | W | A | W | A | W | W |
| Chrl | 11.6 | A | A | A | W | W | A | A | W | A | A | A | A | W | A | A | W | A | W | W | NA | W | A | W | A | A | A | A | W | W | A | W | A | W | W | A | A | W | W | A | W | A | W | A | A | W | A | W | A | W | A | A | A | A | A | A | A | A | A | W | A | A | W | W | W | A | W | W | A | W | W | A | A | W | W | W | A | W | W | A | A | W | A | W | W | A | W | W | A | W | A | W | W |
| Chrl | 11.7 | A | A | A | W | W | A | A | W | A | A | A | A | W | A | A | W | A | W | W | A | W | A | W | A | A | A | A | NA | W | A | W | A | W | W | A | A | W | W | A | W | A | W | A | A | W | A | W | A | W | NA | A | A | A | A | A | A | A | A | W | A | A | W | W | W | A | W | NA | A | W | W | A | A | W | W | W | A | W | W | A | A | W | A | W | W | A | W | W | A | W | A | W | NA |
| Chrl | 11.8 | A | A | A | W | W | A | A | W | A | A | A | A | W | A | A | W | A | W | W | A | W | A | NA | A | A | A | A | W | W | A | W | A | W | W | A | A | W | W | A | W | A | W | A | A | W | A | W | A | W | A | A | A | A | A | A | A | A | A | W | A | A | W | W | W | A | W | W | A | W | W | A | A | W | W | W | A | W | W | A | A | W | A | W | W | A | W | W | A | W | A | W | W |
| Chrl | 11.9 | A | A | A | W | W | A | A | W | A | A | A | A | W | A | A | W | A | W | W | A | W | A | W | A | A | A | A | W | W | A | W | A | W | NA | A | A | W | W | A | W | A | W | A | A | W | A | W | A | W | A | A | A | A | A | A | A | A | A | W | A | A | W | W | W | A | W | W | A | W | W | A | A | W | W | W | A | W | W | A | A | W | A | W | W | A | W | W | A | W | A | W | W |
| Chrl | 12 | A | A | A | W | W | A | A | W | A | A | A | A | W | A | A | W | A | W | W | A | W | A | NA | A | A | A | A | NA | W | A | W | A | W | W | A | A | W | W | A | W | A | W | A | A | W | A | W | A | W | A | A | A | A | A | A | A | A | A | W | A | A | W | W | W | A | W | NA | A | W | W | A | A | W | W | W | A | W | W | A | A | W | A | W | W | A | W | NA | A | W | A | W | W |
| Chrl | 12.1 | A | A | A | W | W | A | A | W | A | A | A | A | W | A | A | W | A | W | W | A | W | A | W | A | A | A | A | W | W | A | W | A | W | NA | A | A | W | W | A | W | A | W | A | A | W | A | W | A | W | A | A | A | A | A | A | A | A | A | W | A | A | W | W | W | A | W | W | A | W | W | A | A | W | W | W | A | W | W | A | A | W | A | W | W | A | W | NA | A | W | A | W | W |
| Chrl | 12.2 | A | A | A | W | W | A | A | W | A | A | A | A | W | A | A | W | A | W | W | A | W | A | W | A | A | A | A | W | W | A | W | A | W | NA | A | A | W | W | A | W | A | W | A | A | W | A | W | A | W | A | A | A | A | A | A | A | A | A | W | A | A | W | W | W | A | W | NA | A | W | W | A | A | W | W | W | A | W | W | A | A | W | A | W | W | A | W | W | A | W | A | W | W |
| Chrl | 12.3 | A | A | A | W | W | A | A | W | A | A | A | A | W | A | A | W | A | W | W | A | W | A | W | A | A | A | A | W | W | A | W | A | W | W | A | A | W | W | A | W | A | W | A | A | W | A | W | A | W | A | A | A | A | A | A | A | A | A | W | A | A | W | W | W | A | W | W | A | W | W | A | A | W | W | W | A | W | W | A | A | W | A | W | W | A | W | W | A | W | A | W | W |
| Chrl | 12.4 | A | A | A | W | W | A | A | W | A | A | A | A | W | A | A | W | A | W | W | A | W | A | W | A | A | A | A | W | W | A | W | A | W | W | A | A | W | W | A | W | A | W | A | A | W | A | W | A | W | A | A | A | A | A | NA | A | A | A | W | A | A | W | W | W | A | W | W | A | W | W | A | A | W | W | W | A | W | W | A | A | W | A | W | W | A | W | W | A | W | A | W | W |
| Chrl | 12.5 | A | A | A | W | W | A | A | W | A | A | A | A | W | A | A | W | A | W | W | A | W | A | W | A | A | A | A | W | W | A | W | A | W | W | A | A | W | W | A | W | A | W | A | A | W | A | W | A | W | A | A | A | A | A | A | A | A | A | W | A | A | W | W | W | A | W | W | A | W | W | A | A | W | W | W | A | W | W | A | A | W | A | W | W | A | W | W | A | W | A | W | W |
| Chrl | 12.6 | A | A | A | W | W | A | A | W | A | A | A | A | W | A | A | W | A | W | W | A | W | A | W | A | A | A | A | W | W | A | W | A | W | W | A | A | W | W | A | W | A | W | A | A | W | A | W | A | W | A | A | A | A | A | A | A | A | A | W | A | A | W | W | W | A | W | W | A | W | W | A | A | W | W | W | A | W | W | A | A | W | A | W | W | A | W | W | A | W | A | W | W |
| Chrl | 12.7 | A | A | A | W | W | A | A | W | A | A | A | A | W | A | A | W | A | W | W | A | W | A | W | A | A | A | A | W | W | A | W | A | W | W | A | A | W | W | A | W | A | W | A | A | W | A | W | A | W | A | A | A | A | A | A | A | A | A | W | A | A | W | W | W | A | W | W | A | W | W | A | A | W | W | W | A | W | W | A | A | W | A | W | W | A | W | W | A | W | A | W | W |
| Chrl | 12.8 | A | A | A | W | W | A | A | W | A | A | A | A | W | A | A | W | A | W | W | A | W | A | W | A | A | A | A | W | W | A | W | A | W | NA | A | A | W | W | A | W | A | W | A | A | W | A | W | A | W | A | A | A | A | A | A | A | A | A | W | A | A | W | W | W | A | W | W | A | W | W | A | A | W | W | W | A | W | W | A | A | W | A | W | W | A | W | W | A | W | A | W | W |
| Chrl | 12.9 | A | A | A | W | W | A | A | W | A | A | A | A | W | A | A | W | A | W | W | A | W | A | W | A | A | A | A | W | W | A | W | A | W | W | A | A | W | W | A | W | A | W | A | A | W | A | W | A | W | A | A | A | A | A | A | A | A | A | W | A | A | W | W | W | A | W | NA | A | W | W | A | A | W | W | W | A | W | W | A | A | W | A | W | W | A | W | W | A | W | A | W | W |
| Chrl | 13 | A | A | A | W | W | A | A | W | A | A | A | A | W | A | A | W | A | W | W | A | W | A | W | A | A | A | A | W | W | A | W | A | W | W | A | A | W | W | A | W | A | W | A | A | W | A | W | A | W | A | A | A | A | A | A | A | A | A | W | A | A | W | W | W | A | W | W | A | W | W | A | A | W | W | W | A | W | W | A | A | W | A | W | W | A | W | W | A | W | A | W | W |
| Chrl | 13.1 | A | A | A | W | W | A | A | W | A | A | A | A | W | A | A | W | A | W | W | A | W | A | W | A | A | A | A | W | W | A | W | A | W | NA | A | A | W | W | A | W | A | W | A | A | W | A | W | A | W | A | A | A | A | A | A | A | A | A | W | A | A | W | W | W | A | W | W | A | W | W | A | A | W | W | W | A | W | W | A | A | W | A | W | W | A | W | W | A | W | A | W | W |
| Chrl | 13.2 | A | A | A | W | W | A | A | W | A | A | A | A | W | A | A | W | A | W | W | A | W | A | W | A | A | A | A | W | W | A | W | A | W | W | A | A | W | W | A | W | A | W | A | A | W | A | W | A | W | A | A | A | A | A | NA | A | A | A | W | A | A | W | W | W | A | W | W | A | W | W | A | A | W | W | W | A | W | W | A | A | W | A | W | W | A | W | W | A | W | A | W | W |
| Chrl | 13.3 | A | A | A | W | W | A | A | W | A | A | A | A | W | A | A | W | A | W | W | A | W | A | W | A | A | A | A | W | W | A | W | A | W | W | A | A | W | W | A | W | A | W | A | A | W | A | W | A | W | A | A | A | A | A | W | A | A | A | W | A | A | W | W | W | A | W | W | A | W | W | A | A | W | W | W | A | W | W | A | A | W | A | W | W | A | W | W | A | W | A | W | W |
| Chrl | 13.4 | A | A | A | W | W | A | A | W | A | A | A | A | W | A | A | W | A | W | W | A | W | A | NA | A | A | A | A | W | W | A | W | A | W | W | A | A | W | W | A | W | A | W | A | A | W | A | W | A | W | A | A | A | A | A | W | A | A | A | W | A | A | W | W | W | A | W | NA | A | W | W | A | A | W | W | W | A | W | W | A | A | W | A | W | W | A | W | W | A | W | A | W | W |
| Chrl | 13.5 | A | A | A | W | W | A | A | W | A | A | A | A | W | A | A | W | A | W | W | A | W | A | W | A | A | A | A | NA | W | A | W | A | W | NA | A | A | W | W | A | W | A | W | A | A | W | A | W | A | W | A | A | A | A | A | W | A | A | A | W | A | A | W | W | W | NA | W | NA | A | W | W | A | A | W | W | W | A | W | W | A | A | W | A | W | W | A | W | W | A | W | A | W | W |
| Chrl | 13.6 | A | A | A | W | W | A | A | W | A | A | A | A | W | A | A | W | A | W | W | A | W | A | W | A | A | A | A | W | W | A | W | A | W | NA | A | A | W | W | A | W | A | W | A | A | W | A | W | A | W | A | A | A | A | A | W | A | A | A | W | A | A | W | W | W | A | W | W | A | W | W | A | A | W | W | W | A | W | W | A | A | W | A | W | W | A | W | W | A | W | A | W | W |
| Chrl | 13.7 | A | A | A | W | W | A | A | W | A | A | A | A | W | A | A | W | A | W | W | A | W | A | W | A | A | A | A | W | W | A | W | A | W | NA | A | A | W | W | A | W | A | W | A | A | W | A | W | A | W | A | A | A | A | A | W | A | A | A | W | A | A | W | W | W | A | W | NA | A | W | W | A | A | W | W | W | A | W | W | A | A | W | A | W | W | A | W | W | A | W | A | W | W |
| Chrl | 13.8 | A | A | A | W | W | A | A | W | A | A | A | A | W | A | A | W | A | W | W | A | W | A | W | A | A | A | A | W | W | A | W | A | W | W | A | A | W | W | A | W | A | W | A | A | W | A | W | A | W | A | A | A | A | A | W | A | A | A | W | A | A | W | W | W | A | W | W | A | W | W | A | A | W | W | W | A | W | W | A | A | W | A | W | W | A | W | W | A | W | A | W | W |
| Chrl | 13.9 | A | A | A | W | W | A | A | W | A | A | A | A | W | A | A | W | A | W | W | A | W | A | W | A | A | A | A | NA | W | A | W | A | W | NA | A | A | W | W | A | W | A | W | A | A | W | A | W | A | W | A | A | A | A | A | W | A | A | A | W | A | A | W | W | W | A | NA | W | A | W | W | A | A | W | W | W | A | W | W | A | A | W | A | W | W | A | W | W | A | W | A | W | W |
| Chrl | 14 | A | A | A | W | W | A | A | W | A | A | A | A | W | A | A | W | A | W | W | A | W | A | W | A | A | A | A | W | W | A | W | A | W | W | A | A | W | W | A | W | A | W | A | A | W | A | W | A | W | A | A | A | A | A | W | A | A | A | W | A | A | W | W | W | A | W | NA | A | W | W | A | A | W | W | W | A | W | W | A | A | W | A | W | W | A | W | W | A | W | A | W | W |
| Chrl | 14.1 | A | A | A | W | W | A | A | W | A | A | A | A | W | A | A | W | A | W | W | A | W | A | W | A | A | A | A | W | W | A | W | A | W | W | A | A | W | W | A | W | A | W | A | A | W | A | W | A | W | A | A | A | A | A | W | A | A | A | W | A | A | W | W | W | A | W | W | A | W | W | A | A | W | W | W | A | W | W | A | A | W | A | W | W | A | W | W | A | W | A | W | W |
| Chrl | 14.2 | NA | A | A | W | W | A | A | W | A | NA | A | A | W | A | A | W | A | W | W | A | W | NA | W | A | A | A | A | NA | W | A | W | A | W | NA | A | NA | W | W | A | W | A | W | NA | A | W | NA | W | A | W | A | A | A | A | A | W | A | A | A | W | A | NA | W | W | W | A | W | W | NA | W | W | A | A | W | W | W | A | NA | W | NA | NA | W | NA | W | W | A | W | NA | A | W | A | W | W |
| Chrl | 14.3 | A | A | A | W | W | A | A | W | A | A | A | A | W | A | A | W | A | W | W | A | W | A | W | A | A | A | A | W | W | A | W | A | W | W | A | A | W | W | A | W | A | W | A | A | W | A | W | A | W | A | A | A | A | A | W | A | A | A | W | A | A | W | W | W | A | W | W | A | W | W | A | A | W | W | W | A | W | W | A | A | W | A | W | W | A | W | W | A | W | A | W | W |
| Chrl | 14.4 | NA | A | A | W | W | A | A | W | A | A | A | A | W | A | A | W | A | W | W | A | W | A | NA | A | A | A | A | NA | W | A | NA | A | W | NA | A | A | W | NA | A | W | A | W | A | A | W | A | W | A | W | A | A | A | A | NA | W | A | A | A | W | A | A | W | W | W | A | NA | NA | A | W | W | A | A | W | W | W | NA | NA | W | A | A | W | A | NA | W | A | W | NA | A | NA | NA | W | NA |
| Chrl | 14.5 | A | A | A | W | W | A | A | W | A | A | A | A | W | A | A | W | A | W | W | A | W | A | W | A | A | A | A | W | W | A | W | A | W | W | A | A | W | W | A | W | A | W | A | A | W | A | W | A | W | A | A | A | A | A | W | A | A | A | W | A | A | W | W | W | A | W | W | A | W | W | A | A | W | W | W | A | W | W | A | A | W | A | W | W | A | W | W | A | W | A | W | W |
| Chrl | 14.6 | A | A | A | W | W | A | A | W | A | A | A | A | W | A | A | W | A | W | W | A | W | A | W | A | A | A | A | W | W | A | W | A | W | W | A | A | W | W | A | W | A | W | A | A | W | A | W | A | W | A | A | A | A | A | W | A | A | A | W | A | A | W | W | W | A | W | W | A | W | W | A | A | W | W | W | A | W | W | A | A | W | A | W | W | A | W | W | A | W | A | W | W |
| Chrl | 14.7 | A | A | A | W | W | A | A | W | A | A | A | A | W | A | A | W | A | W | W | A | W | A | NA | A | A | A | A | NA | W | A | W | A | W | NA | A | A | W | W | A | W | A | W | A | A | W | A | W | A | W | A | A | A | A | A | W | A | A | A | W | A | A | W | W | W | A | W | W | A | W | W | A | A | W | W | W | A | W | W | A | A | W | A | NA | W | A | W | W | A | W | A | W | W |
| Chrl | 14.8 | A | A | A | W | W | A | A | W | A | A | A | A | W | A | A | W | A | W | W | A | W | A | W | A | A | A | A | W | W | A | W | A | W | NA | A | A | W | W | A | W | A | W | A | A | W | A | W | A | W | A | A | A | A | A | W | A | A | A | W | A | A | W | W | W | A | W | W | A | W | W | A | A | W | W | W | A | W | W | A | A | W | A | W | W | A | W | W | A | W | A | W | W |
| Chrl | 14.9 | A | A | A | W | W | A | A | W | A | A | A | A | W | A | A | W | A | W | W | A | W | A | W | A | A | A | A | NA | W | A | W | A | W | W | A | A | W | W | A | W | A | W | A | A | W | A | W | A | W | A | A | A | A | A | W | A | A | A | W | A | A | W | W | W | A | W | NA | A | W | W | A | A | W | W | W | A | W | W | A | A | W | A | W | W | A | W | W | A | W | A | W | W |
| Chrl | 15 | A | A | A | W | W | A | A | W | A | A | A | A | W | A | A | W | A | W | W | A | W | A | W | A | A | A | A | W | W | A | W | A | W | W | A | A | W | W | A | W | A | W | A | A | W | A | W | A | W | A | A | A | A | A | W | A | A | A | W | A | A | W | W | W | A | W | W | A | W | W | A | A | W | W | W | A | W | W | A | A | W | A | W | W | A | W | W | A | W | A | W | W |
| Chrl | 15.1 | A | A | A | W | W | A | A | W | A | A | A | A | W | A | A | W | A | W | W | A | W | A | W | A | A | A | A | W | W | A | W | A | W | W | A | A | W | W | A | W | A | W | A | A | W | A | W | A | W | A | A | A | A | A | W | A | A | A | W | A | A | W | W | W | A | W | W | A | W | W | A | A | W | W | W | A | W | W | A | A | W | A | W | W | A | W | W | A | W | A | W | W |
| Chrl | 15.2 | A | A | A | W | W | A | A | W | A | A | A | A | W | A | A | W | A | W | W | A | W | A | W | A | A | A | A | W | W | A | W | A | W | W | A | A | W | W | A | W | A | W | A | A | W | A | W | A | W | A | A | A | A | A | W | A | A | A | W | A | A | W | W | W | A | W | W | A | W | W | A | A | W | W | W | A | W | W | A | A | W | A | W | W | A | W | NA | A | W | A | W | W |
| Chrl | 15.3 | A | A | A | W | W | A | A | W | A | A | A | A | W | A | A | W | A | W | W | A | W | A | W | A | A | A | A | W | W | A | W | A | W | W | A | A | W | W | A | W | A | W | A | A | W | A | W | A | W | A | A | A | A | A | W | A | A | A | W | A | A | W | W | W | A | W | W | A | W | W | A | A | W | W | W | A | W | W | A | A | W | A | W | W | A | W | W | A | W | A | W | W |
| Chrl | 15.4 | A | A | A | W | W | A | A | W | A | A | A | A | W | A | A | W | A | W | W | A | W | A | W | A | A | A | A | NA | W | A | W | A | W | W | A | A | W | W | A | W | A | W | A | A | W | A | W | A | W | A | A | A | A | A | W | A | A | A | W | A | A | W | W | W | A | W | NA | A | W | W | A | A | W | W | W | A | W | W | A | A | W | A | W | W | A | W | W | A | W | A | W | W |
| Chrl | 15.5 | A | A | A | W | W | A | A | W | A | A | A | A | W | A | A | W | A | W | W | A | W | A | W | A | A | A | A | W | W | A | W | A | W | W | A | A | W | W | A | W | A | W | A | A | W | A | W | A | W | NA | A | A | A | A | W | A | A | A | W | A | A | W | W | W | A | W | W | A | W | W | A | A | W | W | W | A | W | W | A | A | W | A | W | W | A | W | W | A | W | A | W | W |
| Chrl | 15.6 | A | A | A | W | W | A | A | W | A | A | A | A | W | A | A | W | A | W | W | A | W | A | W | A | A | A | A | W | W | A | W | A | W | W | A | A | W | W | A | W | A | W | A | A | W | A | W | A | W | A | A | A | A | A | W | A | A | A | W | A | A | W | W | W | A | W | NA | A | W | W | A | A | W | W | W | A | W | W | A | A | W | A | W | W | A | W | W | A | W | A | W | W |
| Chrl | 15.7 | A | A | A | W | W | A | A | W | A | A | A | A | W | A | A | W | A | W | W | A | W | A | W | A | A | A | A | W | W | A | W | A | W | W | A | A | W | W | A | W | A | W | A | A | W | A | W | A | W | NA | A | A | A | A | W | A | A | A | W | A | A | W | W | W | NA | W | NA | A | W | W | A | A | W | W | W | A | W | W | A | A | W | A | W | W | A | W | NA | A | W | A | W | W |
| Chrl | 15.9 | A | A | A | W | W | A | A | W | A | A | A | A | W | A | A | W | A | W | W | A | W | A | NA | A | A | A | NA | NA | W | A | W | A | W | W | A | A | W | W | A | W | A | W | A | A | W | A | W | A | W | NA | A | A | A | A | W | A | A | A | W | A | A | W | W | W | A | W | W | A | W | W | A | A | W | W | W | A | W | W | A | A | W | A | W | W | A | W | W | A | W | A | W | W |
| Chrl | 16 | A | A | A | W | W | A | A | W | A | A | A | A | W | A | A | W | A | W | W | A | W | A | W | A | A | A | A | W | W | A | W | A | W | W | A | A | W | NA | A | W | A | W | A | A | W | A | W | A | W | A | A | A | A | A | NA | A | A | A | W | A | A | W | W | W | A | W | W | A | W | W | A | A | W | W | W | A | W | W | A | A | W | A | NA | W | A | W | W | A | W | A | W | NA |
| Chrl | 16.1 | A | A | A | W | W | A | A | W | A | A | A | A | W | A | NA | W | A | W | W | A | W | A | W | A | A | A | A | W | W | A | W | A | W | W | A | A | W | W | A | W | A | W | A | A | W | A | W | A | W | A | A | A | A | A | W | A | A | A | W | A | A | W | W | W | A | W | W | A | W | W | A | A | W | W | W | A | W | W | A | A | W | A | W | W | A | W | W | A | W | A | W | W |
| Chrl | 16.2 | A | A | A | W | W | A | A | W | A | A | NA | NA | W | A | A | W | NA | W | W | A | W | A | W | NA | A | A | NA | W | W | A | W | A | W | W | A | A | W | W | A | W | A | W | A | A | W | NA | W | A | W | A | A | A | A | A | W | A | A | A | W | A | A | W | W | W | A | W | W | A | W | W | A | A | W | W | W | A | W | W | NA | A | W | A | W | W | A | W | W | A | W | A | W | W |
| Chrl | 16.3 | A | A | A | W | W | A | A | W | A | A | A | A | W | A | NA | W | A | W | W | A | W | A | W | A | A | A | A | W | W | A | W | A | W | W | A | A | W | W | A | W | A | W | A | A | W | A | W | A | W | NA | A | A | NA | A | W | A | A | A | W | A | A | W | W | W | NA | W | W | A | W | W | A | A | W | W | W | A | W | W | A | NA | W | A | W | W | A | W | W | A | W | A | W | W |
| Chrl | 16.4 | A | A | A | W | W | NA | A | W | A | A | A | NA | W | NA | A | W | A | W | W | NA | W | A | W | A | A | NA | A | W | W | A | W | A | W | W | A | NA | W | W | A | W | NA | W | NA | A | W | A | W | NA | W | A | NA | NA | A | A | W | A | NA | A | W | A | A | W | W | W | NA | NA | W | A | W | W | NA | A | W | W | W | NA | W | W | NA | A | W | A | W | W | NA | W | NA | A | W | A | W | W |
| Chrl | 16.5 | A | A | A | W | W | A | A | W | A | A | A | A | W | A | A | W | A | W | W | A | W | A | W | A | A | A | A | NA | W | A | W | A | W | W | A | A | W | W | A | W | A | W | A | A | W | A | W | A | W | NA | A | A | A | A | W | A | A | A | W | A | A | W | W | W | A | W | NA | A | W | W | A | A | W | W | W | A | W | W | A | A | W | A | W | W | A | W | W | A | W | A | W | W |
| Chrl | 16.6 | A | A | A | W | W | A | A | W | A | A | A | A | W | A | A | W | A | W | W | A | W | A | W | A | A | A | A | W | W | A | W | A | W | W | A | A | W | W | A | W | A | W | A | A | W | A | W | A | W | A | A | A | A | A | W | A | A | A | W | A | A | W | W | W | A | W | W | A | W | W | A | A | W | W | W | A | W | W | A | A | W | A | W | W | A | W | W | A | W | A | W | W |
| Chrl | 16.7 | A | A | A | W | W | A | A | W | A | A | A | A | W | A | A | W | A | W | W | A | W | A | W | A | A | A | A | NA | W | A | W | A | W | NA | A | A | W | W | A | W | A | W | A | A | W | A | W | A | W | A | A | A | A | A | W | A | A | A | W | A | A | W | W | W | A | NA | W | A | W | W | A | A | W | W | W | A | W | W | A | A | W | A | W | W | A | W | W | A | W | A | W | W |
| Chrl | 16.8 | A | A | A | W | W | A | A | W | A | A | A | A | W | A | A | W | A | W | W | A | W | A | NA | A | A | A | A | W | W | A | W | A | W | W | A | A | W | W | A | W | A | W | A | A | W | A | W | A | W | A | A | A | A | A | W | A | A | A | W | A | A | W | W | W | NA | W | W | A | W | W | A | A | W | W | W | A | W | W | A | A | W | A | W | W | A | W | NA | A | W | A | W | NA |
| Chrl | 16.9 | A | A | A | W | W | A | A | W | A | A | A | A | W | A | A | W | A | W | W | A | W | A | W | A | A | A | A | NA | W | A | W | A | W | W | A | A | W | W | A | W | A | W | A | A | W | A | W | A | W | A | A | A | A | A | W | A | A | A | W | A | A | W | W | W | A | W | W | A | W | W | A | A | W | W | W | A | W | W | A | A | W | A | W | W | A | W | NA | A | W | A | W | W |
| Chrl | 17 | A | A | A | W | W | A | A | W | A | A | A | A | W | A | A | W | A | W | W | A | W | A | W | A | A | A | A | W | W | A | W | A | W | W | A | A | W | W | A | W | A | W | A | A | W | A | W | A | W | A | A | A | A | A | W | A | A | A | W | A | A | W | W | W | A | W | W | A | W | W | A | A | W | W | W | A | W | W | A | A | W | A | W | W | A | W | W | A | W | A | W | W |
| Chrl | 17.1 | A | A | A | W | W | A | A | W | A | A | A | A | W | A | A | W | A | W | W | A | W | A | W | A | A | A | A | W | W | A | W | A | W | W | A | A | W | W | A | W | A | W | A | A | W | A | W | A | W | A | A | A | A | A | W | A | A | A | W | A | A | W | W | W | A | W | W | A | W | W | A | A | W | W | W | A | W | W | A | A | W | A | W | W | A | W | W | A | W | A | W | W |
| Chrl | 17.2 | A | A | A | W | W | A | A | W | A | A | A | A | W | A | A | W | A | W | W | A | W | A | W | A | A | A | A | W | W | A | W | A | W | W | A | A | W | W | A | W | A | W | A | A | W | A | W | A | W | A | A | A | A | A | W | A | A | A | W | A | A | W | W | W | A | W | W | A | W | W | A | A | W | W | W | A | W | W | A | A | W | A | W | W | A | W | W | A | W | A | W | W |
| Chrl | 17.3 | A | A | A | W | W | A | A | W | A | A | A | A | W | NA | A | W | NA | W | W | NA | W | A | W | A | A | A | A | W | W | A | W | A | W | W | A | A | W | W | A | W | A | W | A | A | W | A | W | A | W | A | A | A | A | A | W | A | A | A | W | A | A | W | W | W | A | W | W | A | W | W | A | A | W | W | W | A | W | W | NA | A | W | A | W | W | NA | W | W | A | W | A | W | W |
| Chrl | 17.4 | A | A | A | W | W | A | A | W | A | A | A | A | W | A | A | W | A | W | W | A | W | A | NA | A | A | A | A | W | W | A | W | A | W | W | A | A | W | W | A | W | A | W | A | A | W | A | W | A | W | A | A | A | A | A | W | A | A | A | W | A | A | W | W | W | A | W | NA | A | W | W | A | A | W | W | W | A | W | W | A | A | W | A | W | W | A | W | W | A | W | A | W | W |
| Chrl | 17.5 | A | A | A | W | W | A | A | W | A | A | A | A | W | A | NA | W | A | W | W | A | W | A | W | A | A | A | A | W | W | A | W | A | W | W | A | A | W | W | A | W | A | W | A | A | W | A | W | A | W | A | A | A | A | A | W | A | A | A | W | A | A | W | W | W | A | W | NA | A | W | W | A | A | W | W | W | A | W | W | A | A | W | A | W | W | A | W | W | A | W | A | W | W |
| Chrl | 17.6 | A | A | A | W | W | A | A | W | A | A | A | A | W | A | A | W | A | W | W | A | W | A | NA | A | A | A | A | W | W | A | W | A | W | W | A | A | W | W | A | W | A | W | A | A | W | A | W | A | W | A | A | A | A | A | W | A | A | A | W | A | A | W | W | W | A | W | W | A | W | W | A | A | W | W | W | A | W | W | A | A | W | A | W | W | A | W | W | A | W | A | W | W |
| Chrl | 17.7 | A | A | A | W | W | A | A | W | A | A | A | A | W | A | A | W | A | W | W | A | W | A | W | A | A | A | A | W | W | A | W | A | W | W | A | A | W | W | A | W | A | W | A | A | W | A | W | A | W | A | A | A | A | A | W | A | A | A | W | A | A | W | W | W | A | W | W | A | W | W | A | A | W | W | W | A | W | W | A | A | W | A | W | W | A | W | W | A | W | A | W | W |
| Chrl | 17.8 | A | A | A | W | W | A | A | W | A | A | A | A | W | A | A | W | A | W | W | A | W | A | W | A | A | A | A | W | W | A | W | A | W | W | A | A | W | W | A | W | A | W | A | A | W | A | W | A | W | A | A | A | A | A | W | A | A | A | W | A | A | W | W | W | A | W | W | A | W | W | A | A | W | W | W | A | W | W | A | A | W | A | W | W | A | W | W | A | W | A | W | W |
| Chrl | 17.9 | A | A | A | W | W | A | A | W | A | A | A | A | W | A | A | W | A | W | W | A | W | A | NA | A | A | A | A | W | W | A | W | A | W | W | A | A | W | W | A | W | A | W | A | A | W | A | W | A | W | A | A | A | A | A | W | A | A | A | W | A | A | W | W | W | A | W | W | A | W | W | A | A | W | W | W | A | W | W | A | A | W | A | W | W | A | W | W | A | W | A | W | W |

| | | | | | | | | | | | | | | | | | | | | | | | | | | | | | | | | | | | | | | | | | | | | | | | | | | | | | | | | | | | | | | | | | | | | | | | | | | | | | | | | | | | | | | | | | | | | | |
|---|---|---|---|---|---|---|---|---|---|---|---|---|---|---|---|---|---|---|---|---|---|---|---|---|---|---|---|---|---|---|---|---|---|---|---|---|---|---|---|---|---|---|---|---|---|---|---|---|---|---|---|---|---|---|---|---|---|---|---|---|---|---|---|---|---|---|---|---|---|---|---|---|---|---|---|---|---|---|---|---|---|---|---|---|---|---|---|---|---|---|---|---|---|
| Chrl | 18 | A | A | A | W | W | A | A | W | A | A | A | A | W | A | A | W | A | W | W | A | W | A | W | A | A | A | A | W | W | A | W | A | W | W | A | A | W | W | A | W | A | W | A | A | W | A | W | A | W | A | A | A | A | A | W | A | A | A | W | A | A | W | W | W | A | W | W | A | W | W | A | A | W | W | W | A | W | W | A | A | W | A | W | W | A | W | W | A | W | A | W | W |
| Chrl | 18.1 | A | A | A | W | W | A | A | W | A | A | A | A | W | A | A | W | A | W | W | A | W | A | NA | A | A | A | A | W | W | A | W | A | W | W | A | A | W | W | A | W | A | W | A | A | W | A | W | A | W | A | A | A | A | A | W | A | A | A | W | A | A | W | W | W | A | W | NA | A | W | W | A | A | W | W | W | A | W | W | A | A | W | A | W | W | A | W | NA | A | W | A | W | W |
| Chrl | 18.2 | A | A | A | W | W | A | A | W | A | A | A | A | W | A | A | W | A | W | W | A | W | A | W | A | A | A | A | W | W | A | W | A | W | W | A | A | W | W | A | W | A | W | A | A | W | A | W | A | W | A | A | A | A | A | W | A | A | A | W | A | A | W | W | W | A | W | W | A | W | W | A | A | W | W | W | A | W | W | A | A | W | A | W | W | A | W | W | A | W | A | W | W |
| Chrl | 18.3 | A | A | A | W | W | A | A | W | A | A | A | A | W | A | A | W | A | W | W | A | W | A | W | A | A | A | A | W | W | A | W | A | W | W | A | A | W | W | NA | W | A | W | A | A | W | A | W | A | W | A | A | A | A | A | W | A | A | A | W | A | A | W | W | W | A | W | W | A | W | W | A | A | W | W | W | NA | W | W | A | A | W | A | W | W | A | W | W | A | W | A | W | W |
| Chrl | 18.4 | A | A | A | W | W | A | A | W | A | A | A | A | W | A | A | W | A | W | W | A | W | A | W | A | A | A | A | W | W | A | W | A | W | NA | A | A | W | W | A | W | A | W | A | A | W | A | W | A | W | A | A | A | A | A | W | A | A | A | W | A | A | W | W | W | A | W | W | A | W | W | A | A | W | W | W | A | W | W | A | A | W | A | W | W | A | W | W | A | W | A | W | W |
| Chrl | 18.5 | A | A | A | W | W | A | A | W | A | A | A | A | W | A | A | W | A | W | W | A | W | A | W | A | A | A | A | W | W | A | W | A | W | NA | A | A | W | W | A | W | A | W | A | A | W | A | W | A | W | NA | A | A | A | A | W | A | A | A | W | A | A | W | W | W | A | W | NA | A | W | W | A | A | W | W | W | A | W | W | A | A | W | A | W | W | A | W | W | A | W | A | W | W |
| Chrl | 18.6 | A | A | A | W | W | A | A | W | A | A | A | A | W | A | A | W | A | W | W | A | W | A | W | A | A | A | A | NA | W | A | W | A | W | W | A | A | W | W | A | W | A | W | A | A | W | A | W | A | W | A | A | A | A | A | W | A | A | A | W | A | A | W | W | W | A | W | W | A | W | W | A | A | W | W | W | A | W | W | A | A | W | A | W | W | A | W | W | A | W | A | W | W |
| Chrl | 18.7 | A | A | A | W | W | A | A | W | A | A | A | A | W | A | A | W | A | W | W | A | W | A | NA | A | A | A | A | W | W | A | W | A | W | W | A | A | W | W | A | W | A | W | A | A | W | A | W | A | W | A | A | A | A | A | W | A | A | A | W | A | A | W | W | W | A | W | NA | A | W | W | A | A | W | W | W | A | W | W | A | A | W | A | W | W | A | W | NA | A | W | A | W | W |
| Chrl | 18.8 | A | A | A | W | W | A | A | W | A | A | A | A | W | A | A | W | A | W | W | A | W | A | W | A | A | A | A | W | W | A | W | A | W | W | A | A | W | W | A | W | A | W | A | A | W | A | W | A | W | A | A | A | A | A | W | A | A | A | W | A | A | W | W | W | A | W | W | A | W | W | A | A | W | W | W | A | W | W | A | A | W | A | W | W | A | W | W | A | W | A | W | W |
| Chrl | 18.9 | A | A | A | W | W | A | A | W | A | A | A | A | W | A | A | W | A | W | W | A | W | A | NA | A | A | A | A | W | W | A | W | A | W | NA | A | A | W | W | A | W | A | W | A | A | W | A | W | A | W | A | A | A | A | A | W | A | A | A | W | A | A | W | W | W | A | W | W | A | W | W | A | A | W | W | W | A | W | W | A | A | W | A | W | W | A | W | W | A | W | A | W | W |
| Chrl | 19 | A | A | A | W | W | A | A | W | A | A | A | A | W | A | A | W | A | W | W | A | W | A | NA | A | A | A | A | W | W | A | W | A | W | NA | A | A | W | W | A | W | A | W | A | A | W | A | W | A | W | A | A | A | A | A | W | A | A | A | W | A | A | W | W | W | A | NA | W | A | W | W | A | A | W | W | W | A | W | W | A | A | W | A | W | W | A | W | W | A | W | A | W | W |
| Chrl | 19.1 | A | A | A | W | W | A | A | W | A | A | A | A | W | A | A | W | A | W | W | A | W | A | W | A | A | A | A | W | W | A | W | A | NA | W | A | A | W | W | A | W | A | W | A | A | W | A | W | A | W | A | A | A | A | A | W | A | A | A | W | A | A | W | W | W | A | W | W | A | W | W | A | A | W | W | W | A | W | W | A | A | W | A | W | W | A | W | W | A | W | A | W | W |
| Chrl | 19.2 | A | A | A | W | W | A | A | W | A | A | A | A | W | A | A | W | A | W | W | A | W | A | NA | A | A | A | A | W | W | A | W | A | W | NA | A | A | W | W | A | W | A | W | A | A | W | A | W | A | W | A | A | A | A | A | W | A | A | A | W | A | A | W | W | W | A | W | W | A | W | W | A | A | W | W | W | A | W | W | A | A | W | A | W | W | A | W | W | A | W | A | W | W |
| Chrl | 19.3 | A | A | A | W | W | A | A | W | A | A | A | A | W | A | A | W | A | W | W | A | W | A | W | A | A | A | NA | W | W | A | W | A | W | W | A | A | W | W | A | W | A | W | A | A | W | A | W | A | W | A | A | A | A | A | W | A | A | A | W | A | A | W | W | W | A | W | W | A | W | W | A | A | W | W | W | A | W | W | A | A | W | A | W | W | A | W | W | A | W | A | W | W |
| Chrl | 19.4 | A | A | A | W | W | A | A | W | A | A | A | A | W | A | A | W | A | W | W | A | W | A | W | A | A | A | A | W | W | A | W | A | W | W | A | A | W | W | A | W | A | W | A | A | W | A | W | A | W | A | A | A | A | A | W | A | A | A | W | A | A | W | W | W | A | W | W | A | W | W | A | A | W | W | W | A | W | W | A | A | W | A | W | W | A | W | W | A | W | A | W | W |
| Chrl | 19.5 | A | A | A | W | W | NA | A | W | A | A | NA | A | W | A | NA | W | NA | W | W | A | W | A | W | A | NA | NA | A | W | W | NA | W | A | W | W | NA | NA | W | W | A | W | NA | W | A | NA | W | NA | W | A | W | A | NA | A | A | NA | W | A | A | A | W | A | NA | W | W | W | NA | W | W | A | W | W | NA | A | W | W | W | NA | W | W | NA | NA | W | NA | W | W | A | W | W | NA | W | A | W | W |
| Chrl | 19.6 | A | A | A | W | W | A | A | W | A | A | A | A | W | A | A | W | A | W | W | A | W | A | W | A | A | A | A | W | W | A | W | A | W | NA | A | A | W | W | A | W | A | W | A | A | W | A | W | A | W | A | A | A | A | A | W | A | A | A | W | A | A | W | W | W | A | W | W | A | W | W | A | A | W | W | W | A | W | W | A | A | W | A | W | W | A | W | W | A | W | A | W | W |
| Chrl | 19.7 | A | A | A | W | W | A | A | W | A | A | A | A | W | A | A | W | A | W | W | A | W | A | W | A | A | A | A | W | W | A | W | A | W | W | A | A | W | W | A | W | A | W | A | A | W | A | W | A | W | A | A | A | A | A | W | A | A | A | W | A | A | W | W | W | A | W | W | A | W | W | A | A | W | W | W | A | W | W | A | A | W | A | W | W | A | W | W | A | W | A | W | W |
| Chrl | 19.8 | A | A | A | W | W | A | A | W | A | A | A | A | W | A | A | W | A | W | W | A | W | A | W | A | A | A | A | W | W | A | W | A | W | NA | A | A | W | W | A | W | A | W | A | A | W | A | W | A | W | A | A | A | A | A | W | A | A | A | W | A | A | W | W | W | A | W | W | A | W | W | A | A | W | W | W | A | W | W | A | A | W | A | W | W | A | W | W | A | W | A | W | W |
| Chrl | 19.9 | A | A | A | W | W | A | A | W | A | A | A | A | W | A | A | W | A | W | W | A | W | A | W | A | A | A | A | W | W | A | W | A | W | W | A | A | W | W | A | W | A | W | A | A | W | A | W | A | W | A | A | A | A | A | W | A | A | A | W | A | A | W | W | W | A | W | W | A | W | W | A | A | W | W | W | A | W | W | A | A | W | A | W | W | A | W | W | A | W | A | W | W |
| Chrl | 20 | A | A | A | W | W | A | NA | W | A | A | A | A | W | A | A | W | A | W | W | A | W | A | W | A | A | A | A | W | W | A | W | A | W | NA | A | A | W | W | A | W | A | W | A | A | W | A | W | A | W | A | A | A | A | A | W | A | A | A | W | A | A | W | W | W | NA | NA | W | A | W | W | A | A | W | W | W | A | W | W | A | A | W | A | W | W | A | W | W | A | W | A | W | W |
| Chrl | 20.1 | A | A | A | W | W | A | A | W | A | A | A | A | W | A | A | W | A | W | W | A | W | A | W | A | A | NA | A | W | W | A | W | A | W | NA | A | A | W | W | A | W | A | W | A | A | W | A | W | A | W | A | A | A | A | A | W | A | A | A | W | A | A | W | W | W | A | W | W | A | W | W | A | A | W | W | W | A | W | W | A | A | W | A | W | W | A | W | W | A | W | A | W | W |
| Chrl | 20.2 | A | A | A | W | W | A | A | W | A | A | A | A | W | A | A | W | A | W | W | A | W | A | W | A | A | A | A | W | W | A | W | A | W | W | A | A | W | W | A | W | A | W | A | A | W | A | W | A | W | A | A | A | A | A | W | A | A | A | W | A | A | W | W | W | A | W | W | A | W | W | A | A | W | W | W | A | W | W | A | A | W | A | W | W | A | W | W | A | W | A | W | W |
| Chrl | 20.3 | A | A | A | W | W | A | A | W | A | A | A | A | W | A | A | W | A | W | W | A | W | A | W | A | A | A | A | W | W | A | W | A | W | W | A | A | W | W | A | W | A | W | A | A | W | A | W | A | W | A | A | A | A | A | W | A | A | A | W | A | A | W | W | W | A | W | NA | A | W | W | A | A | W | W | W | A | W | W | A | A | W | A | W | W | A | W | W | A | W | A | W | W |
| Chrl | 20.4 | A | A | A | W | W | A | A | W | A | A | A | A | W | A | A | W | A | W | W | NA | W | A | W | A | A | A | A | W | W | A | W | A | W | W | A | A | W | W | A | W | A | W | A | A | W | A | W | A | W | A | A | A | A | A | W | A | A | A | W | A | A | W | W | W | A | W | W | A | W | W | A | A | W | W | W | A | W | W | A | A | W | A | W | W | A | W | W | A | W | A | W | W |
| Chrl | 20.5 | A | A | A | W | W | A | A | W | A | A | A | A | W | A | A | W | A | W | W | NA | W | A | W | A | NA | A | A | W | W | A | W | A | W | W | A | A | W | W | A | W | A | W | A | A | W | A | W | A | W | A | A | A | A | A | W | A | A | A | W | A | A | W | W | W | A | W | NA | A | W | W | A | A | W | W | W | A | W | W | A | A | W | A | W | W | A | W | W | A | W | A | W | W |
| Chrl | 20.6 | A | A | A | W | W | A | A | W | A | A | A | A | W | A | A | W | A | W | W | A | W | A | W | A | A | A | A | W | W | A | W | A | W | NA | A | A | W | W | A | W | A | W | A | A | W | A | W | A | W | A | A | A | A | A | W | A | A | A | W | A | A | W | W | W | A | W | NA | A | W | W | A | A | W | W | W | A | W | W | A | A | W | A | W | W | A | W | W | A | W | A | W | W |
| Chrl | 20.7 | A | A | A | W | W | A | A | W | A | A | A | A | W | A | A | W | A | W | W | A | W | A | W | A | A | A | A | W | W | A | W | A | W | NA | A | A | W | W | A | W | A | W | A | A | W | A | W | A | W | A | A | A | A | A | W | A | A | A | W | A | A | W | W | W | A | W | W | A | W | W | A | A | W | W | W | A | W | W | A | A | W | A | W | W | A | W | W | A | W | A | W | NA |
| Chrl | 20.8 | NA | NA | NA | W | W | NA | A | W | A | NA | NA | NA | W | A | NA | W | A | W | W | A | W | A | W | NA | A | A | A | W | W | A | W | A | W | NA | A | A | W | W | A | W | A | W | A | NA | W | A | W | A | W | NA | A | A | A | A | W | A | A | A | W | A | NA | W | W | W | NA | W | NA | A | W | W | A | A | W | W | W | NA | W | W | NA | NA | W | A | W | W | A | W | NA | NA | W | NA | W | W |
| Chrl | 20.9 | A | A | A | W | W | A | A | W | A | A | A | A | W | A | A | W | A | W | W | A | W | A | NA | A | A | A | A | NA | W | A | W | A | W | W | A | A | W | W | A | W | A | W | A | A | W | A | W | A | W | A | A | A | A | A | W | A | A | A | W | A | A | W | W | W | A | W | W | A | W | W | A | A | W | W | W | A | W | W | A | A | W | A | W | W | A | W | W | A | W | A | W | W |
| Chrl | 21 | A | A | A | W | W | A | A | W | A | A | A | A | W | A | A | W | A | W | W | A | W | A | W | A | A | A | A | W | W | A | W | A | W | W | A | A | W | W | A | W | A | W | A | A | W | A | W | A | W | A | A | A | A | A | W | A | A | A | W | A | A | W | W | W | A | NA | W | A | W | W | A | A | W | W | W | A | NA | W | A | A | W | A | W | W | A | W | W | A | W | A | W | W |
| Chrl | 21.1 | A | A | A | W | W | A | A | W | A | A | A | A | W | A | A | W | A | W | W | A | W | A | NA | A | A | A | A | NA | W | A | W | A | W | NA | A | A | W | W | A | W | A | W | A | A | W | A | W | A | W | A | A | A | A | A | W | A | A | A | W | A | A | W | W | W | A | NA | W | A | W | W | A | A | W | W | W | A | W | W | A | A | W | A | W | W | A | W | W | A | W | A | W | W |
| Chrl | 21.2 | A | A | A | W | W | A | A | W | A | A | A | A | W | A | A | W | A | W | W | A | W | A | W | A | A | A | A | W | W | A | W | A | W | NA | A | A | W | W | A | W | A | W | A | A | W | A | W | A | W | NA | A | A | A | A | W | A | A | A | W | A | A | W | W | W | A | W | NA | A | W | W | A | A | W | W | W | A | W | W | A | A | W | A | W | W | A | W | W | A | W | A | W | W |
| Chrl | 21.3 | A | A | A | W | W | A | A | W | A | A | A | A | W | A | A | W | A | W | W | A | W | A | W | A | A | A | A | NA | W | A | W | A | W | NA | A | A | W | W | A | W | A | W | A | A | W | A | W | A | W | A | A | A | A | A | W | A | A | A | W | A | A | W | W | W | A | W | W | A | W | W | A | A | W | W | W | A | W | W | A | A | W | A | W | W | A | W | W | A | W | A | W | W |
| Chrl | 21.4 | A | A | A | W | W | A | A | W | A | A | A | A | W | A | A | W | A | W | W | A | W | A | W | A | A | A | A | W | W | A | W | A | W | NA | A | A | W | W | A | W | A | W | A | A | W | A | W | A | W | A | A | A | A | A | W | A | A | A | W | A | A | W | W | W | A | W | W | A | W | W | A | A | W | W | W | A | W | W | A | A | W | A | W | W | A | W | W | A | W | A | W | W |
| Chrl | 21.5 | A | A | A | W | W | A | A | W | A | A | A | A | W | A | A | W | A | W | W | A | W | A | W | A | A | A | A | W | W | A | W | A | W | W | A | A | W | W | A | W | A | W | A | A | W | A | W | A | W | A | A | A | A | A | W | A | A | A | W | A | A | W | W | W | A | W | W | A | W | W | A | A | W | W | W | A | W | W | A | A | W | A | W | W | A | W | W | A | W | A | W | W |
| Chrl | 21.6 | A | A | A | W | W | A | A | W | A | A | A | A | W | A | A | W | A | W | W | A | W | A | W | A | A | A | A | W | W | A | W | A | W | W | A | A | W | W | A | W | A | W | A | A | W | A | W | A | W | A | A | A | A | A | W | A | A | A | W | A | A | W | W | W | A | W | W | A | W | W | A | A | W | W | W | A | W | W | A | NA | W | A | W | W | A | W | W | A | W | A | W | W |
| Chrl | 21.7 | A | A | A | W | W | A | A | W | A | A | A | A | W | A | A | W | A | W | W | A | W | A | W | A | A | A | A | W | W | A | W | A | W | W | A | A | W | W | A | W | A | W | A | A | W | A | W | A | W | A | A | A | A | A | W | A | A | A | W | A | A | W | W | W | A | W | W | A | W | W | A | A | W | W | W | A | W | W | A | W | W | A | W | W | A | W | W | A | W | A | W | W |
| Chrl | 21.8 | A | A | A | W | W | A | A | W | A | A | A | A | W | A | A | W | A | W | W | A | W | A | W | A | A | A | A | NA | W | A | W | A | W | W | A | A | W | W | A | W | A | W | A | A | W | A | W | A | W | A | A | A | A | A | W | A | A | A | W | A | A | W | W | W | A | W | W | A | W | W | A | A | W | W | W | A | W | W | A | W | W | A | W | W | A | W | W | A | W | A | W | NA |
| Chrl | 22 | A | A | A | W | W | A | A | W | A | A | A | A | W | A | A | W | A | W | W | A | W | A | W | A | A | A | A | W | W | A | W | A | W | W | A | A | W | W | A | W | A | W | A | A | W | A | W | A | W | A | A | A | A | A | W | A | A | A | W | A | A | W | W | W | A | W | W | A | W | W | A | A | W | W | W | A | W | W | A | W | W | A | W | W | A | W | W | A | W | A | W | W |
| Chrl | 22.1 | A | A | A | W | W | A | A | W | A | A | A | A | W | A | A | W | A | W | W | A | W | A | W | A | A | A | A | NA | W | A | W | A | W | NA | A | A | W | W | A | W | A | W | A | A | W | A | W | A | W | A | A | A | A | A | W | A | A | A | W | A | A | W | W | W | A | W | NA | A | W | W | A | A | W | W | W | A | W | W | A | W | W | A | W | W | A | W | W | A | W | A | W | W |
| Chrl | 22.2 | A | A | A | W | W | A | A | W | A | A | A | A | W | A | A | W | A | W | W | A | W | A | W | A | A | A | A | NA | W | A | W | A | W | W | A | A | W | W | A | W | A | W | A | A | W | A | W | A | W | A | A | A | A | A | W | A | A | A | W | A | A | W | W | W | A | W | W | A | W | W | A | A | W | W | W | A | W | W | A | W | W | A | W | W | A | W | W | A | W | A | W | W |
| Chrl | 22.3 | A | A | A | W | W | A | A | W | A | A | A | A | W | A | A | W | A | W | W | A | W | A | W | A | A | A | A | W | W | A | W | A | W | NA | A | A | W | W | A | W | A | W | A | A | W | A | W | A | W | A | A | A | A | A | W | A | A | A | W | A | A | W | W | W | A | W | W | A | W | W | A | A | W | W | W | A | W | W | A | W | W | A | W | W | A | W | W | A | W | A | W | W |
| Chrl | 22.4 | A | A | A | W | W | A | A | W | A | A | A | A | W | A | A | W | A | W | W | A | W | A | W | A | A | A | A | W | W | A | W | A | W | W | A | A | W | W | A | W | A | W | A | A | W | A | W | A | W | A | A | A | A | A | W | A | A | A | W | A | A | W | W | W | A | W | W | A | W | W | A | A | W | W | W | A | W | W | A | W | W | A | W | W | A | W | W | A | W | A | W | W |
| Chrl | 22.5 | A | A | A | W | W | A | A | W | A | A | A | A | W | A | A | W | A | W | W | A | W | A | W | A | A | A | A | W | W | A | W | A | W | NA | A | A | W | W | A | W | A | W | A | A | W | A | W | A | W | A | A | A | A | A | W | A | A | A | W | A | A | W | W | W | A | W | W | A | W | W | A | A | W | W | W | A | W | W | A | W | W | A | W | W | A | W | NA | A | W | A | W | W |
| Chrl | 22.6 | A | A | A | W | W | A | A | W | A | A | A | A | W | A | A | W | A | W | W | A | W | A | W | A | A | A | A | NA | W | A | W | A | W | NA | A | A | W | W | A | W | A | W | A | A | W | A | W | A | W | A | A | A | A | A | W | A | A | A | W | A | A | W | W | W | A | W | W | A | W | W | A | A | W | W | W | A | W | W | A | W | W | A | W | W | A | W | W | A | W | A | W | W |
| Chrl | 22.7 | A | A | A | W | W | A | A | W | A | A | A | A | W | A | A | W | A | W | W | A | W | A | W | A | A | A | A | W | W | A | W | A | W | W | A | A | W | W | A | W | A | W | A | A | W | A | W | A | W | A | A | A | A | A | W | A | A | A | W | A | A | W | W | W | A | W | NA | A | W | W | A | A | W | W | W | A | W | W | A | W | W | A | W | W | A | W | W | A | W | A | W | W |
| Chrl | 22.9 | A | A | A | W | W | A | A | W | A | A | A | A | W | A | A | W | A | W | W | A | W | A | W | A | A | A | A | W | W | A | W | A | W | W | A | A | W | W | A | W | A | W | A | A | W | A | W | A | W | A | A | A | A | A | W | A | A | A | W | A | A | W | W | W | NA | W | W | A | W | W | A | A | W | W | W | A | W | W | A | W | W | A | W | W | A | W | W | A | W | A | W | W |
| Chrl | 23 | A | A | A | W | W | A | A | W | A | A | A | A | W | A | A | W | A | W | W | A | W | A | W | A | A | A | A | W | W | A | W | A | W | W | A | A | W | W | A | W | A | W | A | A | W | A | W | A | W | A | A | A | A | A | W | A | A | A | W | A | A | W | W | W | A | W | W | A | W | W | A | A | W | W | W | A | W | W | A | W | W | A | W | W | A | W | W | A | W | A | W | W |
| Chrl | 23.1 | A | A | A | W | W | A | A | W | A | A | A | A | W | A | A | W | A | W | W | A | W | A | W | A | A | A | A | W | W | A | W | A | W | NA | A | A | W | W | A | W | A | W | A | A | W | A | W | A | W | A | A | A | A | A | W | A | A | A | W | A | A | W | W | W | NA | W | NA | A | W | W | A | A | W | W | W | A | W | W | A | W | W | A | W | W | A | W | W | A | W | A | W | W |
| Chrl | 23.2 | A | NA | A | W | W | A | A | W | A | A | A | A | W | A | A | W | A | W | W | NA | W | NA | W | A | A | A | A | W | W | A | W | A | W | W | A | A | W | W | A | W | A | W | NA | A | W | A | W | A | W | A | NA | A | A | A | W | A | A | A | W | NA | A | W | W | W | A | W | W | A | W | W | A | A | W | W | W | A | W | W | NA | W | W | A | W | W | A | W | W | A | W | A | W | W |
| Chrl | 23.4 | A | A | A | W | W | A | A | W | A | A | A | A | W | A | A | W | A | W | W | A | W | A | W | A | A | A | A | W | W | A | W | A | W | NA | A | A | W | W | A | W | A | W | A | A | W | A | W | A | W | A | A | A | A | A | W | A | A | A | W | A | A | W | W | W | A | W | NA | A | W | W | A | A | W | W | W | A | W | W | A | W | W | A | W | W | A | W | W | A | W | A | W | W |
| Chrl | 23.5 | A | A | A | W | W | A | A | W | A | A | A | A | W | A | A | W | A | W | W | A | W | A | W | A | A | A | A | NA | W | A | W | A | W | NA | A | A | W | W | A | W | A | W | A | A | W | A | W | A | W | A | A | A | A | A | W | A | A | A | W | A | A | W | W | W | A | W | NA | A | W | W | A | A | W | W | W | A | W | W | A | W | W | A | W | W | A | W | NA | A | W | A | W | W |
| Chrl | 23.6 | A | A | A | W | W | A | A | W | A | A | A | A | W | A | A | W | A | W | W | A | W | A | NA | A | A | A | A | W | W | A | W | A | W | NA | A | A | W | W | A | W | A | W | A | A | W | A | W | A | W | A | A | A | A | A | W | A | A | A | W | A | A | W | W | W | A | W | W | A | W | W | A | A | W | W | W | A | W | W | A | W | W | A | W | W | A | W | W | A | W | A | W | W |
| Chrl | 23.7 | A | A | A | W | W | A | A | W | A | A | A | A | W | A | A | W | A | W | W | NA | W | A | W | A | A | A | A | W | W | A | W | A | W | W | A | A | W | W | A | W | A | W | A | A | W | A | W | A | W | A | A | A | A | A | W | A | A | A | W | A | A | W | W | W | A | W | W | A | W | W | A | A | W | W | W | A | W | W | A | W | W | A | W | W | A | W | W | A | W | A | W | W |
| Chrl | 23.8 | A | A | A | W | W | A | A | W | A | A | A | A | W | A | A | W | A | W | W | A | W | A | W | A | A | A | A | W | W | A | W | A | W | W | A | A | W | W | A | W | A | W | A | A | W | A | W | A | W | A | A | A | A | A | W | A | A | A | W | A | A | W | W | W | A | W | W | A | W | W | A | A | W | W | W | A | W | W | A | W | W | A | W | W | A | W | W | A | W | A | W | W |
| Chrl | 23.9 | A | A | A | W | W | A | A | W | A | A | A | A | W | A | NA | W | A | W | W | A | W | A | W | A | A | A | A | W | W | A | W | A | W | NA | A | A | W | W | A | W | A | W | A | A | W | A | W | A | W | A | A | A | A | A | W | A | A | A | W | A | A | W | W | W | A | W | W | A | W | W | A | A | W | W | W | A | W | W | A | W | W | A | W | W | A | W | W | A | W | A | W | W |
| Chrl | 24 | A | A | A | W | W | A | A | W | A | A | A | A | W | A | NA | W | A | W | W | A | W | A | W | A | A | A | A | W | W | A | W | A | W | NA | A | A | W | W | A | W | A | W | A | A | W | A | W | A | W | A | A | A | A | A | W | A | A | A | W | A | A | W | W | W | A | W | W | A | W | W | A | A | W | W | W | A | W | W | A | W | W | A | W | W | A | W | W | A | W | A | W | W |
| Chrl | 24.1 | A | A | A | W | W | A | A | W | A | A | A | A | W | A | A | W | A | W | W | A | W | A | W | A | A | A | A | W | W | A | W | A | W | W | A | A | W | W | A | W | A | W | A | A | W | A | W | A | W | A | A | A | A | A | W | A | A | A | W | A | A | W | W | W | A | W | W | A | W | W | A | A | W | W | W | A | W | W | A | W | W | A | W | W | A | W | W | A | W | A | W | W |
| Chrl | 24.2 | A | A | A | W | W | A | A | W | A | A | A | A | W | A | A | W | A | W | W | A | W | A | W | A | A | A | A | W | W | A | W | A | W | NA | A | A | W | W | A | W | A | W | A | A | W | A | W | A | W | A | A | A | A | A | W | A | A | A | W | A | A | W | W | W | A | W | W | A | W | W | A | A | W | W | W | A | W | W | A | W | W | A | W | W | A | W | W | A | W | A | W | W |
| Chrl | 24.3 | A | A | A | W | W | A | A | W | A | A | A | A | W | A | A | W | A | W | W | A | W | A | W | A | A | A | A | W | W | A | W | A | W | W | A | A | W | W | A | W | A | W | A | A | W | A | W | A | W | A | A | A | A | A | W | A | A | A | W | A | A | W | W | W | A | W | W | A | W | W | A | A | W | W | W | A | W | W | A | W | W | A | W | W | A | W | W | A | W | A | W | W |
| Chrl | 24.4 | A | A | A | W | W | A | A | W | A | A | A | A | W | A | A | W | A | W | W | A | W | A | W | A | A | A | A | W | W | A | W | A | W | W | A | A | W | W | A | W | A | W | A | A | W | A | W | A | W | A | A | A | A | A | W | A | A | A | W | A | A | W | W | W | A | W | W | A | W | W | A | A | W | W | W | A | W | W | A | W | W | A | W | W | A | W | W | A | W | A | W | W |
| Chrl | 24.5 | A | A | A | W | W | A | A | W | A | A | A | A | W | A | A | W | A | W | W | A | W | A | W | A | A | A | A | W | W | A | W | A | W | W | A | A | W | W | A | W | A | W | A | A | W | A | W | A | W | A | A | NA | A | A | W | A | A | A | W | A | A | W | W | W | A | W | W | A | W | W | A | A | W | W | W | A | W | W | A | W | W | A | W | W | A | W | W | A | W | A | W | W |
| Chrl | 24.6 | A | A | A | W | W | A | A | W | A | A | A | A | W | A | A | W | A | W | W | A | W | A | W | A | A | A | A | W | W | A | W | A | W | NA | A | A | W | W | A | W | A | W | A | A | W | A | W | A | W | A | A | A | A | A | W | A | A | A | W | A | A | W | W | W | A | W | W | A | W | W | A | A | W | W | W | A | W | W | A | W | W | A | W | W | A | W | W | A | W | A | W | W |
| Chrl | 24.7 | A | A | A | W | W | A | A | W | A | A | A | A | W | A | A | W | A | W | W | A | W | A | W | A | A | A | A | W | W | A | W | A | W | NA | A | A | W | W | A | W | A | W | A | A | W | A | W | A | W | NA | A | A | A | A | W | A | A | NA | W | A | A | W | W | W | NA | W | W | A | W | W | A | A | W | W | W | A | W | W | NA | W | W | A | W | W | A | W | W | A | W | A | W | W |
| Chrl | 24.8 | A | A | A | W | W | A | A | W | A | A | A | A | W | A | A | W | A | W | W | A | W | A | W | A | A | A | A | W | W | A | W | A | W | W | A | A | W | W | A | W | A | W | A | A | W | A | W | A | W | A | A | A | A | A | W | A | A | A | W | A | A | W | W | W | A | W | W | A | W | W | A | A | W | W | W | A | W | W | A | W | W | A | W | W | A | W | NA | A | W | A | W | W |
| Chrl | 24.9 | A | A | A | W | W | A | A | W | A | A | A | A | W | A | A | W | A | W | W | A | W | NA | W | A | A | A | A | W | W | A | W | A | W | W | A | A | W | W | A | W | A | W | A | A | W | A | W | A | W | NA | A | A | A | A | W | A | A | A | W | A | A | W | W | W | A | W | NA | A | W | W | A | A | W | W | W | A | W | W | A | W | W | A | W | W | A | W | W | A | W | A | W | W |
| Chrl | 25 | A | A | A | W | W | A | A | W | A | A | A | A | W | A | A | W | A | W | W | A | W | W | W | A | A | A | A | W | W | A | W | A | W | W | A | A | W | W | A | W | A | W | A | A | W | A | W | A | W | A | A | A | A | A | W | A | A | A | W | A | A | W | W | W | A | W | W | A | W | W | A | A | W | W | W | A | W | W | A | W | W | A | W | W | A | W | W | A | W | A | W | W |
| Chrl | 25.1 | A | A | A | W | W | A | A | W | A | A | A | A | W | A | A | W | A | W | W | A | W | W | W | A | A | A | A | W | W | A | W | A | W | W | A | A | W | W | A | W | A | W | A | A | W | A | W | A | W | A | A | A | A | A | W | A | A | A | W | A | A | W | W | W | NA | W | W | A | W | W | A | A | W | W | W | A | W | W | A | W | W | A | W | W | A | W | W | A | W | A | W | W |
| Chrl | 25.2 | A | A | A | W | W | A | A | W | A | A | A | A | W | NA | A | W | A | W | W | A | W | W | W | A | A | A | A | W | W | NA | W | A | W | NA | A | A | W | W | A | W | A | W | A | A | W | A | W | A | W | A | A | A | A | A | W | A | A | A | W | A | A | W | W | W | NA | W | W | A | W | W | A | A | W | W | W | NA | W | W | A | W | W | A | W | W | NA | W | W | A | W | A | W | W |
| Chrl | 25.3 | A | A | A | W | W | A | A | W | A | A | A | A | W | A | A | W | A | W | W | A | W | W | NA | A | A | A | A | NA | W | A | W | A | W | NA | A | A | W | W | A | W | A | W | A | A | W | A | W | A | W | A | A | A | A | A | W | A | A | A | W | A | A | W | W | W | NA | NA | NA | A | W | W | A | A | W | W | W | A | W | W | A | W | W | A | W | W | A | W | W | A | NA | A | W | W |
| Chrl | 25.4 | A | A | A | W | W | A | A | W | A | A | A | A | W | A | A | W | A | W | W | A | W | W | W | A | A | A | A | NA | W | A | W | A | W | W | A | A | W | W | A | W | A | W | A | A | W | A | W | A | W | A | A | A | A | A | W | A | A | A | W | A | A | W | W | W | A | W | NA | A | W | W | A | A | W | W | W | A | W | W | A | W | W | A | W | W | A | W | W | A | W | A | W | W |
| Chrl | 25.5 | A | A | A | W | W | A | A | W | A | A | A | A | W | A | A | W | A | W | W | A | W | W | NA | A | A | A | A | NA | W | A | W | A | W | W | A | A | W | W | A | W | A | W | A | A | W | A | W | A | W | A | A | A | A | A | W | A | A | A | W | A | A | W | W | W | A | W | NA | A | W | W | A | A | W | W | W | A | W | W | A | W | W | A | W | W | A | W | NA | A | W | A | W | W |
| Chrl | 25.6 | A | A | A | W | W | A | A | W | A | A | A | A | W | A | A | W | A | W | W | NA | W | W | W | A | A | A | A | W | W | A | W | A | W | W | A | A | W | W | A | W | A | W | A | A | W | A | W | A | W | NA | A | A | A | A | W | A | A | A | W | A | A | W | W | W | A | W | W | A | W | W | A | A | W | W | W | A | W | W | A | W | W | A | W | W | A | W | NA | A | W | A | W | W |
| Chrl | 25.7 | A | A | A | W | W | A | A | W | A | A | A | A | W | A | A | W | A | W | W | A | W | W | W | A | A | A | A | W | W | A | W | A | W | W | A | A | W | W | A | W | A | W | A | A | W | A | W | A | W | A | A | A | A | A | W | A | A | A | W | A | A | W | W | W | A | W | W | A | W | W | A | A | W | W | W | A | W | W | A | W | W | A | W | W | A | W | W | A | W | A | W | W |
| Chrl | 25.8 | A | A | A | W | W | A | A | W | A | A | A | A | W | A | A | W | A | W | W | A | W | W | W | A | A | A | A | W | W | A | W | A | W | W | A | A | W | W | A | W | A | W | A | A | W | A | W | A | W | A | A | A | A | A | W | A | A | A | W | A | A | W | W | W | A | W | W | A | W | W | A | A | W | W | W | A | W | W | A | W | W | A | W | W | A | W | W | A | W | A | W | W |
| Chrl | 25.9 | A | A | A | W | W | A | A | W | A | A | A | A | W | A | A | W | A | W | W | A | W | W | W | A | A | A | A | W | W | A | W | A | W | NA | A | A | W | W | A | W | A | W | A | A | W | A | W | A | W | A | A | A | A | A | W | A | A | A | W | A | A | W | W | W | A | W | W | A | W | W | A | A | W | W | W | A | W | W | A | W | W | A | W | W | A | W | W | A | W | A | W | W |
| Chrl | 26 | A | A | A | W | W | A | A | W | A | A | A | A | W | A | A | W | A | W | W | A | W | W | W | A | A | A | A | W | W | A | W | A | W | W | A | A | W | W | A | W | A | W | A | A | W | A | W | A | W | A | A | A | A | A | W | A | A | A | W | A | A | W | W | W | NA | W | W | A | W | W | A | A | W | W | W | A | W | W | A | W | W | A | W | W | A | W | W | A | W | A | W | W |
| Chrl | 26.1 | A | A | A | W | W | A | A | W | A | A | A | A | W | A | A | W | A | W | W | A | W | W | W | A | A | A | A | W | W | A | W | A | W | NA | A | A | W | W | A | W | A | W | A | A | W | A | W | A | W | A | A | A | A | A | W | A | A | A | W | A | A | W | W | W | A | W | W | A | W | W | A | A | W | W | W | A | W | W | A | W | W | A | W | W | A | W | W | A | W | A | W | W |
| Chrl | 26.2 | A | A | A | W | W | A | A | W | A | A | A | A | W | A | A | W | A | W | W | A | W | W | W | A | A | A | A | W | W | A | W | A | W | NA | A | A | W | W | A | W | A | W | A | A | W | A | W | A | W | A | A | A | A | A | W | A | A | A | W | A | A | W | W | W | A | W | W | A | W | W | A | A | W | W | W | A | W | W | A | W | W | A | W | W | A | W | W | A | W | A | W | W |
| Chrl | 26.3 | A | A | A | W | W | A | A | W | A | A | A | A | W | A | A | W | A | W | W | A | W | W | W | A | A | A | A | W | W | A | W | A | W | NA | A | A | W | W | A | W | A | W | A | A | W | A | W | A | W | A | A | A | A | A | W | A | A | A | W | A | A | W | W | W | A | W | W | A | W | W | A | A | W | W | W | A | W | W | A | W | W | A | W | W | A | W | W | A | W | A | W | W |
| Chrl | 26.4 | A | A | A | W | W | A | A | W | A | A | A | A | W | A | A | W | A | W | W | A | W | W | W | A | A | A | A | W | W | A | W | A | W | W | A | A | W | W | A | W | A | W | A | A | W | A | W | A | W | A | A | A | A | A | W | A | A | A | W | A | A | W | W | W | A | W | W | A | W | W | A | A | W | W | W | A | W | W | A | W | W | A | W | W | A | W | W | A | W | A | W | W |
| Chrl | 26.5 | A | A | A | W | W | A | A | W | A | A | A | A | W | A | A | W | A | W | W | A | W | W | W | A | A | A | A | W | W | A | W | A | W | W | A | A | W | W | A | W | A | W | A | A | W | A | W | A | W | A | A | A | A | A | W | A | A | A | W | A | A | W | W | W | A | W | W | A | W | W | A | A | W | W | W | A | W | W | A | W | W | A | W | W | A | W | W | A | W | A | W | W |
| Chrl | 26.6 | A | A | A | W | W | A | A | W | A | A | A | A | W | A | A | W | A | W | W | A | W | W | W | A | A | A | A | NA | W | A | W | A | W | W | A | A | W | W | A | W | A | W | A | A | W | A | W | A | W | NA | A | A | A | A | W | A | A | A | W | A | A | W | W | W | A | W | NA | A | W | W | A | A | W | W | W | A | W | W | A | W | W | A | W | W | A | W | NA | A | W | A | W | W |
| Chrl | 26.7 | A | A | A | W | W | A | A | W | A | A | A | A | W | A | A | W | A | W | W | A | W | W | W | A | A | A | A | W | W | A | W | A | W | NA | A | A | W | W | A | W | A | W | A | A | W | A | W | A | W | A | A | A | A | A | W | A | A | A | W | A | A | W | W | W | A | W | NA | A | W | W | A | A | W | W | W | A | W | W | A | W | W | A | W | W | A | W | W | A | W | A | W | W |
| Chrl | 26.8 | A | A | A | W | W | A | A | W | A | A | A | A | W | A | A | W | A | W | W | A | W | W | W | A | A | A | A | NA | W | A | W | A | W | NA | A | A | W | W | A | W | A | W | A | A | W | A | NA | A | W | A | A | A | A | A | W | A | A | A | W | A | A | W | W | W | A | W | NA | A | W | W | A | A | W | W | W | A | W | W | A | W | W | A | W | W | A | W | W | A | W | A | W | W |
| Chrl | 26.9 | A | A | A | W | W | A | A | W | A | A | A | A | W | A | A | W | A | W | W | A | W | W | W | A | A | A | A | NA | W | A | W | A | W | NA | A | A | W | W | A | W | A | W | A | A | W | A | W | A | W | A | A | A | A | A | W | A | A | A | W | A | A | W | W | W | A | W | W | A | W | W | A | A | W | W | W | A | W | W | A | W | W | A | W | W | A | W | NA | A | W | A | W | NA |
| Chrl | 27 | A | A | A | W | W | A | A | W | A | A | A | A | W | A | A | W | A | W | W | A | W | W | W | A | A | A | A | NA | W | A | W | A | W | W | A | A | W | W | A | W | A | W | A | A | W | A | W | A | W | A | A | A | A | A | W | A | A | A | W | A | A | W | W | W | A | W | W | A | W | W | A | A | W | W | W | A | W | W | A | W | W | A | W | W | A | W | W | A | W | A | W | W |

| | | | | | | | | | | | | | | | | | | | | | | | | | | | | | | | | | | | | | | | | | | | | | | | | | | | | | | | | | | | | | | | | | | | | | | | | | | | | | | | | | | | | | | | | | | | | | |
|---|---|---|---|---|---|---|---|---|---|---|---|---|---|---|---|---|---|---|---|---|---|---|---|---|---|---|---|---|---|---|---|---|---|---|---|---|---|---|---|---|---|---|---|---|---|---|---|---|---|---|---|---|---|---|---|---|---|---|---|---|---|---|---|---|---|---|---|---|---|---|---|---|---|---|---|---|---|---|---|---|---|---|---|---|---|---|---|---|---|---|---|---|---|
| Chrl | 27.1 | A | A | A | W | W | A | A | W | A | A | A | A | W | A | A | W | A | W | W | A | W | W | NA | A | A | A | A | NA | W | A | W | A | W | NA | A | A | W | W | A | W | A | W | A | A | W | A | W | A | W | A | A | A | A | A | W | A | A | A | W | A | A | W | W | W | A | W | W | A | W | W | A | A | W | W | W | A | W | W | A | W | W | A | W | W | A | W | W | A | W | A | W | W |
| Chrl | 27.2 | A | A | A | W | W | A | A | W | A | A | A | A | W | A | A | W | A | W | W | A | W | W | NA | A | A | A | A | W | W | A | W | A | W | W | A | A | W | W | A | W | A | W | A | A | W | A | W | A | W | A | A | A | A | A | W | A | A | A | W | A | A | W | W | W | A | W | NA | A | W | W | A | A | W | W | W | A | W | W | A | W | W | A | W | W | A | W | W | A | W | A | W | W |
| Chrl | 27.3 | A | A | A | W | W | A | A | W | A | A | A | A | W | A | A | W | A | W | W | A | W | W | W | A | A | A | A | W | W | A | W | A | W | W | A | A | W | W | A | W | A | W | A | A | W | A | W | A | W | A | A | A | A | A | W | A | A | A | W | A | A | W | W | W | A | W | W | A | W | W | A | A | W | W | W | A | W | W | A | W | W | A | W | W | A | W | W | A | W | A | W | NA |
| Chrl | 27.4 | A | A | A | W | W | A | A | W | A | A | A | A | W | A | A | W | NA | W | W | A | W | W | W | A | A | A | A | W | W | A | W | A | W | W | A | A | W | W | A | W | A | W | A | A | W | A | W | A | W | A | A | A | A | A | W | A | A | A | W | A | A | W | W | W | A | W | W | A | W | W | A | A | W | W | W | A | W | W | A | W | W | A | W | W | A | W | W | A | W | A | W | W |
| Chrl | 27.5 | A | A | A | W | W | A | NA | W | A | A | A | NA | W | A | A | W | A | W | W | NA | W | W | W | NA | NA | A | NA | W | W | A | W | A | W | W | A | A | W | W | A | W | A | W | A | A | W | A | W | NA | W | A | A | NA | NA | A | W | NA | A | NA | W | A | NA | W | W | W | NA | NA | W | A | W | W | NA | A | W | W | W | NA | W | W | A | W | W | A | W | W | NA | W | W | A | W | A | W | W |
| Chrl | 27.6 | A | A | A | W | W | A | A | W | A | A | A | A | W | A | A | W | A | W | W | A | W | W | W | A | A | A | A | W | W | A | W | A | W | NA | A | A | W | W | A | W | A | W | A | A | W | A | W | A | W | A | A | A | A | A | W | A | A | A | W | A | A | W | W | W | A | W | W | A | W | W | A | A | W | W | W | A | W | W | A | W | W | A | W | W | A | W | W | A | W | A | W | W |
| Chrl | 27.7 | A | A | A | W | W | A | A | W | A | A | A | A | W | A | A | W | A | W | W | A | W | W | NA | A | A | A | A | NA | W | A | W | A | W | NA | A | A | W | W | A | W | A | W | A | A | W | A | W | A | W | A | A | A | A | A | W | A | A | A | W | A | A | W | W | W | A | W | W | A | W | W | A | A | W | W | W | A | W | W | A | W | W | A | W | W | A | W | W | A | W | A | W | W |
| Chrl | 27.8 | A | A | A | W | W | A | A | W | A | A | A | A | W | A | A | W | A | W | W | A | W | W | W | A | A | A | A | W | W | A | W | A | W | NA | A | A | W | W | A | W | A | W | A | A | W | A | W | A | W | A | A | A | A | A | W | A | A | A | W | A | A | W | W | W | A | W | W | A | W | W | A | A | W | W | W | A | W | W | A | W | W | A | W | W | A | W | W | A | W | A | W | W |
| Chrl | 27.9 | A | A | A | W | W | A | A | W | A | A | A | A | W | A | A | W | A | W | W | A | W | W | NA | A | A | A | A | NA | W | A | W | A | W | W | A | A | W | W | A | W | A | W | A | A | W | A | W | A | W | A | A | A | A | A | W | A | A | A | W | A | A | W | W | W | A | W | NA | A | W | W | A | A | W | W | W | A | W | W | A | W | W | A | W | W | A | W | W | A | W | A | W | W |
| Chrl | 28 | A | A | A | W | W | A | A | W | A | A | A | A | W | A | A | W | A | W | W | A | W | W | W | A | NA | A | A | W | W | A | W | A | W | W | A | A | W | W | A | W | A | W | A | A | W | A | W | A | W | A | A | NA | A | A | W | A | A | A | W | A | A | W | W | W | A | W | W | A | W | W | A | A | W | W | W | A | W | W | A | W | W | A | W | W | NA | W | W | A | W | A | W | W |
| Chrl | 28.1 | A | A | A | W | W | A | A | W | A | A | A | A | W | A | A | W | NA | W | W | A | W | W | W | A | A | A | A | W | W | A | W | A | W | W | A | A | W | W | A | W | A | W | A | A | W | A | W | A | W | A | A | A | A | A | W | A | A | A | W | A | A | W | W | W | A | W | W | A | W | W | A | A | W | W | W | A | W | W | A | W | W | A | W | W | A | W | W | A | W | A | W | W |
| Chrl | 28.2 | A | A | A | W | W | A | A | W | A | A | A | A | W | A | NA | W | A | W | W | A | W | W | W | A | A | A | A | NA | W | A | W | A | W | NA | A | A | W | W | A | W | A | W | A | A | W | A | W | A | W | A | A | A | A | A | W | A | A | A | W | A | A | W | W | W | A | W | W | A | W | W | A | A | W | W | W | A | W | W | A | W | W | A | W | W | A | W | W | A | W | A | W | W |
| Chrl | 28.3 | A | A | A | W | W | A | A | W | A | A | A | A | W | A | A | W | A | W | W | A | W | W | W | A | A | A | A | W | W | A | W | A | W | NA | A | A | W | W | A | W | A | W | A | A | W | A | W | A | W | A | A | A | A | A | W | A | A | A | W | A | A | W | W | W | A | W | W | A | W | W | A | A | W | W | W | A | W | W | A | W | W | A | W | W | A | W | W | A | W | A | W | W |
| Chrl | 28.4 | A | A | A | W | W | A | A | W | A | A | A | A | W | A | A | W | A | W | W | A | W | W | W | A | A | A | A | NA | W | A | W | A | W | NA | A | A | W | W | A | W | A | W | A | A | W | A | W | A | W | A | A | A | A | A | W | A | A | A | W | A | A | W | W | W | A | W | W | A | W | W | A | A | W | W | W | A | W | W | A | W | W | A | W | W | A | W | W | A | W | A | W | W |
| Chrl | 28.5 | A | A | A | W | W | A | A | W | A | A | A | A | W | A | A | W | A | W | W | A | W | W | W | A | A | A | A | W | W | A | W | A | W | W | A | A | W | W | A | W | A | W | A | A | W | A | W | A | W | NA | A | A | A | A | W | A | A | A | W | A | A | W | W | W | NA | W | NA | A | W | W | A | A | NA | W | W | A | NA | W | A | W | W | A | W | W | A | NA | NA | A | W | A | W | W |
| Chrl | 28.6 | A | A | A | W | W | A | A | W | A | A | A | A | W | A | A | W | A | W | W | A | W | W | W | A | A | A | A | NA | W | A | W | A | W | W | A | A | W | W | A | W | A | W | A | A | W | A | W | A | W | A | A | A | A | A | W | A | A | A | W | A | A | W | W | W | A | W | W | A | W | W | A | A | W | W | W | A | W | W | A | W | W | A | W | W | A | W | W | A | W | A | W | W |
| Chrl | 28.7 | A | A | A | W | W | A | A | W | A | A | A | A | W | A | A | W | A | W | W | A | W | W | W | A | A | A | A | W | W | A | W | A | W | NA | A | A | W | W | A | W | A | W | A | A | W | A | W | A | W | A | A | A | A | A | W | A | A | A | W | A | A | W | W | W | A | W | W | A | W | W | A | A | W | W | W | A | W | W | A | W | W | A | W | W | A | W | W | A | W | A | W | W |
| Chrl | 28.8 | A | A | A | W | W | A | A | W | A | A | A | A | W | A | NA | W | A | W | W | A | W | W | W | A | A | A | A | W | W | A | W | A | W | W | A | A | W | W | A | W | A | W | A | A | W | A | W | A | W | A | A | A | A | A | W | A | A | A | W | A | A | W | W | W | A | W | W | A | W | W | NA | A | W | W | W | A | W | W | A | W | W | A | W | W | A | W | W | A | W | A | W | W |
| Chrl | 28.9 | A | A | A | W | W | A | A | W | A | A | A | A | W | A | A | W | A | W | W | A | W | W | W | A | A | A | A | W | W | A | W | A | W | W | A | A | W | W | A | W | A | W | A | A | W | A | W | A | W | A | A | A | A | A | W | A | A | A | W | A | A | W | W | W | A | W | NA | A | W | W | A | A | W | W | W | A | W | W | A | W | W | A | W | W | A | W | W | A | W | A | W | W |
| Chrl | 29 | A | A | A | W | W | A | A | W | A | A | A | A | W | A | A | W | A | W | W | A | W | W | W | A | A | A | A | W | W | A | W | A | W | W | A | A | W | W | A | W | A | W | A | A | W | A | W | A | W | NA | A | A | A | A | W | A | A | A | W | A | A | W | W | W | A | W | W | A | W | W | A | A | W | W | W | A | W | W | A | W | W | A | W | W | A | W | W | A | W | A | W | W |
| Chrl | 29.1 | A | A | A | W | W | NA | A | W | A | A | A | A | W | A | NA | W | A | W | W | A | W | W | W | A | A | A | A | W | W | A | W | A | W | W | A | A | W | W | A | W | A | W | A | A | W | NA | W | A | W | A | A | A | A | A | W | A | A | A | W | A | A | W | W | W | NA | W | W | A | W | W | A | A | W | W | W | A | W | W | A | W | W | A | W | W | A | W | W | A | W | A | W | W |
| Chrl | 29.2 | A | A | A | W | W | A | A | W | A | A | A | A | W | A | A | W | A | W | W | A | W | W | W | A | A | A | A | W | W | A | W | A | W | W | A | A | W | W | A | W | A | W | A | A | W | A | W | A | W | A | A | A | A | A | W | A | A | A | W | A | A | W | W | W | A | W | W | A | W | W | A | A | W | W | W | A | W | W | A | W | W | A | W | W | A | W | W | A | W | A | W | W |
| Chrl | 29.3 | A | A | A | W | W | A | A | W | A | A | A | A | W | A | A | W | A | W | W | A | W | W | W | A | A | A | A | W | W | A | W | A | W | W | A | A | W | W | A | W | A | W | A | A | W | A | W | A | W | A | A | A | A | A | W | A | A | A | W | A | A | W | W | W | A | W | W | A | W | W | A | A | W | W | W | A | W | W | A | W | W | A | W | W | A | W | W | A | W | A | W | W |
| Chrl | 29.4 | A | A | A | W | W | A | A | W | A | A | A | A | W | A | A | W | A | W | W | A | W | W | W | A | A | A | A | W | W | A | W | A | W | W | A | A | W | W | A | W | A | W | A | A | W | A | W | A | W | A | A | A | A | A | W | A | A | A | W | A | A | W | W | W | A | W | W | A | W | W | A | A | W | W | W | A | W | W | A | W | W | A | W | W | A | W | W | A | W | A | W | W |
| Chrl | 29.5 | A | A | A | W | W | A | A | W | A | A | A | A | W | A | A | W | A | W | W | A | W | W | NA | A | A | A | A | W | W | A | W | A | W | W | A | A | W | W | A | W | A | W | A | A | W | A | W | A | W | A | A | A | A | A | W | A | A | A | W | A | A | W | W | W | A | W | W | A | W | W | A | A | W | W | W | A | W | W | A | W | W | A | W | W | A | W | W | A | W | A | W | W |
| Chrl | 29.6 | A | A | A | W | W | A | A | W | A | A | A | A | W | A | A | W | A | W | W | A | W | W | NA | A | A | A | A | W | W | A | W | A | W | W | A | A | W | W | A | W | A | W | A | A | W | A | W | A | W | A | A | A | A | A | W | A | A | A | W | A | A | W | W | W | A | W | W | A | W | W | A | A | W | W | W | A | W | W | A | W | W | A | W | W | A | W | W | A | W | A | W | W |
| Chrl | 29.7 | A | A | A | W | W | A | A | W | A | A | A | A | W | A | A | W | A | W | W | A | W | W | NA | A | A | A | A | NA | W | A | W | A | W | W | A | A | W | W | A | W | A | W | A | A | W | A | NA | A | W | A | A | A | A | A | W | A | A | A | W | A | A | W | W | W | NA | W | NA | A | W | W | A | A | NA | W | W | A | W | W | A | W | W | A | W | W | A | W | NA | A | W | A | W | NA |
| Chrl | 29.8 | A | A | A | W | W | A | A | W | A | A | A | A | W | A | A | W | A | W | W | A | W | W | NA | A | A | A | A | NA | W | A | W | A | W | W | A | A | W | W | A | W | A | W | A | A | W | A | NA | A | W | A | A | A | A | A | W | A | A | A | W | A | A | W | W | W | NA | NA | NA | A | W | W | A | A | NA | W | W | A | NA | W | A | W | W | A | W | W | A | W | W | A | W | A | W | W |
| Chrl | 29.9 | A | A | A | W | W | A | A | W | A | A | A | A | W | A | A | W | A | W | W | A | W | W | W | A | A | A | A | NA | W | A | W | A | W | W | A | A | W | W | A | W | A | W | A | A | W | A | NA | A | W | A | A | A | A | A | W | A | A | A | W | A | A | W | W | W | A | NA | W | A | W | W | A | A | NA | W | W | A | W | W | A | W | W | A | W | W | A | W | W | A | W | A | W | W |
| Chrl | 30 | A | A | A | W | W | A | A | W | A | A | A | A | W | A | A | W | A | W | W | A | W | W | W | A | A | A | A | W | W | A | W | A | W | W | A | A | W | W | A | W | A | W | A | A | W | A | W | A | W | A | A | A | A | A | W | A | A | A | W | A | A | W | W | W | A | W | W | A | W | W | A | A | W | W | W | A | W | W | A | W | W | A | W | W | A | W | W | A | W | A | W | W |
| Chrl | 30.1 | A | A | A | W | W | A | A | W | A | A | A | A | W | A | A | W | A | W | W | A | W | W | W | A | A | A | A | NA | W | A | W | A | W | W | A | A | W | W | A | W | A | W | A | A | W | A | W | A | W | A | A | A | A | A | W | A | A | A | W | A | A | W | W | W | A | W | NA | A | W | W | A | A | W | W | W | A | W | W | A | W | W | A | W | W | A | W | W | A | W | A | W | W |
| Chrl | 30.2 | A | A | A | W | W | A | A | W | A | A | A | A | W | A | A | W | A | W | W | A | W | W | W | A | A | A | A | W | W | A | W | A | W | W | A | A | W | W | A | W | A | W | A | A | W | A | W | A | W | NA | A | A | A | A | W | A | A | A | W | A | A | W | W | W | A | W | W | A | W | W | A | A | W | W | W | A | W | W | A | W | W | A | W | W | A | W | W | A | W | A | W | W |
| Chrl | 30.3 | NA | A | A | W | W | A | A | W | A | A | A | A | W | A | A | W | A | W | W | A | W | W | W | A | A | A | A | W | W | A | W | A | W | NA | A | A | W | W | A | W | A | W | A | A | W | A | W | A | W | A | A | A | A | A | W | A | A | A | W | A | A | W | W | W | A | W | W | A | W | W | A | A | W | W | W | A | W | W | A | W | W | A | W | W | A | W | W | A | W | A | W | W |
| Chrl | 30.4 | W | A | A | W | W | A | A | W | A | A | A | A | W | A | A | W | A | W | W | A | W | W | W | A | A | A | A | W | W | A | W | A | W | W | A | A | W | W | A | W | A | W | A | A | W | A | W | A | W | A | A | A | A | A | W | A | A | A | W | A | A | W | W | W | A | W | W | A | W | W | A | A | W | W | W | A | W | W | A | W | W | A | W | W | A | W | W | A | W | A | W | W |
| Chrl | 30.5 | W | A | A | W | W | A | A | W | A | A | A | A | W | A | A | W | A | W | W | A | W | W | NA | A | A | A | A | W | W | A | W | A | W | NA | A | A | W | W | A | W | A | W | A | A | W | A | W | A | W | NA | A | A | A | A | W | A | A | A | W | A | A | W | W | W | A | NA | NA | A | W | W | A | A | W | W | W | A | W | W | A | W | W | A | W | W | A | W | W | A | W | A | W | W |
| Chrl | 30.6 | W | A | A | W | W | A | A | W | A | A | A | A | W | A | A | W | A | W | W | A | W | W | W | A | A | A | A | W | W | A | W | A | W | NA | A | A | W | W | A | W | A | W | A | A | W | A | W | A | W | A | A | A | A | A | W | A | A | A | W | A | A | W | W | W | A | W | NA | A | W | W | A | A | W | W | W | A | W | W | A | W | W | A | W | W | A | W | W | A | W | A | W | W |
| Chrl | 30.7 | W | A | A | W | W | A | A | W | A | A | A | A | W | A | A | W | A | W | W | A | W | W | W | A | A | A | A | W | W | A | W | A | W | W | A | A | W | W | A | W | A | W | A | A | W | A | W | A | W | A | A | A | A | A | W | A | A | A | W | A | A | W | W | W | A | W | W | A | W | W | A | A | W | W | W | A | W | W | A | W | W | A | W | W | A | W | W | A | W | A | W | W |
| Chrl | 30.8 | W | A | A | W | W | A | A | W | A | A | A | A | W | A | A | W | A | W | W | A | W | W | NA | A | A | A | A | W | W | A | W | A | W | NA | A | A | W | W | A | W | A | W | A | A | W | A | W | A | W | A | A | A | A | A | W | A | A | A | W | A | A | W | W | W | A | W | NA | A | W | W | A | A | W | W | W | A | W | W | A | W | W | A | W | W | A | W | W | A | W | A | W | W |
| Chrl | 30.9 | W | A | A | W | W | A | A | W | A | A | A | A | W | A | A | W | A | W | W | A | W | W | W | A | A | A | A | W | W | A | W | A | W | W | A | A | W | W | A | W | A | W | A | A | W | A | W | A | W | NA | A | A | A | A | W | A | A | A | W | A | A | W | W | W | A | W | W | A | W | W | A | A | W | W | W | A | W | W | A | W | W | A | W | W | A | W | W | A | W | A | W | W |
| Chrl | 31 | W | A | A | W | W | A | A | W | A | A | A | A | W | A | A | W | A | W | A | A | W | W | W | A | A | A | A | W | W | A | W | A | W | W | A | A | W | W | A | W | A | W | A | A | W | A | NA | A | W | A | A | A | A | A | W | A | A | A | W | A | A | W | W | W | A | W | W | A | W | W | A | A | W | W | W | A | W | W | A | W | W | A | W | W | A | W | W | A | W | A | W | W |
| Chrl | 31.1 | W | A | A | W | W | A | A | W | A | A | A | A | W | A | A | W | A | W | A | A | W | W | W | A | A | A | A | W | W | A | W | A | W | W | A | A | W | W | A | W | A | W | A | A | W | A | W | A | W | A | NA | A | A | A | W | A | A | A | W | A | A | W | W | W | A | W | W | A | W | W | A | A | W | W | W | A | W | W | A | W | W | A | W | W | A | W | W | A | W | A | W | W |
| Chrl | 31.2 | W | A | A | W | W | A | A | W | A | A | A | A | W | A | A | W | A | W | A | A | W | W | W | A | A | A | A | NA | W | A | W | A | W | NA | A | A | W | W | A | W | A | W | A | A | W | A | W | A | W | A | A | A | A | A | W | A | A | A | W | A | A | W | W | W | A | W | W | A | W | W | A | A | W | W | W | A | W | W | A | W | W | A | W | W | A | W | W | A | W | A | W | W |
| Chrl | 31.3 | W | A | A | W | W | A | A | W | A | A | A | A | W | A | A | W | A | W | A | A | W | W | NA | A | A | A | A | W | W | A | W | A | W | W | A | A | W | W | A | W | A | W | A | A | W | A | W | A | W | A | A | A | A | A | W | A | A | A | W | A | A | W | W | W | A | W | W | A | W | W | A | A | NA | W | W | A | W | W | A | W | W | A | W | W | A | W | W | A | W | A | W | W |
| Chrl | 31.4 | W | A | A | W | W | A | A | W | A | A | A | A | W | A | A | W | A | W | A | A | W | W | NA | A | A | A | A | W | W | A | W | A | W | W | A | A | W | W | A | W | A | W | A | A | W | A | NA | A | W | A | A | A | A | A | W | A | A | A | W | A | A | W | W | W | NA | W | NA | A | W | W | A | A | NA | W | W | NA | NA | W | A | W | W | A | W | W | A | W | W | A | W | A | W | W |
| Chrl | 31.5 | W | A | A | W | W | A | A | W | A | A | A | A | W | A | A | W | A | W | A | A | W | W | NA | A | A | A | A | W | W | A | W | A | W | W | A | A | W | W | A | W | A | W | A | A | W | A | W | A | W | A | A | A | A | A | W | A | A | A | W | A | A | W | W | W | A | W | NA | A | W | W | A | A | W | W | W | A | W | W | A | W | W | A | W | W | A | W | W | A | W | A | W | W |
| Chrl | 31.6 | W | A | A | W | W | A | A | W | A | A | A | A | W | A | A | W | A | W | A | A | W | W | W | A | A | A | A | W | W | A | W | A | W | W | A | A | W | W | A | W | A | W | A | A | W | A | W | A | W | A | A | A | A | A | W | A | A | A | W | A | A | W | W | W | A | W | NA | A | W | W | A | A | W | W | W | A | W | W | A | W | W | A | W | W | A | W | W | A | W | A | W | W |
| Chrl | 31.7 | W | A | A | W | W | A | A | W | A | A | A | A | W | A | A | W | A | W | A | A | W | W | W | A | A | A | A | W | W | A | W | A | W | NA | A | A | W | W | A | W | A | W | A | A | W | A | W | A | W | A | A | A | A | A | W | A | A | A | W | A | A | W | W | W | A | W | W | A | W | W | A | A | W | W | W | A | W | W | A | W | W | A | W | W | A | W | W | A | W | A | W | W |
| Chrl | 31.8 | W | A | A | W | W | A | A | W | A | A | A | A | W | A | A | W | A | W | A | A | W | W | W | A | A | A | A | W | W | A | W | A | W | W | A | A | W | W | A | W | A | W | A | A | W | A | W | A | W | A | A | A | A | A | W | A | A | A | W | A | A | W | W | W | A | W | W | A | W | W | A | A | W | W | W | A | W | W | A | W | W | A | W | W | A | W | W | A | W | A | W | W |
| Chrl | 31.9 | W | A | A | W | W | A | A | W | A | A | A | A | W | A | A | W | A | W | A | A | W | W | W | A | A | A | A | W | W | A | W | A | W | W | A | A | W | W | A | W | A | W | A | A | W | A | W | A | W | A | A | A | A | A | W | A | A | A | W | A | A | W | W | W | A | W | W | A | W | W | A | A | W | W | W | A | W | W | A | W | W | A | W | W | A | W | NA | A | W | A | W | W |
| Chrl | 32 | W | A | A | W | W | A | A | W | A | A | A | A | W | A | A | W | A | W | A | NA | W | W | W | A | A | A | A | W | W | A | W | A | W | W | A | A | W | W | A | W | A | W | A | A | W | A | W | A | W | NA | A | A | A | A | W | A | A | A | W | A | A | W | W | W | NA | W | W | A | W | W | NA | A | W | W | W | A | W | W | A | W | W | A | W | W | A | W | W | A | W | A | W | W |
| Chrl | 32.1 | W | A | A | W | W | A | A | W | A | A | A | A | W | A | A | W | A | W | A | A | W | W | W | A | A | A | A | NA | W | A | W | A | W | NA | A | A | W | W | A | W | A | W | A | A | W | A | W | A | W | A | A | A | A | A | W | A | A | A | W | A | A | W | W | W | A | W | NA | A | W | W | A | A | W | W | W | A | W | W | A | W | W | A | W | W | A | W | W | A | W | A | W | W |
| Chrl | 32.2 | W | A | A | W | W | A | A | W | A | A | A | A | W | A | A | W | A | W | A | A | W | W | W | A | A | A | A | W | W | A | W | A | W | W | A | A | W | W | A | W | A | W | A | A | W | A | W | A | W | A | A | A | A | A | W | A | A | A | W | A | A | W | W | W | A | W | W | A | W | W | A | A | W | W | W | A | W | W | A | W | W | A | W | W | NA | W | W | A | W | A | W | W |
| Chrl | 32.3 | W | A | A | W | W | A | A | W | A | A | A | A | W | A | A | W | A | W | A | A | W | W | NA | A | A | A | A | W | W | A | W | A | W | W | A | A | W | W | A | W | A | W | A | A | W | A | W | A | W | A | A | A | A | A | W | A | A | A | W | A | A | W | W | W | A | W | W | A | W | W | A | A | W | W | W | A | W | W | A | W | W | A | W | W | A | W | W | A | W | A | W | W |
| Chrl | 32.4 | W | A | A | W | W | A | A | W | A | A | A | A | W | A | A | W | A | W | A | A | W | W | NA | A | A | A | A | NA | W | A | W | A | W | W | A | A | W | W | A | W | A | W | A | A | W | A | W | A | W | A | A | A | A | A | W | A | A | A | W | A | A | W | W | W | A | W | NA | A | W | W | A | A | W | W | W | A | W | W | A | W | W | A | W | W | A | W | W | A | W | A | W | W |
| Chrl | 32.5 | W | A | A | W | W | A | A | W | A | A | A | A | W | A | A | W | A | W | A | A | W | W | W | A | A | A | A | W | W | A | W | A | W | A | A | A | W | W | A | W | A | W | A | A | W | A | W | A | W | A | A | A | A | A | W | A | A | A | W | A | A | W | W | W | A | W | W | A | W | W | A | A | W | W | W | A | W | W | A | W | W | A | W | W | A | W | W | A | W | A | W | W |
| Chrl | 32.6 | W | A | A | W | W | A | A | W | A | A | A | A | W | A | NA | W | A | W | A | A | W | W | W | A | A | A | NA | W | W | A | W | A | W | A | A | A | W | W | A | W | A | W | A | A | W | A | W | A | W | A | A | A | A | A | W | A | A | A | W | A | A | W | W | W | A | W | W | A | W | W | A | A | W | W | W | A | W | W | A | W | W | A | W | W | A | W | W | A | W | A | W | W |
| Chrl | 32.7 | W | A | A | W | W | A | A | W | A | A | A | A | W | A | A | W | A | W | A | A | W | W | W | A | A | A | NA | W | W | A | W | A | W | A | A | A | W | W | A | W | A | W | A | A | W | A | NA | A | W | A | A | A | A | A | W | A | A | A | W | A | A | W | W | W | A | W | W | A | W | W | A | A | W | W | W | A | W | W | A | W | W | A | W | W | A | W | W | A | W | A | W | W |
| Chrl | 32.8 | W | A | A | W | W | A | A | W | A | A | A | A | W | A | A | W | A | W | A | A | W | W | W | A | A | A | W | NA | W | A | W | A | W | A | A | A | W | NA | A | W | A | W | A | A | W | A | W | A | W | A | A | A | A | A | W | A | A | A | W | A | A | W | W | W | A | W | W | A | W | W | A | A | NA | W | W | NA | NA | W | A | W | W | A | W | W | NA | NA | W | A | NA | A | W | NA |
| Chrl | 32.9 | W | A | A | W | W | A | A | W | A | A | A | A | W | NA | A | W | A | W | A | A | W | W | W | A | A | A | W | NA | W | A | W | A | W | A | A | A | W | W | A | W | A | W | A | A | W | A | W | A | W | A | A | A | A | A | W | A | A | A | W | A | A | W | W | W | NA | W | W | A | W | W | A | A | W | W | W | A | W | W | A | W | W | A | W | W | A | W | W | A | W | A | W | W |
| Chrl | 33 | W | A | A | W | W | A | A | W | A | A | A | A | W | A | A | W | A | W | A | A | W | W | W | A | A | A | W | W | W | A | W | A | W | A | A | A | W | W | A | W | A | W | A | A | W | A | W | A | W | A | A | A | A | A | W | A | A | A | W | A | A | W | W | W | A | W | W | A | W | W | A | A | W | W | W | A | W | W | A | W | W | W | W | W | A | W | W | A | W | A | W | W |
| Chrl | 33.1 | W | A | A | W | W | A | A | W | A | A | A | A | W | A | NA | W | A | W | A | A | W | W | W | A | A | A | W | W | W | A | W | A | W | A | A | A | W | W | A | W | A | W | A | A | W | A | W | A | W | NA | A | A | A | A | W | A | A | A | W | A | A | W | W | W | A | W | W | A | W | W | A | A | W | W | W | A | W | W | A | W | W | W | W | W | A | W | W | A | W | A | W | W |
| Chrl | 33.2 | W | A | A | W | W | A | A | W | A | A | A | A | W | A | A | W | A | W | A | A | W | W | W | A | A | A | W | W | W | A | W | A | W | A | A | A | W | W | A | W | A | W | A | A | W | A | W | A | W | A | A | A | A | A | W | A | A | A | W | A | A | W | W | W | A | W | NA | A | W | W | A | A | W | W | W | A | W | W | A | W | W | W | W | W | A | W | W | A | W | A | W | W |
| Chrl | 33.3 | W | A | A | W | W | A | A | W | A | A | A | A | W | A | A | W | A | W | A | A | W | W | W | A | A | A | W | W | W | A | W | A | W | A | A | A | W | W | A | W | A | W | A | A | W | A | W | A | W | A | A | A | A | A | W | A | A | A | W | A | A | W | W | W | A | W | W | A | W | W | A | A | W | W | W | A | W | W | A | W | W | W | W | W | A | W | W | A | W | A | W | W |
| Chrl | 33.4 | W | A | A | W | W | A | A | W | A | A | A | A | W | A | NA | W | A | W | A | A | W | W | NA | A | A | A | W | W | W | A | W | A | W | A | A | A | W | W | A | W | A | W | A | A | W | A | W | A | W | A | A | A | A | A | W | A | A | A | W | A | A | W | W | W | A | W | W | A | W | W | A | A | W | W | W | A | W | NA | A | W | W | W | W | W | NA | W | W | A | W | A | W | W |
| Chrl | 33.5 | W | A | A | W | W | A | A | W | A | A | A | A | W | A | A | W | A | W | A | A | W | W | W | A | A | A | W | W | W | A | W | A | W | A | A | A | W | W | A | W | A | W | A | A | W | A | W | A | W | A | A | A | A | A | W | A | A | A | W | A | A | W | W | W | A | W | W | A | W | W | A | A | W | W | W | A | W | W | A | W | W | W | W | W | A | W | W | A | W | A | W | W |
| Chrl | 33.6 | W | A | A | W | W | A | A | W | A | A | A | A | W | A | A | W | A | W | A | A | W | W | NA | A | A | A | W | W | W | A | W | A | W | A | A | A | W | W | A | W | A | W | A | A | W | A | W | A | W | A | A | A | A | A | W | A | A | A | W | A | A | W | W | W | A | W | W | A | W | W | A | A | W | W | W | A | W | W | A | W | W | W | W | W | A | W | W | A | W | A | W | W |
| Chrl | 33.7 | W | A | A | W | W | A | A | W | A | A | A | A | W | A | A | W | A | W | A | A | W | W | W | A | A | A | W | W | W | A | W | A | W | A | A | A | W | W | A | W | A | W | A | A | W | A | W | A | W | A | A | A | A | A | W | A | A | A | W | A | A | W | W | W | A | W | W | A | W | W | A | A | W | W | W | A | W | W | A | W | W | W | W | W | A | W | W | A | W | A | W | W |
| Chrl | 33.8 | W | A | A | W | W | A | A | W | A | A | A | A | W | A | A | W | A | W | A | A | W | W | W | A | A | A | W | W | W | A | W | A | W | A | A | A | W | W | A | W | A | W | A | A | W | A | W | A | W | A | A | A | A | A | W | A | A | A | W | A | A | W | W | W | A | W | W | A | W | W | A | A | W | W | W | A | W | W | A | W | W | W | W | W | A | W | W | A | W | A | W | W |
| Chrl | 33.9 | W | A | A | W | W | A | A | W | A | A | A | A | W | A | A | W | A | W | A | A | W | W | W | A | A | A | W | W | W | A | W | A | W | A | A | A | W | W | A | W | A | W | A | A | W | A | W | A | W | A | A | A | A | A | W | A | A | A | W | A | A | W | W | W | A | W | W | A | W | W | A | A | W | W | W | A | W | W | A | W | W | W | W | W | A | W | W | A | W | A | W | W |
| Chrl | 34 | W | A | A | W | W | A | A | W | A | A | A | A | W | A | A | W | A | W | A | A | W | W | W | A | A | A | W | W | W | A | W | A | W | A | A | A | W | W | A | W | A | W | A | A | W | A | W | A | W | A | A | A | A | A | W | A | A | A | W | A | A | W | W | W | A | W | W | A | W | W | A | A | W | W | W | A | W | W | A | W | W | W | W | W | A | W | W | A | W | A | W | W |
| Chrl | 34.1 | W | A | A | W | W | A | A | W | A | A | A | A | W | A | A | W | A | W | A | A | W | W | NA | A | A | A | W | W | W | A | W | A | W | A | A | A | W | W | A | W | A | W | A | A | W | A | W | A | W | A | A | A | A | A | W | A | A | A | W | A | A | W | W | W | A | W | NA | A | W | W | A | A | W | W | W | A | W | W | A | W | W | W | W | W | A | W | W | A | W | A | W | W |
| Chrl | 34.2 | W | A | A | W | W | A | A | W | A | A | A | A | W | A | A | W | A | W | A | A | W | W | W | A | A | A | W | W | W | A | W | A | W | A | A | A | W | W | A | W | A | W | A | A | W | A | W | A | W | A | A | A | A | A | W | A | A | A | W | A | A | W | W | W | A | W | W | A | W | W | A | A | W | W | W | A | W | W | A | W | W | W | W | W | A | W | W | A | W | A | W | W |
| Chrl | 34.3 | W | A | A | W | W | A | A | W | A | A | A | A | W | A | A | W | A | W | A | A | W | W | W | A | A | A | W | W | W | A | W | A | W | A | A | A | W | W | A | W | A | W | A | A | W | NA | W | A | W | A | A | A | A | A | W | A | A | A | W | A | A | W | W | W | A | W | W | A | NA | W | A | A | W | W | NA | A | A | W | A | W | W | W | W | W | A | W | W | A | W | A | W | W |
| Chrl | 34.4 | W | A | A | W | W | A | A | W | A | A | A | A | W | A | A | W | A | W | A | A | W | W | W | A | A | A | W | W | W | A | W | A | W | NA | A | A | W | W | A | W | A | W | A | A | W | W | W | A | W | A | A | A | A | A | W | A | A | A | W | A | A | W | W | W | A | W | W | A | W | W | A | A | W | W | W | A | A | W | A | W | W | W | W | W | A | W | W | A | W | A | W | W |
| Chrl | 34.5 | W | A | A | W | W | A | A | W | A | A | A | A | W | A | A | W | A | W | A | A | W | W | W | A | A | A | W | W | W | A | W | A | W | A | A | A | W | W | A | W | A | W | A | A | W | W | W | A | W | A | A | A | A | A | W | A | A | A | W | A | A | W | W | W | NA | W | W | A | W | W | A | A | NA | W | W | A | A | W | A | W | W | W | W | W | A | W | W | A | W | A | W | W |
| Chrl | 34.6 | W | A | A | W | W | A | A | W | A | A | A | A | W | A | A | W | A | W | A | A | W | W | W | A | A | A | W | W | W | A | W | A | W | A | A | A | W | W | A | W | A | W | A | A | W | W | W | A | W | A | A | A | A | A | W | A | A | A | W | A | A | W | W | W | A | W | W | A | W | W | A | A | W | W | W | A | A | W | A | W | W | W | W | W | A | W | W | A | W | A | W | W |
| Chrl | 34.7 | W | A | A | W | W | A | NA | W | A | W | A | A | W | A | A | W | A | W | A | NA | W | W | NA | A | A | A | W | W | W | A | NA | A | W | A | A | A | W | NA | A | W | A | W | A | A | W | W | W | A | W | A | A | A | A | A | W | A | A | A | W | A | A | W | W | W | NA | W | NA | A | NA | W | A | A | NA | NA | W | A | A | NA | A | W | NA | W | W | W | A | W | W | A | W | A | W | NA |
| Chrl | 34.8 | W | A | A | A | W | A | A | W | A | W | A | A | W | A | A | W | A | W | A | A | W | W | W | A | A | A | W | W | W | A | W | A | W | A | A | A | W | W | A | W | A | W | A | A | W | W | W | A | W | A | A | A | A | A | W | A | A | A | W | A | A | W | W | W | A | W | W | A | W | W | A | A | W | A | W | NA | NA | NA | A | W | A | W | W | W | A | NA | W | A | W | A | W | W |
| Chrl | 34.9 | W | A | A | A | W | A | A | W | A | W | A | A | W | A | NA | W | A | W | A | A | W | W | W | A | A | A | W | W | W | A | W | A | W | A | A | A | W | W | A | W | A | W | A | A | W | W | W | A | W | A | A | A | A | A | W | A | A | A | W | A | A | W | W | W | A | W | W | A | W | W | A | A | W | A | W | A | A | W | A | W | A | W | W | W | A | W | W | A | W | A | W | W |
| Chrl | 35 | W | A | A | A | W | A | A | W | A | W | A | A | W | A | A | W | A | W | A | A | W | W | W | A | A | A | W | W | W | A | W | A | W | A | A | A | W | W | A | W | A | W | A | A | W | W | W | A | W | A | A | A | A | A | W | A | A | A | W | A | A | W | W | W | A | W | W | A | W | W | A | A | W | A | W | A | A | W | A | W | A | W | W | W | A | W | W | A | W | A | W | W |
| Chrl | 35.1 | W | A | A | A | W | A | A | W | A | W | A | A | W | A | A | W | A | W | A | A | W | W | W | A | A | A | W | W | W | A | W | A | W | A | A | A | W | W | A | W | A | W | A | A | W | W | W | A | W | A | A | A | A | A | W | A | A | A | W | A | A | W | W | W | A | W | W | A | W | W | A | A | W | A | W | A | A | W | A | W | A | W | W | W | A | W | NA | A | W | A | W | W |
| Chrl | 35.2 | W | A | A | A | W | A | NA | W | A | W | A | A | W | A | A | W | A | W | A | A | W | W | NA | A | A | A | W | W | W | A | W | A | W | A | A | A | W | W | A | W | A | W | A | A | W | W | W | A | W | A | A | A | A | A | W | A | A | A | W | A | A | W | W | W | NA | W | W | A | W | W | A | A | NA | A | NA | A | NA | W | A | W | A | W | W | W | A | W | W | A | W | A | W | W |
| Chrl | 35.3 | W | A | A | A | W | A | A | W | A | W | A | A | W | A | A | A | A | W | A | A | NA | W | NA | A | A | A | W | W | W | A | W | A | W | A | A | A | W | W | A | W | A | W | A | A | W | W | W | A | W | A | A | A | A | A | W | A | A | A | W | A | A | W | W | W | A | W | W | A | NA | W | A | A | NA | A | W | A | A | NA | A | W | A | W | W | W | A | W | NA | A | W | A | W | A |
| Chrl | 35.4 | W | A | A | A | W | A | A | W | A | W | A | A | W | A | A | A | A | W | A | A | W | W | NA | A | A | A | W | W | W | A | W | A | W | A | A | A | W | W | A | W | A | W | A | A | W | W | W | A | W | A | A | A | A | A | W | A | A | A | W | A | W | W | W | A | A | W | W | A | W | W | A | A | W | A | W | A | A | NA | W | W | A | W | W | W | A | W | W | A | W | A | W | A |
| Chrl | 35.5 | W | A | A | A | W | A | A | W | A | W | A | A | W | A | A | A | A | W | A | A | W | W | W | A | A | A | W | W | W | A | W | A | W | A | A | A | W | W | A | W | A | W | A | A | W | W | W | A | W | A | A | A | A | A | W | A | A | A | W | A | W | W | W | A | A | W | W | A | W | W | A | A | W | A | W | A | A | W | W | W | A | W | W | W | A | W | W | A | W | A | W | A |
| Chrl | 35.6 | W | A | A | A | W | A | A | W | A | W | A | A | W | A | A | A | A | W | A | A | W | W | W | A | A | A | W | W | W | A | W | A | W | A | A | A | W | W | A | W | A | W | A | A | W | W | W | A | W | A | A | A | A | A | W | A | A | A | W | A | W | W | W | A | A | W | W | A | W | W | A | A | W | A | W | A | A | W | W | W | A | W | W | W | A | W | W | A | W | A | W | A |
| Chrl | 35.7 | W | A | A | A | W | A | A | W | A | W | A | A | W | A | A | A | A | W | A | A | W | W | W | A | A | A | W | W | W | A | W | A | NA | A | A | A | W | W | A | W | A | W | NA | A | W | W | W | A | W | A | A | A | A | A | W | A | A | A | W | A | W | W | W | A | A | W | W | A | NA | W | A | A | W | A | NA | A | A | W | W | W | A | W | W | W | A | W | NA | A | W | A | W | A |
| Chrl | 35.8 | W | A | A | A | NA | A | NA | W | A | W | A | A | W | A | A | A | A | W | A | A | W | W | NA | A | A | A | W | W | W | A | W | A | W | A | A | A | W | W | A | W | A | W | W | A | W | W | W | A | W | NA | A | NA | A | A | W | A | A | A | W | A | W | W | W | A | A | W | W | A | W | W | A | A | W | A | W | A | A | W | W | W | A | W | W | W | A | W | W | A | W | A | W | A |

| | | | | | | | | | | | | | | | | | | | | | | | | | | | | | | | | | | | | | | | | | | | | | | | | | | | | | | | | | | | | | | | | | | | | | | | | | | | | | | | | | | | | | | | | | | | | | |
|---|---|---|---|---|---|---|---|---|---|---|---|---|---|---|---|---|---|---|---|---|---|---|---|---|---|---|---|---|---|---|---|---|---|---|---|---|---|---|---|---|---|---|---|---|---|---|---|---|---|---|---|---|---|---|---|---|---|---|---|---|---|---|---|---|---|---|---|---|---|---|---|---|---|---|---|---|---|---|---|---|---|---|---|---|---|---|---|---|---|---|---|---|---|
| ChrI | 35.9 | W | A | A | A | NA | A | A | W | A | W | A | A | W | A | A | A | A | W | A | A | W | W | W | A | A | A | W | W | W | A | W | A | W | A | A | A | W | W | A | W | A | W | W | A | W | W | W | A | W | A | A | A | A | A | W | A | A | A | W | A | W | W | W | A | A | W | W | A | W | W | A | A | NA | A | W | A | A | NA | W | W | A | W | W | W | A | NA | NA | A | W | A | W | A |
| ChrI | 36.1 | W | A | A | A | W | A | A | W | A | W | A | A | W | A | A | A | A | W | A | A | W | W | W | A | A | A | W | W | W | A | W | A | W | A | A | W | W | W | A | W | A | W | W | A | W | W | W | A | W | A | A | A | A | A | W | A | A | A | W | A | W | W | W | A | A | W | W | A | W | W | A | A | W | A | W | A | A | W | W | W | A | W | W | W | A | W | W | A | W | A | W | A |
| ChrI | 36.2 | W | A | A | A | W | A | A | W | A | W | A | A | W | A | A | A | A | W | A | A | W | W | W | A | A | A | W | W | W | A | W | A | W | A | A | W | W | W | A | W | A | W | W | A | W | W | W | A | W | A | A | A | A | A | W | A | A | A | NA | A | W | W | W | A | A | W | W | A | W | W | A | A | NA | A | W | A | A | NA | NA | W | A | W | W | W | A | W | W | A | W | A | W | A |
| ChrI | 36.3 | W | A | A | A | W | A | A | W | A | W | A | A | W | A | A | A | A | W | A | A | W | W | W | A | A | A | W | W | W | A | W | A | W | A | A | W | W | W | A | W | A | W | W | A | W | W | W | A | W | A | A | A | A | A | W | A | A | A | A | A | W | W | W | A | A | W | W | A | W | W | A | A | W | A | W | A | A | W | W | W | A | W | W | W | A | W | W | A | W | A | W | A |
| ChrI | 36.4 | W | A | A | A | W | A | A | W | A | W | A | A | W | A | A | A | NA | W | A | A | W | W | W | A | A | A | W | W | W | A | W | A | W | A | A | W | W | W | A | W | A | W | W | A | W | W | W | A | W | A | A | A | A | A | W | A | A | A | A | A | W | W | W | A | A | W | W | A | W | W | A | A | W | A | W | A | A | W | W | W | A | W | W | W | A | W | W | A | W | A | W | A |
| ChrI | 36.5 | W | A | A | A | W | A | A | W | A | W | A | A | W | A | A | A | A | W | A | A | W | W | W | A | A | A | W | W | W | A | W | A | NA | A | A | W | W | W | A | W | A | W | W | A | W | W | W | A | W | A | A | A | A | A | W | A | A | A | A | A | W | W | W | A | A | W | W | A | W | W | A | A | W | A | W | A | A | W | W | W | A | W | W | W | A | W | W | A | W | A | W | A |
| ChrI | 36.6 | W | A | A | A | W | A | A | W | A | W | A | A | W | A | A | A | A | W | A | A | NA | W | W | A | A | A | W | W | W | A | W | A | W | A | A | W | W | W | A | W | A | W | W | A | W | W | NA | A | W | A | A | A | A | A | W | A | A | A | A | A | W | W | W | A | A | W | W | NA | W | W | A | A | W | A | W | A | A | W | W | W | A | W | W | W | A | W | W | A | W | A | W | A |
| ChrI | 36.7 | W | A | A | A | W | A | A | W | A | W | A | A | W | A | A | A | A | NA | A | A | W | W | W | A | A | A | NA | W | W | A | W | A | W | A | A | W | W | W | A | W | A | W | W | A | W | W | A | A | NA | A | A | A | A | A | W | A | A | A | A | A | W | W | W | A | A | W | W | W | W | W | A | A | W | A | W | A | A | W | W | W | A | W | W | W | A | W | W | A | W | A | W | NA |
| ChrI | 37 | W | A | A | A | NA | A | A | W | A | W | A | A | W | A | A | A | A | NA | A | A | W | W | W | A | A | A | W | W | W | A | W | A | NA | A | A | W | W | W | A | W | A | W | W | A | W | W | A | A | W | A | A | A | A | A | NA | A | A | A | A | A | W | W | W | A | A | W | W | W | NA | W | A | A | NA | A | NA | A | A | NA | NA | W | A | W | W | W | A | W | W | A | W | A | W | A |
| ChrI | 37.3 | W | A | A | A | W | A | A | W | A | W | A | A | W | A | A | A | A | W | A | A | W | W | W | A | A | A | W | W | W | A | W | A | W | A | A | W | W | W | A | W | A | W | W | A | W | W | A | A | W | A | A | NA | A | A | W | A | A | A | A | A | W | W | W | A | A | W | W | W | W | W | A | A | W | A | W | A | A | W | W | W | A | W | W | W | A | W | W | A | W | A | W | A |
| ChrI | 37.4 | W | A | A | A | W | A | A | W | A | W | A | A | W | A | A | A | A | W | A | A | W | W | W | A | A | A | W | W | W | A | NA | A | W | A | A | W | W | W | A | W | A | W | W | A | W | W | A | A | W | A | A | A | A | A | W | A | A | A | A | A | W | W | W | A | A | W | W | W | W | W | A | A | NA | A | W | A | A | NA | NA | W | A | W | W | W | A | NA | W | A | W | A | W | A |
| ChrI | 37.5 | W | A | A | A | W | A | NA | W | A | W | A | A | W | A | A | A | A | W | A | A | W | W | W | A | A | A | W | W | W | A | W | A | W | A | A | W | W | W | A | W | A | W | W | A | W | W | A | A | W | A | A | A | A | A | W | A | A | A | A | A | W | W | W | A | A | W | W | W | W | W | A | A | W | A | W | A | A | W | W | W | A | W | W | W | A | W | W | A | W | A | W | A |
| ChrI | 37.6 | W | A | A | A | W | A | A | W | A | W | A | A | W | A | A | A | A | NA | A | A | W | W | W | A | A | A | W | W | W | A | W | A | W | A | A | W | W | W | A | W | A | W | W | A | W | W | A | A | W | A | A | A | A | A | W | A | A | A | A | A | W | W | W | A | A | W | W | W | NA | W | A | A | W | A | W | A | A | W | W | W | A | W | W | W | A | W | NA | A | W | A | W | A |
| ChrI | 37.7 | W | A | A | A | W | A | A | W | A | W | A | A | W | A | A | A | A | W | A | A | W | W | W | A | A | A | W | W | W | A | W | A | W | A | A | W | W | W | A | W | A | W | W | A | W | W | A | A | W | A | A | A | A | A | W | A | A | A | A | A | W | W | W | A | A | W | W | W | W | W | A | A | W | A | W | A | A | W | W | W | A | W | W | W | A | W | W | A | W | A | W | A |
| ChrI | 37.8 | W | A | A | A | W | A | A | W | A | W | A | A | W | A | A | A | A | NA | A | A | W | W | W | A | A | NA | W | W | W | A | W | A | W | A | A | W | W | W | A | W | A | W | W | A | W | W | A | A | W | NA | A | A | A | A | W | A | A | A | A | A | W | W | W | A | A | W | W | W | W | W | A | A | NA | A | W | A | A | W | W | W | A | W | W | W | A | W | W | A | W | A | W | A |
| ChrI | 37.9 | W | A | A | A | W | A | A | W | A | W | A | A | W | A | A | A | A | W | A | A | W | W | W | A | A | W | W | W | W | A | W | A | NA | A | A | W | W | W | A | W | A | W | W | A | W | W | A | A | W | A | A | A | A | A | W | A | A | A | A | A | W | W | W | A | A | W | W | W | W | W | A | A | W | A | W | A | A | W | W | W | A | W | W | W | A | W | W | A | W | A | W | A |
| ChrI | 38 | W | A | A | A | W | A | A | W | A | W | A | A | W | A | A | A | A | W | A | A | W | W | W | A | A | W | W | W | W | A | W | A | NA | A | A | W | W | W | A | W | A | W | W | A | W | W | A | A | W | A | A | A | A | A | W | A | A | A | NA | A | W | W | W | A | A | W | W | W | W | W | A | A | W | A | W | A | A | NA | W | W | A | W | W | W | A | W | NA | A | W | A | W | NA |
| ChrI | 38.1 | W | A | A | A | W | A | A | W | A | W | A | A | W | A | A | A | A | W | A | A | W | W | W | A | A | W | W | W | W | A | W | A | W | A | A | W | W | W | A | W | A | W | W | A | W | W | A | A | W | A | A | A | A | A | W | A | A | A | A | A | W | W | W | A | A | W | W | W | W | W | A | A | W | A | W | A | A | W | W | W | A | W | W | W | A | W | W | A | W | W | W | A |
| ChrI | 38.2 | W | A | A | A | NA | A | A | W | A | W | A | A | W | A | A | A | A | NA | A | A | W | W | W | A | A | W | W | W | W | A | W | A | W | A | A | W | W | W | A | W | A | W | W | A | W | W | A | A | W | A | A | A | A | A | W | A | A | A | A | A | W | W | W | A | A | W | W | W | NA | W | A | A | NA | A | W | A | A | W | W | W | A | W | W | W | A | W | W | A | W | W | W | A |
| ChrI | 38.4 | W | A | A | A | W | A | A | W | A | W | A | A | W | A | A | A | A | W | A | A | W | W | W | A | A | W | W | W | W | A | W | A | NA | A | A | W | W | W | A | W | A | W | W | A | W | W | A | A | W | A | A | A | A | A | W | A | A | A | A | A | W | W | W | A | A | W | W | W | W | W | A | A | W | A | W | A | A | A | W | W | A | W | W | W | A | W | W | A | W | W | W | A |
| ChrI | 38.5 | W | A | A | A | W | A | A | W | A | W | A | A | W | A | A | A | A | W | A | A | W | W | W | A | A | W | W | W | W | A | W | A | NA | A | A | W | W | W | A | W | A | W | W | A | W | W | A | A | W | A | A | A | A | A | W | A | A | A | A | A | W | W | W | A | A | W | W | W | W | W | A | A | W | A | W | A | A | A | W | W | A | W | W | W | A | W | W | A | W | W | W | A |
| ChrI | 38.6 | W | A | A | A | W | A | A | W | A | W | A | A | W | A | A | A | A | W | A | A | W | W | W | A | A | W | W | W | W | A | W | A | W | A | A | W | W | W | A | W | A | W | W | A | W | W | A | A | W | A | A | A | A | A | W | A | A | A | A | A | W | W | W | A | A | W | W | W | W | W | A | A | W | A | W | A | A | A | W | W | A | W | W | W | A | W | W | A | W | W | W | A |
| ChrI | 38.8 | W | A | A | A | W | A | A | W | A | W | A | A | W | A | A | A | A | W | A | A | W | W | W | A | A | W | W | W | W | A | W | A | W | A | A | W | W | W | A | W | A | W | W | A | W | W | A | A | W | A | A | A | A | A | W | A | A | A | A | A | W | W | W | A | A | W | W | W | W | W | A | A | W | A | W | A | A | A | W | W | A | W | W | W | A | W | W | A | W | W | W | A |
| ChrI | 38.9 | W | A | A | A | W | A | A | W | A | W | A | A | W | A | A | A | A | W | A | A | W | W | W | A | A | W | W | W | W | A | W | A | W | A | A | W | W | W | A | W | A | W | W | A | W | W | A | A | W | A | A | A | A | A | W | A | A | A | A | A | W | W | W | A | A | W | W | W | W | W | A | A | W | A | W | A | A | A | W | W | A | W | W | W | A | W | W | A | W | W | W | A |
| ChrI | 39.1 | W | A | A | A | W | A | A | W | A | W | A | A | W | NA | A | NA | A | W | A | A | W | W | W | A | A | W | W | W | W | A | W | A | W | NA | A | W | W | W | A | W | A | W | W | A | W | W | A | A | W | A | A | A | A | A | W | A | A | A | A | A | W | W | W | A | A | W | W | W | W | W | NA | A | W | A | W | A | A | A | W | W | A | W | W | W | A | W | W | A | W | W | W | NA |
| ChrI | 39.2 | W | A | A | A | NA | A | NA | W | A | W | A | A | W | A | A | A | A | W | A | A | W | W | W | A | A | W | W | W | W | A | W | A | W | A | A | W | W | W | A | W | A | W | W | A | W | W | A | A | W | A | A | A | A | A | W | A | A | A | A | A | W | W | W | A | A | W | W | W | W | W | A | A | W | A | W | A | A | A | W | W | A | W | W | W | A | W | W | A | W | W | W | A |
| ChrI | 39.3 | W | A | A | A | NA | A | NA | W | A | W | A | A | W | A | A | A | A | W | A | A | W | W | W | A | A | W | W | W | W | A | W | A | NA | A | A | W | W | W | A | W | A | W | W | A | W | W | A | A | W | A | A | A | A | A | W | A | A | A | A | A | W | W | W | A | A | W | W | W | W | W | A | A | NA | A | W | A | A | A | W | W | A | W | W | W | A | W | NA | A | W | W | W | A |
| ChrI | 39.4 | W | A | A | A | W | A | A | W | A | W | A | A | W | A | A | A | A | W | A | A | W | W | W | A | A | W | W | W | W | A | W | A | W | A | A | W | W | W | A | W | A | W | W | A | W | W | A | A | W | A | A | A | A | A | W | A | A | A | A | A | W | W | W | A | A | W | W | W | W | W | A | A | W | A | W | A | A | A | W | W | A | W | W | W | A | W | W | A | W | W | W | A |
| ChrI | 39.5 | W | A | A | A | W | A | NA | W | A | W | A | A | W | A | A | A | A | W | A | A | W | W | W | A | A | W | W | W | W | A | W | A | W | NA | A | W | W | W | A | W | A | W | W | A | W | W | A | A | W | A | NA | A | A | A | W | A | A | A | A | A | W | W | W | NA | A | W | W | W | W | W | A | A | W | A | W | A | A | A | W | W | A | W | W | W | A | W | W | A | W | W | W | A |
| ChrII | 0.1 | A | W | W | A | W | A | A | A | W | W | W | A | A | W | W | W | A | A | W | A | A | A | A | A | A | A | A | A | W | W | A | W | W | A | A | NA | A | A | W | W | A | W | A | W | W | W | W | A | W | W | A | A | A | A | A | A | A | A | W | A | A | A | W | W | A | A | W | A | W | W | A | W | A | A | W | W | W | A | W | A | A | A | A | A | A | W | A | W | W | W | A | W |
| ChrII | 0.2 | A | W | W | A | W | A | A | A | W | W | W | A | A | W | W | W | A | A | W | A | A | A | A | A | NA | A | A | A | W | W | A | W | W | NA | A | W | A | A | W | W | A | NA | A | W | W | W | W | A | W | W | A | A | A | A | A | A | A | A | W | A | A | A | NA | NA | A | A | W | A | W | W | A | W | A | A | W | W | W | A | W | A | A | A | A | A | A | NA | A | W | W | W | A | W |
| ChrII | 0.3 | A | W | W | A | W | A | A | A | W | W | W | A | A | W | W | W | A | A | NA | A | A | A | A | A | A | A | A | A | NA | W | A | W | NA | A | A | NA | A | A | W | NA | A | NA | A | W | W | W | W | A | W | W | A | A | A | A | A | A | A | A | W | A | A | A | NA | NA | A | A | NA | A | W | W | A | W | A | A | W | W | W | A | W | A | A | A | A | A | A | NA | A | W | W | W | A | W |
| ChrII | 0.5 | A | W | W | A | W | A | A | A | W | W | W | A | A | W | W | W | A | A | NA | A | A | A | A | A | A | A | A | A | NA | W | A | W | NA | A | A | NA | A | A | W | W | A | NA | A | W | W | W | W | A | W | W | A | A | A | A | A | A | A | A | W | A | A | A | NA | NA | A | A | W | NA | W | W | A | W | A | A | W | W | W | A | W | A | A | A | A | A | A | NA | A | W | W | W | A | W |
| ChrII | 0.6 | A | W | W | A | W | A | A | A | W | W | W | A | A | W | W | W | A | A | W | A | A | A | A | A | A | A | A | A | W | W | A | W | NA | A | A | W | A | A | W | W | A | W | A | W | W | W | W | A | W | NA | A | A | A | A | A | A | A | A | W | A | A | A | W | W | A | A | W | A | W | W | A | W | A | A | W | W | W | A | W | A | A | A | A | A | A | W | A | W | W | W | A | W |
| ChrII | 0.7 | A | W | W | A | W | A | A | A | W | W | W | A | A | W | W | W | A | A | NA | A | A | A | A | A | A | A | A | A | W | W | A | W | NA | A | A | NA | A | A | W | NA | A | W | A | W | W | W | W | A | W | NA | A | A | A | A | A | A | A | A | W | A | A | A | NA | NA | A | A | W | A | W | W | A | W | A | A | W | W | W | A | W | A | A | A | A | A | A | W | A | W | W | W | A | W |
| ChrII | 0.8 | A | W | W | A | W | A | A | A | W | W | W | A | A | W | W | W | A | A | NA | A | A | A | A | A | A | A | A | A | NA | W | A | W | NA | A | A | W | NA | A | W | NA | A | NA | A | W | W | W | W | A | W | NA | A | NA | A | A | A | A | A | A | W | A | A | A | NA | W | A | A | W | NA | W | W | NA | W | A | A | W | W | W | A | W | A | A | A | A | A | A | NA | NA | W | W | W | A | W |
| ChrII | 0.9 | A | W | W | A | W | A | A | A | W | W | W | A | A | W | W | W | A | NA | NA | A | A | A | A | A | A | A | A | A | W | W | A | W | NA | A | A | W | A | A | W | W | A | W | A | W | W | W | W | A | W | W | A | A | A | A | A | A | A | A | W | A | A | A | W | NA | A | A | W | A | W | W | A | W | A | A | W | W | W | A | W | A | A | A | A | A | A | W | A | W | W | W | A | W |
| ChrII | 1 | A | W | W | A | W | A | A | A | W | W | W | A | A | W | W | W | A | W | W | A | A | A | NA | A | A | A | A | A | W | W | A | W | W | A | A | W | A | A | W | W | A | W | A | W | W | W | W | A | W | W | A | A | A | A | NA | A | A | A | W | A | NA | NA | W | W | A | A | W | A | W | W | A | W | A | A | W | W | W | A | W | A | A | A | A | A | A | W | A | W | W | W | A | W |
| ChrII | 1.1 | A | W | W | A | W | A | A | A | W | W | W | A | A | W | W | W | A | W | W | A | A | A | A | A | A | A | A | A | W | W | A | W | NA | A | A | NA | A | A | W | W | A | W | A | W | W | W | W | A | W | NA | A | A | A | A | A | A | A | A | W | A | A | A | W | NA | A | A | W | NA | W | W | NA | W | A | A | W | W | W | A | W | A | A | A | A | A | A | W | A | W | W | W | A | W |
| ChrII | 1.2 | A | W | W | A | W | A | A | A | W | W | W | A | A | W | W | W | A | W | W | A | NA | A | A | A | A | A | A | A | W | W | A | W | W | A | A | W | A | A | W | W | A | W | A | W | W | W | W | A | W | NA | A | A | A | A | A | A | A | A | W | A | A | A | W | W | A | A | W | A | W | W | A | W | A | A | W | W | W | A | W | A | A | A | A | A | A | NA | A | W | W | W | A | W |
| ChrII | 1.3 | A | W | W | A | W | A | A | A | W | W | W | A | A | W | W | W | A | W | NA | A | A | A | A | A | A | A | A | A | NA | W | A | W | W | A | A | NA | A | A | W | W | A | W | A | W | W | W | W | A | W | NA | A | A | A | A | A | A | A | A | W | A | A | A | W | W | A | A | W | A | W | W | A | W | A | A | W | W | W | A | W | A | A | A | A | A | A | W | A | W | W | W | A | W |
| ChrII | 1.4 | A | W | W | A | W | A | A | A | W | W | W | A | A | W | W | W | A | W | W | A | A | A | A | A | A | A | A | A | W | W | A | W | W | A | A | W | A | A | W | W | A | W | A | W | W | W | W | A | W | NA | A | A | A | A | A | A | A | A | W | A | NA | A | W | W | A | A | W | A | W | W | A | W | A | A | W | W | W | A | W | A | A | A | A | A | A | W | A | W | W | W | A | W |
| ChrII | 1.5 | A | W | W | A | W | A | A | A | W | W | W | A | A | W | W | W | A | W | W | A | A | A | A | A | A | A | A | A | W | W | A | W | NA | A | A | W | A | A | W | W | A | W | A | W | W | W | W | A | W | W | A | A | A | A | A | A | A | A | W | A | A | A | W | W | A | A | W | A | W | W | A | W | A | A | W | W | W | A | W | A | A | A | A | A | A | W | A | W | W | W | A | W |
| ChrII | 1.6 | A | W | W | A | W | A | A | A | W | W | W | A | A | W | W | W | A | W | W | A | A | A | A | A | A | A | A | A | W | W | A | W | NA | A | A | NA | A | A | W | W | A | W | A | W | W | W | W | A | W | W | A | A | A | A | A | A | A | A | W | A | A | A | W | W | A | A | W | A | W | W | A | W | A | A | W | W | W | A | W | A | A | A | A | A | A | W | A | W | W | W | A | W |
| ChrII | 1.7 | A | W | W | A | W | A | A | A | W | W | W | A | A | W | W | W | A | W | NA | A | A | A | A | A | A | A | A | A | NA | W | A | W | NA | A | A | NA | NA | A | W | NA | A | W | A | W | W | W | W | A | W | NA | A | NA | A | A | A | A | A | A | W | A | A | A | NA | NA | A | A | W | NA | W | W | NA | W | A | A | W | W | W | A | W | A | A | A | A | A | A | NA | A | W | W | W | A | W |
| ChrII | 1.8 | A | W | W | A | W | A | A | A | W | W | W | A | A | W | W | W | A | W | NA | A | A | A | A | A | A | A | A | A | NA | NA | A | W | NA | A | A | NA | A | A | W | NA | A | W | A | W | W | W | W | A | W | W | A | A | A | A | A | A | A | A | W | A | A | A | NA | NA | A | A | W | A | W | W | A | W | A | A | W | W | W | A | W | A | A | A | A | A | A | NA | A | W | W | W | A | W |
| ChrII | 1.9 | A | W | W | A | W | A | A | A | W | W | W | A | A | W | W | W | A | W | W | A | A | A | A | A | A | A | A | A | W | W | A | W | W | A | A | W | A | A | W | W | A | W | A | W | W | W | W | A | W | W | A | A | A | A | A | A | A | A | W | A | A | A | W | NA | A | A | W | A | W | W | NA | W | A | A | W | W | W | A | W | A | A | A | A | A | A | W | A | W | W | W | A | W |
| ChrII | 2 | A | W | W | A | W | A | A | A | W | W | W | A | A | W | W | W | NA | W | W | A | A | A | A | A | A | A | A | A | NA | W | A | W | NA | A | A | NA | A | A | W | NA | A | NA | A | W | W | W | W | A | W | W | A | A | A | A | A | A | A | A | W | A | A | A | W | NA | A | A | W | A | W | W | A | W | A | A | W | W | W | A | W | A | A | A | A | A | A | W | A | W | W | W | A | W |
| ChrII | 2.1 | A | W | W | A | W | A | A | A | W | W | W | A | A | W | W | W | A | W | W | A | A | A | A | A | A | A | A | A | W | W | A | W | W | A | A | W | A | A | W | W | A | W | A | W | W | W | W | A | W | W | A | A | A | A | A | A | A | A | W | A | A | A | W | NA | A | A | W | A | W | W | A | W | A | A | W | W | W | A | W | A | A | A | A | A | A | NA | A | W | W | W | A | W |
| ChrII | 2.2 | A | W | W | A | W | A | A | A | W | W | W | NA | A | W | W | W | A | W | W | A | A | A | A | A | A | A | A | A | W | W | A | W | W | A | A | NA | A | A | W | W | A | W | A | W | W | W | W | NA | W | NA | A | NA | A | A | A | A | A | A | W | A | A | A | NA | NA | A | A | W | A | W | W | A | W | A | A | W | W | W | A | W | A | A | A | A | A | A | W | NA | W | W | W | A | W |
| ChrII | 2.3 | A | W | W | A | W | A | A | A | W | W | W | A | A | W | W | W | NA | W | W | A | A | A | A | A | A | A | A | A | W | W | A | W | NA | A | A | NA | A | A | W | NA | A | W | A | W | W | W | W | A | W | NA | A | A | A | A | A | A | A | A | W | A | A | A | NA | NA | A | A | W | A | W | W | A | W | A | A | W | W | W | A | W | A | A | A | A | A | A | NA | W | W | W | W | A | W |
| ChrII | 2.5 | A | W | W | A | W | A | A | A | W | W | W | A | A | W | W | W | A | W | W | NA | A | A | A | A | A | NA | A | A | W | W | A | W | W | A | A | W | A | A | W | W | A | W | A | W | W | W | W | A | W | NA | A | A | A | A | A | A | A | A | W | A | A | NA | W | W | A | A | W | A | W | W | NA | W | A | A | W | W | W | A | W | A | A | A | A | A | A | W | W | W | W | W | A | W |
| ChrII | 2.6 | A | W | W | A | W | A | A | A | W | W | W | A | A | W | W | W | A | W | W | A | A | A | A | A | A | A | A | A | W | W | A | W | NA | A | A | NA | A | A | W | W | A | NA | A | W | W | W | W | A | W | NA | A | A | A | A | A | A | A | A | W | A | A | A | NA | W | A | A | W | A | W | W | A | W | A | A | W | W | W | A | W | A | A | A | A | A | A | W | W | W | W | W | A | W |
| ChrII | 2.7 | A | W | W | A | W | A | A | A | W | W | W | A | A | W | W | W | A | W | W | A | A | A | A | A | A | A | A | A | W | W | A | W | W | A | A | W | A | A | W | W | A | W | A | W | W | W | W | A | W | W | A | A | A | A | A | A | A | A | W | A | A | A | W | W | A | A | W | A | W | W | A | W | A | A | W | W | W | A | W | A | A | A | A | A | A | W | W | W | W | W | A | W |
| ChrII | 2.8 | A | W | W | A | W | A | A | A | W | W | W | A | A | W | W | W | A | W | W | A | A | A | A | A | A | A | A | A | W | W | A | W | NA | A | A | W | A | A | W | NA | A | W | A | W | W | W | W | A | W | W | A | A | A | A | A | A | A | A | W | A | A | A | W | W | A | A | W | A | W | W | A | W | A | A | W | W | W | A | W | A | A | A | A | A | A | W | W | W | W | W | A | W |
| ChrII | 3 | A | W | W | A | W | A | A | A | W | W | W | A | A | W | W | W | A | W | W | A | A | A | A | A | A | A | A | A | W | W | A | W | NA | A | A | W | A | A | W | W | A | W | A | W | W | W | W | A | W | NA | A | A | A | A | A | A | A | A | W | A | A | A | W | W | A | A | W | A | W | W | A | W | A | A | W | W | W | A | W | A | A | A | A | A | A | W | W | W | W | W | A | W |
| ChrII | 3.1 | A | W | W | A | W | A | A | A | W | W | W | A | A | W | W | W | A | W | W | A | A | A | A | A | A | A | A | A | W | W | A | W | NA | A | A | W | A | A | W | W | A | W | A | W | W | W | W | A | W | NA | A | A | A | A | A | A | A | A | W | A | A | A | W | W | A | A | W | A | W | W | A | W | A | A | W | W | W | A | W | A | A | A | A | A | A | W | W | W | W | W | A | W |
| ChrII | 3.2 | A | W | W | A | W | A | NA | A | W | W | W | A | A | W | W | W | NA | W | W | A | NA | A | A | A | NA | A | A | A | W | W | A | W | NA | NA | A | NA | A | A | W | NA | A | W | A | W | W | W | W | A | W | W | A | A | A | NA | A | A | A | A | W | A | NA | A | W | W | A | NA | W | A | W | W | A | W | A | A | W | W | W | A | W | NA | A | A | A | A | A | W | W | W | W | W | A | W |
| ChrII | 3.3 | A | W | W | A | W | A | A | A | W | W | W | A | A | W | W | W | A | W | W | A | A | A | A | A | A | A | A | A | W | W | A | W | W | A | A | W | A | A | W | A | A | W | A | W | W | W | W | A | W | W | A | A | A | A | A | A | A | A | W | A | A | A | W | W | A | A | W | A | W | W | A | W | A | A | W | W | W | A | W | A | A | A | A | A | A | NA | W | W | W | W | A | W |
| ChrII | 3.4 | A | W | W | A | W | A | A | A | W | W | W | A | A | W | W | W | A | W | W | A | A | A | A | A | A | A | A | A | W | W | A | W | NA | A | A | W | A | A | W | A | A | W | A | W | W | W | W | A | W | NA | A | A | A | A | A | A | A | A | W | A | A | A | W | W | A | A | W | A | W | W | A | W | A | A | W | W | W | A | W | A | A | A | A | A | A | W | W | W | W | W | A | W |
| ChrII | 3.5 | A | W | W | A | A | A | A | A | W | W | W | A | A | W | W | W | A | W | W | A | A | A | A | A | A | A | A | A | W | W | A | W | W | A | A | NA | A | A | W | A | A | W | A | W | W | W | W | A | W | NA | A | A | A | A | A | A | A | A | W | A | A | A | W | W | A | A | W | A | W | W | A | W | A | A | W | W | W | A | W | A | A | A | A | A | A | W | W | W | W | W | A | W |
| ChrII | 3.6 | A | W | W | A | A | A | A | A | W | W | W | A | A | W | W | W | A | W | NA | A | A | A | A | A | A | A | A | A | W | W | A | W | NA | A | A | NA | A | A | W | A | A | W | A | W | W | W | W | A | W | NA | A | A | A | A | A | A | A | A | W | A | A | A | NA | W | A | A | W | A | W | W | A | W | A | A | W | W | W | A | W | A | A | A | A | A | A | W | W | W | W | W | A | W |
| ChrII | 3.7 | W | W | W | A | A | A | A | A | W | W | W | A | A | W | W | W | A | W | W | A | A | A | A | A | A | A | A | A | NA | W | A | W | W | A | A | W | A | A | W | A | A | W | A | W | W | W | W | A | W | W | A | A | A | A | A | A | A | A | W | A | A | A | W | W | A | A | W | A | W | W | A | W | A | A | W | W | W | A | W | A | A | A | A | A | A | W | W | W | W | W | A | W |
| ChrII | 3.8 | W | W | W | A | A | A | A | A | W | W | W | A | A | W | W | W | A | W | NA | A | A | A | A | A | A | A | A | A | W | W | A | W | NA | A | A | W | A | A | W | A | A | W | A | W | W | W | W | A | W | W | A | A | A | A | A | NA | A | A | W | A | A | A | NA | W | A | A | W | A | W | W | A | W | NA | A | W | W | W | A | W | A | A | A | A | A | A | W | W | W | W | W | A | W |
| ChrII | 3.9 | W | W | W | A | A | A | A | A | W | W | W | A | A | W | W | W | A | W | NA | A | A | A | A | A | A | A | A | A | NA | NA | A | W | NA | NA | A | NA | A | A | W | A | A | W | A | W | W | W | W | A | W | W | A | A | A | A | A | A | A | A | W | A | A | A | NA | W | A | A | W | A | W | W | A | W | A | A | W | W | W | A | W | A | A | A | A | A | A | W | W | W | W | W | A | W |
| ChrII | 4 | W | W | W | A | A | A | A | A | W | W | W | A | A | W | W | W | A | W | NA | A | A | A | A | A | A | A | A | A | NA | W | A | W | NA | A | A | NA | A | A | W | A | A | W | A | W | W | W | W | A | W | W | A | A | A | A | A | A | A | A | W | A | A | A | W | W | A | A | W | A | W | W | A | W | A | A | W | W | W | A | W | A | A | A | A | A | A | NA | W | W | W | W | A | W |
| ChrII | 4.1 | W | W | W | A | A | A | A | A | W | W | W | A | A | W | W | W | A | W | W | A | A | A | A | A | A | A | A | A | NA | W | A | W | W | A | A | NA | A | A | W | A | A | W | A | W | W | W | W | A | W | NA | A | A | A | A | A | A | A | A | W | A | A | A | W | W | A | A | W | A | W | W | A | W | A | A | W | W | W | A | W | A | A | A | A | A | A | NA | W | W | W | W | A | W |
| ChrII | 4.2 | W | W | W | A | A | A | A | A | W | W | W | A | A | W | W | W | A | W | W | A | A | A | A | A | A | A | A | A | W | W | A | W | W | A | A | W | A | A | W | A | A | W | A | W | W | W | A | A | W | W | A | A | A | A | A | A | A | A | W | A | A | A | W | W | A | A | W | A | W | A | A | W | A | W | W | W | W | A | W | A | A | A | A | A | A | W | W | W | W | W | A | W |
| ChrII | 4.3 | W | W | W | A | A | A | A | A | W | W | W | A | A | W | W | W | A | W | W | A | A | A | A | A | A | A | A | A | W | W | A | W | W | A | A | W | A | A | W | A | A | W | A | W | W | W | A | A | W | W | A | A | A | A | A | A | A | A | W | A | A | A | W | W | A | A | W | A | W | A | A | W | A | W | W | W | W | A | W | A | A | A | A | A | A | W | W | W | W | W | A | W |
| ChrII | 4.4 | W | W | W | A | A | A | A | A | W | W | W | A | A | W | W | W | A | W | W | A | A | A | A | A | A | A | A | A | W | W | A | W | NA | A | A | NA | A | A | W | A | A | W | A | W | W | W | A | A | W | W | A | A | A | A | A | A | A | A | W | A | A | A | NA | W | A | A | W | A | W | A | A | W | A | W | W | W | W | A | W | A | A | A | A | A | A | W | W | W | W | W | A | W |
| ChrII | 4.5 | W | W | W | A | A | A | A | A | W | W | W | A | A | W | W | W | A | W | W | A | A | A | A | A | A | A | A | A | W | W | A | W | NA | A | A | W | A | A | W | A | A | W | A | W | W | W | A | A | W | NA | A | A | A | A | A | A | A | A | W | A | A | A | W | W | A | A | W | A | W | A | A | W | A | W | W | W | W | A | W | A | A | A | A | A | A | W | W | W | W | W | A | W |
| ChrII | 4.6 | W | W | W | A | A | A | A | A | W | W | W | A | A | W | W | W | A | W | W | A | A | A | A | A | A | A | A | A | W | W | A | W | NA | A | A | W | A | A | W | A | A | W | A | W | W | W | A | A | W | W | A | A | A | A | A | A | A | A | W | A | A | A | W | W | A | A | W | A | W | A | A | W | A | W | W | W | W | A | W | A | A | A | A | A | A | W | W | W | W | W | A | W |
| ChrII | 4.7 | W | W | W | A | A | A | A | A | W | W | W | A | A | W | W | W | A | W | W | A | A | A | A | A | A | A | A | A | NA | W | A | W | NA | A | A | NA | A | A | W | A | A | W | A | W | W | W | A | A | W | W | A | A | A | A | A | A | A | A | W | A | A | A | W | W | A | A | W | A | W | A | A | W | A | W | W | W | W | A | W | A | A | A | A | A | A | W | W | W | W | W | A | W |
| ChrII | 4.8 | W | W | W | A | A | A | A | A | W | W | W | A | A | W | W | W | A | W | W | A | A | A | A | A | A | A | A | A | W | W | A | W | NA | A | A | W | A | A | W | A | A | W | A | W | W | W | A | A | W | NA | A | A | A | A | A | A | A | A | W | A | A | A | W | W | A | A | W | A | W | A | A | W | A | W | W | W | W | A | W | A | A | A | A | A | A | NA | W | W | W | W | A | W |
| ChrII | 4.9 | W | W | W | A | A | A | A | A | W | W | W | A | A | W | W | W | A | W | W | A | A | A | A | A | A | A | A | A | NA | W | A | W | W | A | A | W | A | A | W | A | A | W | A | W | W | W | A | A | W | NA | A | A | A | A | A | A | A | A | W | A | A | A | NA | W | A | A | W | A | W | A | A | W | A | W | W | W | W | A | W | A | A | A | A | A | A | W | W | W | W | W | A | W |
| ChrII | 5 | W | W | W | A | A | A | A | A | W | W | W | A | A | W | W | W | A | W | W | A | A | A | A | A | A | A | NA | A | W | W | A | W | W | A | A | W | A | A | W | A | A | W | A | W | W | W | A | A | W | W | A | A | A | A | A | A | A | A | W | A | A | A | W | W | A | A | W | A | W | A | A | W | A | W | W | NA | W | A | W | A | A | A | A | A | A | NA | W | W | W | W | A | W |
| ChrII | 5.1 | W | W | W | A | A | A | A | A | W | W | W | A | A | W | W | W | A | W | NA | A | A | A | A | A | A | NA | A | A | NA | NA | A | W | W | A | A | NA | A | A | W | A | A | W | A | W | W | W | A | A | W | NA | A | A | A | A | A | A | A | A | W | A | A | A | NA | NA | W | A | W | NA | W | A | A | W | A | W | W | A | W | A | W | A | A | A | A | A | A | W | W | W | W | W | NA | W |
| ChrII | 5.2 | W | W | W | A | A | A | A | A | W | W | W | A | A | W | W | W | A | W | W | A | A | A | A | A | A | A | A | A | W | W | A | W | NA | A | A | W | A | A | W | A | A | W | A | W | W | W | A | A | W | W | A | A | A | A | A | A | A | A | W | A | A | A | W | W | W | A | W | A | W | A | A | W | A | W | W | A | W | A | W | A | A | A | A | A | A | W | W | W | W | W | A | W |
| ChrII | 5.3 | W | W | W | A | A | A | A | A | W | W | W | A | A | W | W | W | A | W | W | NA | A | A | A | A | A | A | A | A | W | W | A | W | W | A | A | W | A | A | W | A | A | W | A | W | W | W | A | A | W | W | A | A | A | A | A | A | A | A | W | A | A | A | W | W | W | A | W | A | W | A | A | W | A | W | W | A | W | A | W | A | A | A | A | A | A | W | W | W | W | W | A | W |
| ChrII | 5.4 | W | W | W | A | A | A | A | A | W | W | W | A | A | W | W | W | A | W | NA | A | A | A | A | A | A | A | A | A | NA | W | A | W | W | A | A | NA | A | A | W | A | A | W | A | W | W | W | A | A | W | NA | A | A | A | A | A | A | A | A | W | A | A | A | W | W | W | A | W | A | W | A | A | W | A | W | W | A | W | A | W | A | A | A | A | A | A | W | W | W | W | W | A | W |
| ChrII | 5.5 | W | W | W | A | A | A | A | A | W | W | W | A | A | W | W | W | A | W | W | A | A | A | A | A | A | A | A | A | W | W | A | W | NA | A | A | W | A | A | W | A | A | W | A | W | W | W | A | A | W | W | A | A | A | A | A | A | A | A | W | A | A | A | W | W | W | A | W | A | W | A | A | W | A | W | W | A | W | A | W | A | A | A | A | A | A | W | W | W | W | W | A | W |
| ChrII | 5.6 | W | W | W | A | A | A | A | A | W | W | W | A | A | W | W | W | A | W | W | A | A | A | A | A | A | A | A | A | W | W | A | W | W | A | A | W | A | A | W | A | A | W | A | W | W | W | A | A | W | W | A | A | A | A | A | A | A | A | W | A | A | A | W | W | W | NA | W | A | W | A | A | W | A | W | W | A | W | A | W | A | A | A | A | A | A | W | W | W | W | W | A | W |
| ChrII | 5.8 | W | W | W | A | A | A | A | A | W | W | W | A | A | W | W | W | A | W | NA | A | A | A | A | A | A | A | A | A | W | NA | A | W | NA | A | A | NA | A | A | W | A | A | W | A | W | W | W | A | A | W | NA | A | A | A | A | A | A | A | A | W | A | A | A | NA | W | W | A | W | A | W | A | A | W | A | W | W | A | W | A | W | A | A | A | A | A | A | W | W | W | W | W | A | W |
| ChrII | 5.9 | W | W | W | A | A | A | A | A | W | W | W | A | A | W | W | W | A | W | W | A | A | A | A | A | A | A | A | A | NA | W | A | W | NA | A | A | W | A | A | W | A | A | W | A | W | W | W | A | A | W | NA | A | A | A | A | A | A | A | A | W | A | A | A | W | W | W | A | W | A | W | A | A | W | A | W | W | A | W | A | W | A | A | A | A | A | A | W | W | W | W | W | A | W |
| ChrII | 6 | W | W | W | A | A | A | A | A | W | W | W | A | A | W | W | W | A | W | W | A | A | A | A | A | A | A | A | A | W | W | A | W | W | A | A | NA | A | A | NA | A | A | W | A | W | W | W | A | A | W | NA | A | A | A | A | A | A | A | A | W | A | A | A | W | W | W | A | W | A | W | A | A | W | A | W | W | A | W | A | W | A | A | A | A | A | A | W | W | W | W | W | A | W |
| ChrII | 6.1 | W | W | W | A | A | A | A | A | W | W | W | A | A | W | W | W | NA | W | NA | A | A | A | A | A | A | A | A | A | NA | W | A | W | NA | A | A | NA | NA | A | A | A | A | W | A | W | W | W | A | A | W | W | A | A | A | A | A | A | A | A | W | A | A | A | NA | W | W | A | W | A | W | A | A | W | A | W | W | A | W | A | W | A | A | A | A | A | A | W | W | W | W | W | A | W |
| ChrII | 6.2 | W | W | W | A | A | A | NA | A | W | W | W | A | NA | W | W | W | A | W | W | NA | A | A | A | A | A | A | A | A | W | W | A | W | W | A | NA | W | A | A | A | A | A | W | A | W | W | W | A | A | W | W | NA | A | A | A | A | A | A | A | W | A | NA | A | W | W | W | NA | W | A | W | A | A | W | A | W | W | NA | W | A | W | NA | A | A | A | A | A | W | W | W | W | W | A | W |
| ChrII | 6.3 | W | W | W | A | A | A | A | A | W | W | W | A | A | W | W | W | A | W | W | A | A | A | A | A | A | A | A | A | W | W | A | NA | NA | A | A | W | A | A | A | A | A | W | A | W | W | W | A | A | W | W | A | A | A | A | A | A | A | A | W | A | A | A | W | W | W | A | W | A | W | A | A | W | A | W | W | A | W | A | W | A | A | A | A | A | A | W | W | W | W | W | A | W |

| | | | | | | | | | | | | | | | | | | | | | | | | | | | | | | | | | | | | | | | | | | | | | | | | | | | | | | | | | | | | | | | | | | | | | | | | | | | | | | | | | | | | | | | | | | | | | |
|---|---|---|---|---|---|---|---|---|---|---|---|---|---|---|---|---|---|---|---|---|---|---|---|---|---|---|---|---|---|---|---|---|---|---|---|---|---|---|---|---|---|---|---|---|---|---|---|---|---|---|---|---|---|---|---|---|---|---|---|---|---|---|---|---|---|---|---|---|---|---|---|---|---|---|---|---|---|---|---|---|---|---|---|---|---|---|---|---|---|---|---|---|---|
| ChrII | 6.4 | W | W | W | A | A | A | A | A | W | W | W | A | A | W | W | W | A | W | W | A | A | A | A | A | A | A | A | A | W | W | A | W | W | A | A | W | A | A | A | A | A | W | A | W | W | W | A | A | W | W | A | A | A | A | A | A | A | A | W | A | A | A | W | W | W | A | W | A | W | A | A | W | A | W | W | A | W | A | W | A | A | A | A | A | A | W | W | W | W | W | A | W |
| ChrII | 6.5 | W | W | W | A | A | A | A | A | W | W | W | A | A | W | W | W | A | W | W | A | A | A | A | A | A | A | A | A | NA | W | A | W | W | A | A | NA | A | A | A | A | A | W | A | W | W | W | A | A | W | NA | A | A | A | A | A | A | A | A | W | A | A | A | NA | W | W | A | W | A | W | A | A | W | A | W | W | A | W | A | W | A | A | A | A | A | A | W | W | W | W | W | A | W |
| ChrII | 6.6 | W | W | W | A | A | A | A | A | W | W | W | A | A | W | W | W | A | W | W | A | A | A | A | A | A | NA | A | A | W | W | A | W | W | A | A | W | A | A | A | A | A | W | A | W | NA | W | A | A | W | W | A | A | A | A | A | A | A | A | W | A | A | A | W | W | W | A | W | A | W | A | A | W | A | W | W | A | W | A | W | A | A | A | A | A | A | W | W | W | W | W | A | W |
| ChrII | 6.7 | W | W | W | A | A | A | A | A | W | W | W | A | A | W | W | W | A | W | W | A | A | A | A | A | A | A | A | A | W | W | A | W | W | A | A | W | A | A | A | A | A | W | A | W | A | W | A | A | W | W | A | A | A | A | A | A | A | A | W | A | A | A | NA | W | W | A | W | A | W | A | A | W | A | W | W | A | W | A | W | A | A | A | A | A | A | W | W | W | W | W | A | W |
| ChrII | 6.8 | W | W | W | A | A | A | A | A | W | W | W | A | A | W | W | W | A | W | W | A | A | A | A | A | A | A | A | A | W | W | A | W | W | A | A | W | A | A | A | A | A | W | A | W | A | W | A | A | W | NA | A | A | A | A | A | A | A | A | W | A | A | A | NA | W | W | A | W | A | W | A | A | W | A | W | W | A | W | A | W | A | A | A | A | A | A | W | W | W | W | W | A | W |
| ChrII | 6.9 | W | W | W | A | A | A | A | A | W | W | W | A | A | W | W | W | A | W | W | A | A | A | A | A | A | A | A | A | W | A | A | W | W | A | A | NA | A | A | A | A | A | W | A | W | A | W | A | A | W | NA | A | A | A | A | A | A | A | A | W | A | A | A | W | W | W | A | W | A | W | A | A | W | A | W | W | A | W | A | W | A | A | A | A | A | A | W | W | W | W | W | A | W |
| ChrII | 7 | W | W | W | A | A | A | A | A | W | W | W | A | A | W | W | W | A | W | NA | A | A | A | A | A | A | A | A | A | W | A | A | NA | W | A | A | NA | A | A | A | A | A | W | A | W | A | W | A | A | W | W | A | A | A | A | A | A | A | A | W | A | A | A | NA | W | W | A | W | A | W | A | A | W | A | W | W | A | W | A | W | A | A | A | A | A | A | W | W | W | W | W | A | W |
| ChrII | 7.1 | W | W | W | A | A | A | A | A | W | W | W | A | A | W | W | W | A | W | W | A | A | A | A | A | A | A | A | A | W | A | A | W | W | A | A | W | A | A | A | A | A | W | A | W | A | W | A | A | W | W | A | A | A | A | A | A | A | A | W | A | A | A | W | W | W | A | W | A | W | A | A | W | A | W | W | A | W | A | W | A | A | A | A | A | A | W | W | W | W | W | A | W |
| ChrII | 7.2 | W | W | W | A | A | A | A | A | W | W | W | A | A | W | W | W | A | W | NA | A | A | A | A | A | A | A | A | A | W | A | A | W | W | A | A | NA | A | A | A | A | A | W | A | W | A | W | A | A | W | W | A | A | A | A | A | A | A | A | W | A | A | A | W | W | W | A | W | A | W | A | A | W | A | W | W | A | W | A | W | A | A | A | A | A | A | W | W | W | W | W | A | W |
| ChrII | 7.3 | W | W | W | A | A | A | A | A | W | W | W | A | A | W | W | W | A | W | NA | A | A | A | A | A | A | A | A | A | NA | A | A | W | W | NA | A | NA | A | A | A | A | A | W | A | W | A | W | A | A | W | W | A | A | A | A | A | A | A | A | W | A | A | NA | NA | W | W | A | W | A | W | NA | A | W | A | W | W | A | W | NA | W | A | NA | A | A | A | A | W | NA | W | W | W | A | W |
| ChrII | 7.4 | W | W | W | A | A | A | A | A | W | W | W | A | A | W | W | W | A | W | NA | A | A | A | A | A | A | A | A | A | NA | A | A | W | NA | A | A | NA | A | A | A | A | A | W | A | W | A | W | A | A | W | NA | A | A | A | A | A | A | A | A | W | A | A | A | NA | W | W | A | W | A | W | A | A | W | A | W | W | A | W | A | W | A | A | A | A | A | A | W | W | W | W | W | A | W |
| ChrII | 7.5 | W | W | W | A | A | A | A | A | W | W | W | A | A | W | W | W | A | W | W | A | A | A | A | A | A | A | A | A | NA | A | A | W | W | A | A | W | A | A | A | A | A | W | A | W | A | W | A | A | W | W | A | A | A | A | NA | A | A | A | W | A | A | A | NA | W | W | A | W | A | W | A | NA | W | A | W | W | A | W | A | W | A | A | A | A | A | A | W | W | W | W | W | A | W |
| ChrII | 7.6 | W | W | W | A | A | A | A | A | W | W | W | A | A | W | W | W | A | W | W | A | A | A | A | A | A | A | A | A | NA | A | A | NA | W | A | A | W | A | A | A | A | A | W | A | W | A | W | A | A | W | W | A | A | A | A | A | A | A | A | W | A | A | A | NA | W | W | A | W | A | W | A | A | W | A | W | W | A | W | A | W | A | A | A | A | A | A | W | W | W | W | W | A | W |
| ChrII | 7.7 | W | W | W | A | A | A | A | NA | W | W | W | A | A | W | W | W | A | W | W | A | A | A | A | A | A | A | A | A | W | A | A | W | W | A | A | W | A | A | A | A | A | W | A | W | A | W | A | A | W | W | A | NA | A | A | A | A | NA | A | W | NA | A | A | W | W | W | NA | W | A | W | NA | A | W | NA | W | W | A | W | A | W | NA | A | A | NA | A | A | W | W | W | W | W | A | W |
| ChrII | 7.8 | W | W | W | A | A | A | A | A | W | W | W | A | A | W | W | W | A | W | W | A | A | A | A | A | A | A | A | A | NA | A | A | W | W | A | A | NA | A | A | A | A | A | W | A | W | A | W | A | A | W | W | A | A | A | A | A | A | A | A | W | A | A | A | W | W | W | A | W | A | W | A | NA | W | A | W | W | A | W | A | W | A | A | A | A | A | A | W | W | W | W | W | A | W |
| ChrII | 7.9 | W | W | W | A | A | A | A | A | W | W | W | A | A | W | W | W | A | W | W | A | A | A | A | A | A | A | A | A | W | A | A | NA | W | A | A | NA | A | A | A | A | A | W | A | W | A | W | A | A | W | NA | A | A | A | A | A | A | A | A | W | A | A | A | W | W | W | A | W | A | W | A | A | W | A | W | W | A | W | A | W | A | A | A | A | A | A | W | W | W | W | W | A | W |
| ChrII | 8 | W | W | W | A | A | A | A | A | W | W | W | A | A | W | W | W | A | W | W | A | A | A | A | A | A | A | A | A | W | A | A | W | W | A | A | W | A | A | A | A | A | W | A | W | A | W | A | A | W | NA | A | A | A | A | A | A | A | A | W | A | A | A | W | W | W | A | W | A | W | A | A | W | A | W | W | A | W | A | W | A | A | A | A | A | A | W | W | W | W | W | A | W |
| ChrII | 8.1 | W | W | W | A | A | A | A | A | W | W | W | A | A | W | W | W | NA | W | W | A | A | A | A | A | A | A | A | A | W | A | A | W | W | A | A | NA | A | A | A | A | A | W | A | W | A | W | A | A | W | W | A | A | A | A | A | A | A | A | W | A | A | A | W | W | W | A | W | A | W | A | A | W | A | W | W | A | W | A | W | A | A | A | A | A | A | W | W | W | W | W | A | W |
| ChrII | 8.2 | W | W | W | A | A | A | A | A | W | W | W | A | A | W | W | W | A | W | W | A | A | A | A | A | A | A | A | A | NA | A | A | NA | W | A | A | NA | A | A | A | A | A | W | A | W | A | W | A | A | W | NA | A | A | A | A | A | A | A | A | W | A | A | A | NA | W | W | A | W | A | W | A | A | W | A | W | W | A | W | A | W | A | A | A | A | A | A | W | W | W | W | W | A | W |
| ChrII | 8.3 | W | W | W | A | A | A | A | A | W | W | W | A | A | W | W | W | A | W | W | A | A | A | A | A | A | A | A | A | NA | A | A | W | W | A | A | NA | A | A | A | A | A | W | A | W | A | W | A | A | W | NA | A | A | A | A | A | A | A | A | W | A | A | A | NA | W | W | A | W | A | W | A | A | W | A | W | W | A | W | A | W | A | A | A | A | A | A | W | W | W | W | W | A | W |
| ChrII | 8.4 | W | W | W | A | A | A | A | A | W | W | W | A | A | W | W | W | A | W | W | A | A | A | A | A | A | A | A | A | W | A | A | W | W | A | A | W | A | A | A | A | A | W | A | W | A | W | A | A | W | NA | A | A | A | A | A | A | A | A | W | A | A | A | NA | W | W | A | W | A | W | A | A | W | A | W | W | A | W | A | W | A | A | A | A | A | A | W | W | W | W | W | A | W |
| ChrII | 8.5 | W | W | W | A | A | A | A | A | W | W | W | A | A | W | W | W | A | W | W | A | A | A | A | A | A | A | A | A | W | A | A | W | W | A | A | W | A | A | A | A | A | W | A | W | A | W | A | A | W | W | A | A | A | A | A | A | A | A | W | A | A | A | W | W | W | A | W | A | W | A | A | W | A | W | W | A | W | A | W | A | A | A | A | A | A | W | W | W | W | W | A | W |
| ChrII | 8.6 | W | W | W | A | A | A | A | A | W | W | W | A | A | W | W | W | A | W | W | A | A | A | A | A | A | A | A | A | W | A | A | W | W | A | A | NA | A | A | A | A | A | W | A | W | A | W | A | A | W | W | A | A | A | A | A | A | A | A | W | A | A | A | NA | W | W | A | W | A | W | A | A | W | A | W | W | A | W | A | W | A | A | A | A | A | A | W | W | W | W | W | A | W |
| ChrII | 8.7 | W | W | W | A | A | A | A | A | W | W | W | A | A | W | W | W | A | W | W | A | A | A | A | A | A | A | A | A | NA | A | A | W | W | W | A | W | A | A | A | A | A | W | A | W | A | W | A | A | W | NA | A | A | A | A | NA | A | A | A | W | A | A | A | W | W | W | A | A | A | W | A | NA | W | A | W | W | A | W | A | W | A | A | A | A | A | A | W | W | W | W | W | A | W |
| ChrII | 8.8 | W | W | W | A | A | A | A | A | W | W | W | A | A | W | W | W | A | W | W | A | A | A | A | A | A | A | A | A | W | A | A | W | W | W | A | W | A | A | A | A | A | W | A | W | A | W | A | A | W | NA | A | A | A | A | A | A | A | A | W | A | A | A | W | W | W | A | A | A | W | A | A | W | A | W | W | A | W | A | W | A | A | A | A | A | A | W | W | W | W | W | A | W |
| ChrII | 8.9 | W | W | W | A | A | A | A | A | W | W | W | A | A | W | W | W | A | W | W | A | A | A | A | A | A | A | A | A | NA | A | A | W | W | W | A | W | A | A | A | A | A | W | A | W | A | W | A | A | W | NA | A | A | A | A | A | A | A | A | W | A | A | A | W | W | W | A | A | A | W | A | A | W | A | W | W | A | W | A | W | A | A | A | A | A | A | W | W | W | W | W | A | W |
| ChrII | 9 | W | W | W | A | A | A | A | A | W | W | W | A | A | W | W | W | A | W | W | A | A | A | A | A | A | A | A | A | W | A | A | W | W | W | A | NA | A | A | A | A | A | W | A | W | A | W | A | A | W | W | A | A | A | A | A | A | A | A | W | A | A | A | W | W | W | A | A | A | W | A | A | W | A | W | W | A | W | A | W | A | A | A | A | A | A | W | W | W | W | W | A | W |
| ChrII | 9.1 | W | W | W | A | A | A | A | A | W | W | W | A | A | W | W | W | A | W | NA | A | A | A | A | A | A | A | A | A | W | A | A | W | W | W | A | NA | A | A | A | A | A | W | A | W | A | W | A | A | W | NA | A | A | A | A | A | A | A | A | W | A | A | A | W | W | W | A | A | A | W | A | A | W | A | W | W | A | W | A | W | A | A | A | A | A | A | W | W | W | W | W | A | W |
| ChrII | 9.2 | W | W | W | A | A | A | A | A | W | W | W | A | A | W | W | W | A | W | W | A | A | A | A | A | A | A | A | A | W | A | A | NA | W | W | A | W | A | A | A | A | A | W | A | W | A | W | A | A | W | NA | A | A | A | A | A | A | A | A | W | A | A | A | W | W | W | A | A | A | W | A | A | W | A | W | W | A | W | A | W | A | A | A | A | A | A | W | W | W | W | W | A | W |
| ChrII | 9.3 | W | W | W | A | A | A | A | A | W | W | W | A | A | W | W | W | A | W | W | A | A | A | A | A | A | A | A | A | W | A | A | W | W | W | A | W | A | A | A | A | A | W | A | W | A | W | A | A | W | W | A | A | A | A | A | A | A | A | W | A | A | A | W | W | W | A | A | A | W | A | A | W | A | W | W | A | W | A | W | A | A | A | A | A | A | W | W | W | W | W | A | W |
| ChrII | 9.4 | W | W | W | A | A | A | A | A | W | W | W | A | A | W | W | W | A | W | W | A | A | A | A | A | A | A | A | A | W | A | A | W | W | W | A | W | A | A | A | A | A | W | A | W | A | W | A | A | W | W | A | A | A | A | A | A | A | A | W | A | A | A | W | W | W | A | A | A | W | A | A | W | A | W | W | A | W | A | W | A | A | A | A | A | A | W | W | W | W | W | A | W |
| ChrII | 9.5 | W | W | W | A | A | A | A | A | W | W | W | A | A | W | W | W | A | W | W | A | A | A | A | A | A | A | A | A | W | A | A | W | W | W | A | W | A | A | A | A | A | W | A | W | A | W | A | A | W | NA | A | A | A | A | A | A | A | A | W | A | A | A | W | W | W | A | A | A | W | A | A | W | A | W | W | A | W | A | W | A | A | A | A | A | A | W | W | W | W | W | A | W |
| ChrII | 9.6 | W | W | W | A | A | A | A | A | W | W | W | A | A | W | W | W | A | W | W | A | A | A | A | A | A | A | A | A | W | A | A | W | W | W | A | W | A | A | A | A | A | W | A | W | A | W | A | A | W | NA | A | A | A | A | A | A | A | A | W | A | A | A | W | W | W | A | A | A | W | A | A | W | A | W | W | A | W | A | W | A | A | A | A | A | A | W | W | W | W | W | A | W |
| ChrII | 9.7 | W | W | W | A | A | A | A | A | W | W | W | A | A | W | W | W | A | W | W | A | A | A | A | A | A | A | A | A | W | A | A | W | W | W | A | W | A | A | A | A | A | W | A | W | A | W | A | A | W | W | A | A | A | A | A | A | A | A | W | A | A | A | W | W | W | A | A | A | W | A | A | W | A | W | W | A | W | A | W | A | A | A | A | A | A | W | W | W | W | W | A | W |
| ChrII | 9.8 | W | W | W | A | A | A | A | A | W | W | W | A | A | W | W | W | A | W | W | A | NA | A | A | A | A | A | A | A | W | A | A | W | W | W | A | W | A | A | A | A | A | W | A | W | A | W | A | A | W | W | A | A | A | A | A | A | A | A | W | A | A | A | W | W | W | A | A | A | W | A | A | W | A | W | W | A | W | A | W | A | A | A | A | A | A | W | W | W | W | W | A | W |
| ChrII | 9.9 | W | W | W | A | A | A | A | A | W | W | W | A | A | W | W | W | A | W | W | A | A | A | A | A | NA | A | A | A | W | A | A | W | W | W | A | W | A | A | A | A | A | W | A | W | A | W | A | A | W | NA | A | A | A | A | A | A | A | A | W | A | A | A | W | W | W | A | A | A | W | A | A | W | A | W | W | A | W | A | W | A | A | A | A | A | A | W | W | W | W | W | A | W |
| ChrII | 10 | W | W | W | A | A | A | A | A | W | W | W | A | A | W | W | W | A | W | W | A | NA | A | A | A | A | A | A | A | W | A | A | W | W | W | A | W | A | A | A | A | A | W | A | W | A | W | A | A | W | W | A | A | A | A | NA | A | A | A | W | A | A | A | W | W | W | A | A | A | W | A | A | W | A | W | W | A | W | A | W | A | A | A | A | A | A | W | W | W | W | W | A | W |
| ChrII | 10.1 | W | W | W | A | A | A | A | A | W | W | W | A | A | W | W | W | A | W | W | A | A | A | A | A | A | A | A | A | W | A | A | W | W | W | A | W | A | A | A | A | A | W | A | W | A | W | A | A | W | W | A | A | A | A | A | A | A | A | W | A | A | A | W | W | W | A | A | A | W | A | A | W | A | W | W | A | W | A | W | A | A | A | A | A | A | W | W | W | W | W | A | W |
| ChrII | 10.2 | W | W | W | A | A | A | A | A | W | W | W | A | A | W | W | W | A | W | W | A | A | A | A | A | A | A | A | A | W | A | A | W | W | W | A | W | A | A | A | A | A | W | A | W | A | W | A | A | W | W | A | A | A | A | A | A | A | A | W | A | A | A | W | W | W | A | A | A | W | A | A | W | A | W | W | A | W | A | W | A | A | A | A | A | A | W | W | W | W | W | A | W |
| ChrII | 10.3 | W | W | W | A | A | A | A | A | W | W | W | A | A | W | W | W | A | W | W | A | A | A | A | A | A | A | A | A | W | A | A | W | W | W | A | NA | A | A | A | A | A | W | A | W | A | W | A | A | W | NA | A | A | A | A | A | A | A | A | W | A | A | A | W | W | W | A | A | A | W | A | A | W | A | W | W | A | W | A | W | A | A | A | A | A | A | W | W | W | W | W | A | W |
| ChrII | 10.4 | W | W | W | A | A | A | A | A | W | W | W | A | A | W | W | W | NA | W | NA | A | A | A | A | A | A | NA | A | A | NA | A | A | NA | W | W | A | NA | A | A | A | A | A | W | A | W | A | W | A | A | W | W | A | A | A | A | A | A | A | A | W | A | A | A | W | W | W | A | A | A | W | A | A | W | A | W | W | A | W | A | W | A | A | A | A | A | A | W | W | W | W | W | A | W |
| ChrII | 10.5 | W | W | W | A | A | A | A | A | W | W | W | A | A | W | W | W | A | W | W | A | A | A | A | A | A | A | A | A | W | A | A | W | W | W | A | W | A | A | A | A | A | W | A | W | A | W | A | A | W | W | A | A | A | A | A | A | A | A | W | A | A | A | W | W | W | A | A | A | W | A | A | W | A | W | W | A | W | A | W | NA | A | A | A | A | A | W | W | W | W | W | A | W |
| ChrII | 10.6 | W | W | W | A | A | A | A | A | W | W | W | A | A | W | W | W | A | W | W | A | A | A | A | A | A | A | A | A | W | A | A | W | W | W | A | W | A | A | A | A | A | W | A | W | A | W | A | A | W | W | A | A | A | A | A | A | A | A | W | A | A | A | W | W | W | A | A | A | W | A | A | W | A | W | W | A | W | A | W | A | A | A | A | A | A | W | W | W | W | W | A | W |
| ChrII | 10.7 | W | W | W | A | A | A | A | A | W | W | W | A | A | W | W | W | A | W | W | A | A | A | A | A | A | A | A | A | W | A | A | W | W | W | A | W | A | A | A | A | A | W | A | W | A | W | A | A | W | W | A | A | A | A | A | A | A | A | W | A | A | A | W | W | W | A | A | A | W | A | A | W | A | W | W | A | W | A | W | A | A | A | A | A | A | W | W | W | W | W | A | W |
| ChrII | 10.8 | W | W | W | A | A | A | A | A | W | W | W | A | A | NA | W | W | A | W | NA | A | A | A | A | A | A | A | A | A | NA | A | A | W | W | W | A | NA | NA | A | A | A | A | W | A | W | A | W | A | A | NA | NA | A | A | A | A | A | A | A | A | W | A | A | A | W | W | W | A | A | A | W | A | A | W | A | W | W | A | W | A | W | A | A | A | A | A | A | W | W | W | W | W | NA | W |
| ChrII | 10.9 | W | W | W | A | A | A | A | A | W | W | W | A | A | W | W | W | A | W | NA | A | A | A | A | A | A | A | A | A | NA | A | A | NA | W | W | A | NA | A | A | A | A | A | W | A | W | A | W | A | A | NA | NA | A | A | A | A | A | A | A | A | W | A | A | A | W | W | W | A | A | A | W | A | A | W | A | W | W | A | W | A | W | A | A | A | A | A | A | W | W | W | W | W | A | W |
| ChrII | 11.1 | W | W | W | A | A | A | A | A | W | W | W | A | A | W | W | W | A | W | W | A | A | A | A | A | A | NA | A | A | W | A | A | NA | W | W | A | NA | A | A | A | A | A | W | A | W | A | W | A | A | W | W | A | A | A | A | A | A | A | A | W | A | A | A | W | W | W | A | A | A | W | A | A | W | A | W | W | A | W | A | W | A | A | A | A | A | A | W | W | W | W | W | A | W |
| ChrII | 11.2 | W | W | W | A | A | A | A | A | W | W | W | A | A | W | W | W | A | W | W | A | A | A | A | A | A | A | NA | A | W | A | A | W | W | W | A | W | A | A | A | A | A | W | A | W | A | W | A | A | W | W | A | A | A | A | A | A | A | A | W | NA | A | A | W | W | W | NA | NA | A | W | A | A | W | A | W | W | A | W | A | W | A | A | A | A | A | A | W | W | W | W | W | A | W |
| ChrII | 11.3 | W | W | W | A | A | NA | NA | A | W | W | W | NA | A | W | W | W | NA | W | W | A | A | A | A | A | NA | A | NA | A | W | A | A | W | NA | W | NA | NA | A | A | A | NA | A | W | A | W | NA | W | NA | NA | W | W | A | A | NA | A | A | A | A | A | W | A | A | A | W | W | W | A | NA | A | W | NA | NA | W | A | W | W | NA | W | NA | W | NA | NA | A | A | A | A | W | W | W | W | W | NA | W |
| ChrII | 11.4 | W | W | W | A | A | A | A | A | W | W | W | A | A | W | W | W | A | W | W | NA | A | A | A | A | A | A | A | A | W | A | A | W | NA | W | A | W | A | A | A | A | A | W | A | W | A | W | A | A | W | W | A | A | A | A | A | A | A | A | W | A | A | A | W | W | W | A | A | A | W | A | A | W | A | W | W | A | W | A | W | A | A | A | A | NA | A | W | W | W | W | W | A | W |
| ChrII | 11.5 | W | W | W | A | A | A | A | A | W | W | W | A | A | NA | W | W | A | W | NA | A | A | A | A | A | A | A | A | A | NA | A | A | NA | W | W | A | NA | A | A | A | A | A | W | A | W | A | W | A | A | W | NA | A | A | A | A | A | A | A | A | W | A | A | A | W | W | W | A | A | A | W | A | NA | W | A | W | W | A | W | A | W | A | A | A | A | A | A | W | W | W | W | W | NA | W |
| ChrII | 11.6 | W | W | W | A | A | A | A | A | W | W | W | A | A | W | W | W | A | W | W | A | A | A | A | A | A | A | A | A | W | A | A | W | W | W | A | NA | A | A | A | A | A | W | A | W | A | W | A | A | W | NA | A | A | A | A | A | A | A | A | W | A | A | A | W | W | W | A | A | A | W | A | A | W | A | W | W | A | W | A | W | A | A | A | A | A | A | W | W | W | W | W | A | W |
| ChrII | 11.7 | W | W | W | A | A | A | A | A | W | W | W | A | A | W | W | W | A | W | NA | A | A | A | A | A | A | A | A | A | W | A | A | W | W | W | A | NA | A | A | A | A | A | W | A | W | A | W | A | A | W | NA | A | A | A | A | A | A | A | A | W | A | A | A | W | W | W | A | A | A | W | A | A | W | A | W | W | A | W | A | W | A | A | A | A | A | A | W | W | W | W | W | A | W |
| ChrII | 11.8 | W | W | W | A | A | A | A | A | W | W | W | A | A | W | W | W | A | W | W | A | A | A | A | A | A | A | A | A | W | A | A | W | W | W | A | W | A | A | A | A | A | W | A | W | A | W | A | A | W | NA | A | A | A | A | A | A | A | A | W | A | A | A | W | W | W | A | A | A | W | A | A | W | A | W | W | A | W | A | W | A | A | A | A | A | A | W | W | W | W | W | A | W |
| ChrII | 11.9 | W | W | W | A | A | A | A | A | W | W | W | A | A | W | W | W | A | W | W | A | A | A | A | A | A | A | A | A | W | A | A | W | W | W | A | W | A | A | A | A | A | W | A | W | A | W | A | A | W | NA | A | A | A | A | A | A | A | A | W | A | A | A | W | W | W | A | A | A | W | A | A | W | A | W | W | A | W | A | W | A | A | A | A | A | A | W | W | W | W | W | A | W |
| ChrII | 12 | W | W | W | A | A | A | A | A | W | W | W | A | A | W | W | W | A | W | NA | A | A | A | A | A | A | A | A | A | W | A | A | W | W | W | A | NA | A | A | A | A | A | W | A | W | A | W | A | A | W | W | A | A | A | A | A | A | A | A | W | A | A | A | W | W | W | A | A | A | W | A | A | W | A | W | W | A | W | A | W | A | A | A | A | A | A | W | W | W | W | W | A | W |
| ChrII | 12.1 | W | W | W | A | A | A | A | A | W | W | W | A | A | W | W | W | A | W | W | A | A | A | A | A | A | A | A | A | NA | A | A | W | W | W | A | NA | A | A | A | A | A | W | A | W | A | W | A | A | W | NA | A | A | A | A | A | A | A | A | W | A | A | A | W | W | W | A | A | A | W | A | A | W | A | W | W | A | W | A | W | A | A | A | A | A | A | W | W | W | W | W | A | W |
| ChrII | 12.2 | W | W | W | A | A | A | A | A | W | W | W | A | A | W | W | W | A | W | W | A | A | A | A | A | A | A | A | NA | W | A | A | W | W | W | A | W | A | A | A | A | A | W | A | W | A | W | A | A | W | W | A | A | A | A | A | A | A | A | W | A | A | A | W | W | W | A | A | A | W | A | NA | W | A | W | W | A | W | A | W | A | A | A | A | A | A | W | W | W | W | W | A | W |
| ChrII | 12.3 | W | W | W | A | A | A | A | A | W | W | W | A | A | W | W | W | A | W | W | NA | NA | A | NA | A | A | A | A | A | W | A | A | W | W | W | A | W | A | A | A | A | A | W | A | W | A | W | A | A | W | W | A | A | A | A | A | A | A | A | W | A | A | A | W | W | W | A | NA | A | W | A | A | W | A | W | W | A | W | A | W | A | A | NA | A | A | A | W | W | W | W | W | A | W |
| ChrII | 12.4 | W | W | W | A | A | A | A | A | W | W | W | A | A | W | W | W | A | W | W | A | A | A | A | A | A | A | A | A | W | A | A | W | W | W | A | W | A | A | A | A | A | W | A | W | A | W | A | A | W | W | A | A | A | A | A | A | A | A | W | A | A | A | W | W | W | A | A | A | W | A | A | W | A | W | W | A | W | A | W | A | A | A | A | A | A | W | W | W | W | W | A | W |
| ChrII | 12.5 | W | W | W | A | A | A | A | A | W | W | W | A | A | W | W | W | A | W | W | A | A | A | A | A | A | A | NA | A | W | A | A | W | W | W | A | W | A | A | A | A | A | W | A | W | A | W | A | A | W | W | A | A | A | A | A | A | A | A | W | A | A | A | W | W | W | A | A | A | W | A | A | W | A | W | W | A | W | A | W | A | A | A | A | A | A | W | W | W | W | W | A | W |
| ChrII | 12.6 | W | W | W | A | A | A | A | A | W | W | W | A | A | W | W | W | A | W | W | A | A | A | A | A | A | A | A | A | W | A | A | NA | W | W | A | NA | A | A | A | A | A | W | A | W | A | W | A | A | W | W | A | A | A | A | A | A | A | A | W | A | A | A | W | W | W | A | A | A | W | A | A | W | A | W | W | A | W | A | W | A | A | A | A | A | A | W | W | W | W | W | A | W |
| ChrII | 12.7 | W | W | W | A | A | A | A | A | W | W | W | A | A | W | W | W | NA | W | NA | A | A | A | A | A | A | NA | A | A | W | NA | A | NA | W | W | A | NA | A | A | A | A | A | W | A | W | A | W | A | A | W | W | A | A | A | A | A | A | A | A | W | A | A | A | W | W | W | A | A | NA | W | A | A | W | A | W | W | A | W | A | W | A | A | A | A | A | A | W | W | W | W | W | A | W |
| ChrII | 12.8 | W | W | W | A | A | A | A | A | W | W | W | A | A | W | W | W | A | W | NA | A | A | A | A | A | A | A | A | A | NA | A | A | W | W | W | A | NA | A | A | A | A | A | W | A | W | A | W | A | A | W | W | A | A | A | A | A | A | A | A | W | A | A | A | W | W | W | A | A | A | W | A | A | W | A | W | W | A | W | A | W | A | A | A | A | A | A | W | W | W | W | W | A | W |
| ChrII | 12.9 | W | W | W | A | A | A | A | A | W | W | W | A | A | W | W | W | NA | W | W | A | A | A | A | A | A | A | A | A | NA | A | A | W | W | W | A | NA | A | A | A | A | A | W | A | W | A | W | A | A | W | NA | A | A | A | A | A | A | A | A | W | A | A | A | W | W | W | A | A | A | W | A | A | W | A | W | W | A | W | A | W | A | A | A | A | A | A | W | W | W | W | W | A | W |
| ChrII | 13 | W | W | W | A | A | A | A | A | W | W | W | A | A | W | NA | W | A | W | NA | A | A | A | NA | A | A | A | A | A | W | A | A | W | NA | W | A | NA | A | A | A | A | A | W | A | W | A | W | A | A | W | NA | A | A | A | A | A | A | A | A | W | A | A | A | W | W | W | A | A | A | W | A | A | W | A | W | W | A | W | A | W | A | A | A | NA | A | A | W | W | W | W | W | A | W |
| ChrII | 13.1 | W | W | W | A | A | A | A | A | W | W | W | A | A | W | W | W | A | W | NA | A | A | A | A | A | A | A | A | A | NA | A | A | W | W | W | A | NA | A | A | A | A | A | W | A | W | A | W | A | A | W | W | A | A | A | A | A | A | A | A | W | A | A | A | W | W | W | A | A | A | W | A | A | W | A | W | W | A | W | A | W | A | A | A | A | A | A | W | W | W | W | W | A | W |
| ChrII | 13.3 | W | W | W | A | A | A | A | A | W | W | W | A | A | W | W | W | A | W | W | A | A | A | A | A | A | A | A | A | NA | A | A | NA | W | W | A | NA | A | A | A | A | A | W | A | W | A | W | A | A | W | NA | A | A | A | A | NA | A | A | A | W | A | A | A | W | W | W | A | A | A | W | A | A | W | A | W | W | A | W | A | W | A | A | A | A | A | A | W | W | W | W | W | A | W |
| ChrII | 13.4 | W | W | W | A | A | A | A | A | W | W | W | A | A | W | W | W | A | W | NA | A | A | A | A | A | A | A | A | A | NA | A | A | W | W | W | A | NA | A | A | A | A | A | W | A | W | A | W | A | A | W | W | A | A | A | A | A | A | A | A | W | A | A | A | W | W | W | A | A | A | W | A | A | W | A | W | W | A | W | A | W | A | A | A | A | A | A | W | W | W | W | W | A | W |
| ChrII | 13.5 | W | W | W | A | A | A | A | A | W | W | W | A | A | W | W | W | A | W | W | A | A | A | A | A | A | A | A | A | W | A | A | W | W | W | A | W | A | A | A | A | A | W | A | W | A | W | A | A | W | W | A | A | A | A | A | A | A | A | W | A | A | A | W | W | W | A | A | A | W | A | A | W | A | W | W | A | W | A | W | A | A | A | A | A | A | W | W | W | W | W | A | W |
| ChrII | 13.6 | W | W | W | A | A | A | A | A | W | W | W | A | A | W | W | W | A | W | W | A | A | A | A | A | A | A | A | A | W | A | A | W | W | W | A | W | A | A | A | A | A | W | A | W | A | W | A | A | W | NA | A | A | A | A | A | A | A | A | W | A | A | A | W | W | W | A | A | A | W | A | A | W | A | W | W | A | W | A | W | A | A | A | A | A | A | W | W | W | W | W | A | W |
| ChrII | 13.7 | W | W | W | A | A | A | A | A | W | W | W | A | A | W | W | W | A | W | W | A | A | A | A | A | A | A | A | A | NA | A | A | W | W | W | A | NA | A | A | A | A | A | W | A | W | A | W | A | A | W | W | A | A | A | A | A | A | A | A | W | A | A | A | W | W | W | A | A | A | W | A | A | W | A | W | W | A | W | A | W | A | A | A | A | A | A | W | W | W | W | W | A | W |
| ChrII | 13.8 | W | W | W | A | A | A | A | A | W | W | W | A | A | W | W | W | A | W | W | A | A | A | A | A | A | A | A | A | W | A | A | W | W | W | A | NA | A | A | A | A | A | W | A | W | A | W | A | A | W | W | A | A | A | A | NA | A | A | A | W | A | A | A | W | W | W | A | A | A | W | A | A | W | A | W | W | A | W | A | W | A | A | A | A | A | A | W | W | W | W | W | A | W |
| ChrII | 13.9 | W | W | W | A | A | A | A | A | W | W | W | A | A | W | W | W | A | W | W | A | A | A | A | A | A | A | A | A | W | A | A | NA | W | W | A | W | A | A | A | A | A | W | A | W | A | W | A | A | W | NA | A | A | A | A | A | A | A | A | W | A | A | A | W | W | W | A | A | A | W | A | A | W | A | W | W | A | W | A | W | A | A | A | A | A | A | W | W | W | W | W | A | W |
| ChrII | 14 | W | W | W | A | A | A | A | A | W | W | W | A | A | W | W | W | A | W | NA | A | A | A | A | A | A | A | A | A | NA | A | A | NA | W | W | A | NA | A | A | A | A | A | W | A | W | A | W | A | A | W | NA | A | A | A | A | A | A | A | A | W | A | A | A | W | W | W | A | A | A | W | A | A | W | A | W | W | A | W | A | W | A | A | A | A | A | A | W | W | W | W | W | A | W |
| ChrII | 14.1 | W | W | W | A | A | A | A | A | W | W | W | A | A | W | W | W | A | W | W | A | A | A | A | A | A | A | A | A | NA | A | A | W | W | W | A | NA | A | A | A | A | A | W | A | W | A | W | A | A | W | NA | A | A | A | A | A | A | A | A | W | A | A | A | W | W | W | A | A | A | W | A | A | W | A | W | W | A | W | A | W | A | A | A | A | A | A | W | W | W | W | W | A | W |
| ChrII | 14.2 | W | W | W | A | A | A | A | A | W | W | W | A | A | W | W | W | A | W | W | A | A | A | A | A | A | A | A | A | W | A | A | W | W | W | A | W | A | A | A | A | A | W | A | W | A | W | A | A | W | NA | A | A | A | A | A | A | A | A | W | A | A | A | W | W | W | A | A | A | W | A | A | W | A | W | W | A | W | A | W | A | A | A | A | A | A | W | W | W | W | W | A | W |
| ChrII | 14.3 | W | W | W | A | A | A | A | A | W | W | W | A | A | W | W | W | A | W | W | A | A | A | A | A | A | A | A | A | NA | A | A | NA | W | W | A | NA | A | A | A | A | A | W | A | W | A | W | A | A | W | NA | A | A | A | A | A | A | A | A | W | A | A | A | W | W | W | A | A | A | W | A | A | W | A | W | W | A | W | A | W | A | A | A | A | A | A | W | W | W | W | W | A | W |
| ChrII | 14.4 | W | W | W | A | A | A | A | A | W | W | W | A | A | W | W | W | A | W | W | A | A | A | A | A | A | A | A | A | W | A | A | W | W | W | A | W | A | A | A | A | A | W | A | W | A | W | A | A | W | W | A | A | A | A | A | A | A | A | W | A | A | A | W | W | W | A | A | A | W | A | A | W | A | W | W | A | W | A | W | A | A | A | A | A | A | W | W | W | W | W | A | W |
| ChrII | 14.5 | W | W | W | A | A | A | A | A | W | W | W | A | A | W | W | W | A | W | W | A | A | A | A | A | A | A | A | A | W | A | A | W | W | W | A | W | A | A | A | A | A | W | A | W | A | W | A | A | W | NA | A | A | A | A | A | A | A | A | W | A | A | A | W | W | W | A | A | A | W | A | A | W | A | W | W | A | W | A | W | A | A | A | W | A | A | W | W | W | W | W | A | W |
| ChrII | 14.6 | W | W | W | A | A | A | A | A | W | W | W | A | A | W | W | W | A | W | W | A | A | A | A | A | A | A | A | A | W | A | A | W | W | W | A | W | A | A | A | A | A | W | A | W | A | W | A | A | W | W | A | A | A | A | A | A | A | A | W | A | A | A | W | W | W | A | A | A | W | A | A | W | A | W | W | A | W | A | W | A | A | A | W | A | A | W | W | W | W | W | A | W |
| ChrII | 14.7 | W | W | W | A | A | A | A | A | W | W | W | A | A | W | W | W | NA | W | W | A | A | A | A | A | A | A | A | A | NA | A | A | NA | W | W | A | NA | NA | A | A | A | A | W | A | W | A | W | A | A | W | W | A | A | A | A | A | A | A | A | W | A | A | A | W | W | W | A | A | A | W | A | A | W | A | W | W | A | W | A | W | A | A | A | W | A | A | W | W | W | W | W | A | W |
| ChrII | 14.8 | W | W | W | A | A | A | A | A | W | W | W | A | A | W | W | W | A | W | NA | A | NA | A | A | A | A | A | A | A | W | A | A | W | W | W | A | NA | A | A | A | A | A | W | A | W | A | W | A | A | W | NA | A | A | A | A | A | A | A | A | W | A | A | A | W | W | W | A | A | A | W | A | A | W | A | W | W | A | W | A | W | A | A | A | W | A | A | W | W | W | W | W | A | W |
| ChrII | 14.9 | W | W | W | A | A | A | A | A | W | W | W | A | A | W | W | W | A | W | W | A | A | A | A | A | A | A | A | A | NA | A | A | W | W | W | A | NA | A | A | A | A | A | W | A | W | A | W | A | A | W | NA | A | A | A | A | A | A | A | A | W | A | A | A | W | W | W | A | A | A | W | A | A | W | A | W | W | A | W | A | W | A | A | A | W | A | A | W | W | W | W | W | A | W |
| ChrII | 15 | W | W | W | A | A | A | A | A | W | W | W | A | A | W | W | W | A | W | W | A | A | A | NA | A | A | A | A | A | NA | A | A | W | W | W | A | W | A | A | A | A | A | W | A | W | A | W | A | A | W | W | A | A | A | A | A | A | A | A | W | A | A | A | W | W | W | NA | A | A | W | A | A | W | A | W | W | A | W | A | W | A | A | A | W | A | A | W | W | W | W | W | A | W |
| ChrII | 15.1 | W | W | W | A | A | A | A | A | W | W | W | A | A | W | W | W | A | W | W | A | A | A | A | A | A | A | A | A | W | A | A | W | W | W | A | W | A | A | A | A | A | W | A | W | A | W | A | A | W | W | A | A | A | A | A | A | A | A | W | A | A | A | W | W | W | A | A | A | W | A | A | W | A | W | W | A | W | A | W | A | A | A | W | A | A | W | W | W | W | W | A | W |
| ChrII | 15.2 | W | W | W | A | A | A | A | A | W | W | W | A | A | W | W | W | A | W | W | A | NA | A | A | A | A | A | A | A | W | A | A | W | W | W | A | W | A | A | A | A | A | W | A | W | A | W | A | A | W | NA | A | A | A | A | A | A | A | A | W | A | A | A | W | W | W | A | A | A | W | A | A | W | A | W | W | A | W | A | W | A | A | A | W | A | A | W | W | W | W | W | A | W |
| ChrII | 15.3 | W | W | W | A | A | A | A | A | W | W | W | A | A | W | W | W | A | W | W | A | A | A | A | A | A | A | A | A | W | A | A | W | W | W | A | W | A | A | A | A | A | W | A | W | A | W | A | A | W | W | A | A | A | A | A | A | A | A | W | A | A | A | W | W | W | A | A | A | W | A | A | W | A | W | W | A | W | A | W | A | A | A | W | A | A | W | W | W | W | W | A | W |

| | | | | | | | | | | | | | | | | | | | | | | | | | | | | | | | | | | | | | | | | | | | | | | | | | | | | | | | | | | | | | | | | | | | | | | | | | | | | | | | | | | | | | | | | | | | | | |
|---|---|---|---|---|---|---|---|---|---|---|---|---|---|---|---|---|---|---|---|---|---|---|---|---|---|---|---|---|---|---|---|---|---|---|---|---|---|---|---|---|---|---|---|---|---|---|---|---|---|---|---|---|---|---|---|---|---|---|---|---|---|---|---|---|---|---|---|---|---|---|---|---|---|---|---|---|---|---|---|---|---|---|---|---|---|---|---|---|---|---|---|---|---|
| ChrII | 15.4 | W | W | W | A | A | A | A | A | W | W | W | A | A | W | W | W | A | W | W | A | A | A | A | A | A | A | A | A | NA | A | A | W | W | W | A | W | A | A | A | A | A | W | A | W | A | W | A | A | W | NA | A | A | A | A | A | A | A | A | W | A | A | A | W | W | W | A | A | A | W | A | A | W | A | W | W | A | W | A | W | A | A | A | A | W | A | W | W | W | W | W | A | W |
| ChrII | 15.5 | W | W | W | A | A | A | A | A | W | W | W | A | A | W | W | W | A | W | W | A | A | A | A | A | A | A | A | A | W | A | A | W | W | W | A | W | A | A | A | A | A | W | A | W | A | W | A | A | W | W | A | A | A | A | A | A | A | A | W | A | A | A | W | W | W | A | A | A | W | A | A | W | A | W | W | A | W | A | W | A | A | A | A | W | A | W | W | W | W | W | A | W |
| ChrII | 15.6 | W | W | W | A | A | A | NA | A | W | W | W | A | A | W | W | W | A | W | W | A | A | A | A | A | A | A | A | A | W | A | A | W | W | W | A | W | A | A | A | A | A | W | A | W | A | W | A | A | W | W | A | A | A | A | A | A | A | A | W | A | A | A | W | W | W | A | A | A | W | A | A | W | A | W | W | A | W | A | W | A | A | A | A | W | A | W | NA | W | W | W | A | W |
| ChrII | 15.7 | W | W | W | A | A | A | A | A | W | W | W | A | A | W | W | W | A | W | W | A | A | A | A | A | A | A | A | A | W | A | A | W | W | W | A | W | A | A | A | A | A | W | A | W | A | W | A | A | W | NA | A | A | A | A | A | A | A | A | W | A | A | A | W | W | W | A | A | A | W | A | A | W | A | W | W | A | W | A | W | A | A | A | A | W | A | W | W | W | W | W | A | W |
| ChrII | 15.8 | W | W | W | A | A | A | A | A | W | W | W | A | A | W | W | W | A | W | W | A | A | A | A | A | A | A | A | A | W | A | A | W | W | W | A | W | A | A | A | A | A | W | A | W | A | W | A | A | W | W | A | A | A | A | NA | A | A | A | W | A | A | A | W | W | W | A | NA | A | W | A | A | W | A | W | W | A | W | A | W | A | A | A | A | W | A | W | W | W | W | W | A | W |
| ChrII | 15.9 | W | W | W | A | A | A | A | A | W | W | W | A | A | W | W | W | NA | W | W | A | A | A | A | A | A | A | A | A | W | A | A | W | W | W | A | W | A | A | A | A | A | W | A | W | A | W | A | A | W | W | A | A | A | A | A | A | A | A | W | A | A | A | W | W | W | A | A | A | W | A | A | W | A | W | W | A | W | A | W | A | A | A | A | W | A | W | W | W | W | W | A | W |
| ChrII | 16 | W | W | W | A | A | A | A | A | W | W | W | A | A | W | W | W | A | W | W | A | A | A | A | A | A | A | A | A | W | A | A | W | W | W | A | W | NA | A | A | A | A | W | A | W | A | W | A | A | W | W | A | A | A | A | A | A | A | A | W | A | A | A | W | W | W | A | A | A | W | A | A | W | A | W | W | A | W | A | W | A | A | A | A | W | A | W | W | W | W | W | A | W |
| ChrII | 16.1 | W | W | W | A | A | A | A | A | W | W | W | A | A | W | W | W | A | W | W | A | A | A | A | A | A | A | A | A | W | A | A | W | W | W | A | NA | A | A | A | A | A | W | A | W | A | W | A | A | W | NA | A | A | A | A | A | A | A | A | W | A | A | A | W | W | W | A | A | A | W | A | A | W | A | W | W | A | W | A | W | A | A | A | A | W | A | W | W | W | W | W | A | W |
| ChrII | 16.2 | W | W | W | A | A | A | A | A | NA | W | W | A | A | W | NA | W | A | W | W | A | A | A | A | A | NA | A | A | A | W | A | A | W | W | W | A | W | A | A | A | A | A | W | A | W | A | W | A | A | W | NA | A | A | A | A | A | A | A | A | W | A | A | A | W | W | W | A | A | A | W | A | A | W | A | W | W | A | W | A | W | A | A | A | A | W | A | W | W | W | W | W | A | W |
| ChrII | 16.3 | W | W | W | A | A | A | A | A | A | W | W | A | A | W | W | W | A | W | NA | A | A | A | A | A | W | A | A | A | W | A | A | NA | W | W | A | NA | A | A | A | A | A | W | A | W | A | W | A | A | W | NA | A | A | A | A | NA | A | A | A | W | A | A | W | W | W | W | A | A | A | W | A | A | W | A | W | W | A | W | A | W | A | A | A | A | W | A | W | W | W | W | W | A | W |
| ChrII | 16.4 | W | W | W | A | A | A | A | A | A | W | W | A | A | W | W | W | A | W | W | A | A | A | A | A | W | A | A | A | W | A | A | W | W | W | A | W | A | A | A | A | A | W | A | W | A | W | A | A | W | NA | A | A | A | A | A | A | A | A | W | A | A | W | W | W | W | A | A | A | W | A | A | W | A | W | W | A | W | A | W | A | A | A | A | W | A | W | W | W | W | W | A | W |
| ChrII | 16.5 | W | W | W | A | A | A | A | A | A | NA | W | A | A | W | W | W | A | W | W | A | A | A | A | A | W | A | A | A | W | A | A | W | W | W | A | W | A | A | A | A | A | W | A | W | A | W | A | A | W | NA | A | A | A | A | A | A | A | A | W | A | A | W | W | W | W | A | A | A | W | A | A | W | A | W | W | A | W | A | W | A | A | A | A | W | A | W | W | W | W | W | A | W |
| ChrII | 16.6 | W | W | W | A | A | A | A | A | A | W | W | A | A | W | W | W | A | W | W | A | A | A | A | A | W | A | A | A | W | A | A | W | W | W | A | W | A | A | A | A | A | W | A | W | A | W | A | A | W | W | A | A | A | A | A | A | A | A | W | A | A | W | W | W | W | A | A | A | W | A | A | W | A | W | W | A | W | A | W | A | A | A | A | W | A | W | W | W | W | W | A | W |
| ChrII | 16.7 | W | W | W | A | A | A | A | A | A | W | W | A | A | W | W | W | A | W | W | A | A | A | A | A | W | A | A | A | W | A | A | W | W | W | A | W | A | A | A | A | A | W | A | W | A | W | A | A | W | NA | A | A | A | A | A | A | A | A | W | NA | A | W | W | W | W | A | A | A | W | A | A | W | A | W | W | A | W | A | W | A | A | A | A | W | A | W | W | W | W | W | A | W |
| ChrII | 16.8 | W | W | W | A | A | NA | NA | NA | A | W | W | A | A | W | W | W | NA | W | NA | A | A | A | A | A | W | NA | A | A | W | NA | A | NA | W | W | A | NA | NA | A | A | NA | A | W | A | W | A | W | A | A | W | NA | A | A | NA | A | NA | A | A | A | W | NA | A | W | W | W | W | NA | NA | NA | W | NA | A | W | A | W | W | A | W | A | W | NA | A | A | NA | W | A | W | W | W | W | W | NA | W |
| ChrII | 16.9 | W | W | W | A | A | A | A | A | A | NA | W | NA | A | W | W | W | A | W | NA | A | A | A | A | A | W | A | A | A | NA | A | A | W | W | W | A | NA | A | A | A | A | A | W | A | W | A | W | A | A | W | NA | A | A | A | A | A | A | A | A | W | NA | A | W | W | W | NA | A | A | A | W | A | A | W | A | W | W | A | W | A | W | A | NA | A | A | W | A | W | W | W | W | W | A | W |
| ChrII | 17 | W | W | W | A | A | A | A | A | A | W | W | W | A | W | W | W | A | W | W | A | A | A | A | A | W | A | A | A | W | A | A | W | W | W | A | W | A | A | A | A | A | W | A | W | A | W | A | A | W | W | A | A | A | A | A | A | A | A | W | A | A | W | W | W | W | A | A | A | W | A | A | W | A | W | W | A | W | A | W | A | W | A | A | W | A | W | W | W | W | W | A | W |
| ChrII | 17.1 | W | W | W | A | A | A | A | A | A | W | W | W | A | W | W | W | A | W | W | A | A | A | A | A | W | A | A | A | W | A | A | NA | W | W | A | NA | A | A | A | A | A | W | A | W | A | W | A | A | W | NA | A | A | A | A | A | A | A | A | W | A | A | W | W | W | NA | A | A | A | W | A | A | W | A | W | W | A | W | A | W | A | W | A | A | W | A | W | W | W | W | W | A | W |
| ChrII | 17.2 | W | W | W | A | A | A | A | A | A | W | W | W | A | W | NA | W | A | W | W | A | A | A | A | A | W | A | A | A | NA | A | A | NA | W | W | A | NA | A | A | A | A | A | W | A | W | A | W | A | A | W | NA | A | A | A | A | A | A | A | A | W | A | A | W | W | W | NA | A | A | A | W | A | A | W | A | W | W | A | W | A | W | A | W | A | A | W | A | W | W | W | W | W | A | W |
| ChrII | 17.4 | W | W | W | A | A | A | A | A | A | W | W | W | A | W | W | W | A | W | W | A | A | A | A | A | W | A | NA | A | A | A | A | W | W | W | A | W | A | A | A | A | A | W | A | W | A | W | A | W | W | W | A | NA | A | A | A | A | A | A | W | A | A | W | W | W | NA | A | A | A | W | A | A | W | A | W | W | A | W | A | W | A | W | A | A | W | A | W | W | W | W | W | A | W |
| ChrII | 17.5 | W | W | W | A | A | A | A | A | A | W | W | W | A | W | W | W | A | W | W | A | A | A | A | A | W | A | W | A | A | A | A | W | W | W | A | W | A | A | A | A | A | W | A | W | A | W | A | W | W | W | A | A | A | A | A | A | A | A | W | A | A | W | W | W | W | A | A | W | W | A | A | W | A | W | W | A | W | A | W | A | W | A | A | W | A | W | W | W | W | W | A | W |
| ChrII | 17.6 | W | W | W | A | A | A | A | A | A | W | W | W | A | NA | NA | W | A | W | W | A | A | A | A | A | W | A | W | A | A | A | A | W | W | W | A | NA | A | A | A | A | A | W | A | W | A | W | A | W | W | NA | A | A | A | A | A | A | A | A | W | A | A | W | W | W | NA | A | A | W | W | A | A | W | A | W | W | A | W | A | W | A | W | A | A | W | A | W | W | W | W | W | A | W |
| ChrII | 17.7 | W | W | W | A | A | A | A | A | A | NA | W | W | A | W | NA | W | A | W | W | A | A | A | A | A | W | A | W | A | A | A | A | NA | W | W | A | NA | NA | A | A | A | A | W | A | W | A | W | A | W | W | W | A | A | A | A | A | A | A | A | W | A | A | W | W | W | NA | A | A | NA | W | A | A | W | A | W | W | A | W | A | W | A | W | A | A | W | A | W | W | W | W | W | A | W |
| ChrII | 17.8 | W | W | W | A | A | A | A | A | A | W | W | W | A | W | W | W | A | W | W | A | A | A | A | A | W | A | W | A | A | A | A | W | W | W | A | W | A | A | A | A | A | W | A | W | A | W | A | W | W | NA | A | A | A | A | A | A | A | A | W | A | A | W | W | W | W | A | A | W | W | A | A | W | A | W | W | A | W | A | W | W | W | A | A | W | A | W | W | W | W | W | A | W |
| ChrII | 17.9 | W | W | W | A | A | A | A | A | A | NA | W | W | A | W | W | W | A | W | W | A | A | A | A | A | W | A | W | A | A | A | A | W | W | W | A | NA | A | A | A | A | A | W | A | W | A | W | A | W | W | NA | A | A | A | A | A | A | A | A | W | A | A | W | W | W | NA | A | A | NA | W | A | A | W | A | W | W | A | W | A | W | W | W | A | A | W | A | W | W | W | W | W | A | W |
| ChrII | 18 | W | W | W | A | A | A | A | A | A | W | W | W | A | W | W | W | A | W | W | A | A | A | A | A | W | A | W | A | A | A | A | W | W | W | A | W | A | A | A | A | A | W | A | W | A | W | A | W | W | W | A | A | A | A | NA | A | A | A | W | A | A | W | W | W | NA | A | A | W | W | A | A | W | A | W | W | A | W | A | W | W | W | A | A | W | A | W | W | W | W | W | A | W |
| ChrII | 18.1 | W | W | W | A | A | A | A | A | A | NA | W | W | A | W | W | W | A | W | W | A | A | A | A | A | W | NA | W | A | A | A | A | W | W | W | A | NA | A | A | A | A | A | W | A | W | A | W | A | W | W | W | A | A | A | A | A | A | A | A | W | A | A | W | W | W | NA | A | A | NA | W | A | A | W | A | W | W | A | W | A | W | W | W | A | A | W | A | W | W | W | W | W | A | W |
| ChrII | 18.2 | W | W | W | A | A | A | A | A | A | W | W | W | A | W | W | W | A | W | W | A | A | A | A | A | W | A | W | A | A | A | A | W | W | W | A | W | W | A | A | A | A | W | A | W | A | W | A | W | W | NA | NA | A | A | A | A | A | A | A | W | A | A | W | W | W | W | A | A | W | W | A | A | W | A | W | W | A | W | A | W | W | W | A | A | W | A | W | W | W | W | W | A | W |
| ChrII | 18.3 | W | W | W | A | A | A | A | A | A | W | W | W | A | NA | NA | W | A | W | W | A | A | A | A | A | W | A | W | A | A | A | A | W | W | W | A | NA | W | A | A | A | A | W | A | W | A | W | A | W | W | W | W | A | A | A | A | A | A | A | W | A | A | W | W | W | NA | A | A | W | W | A | A | W | A | W | W | A | W | A | W | W | W | A | A | W | A | W | W | W | W | W | A | W |
| ChrII | 18.4 | W | W | W | A | A | A | A | A | A | W | W | W | A | W | NA | W | A | W | W | A | A | A | A | A | W | A | W | A | A | A | A | W | W | W | A | A | W | A | A | A | A | W | A | W | A | W | A | W | W | W | W | A | A | A | A | A | A | A | W | A | A | W | W | W | W | A | A | W | W | A | A | W | A | W | W | A | W | A | W | W | W | A | A | W | A | W | W | W | W | W | A | W |
| ChrII | 18.5 | W | W | W | A | A | A | A | A | A | W | W | W | A | W | W | W | A | W | W | NA | A | A | A | A | W | A | W | A | A | A | A | W | W | W | A | A | W | A | A | A | A | W | A | W | A | W | A | W | W | W | W | A | A | A | A | A | A | A | W | A | A | W | W | W | W | A | A | W | W | A | NA | W | NA | W | W | A | W | A | W | W | W | A | A | W | A | W | W | W | W | W | A | W |
| ChrII | 18.6 | W | W | W | A | A | A | A | A | A | W | W | W | A | W | W | W | NA | W | W | NA | A | W | NA | A | W | A | W | A | A | A | A | W | W | W | A | A | W | A | A | A | A | W | A | W | A | W | A | W | W | W | W | A | A | A | A | A | A | A | W | NA | A | W | W | W | NA | A | A | W | W | A | A | W | A | W | W | A | W | A | W | W | W | A | A | W | A | W | W | W | W | W | A | W |
| ChrII | 18.7 | W | W | W | A | A | A | A | A | A | W | W | W | A | W | W | W | A | W | W | A | A | W | A | A | W | A | W | A | A | A | A | W | W | W | A | A | W | A | A | A | A | W | A | W | A | W | A | W | W | NA | W | A | A | A | A | A | A | A | W | A | A | W | W | W | W | A | A | W | W | A | A | W | A | W | W | A | W | A | W | W | W | A | A | W | A | W | W | W | W | W | A | W |
| ChrII | 18.8 | W | W | W | A | A | A | A | A | A | W | NA | W | A | W | W | W | A | W | W | A | A | W | A | A | W | A | W | A | A | A | A | W | W | W | A | A | W | A | A | A | A | W | A | W | A | W | A | W | W | W | W | A | A | A | A | A | A | A | W | A | A | W | W | W | W | A | A | W | W | A | A | W | A | W | W | A | W | A | W | W | W | A | A | W | A | W | W | W | W | W | A | W |
| ChrII | 18.9 | W | W | W | A | A | A | A | A | A | W | A | W | A | W | NA | W | A | W | W | A | A | W | A | A | W | A | W | A | A | A | A | NA | W | W | A | A | W | A | A | A | A | W | A | W | A | W | A | W | W | NA | W | A | A | A | A | A | A | A | W | A | A | W | W | W | NA | A | A | W | W | A | A | W | A | W | W | A | W | A | W | W | W | A | A | W | A | W | W | W | W | W | A | W |
| ChrII | 19 | W | W | W | A | A | A | A | A | NA | W | NA | W | A | W | NA | W | A | W | W | A | A | W | NA | A | W | A | W | A | A | A | NA | NA | W | W | A | A | NA | NA | NA | A | A | W | NA | W | A | W | NA | W | W | W | W | A | A | A | A | A | A | A | W | A | NA | W | W | W | NA | A | NA | W | W | NA | A | W | A | W | W | NA | W | A | W | W | W | A | NA | W | NA | W | W | W | W | W | A | W |
| ChrII | 19.1 | W | W | W | A | A | A | A | A | A | W | A | W | A | W | W | W | A | W | W | A | A | W | A | A | W | A | W | A | A | A | A | W | W | W | A | A | W | A | A | A | A | W | A | W | A | W | A | W | W | W | W | A | A | A | A | A | A | A | W | A | A | W | W | W | W | A | A | W | W | A | A | W | A | W | W | A | W | A | W | W | W | A | A | W | A | W | W | W | W | W | A | W |
| ChrII | 19.2 | W | W | W | A | A | A | A | A | A | W | A | W | A | W | NA | W | A | W | W | A | A | W | A | A | W | A | W | A | A | A | A | W | W | W | A | A | W | A | A | A | NA | W | A | W | A | NA | A | W | W | W | W | A | A | A | A | A | A | A | W | NA | A | W | W | W | NA | A | A | W | W | A | A | W | A | W | W | A | W | A | W | W | W | A | A | W | A | W | W | W | W | W | A | W |
| ChrII | 19.3 | W | W | W | A | A | A | A | A | A | W | A | W | A | W | W | W | A | W | W | A | A | W | A | A | NA | A | W | A | A | A | A | W | W | W | A | A | W | A | A | A | W | W | A | W | A | A | A | W | W | NA | W | A | A | A | A | A | A | A | W | W | A | W | W | W | NA | A | A | W | W | A | A | W | A | W | W | A | W | A | W | W | W | A | A | W | A | W | W | W | W | W | A | W |
| ChrII | 19.4 | W | W | W | A | A | A | A | A | A | W | A | W | A | W | W | W | A | W | W | A | A | W | A | A | W | A | W | A | A | A | A | W | W | W | A | A | W | A | A | A | W | W | A | W | A | A | A | W | W | NA | W | A | A | A | A | A | A | A | W | W | A | W | W | W | W | A | A | W | W | A | A | W | A | W | W | A | W | A | W | W | W | A | A | W | A | W | W | W | A | W | A | W |
| ChrII | 19.5 | W | NA | W | NA | A | A | NA | NA | A | W | NA | W | A | W | NA | W | A | W | W | NA | A | W | NA | A | NA | A | W | A | A | A | A | W | W | W | A | A | W | A | A | A | W | W | A | W | A | A | A | W | W | W | W | A | A | A | A | NA | A | A | W | W | A | W | W | W | NA | A | NA | W | W | A | A | W | NA | W | W | A | W | A | W | W | W | A | A | W | A | W | NA | W | A | W | A | W |
| ChrII | 19.6 | W | W | W | A | A | A | A | A | A | W | A | W | A | W | NA | W | A | W | W | A | A | W | A | A | W | A | W | A | A | A | A | W | W | W | A | A | W | A | A | A | W | W | A | W | A | A | A | W | W | W | W | A | A | A | A | A | A | A | W | W | A | W | W | W | NA | A | A | W | W | A | A | W | A | W | W | A | W | A | W | W | W | A | A | W | A | W | W | W | A | W | A | W |
| ChrII | 19.7 | W | W | W | A | A | NA | A | A | A | W | A | W | A | W | W | W | A | W | W | A | A | W | A | NA | W | A | W | NA | A | A | A | W | W | W | A | A | W | NA | A | A | W | W | A | W | NA | A | NA | W | W | W | W | NA | A | A | A | NA | A | NA | W | W | A | W | W | W | W | A | NA | W | W | A | A | W | NA | W | W | A | W | A | W | W | W | A | A | W | A | W | W | W | A | W | A | W |
| ChrII | 20.2 | NA | W | W | A | NA | A | A | NA | NA | A | A | W | A | W | NA | NA | A | W | W | A | A | W | A | W | NA | A | W | W | A | A | A | W | W | W | A | A | W | A | A | A | W | A | A | W | A | A | A | W | W | W | W | A | A | A | A | W | NA | A | W | W | A | W | W | W | W | A | A | W | W | NA | A | W | A | W | W | A | W | A | A | W | W | A | NA | W | A | W | NA | W | NA | W | A | W |
| ChrII | 20.3 | NA | W | W | A | NA | A | A | A | A | A | A | W | A | W | W | W | A | W | W | A | A | W | A | W | W | A | W | W | A | A | A | W | W | W | A | A | W | A | A | A | W | A | A | W | A | A | A | W | W | NA | W | A | A | A | A | W | A | A | W | W | A | W | W | W | W | A | A | W | W | A | A | W | A | W | W | A | W | A | A | W | W | A | A | W | A | W | W | W | A | W | A | W |
| ChrII | 20.4 | W | W | W | A | A | A | A | A | A | A | A | W | A | W | NA | NA | A | W | W | A | A | W | A | W | NA | A | W | W | A | A | A | W | W | W | A | A | W | A | A | A | W | A | A | W | A | A | A | W | W | W | W | A | A | A | A | W | A | A | W | W | A | W | W | W | NA | A | A | W | W | A | A | W | A | W | W | A | W | A | A | W | W | A | A | W | A | W | W | W | A | W | A | W |
| ChrII | 20.5 | W | W | W | A | A | A | A | A | A | A | A | W | A | W | W | W | A | W | W | A | A | W | A | W | W | A | W | W | A | A | A | W | W | W | A | A | W | A | A | A | W | A | A | W | A | A | A | W | W | W | W | A | A | A | A | W | A | A | W | W | A | W | W | W | W | A | A | W | W | A | A | W | A | W | W | A | W | A | A | W | W | A | A | W | A | W | W | W | A | W | A | W |
| ChrII | 20.6 | W | NA | W | A | A | A | A | A | A | A | A | W | A | W | NA | W | A | W | NA | A | NA | W | A | W | W | A | W | W | A | A | A | NA | W | W | A | A | W | A | A | A | W | A | A | W | A | A | A | W | W | NA | W | A | A | A | A | W | A | A | W | NA | A | W | W | W | NA | A | A | NA | W | A | A | W | A | W | W | A | W | A | A | W | W | A | A | W | A | W | NA | W | A | W | A | NA |
| ChrII | 20.7 | NA | A | W | A | A | A | A | A | A | A | A | W | A | NA | W | NA | A | W | A | A | A | W | A | W | W | A | W | W | A | A | A | NA | W | W | A | A | W | A | A | A | W | A | A | W | A | A | A | W | W | W | W | A | A | A | A | W | A | A | W | NA | A | W | W | W | NA | A | A | W | W | A | A | W | A | W | W | A | W | A | A | W | W | A | A | W | A | W | W | W | A | NA | A | NA |
| ChrII | 20.8 | W | A | NA | A | A | A | A | A | A | A | A | W | A | NA | W | NA | NA | W | A | A | A | W | A | W | NA | A | W | W | A | A | A | NA | W | W | A | A | NA | A | A | A | W | A | A | W | A | A | A | W | W | NA | W | A | A | A | A | W | A | A | W | W | A | W | W | W | W | A | A | NA | W | A | A | W | A | W | W | A | W | A | A | W | W | A | A | W | A | W | W | W | A | W | A | W |
| ChrII | 21 | NA | A | W | A | A | A | A | A | NA | A | A | W | A | NA | W | W | A | W | NA | A | A | W | A | W | W | A | W | W | A | A | A | W | W | W | A | A | W | A | A | A | W | NA | A | W | A | A | A | W | W | NA | W | A | A | A | A | W | NA | A | W | W | A | W | W | W | W | A | NA | W | W | A | NA | W | A | W | W | A | W | A | NA | W | W | A | NA | W | A | W | W | W | NA | NA | A | W |
| ChrII | 21.2 | W | A | W | A | A | A | A | A | A | A | A | W | A | NA | W | W | A | W | A | A | A | W | A | W | NA | A | W | W | A | A | A | NA | W | W | A | A | NA | A | A | A | W | A | A | W | A | A | A | W | W | NA | W | A | A | A | A | W | A | A | W | NA | A | W | W | W | NA | A | A | NA | W | A | A | W | A | W | W | A | W | A | A | W | W | A | A | W | W | W | NA | W | A | W | NA | W |
| ChrII | 21.3 | W | A | W | A | A | A | A | A | A | A | A | W | A | W | W | W | A | W | A | A | A | W | A | W | NA | A | W | W | A | A | A | NA | W | W | A | A | W | A | A | A | W | A | A | W | A | A | A | W | W | W | W | A | A | A | NA | W | A | A | W | NA | A | W | W | W | NA | A | A | W | W | A | A | W | A | W | W | A | W | A | A | W | W | A | A | W | W | W | W | W | A | NA | NA | W |
| ChrII | 21.4 | W | A | W | A | A | A | A | A | A | A | A | W | A | W | W | W | A | W | A | A | A | W | A | W | W | A | W | W | A | A | A | W | NA | W | A | A | W | A | A | A | W | A | A | W | A | A | A | W | W | W | W | A | A | A | A | W | A | A | W | W | A | W | W | W | W | A | A | W | W | A | A | W | A | W | W | A | W | A | A | W | W | A | A | W | W | W | W | W | A | W | W | W |
| ChrII | 21.5 | W | A | W | A | A | A | A | A | A | A | A | W | A | W | W | W | A | W | A | A | A | W | A | W | W | A | NA | NA | A | A | A | W | W | W | A | A | W | A | A | A | W | A | A | W | A | A | A | NA | W | W | W | A | A | A | A | W | A | A | W | W | A | W | W | W | W | A | A | NA | W | A | A | W | W | W | W | A | W | A | A | W | W | A | A | NA | W | W | W | W | A | W | W | W |
| ChrII | 21.6 | W | A | W | A | A | A | A | A | A | A | A | W | A | W | W | W | A | W | A | A | A | W | A | W | W | A | W | W | A | A | A | W | W | W | A | A | W | A | A | A | W | A | A | W | A | A | A | W | W | NA | W | A | A | A | A | W | A | A | W | W | A | W | W | W | W | A | A | W | W | A | A | W | W | W | W | A | W | A | A | W | W | A | A | W | W | W | W | W | A | W | W | NA |
| ChrII | 21.7 | NA | A | NA | A | A | A | A | A | A | A | A | W | A | W | NA | NA | A | W | A | A | A | W | A | W | W | A | W | W | A | A | A | W | W | W | A | A | NA | A | A | A | W | A | A | W | A | A | A | W | W | NA | W | A | A | A | A | W | A | A | W | NA | A | W | W | W | W | NA | A | W | W | A | A | W | NA | W | W | A | W | A | A | W | W | A | A | NA | W | W | NA | W | A | W | NA | NA |
| ChrII | 21.9 | W | A | W | A | A | A | A | NA | A | A | A | W | A | W | W | W | NA | W | A | A | A | W | A | W | W | A | W | W | A | A | A | NA | W | W | A | A | W | A | A | A | W | A | A | W | A | A | A | W | W | W | NA | A | A | A | A | W | A | A | W | NA | A | W | W | W | W | A | A | W | W | A | A | W | W | W | W | A | W | A | A | W | W | A | A | W | W | W | W | W | A | W | W | W |
| ChrII | 22 | W | A | NA | A | A | A | A | A | A | A | A | W | A | NA | NA | W | A | W | A | A | A | W | A | W | W | A | W | W | A | A | A | NA | W | W | A | A | W | A | A | A | W | A | A | W | A | A | A | W | W | W | W | A | A | A | A | W | A | A | W | W | A | W | W | W | NA | A | A | W | W | A | A | W | NA | W | W | A | W | A | A | W | W | A | A | W | W | W | W | W | A | W | W | W |
| ChrII | 22.1 | W | A | W | A | A | A | A | A | A | A | A | W | A | W | W | W | A | W | A | A | A | W | A | W | W | A | W | W | A | A | A | W | W | W | A | A | W | A | A | A | W | A | A | W | A | A | A | W | W | W | W | A | A | A | A | W | A | A | W | W | A | W | W | W | W | A | A | W | W | A | A | W | W | W | W | A | W | A | A | W | W | A | A | W | W | W | W | W | A | W | W | W |
| ChrII | 22.2 | W | A | W | A | A | A | A | A | A | A | A | W | A | W | W | W | A | W | A | NA | A | W | A | W | W | A | W | W | A | A | A | W | W | W | A | A | W | A | A | A | W | A | A | W | A | A | A | W | W | NA | W | A | A | A | A | W | A | A | W | W | A | W | W | W | W | A | A | W | W | A | A | W | W | W | NA | A | W | A | A | W | W | A | A | W | W | W | W | W | A | W | W | W |
| ChrII | 22.3 | W | A | W | A | A | A | A | A | A | A | A | W | A | W | W | W | A | W | A | A | A | W | A | W | W | A | W | W | A | A | A | W | W | W | A | A | W | A | A | A | W | A | A | W | A | A | A | W | W | W | W | A | A | A | A | W | A | A | W | W | A | W | W | W | W | A | A | W | W | A | A | W | W | W | W | A | W | A | A | W | W | A | A | W | W | W | W | W | A | W | W | W |
| ChrII | 22.4 | W | A | W | A | A | A | A | A | A | A | A | W | A | W | W | W | A | W | A | A | A | W | A | W | W | A | W | W | A | A | A | W | W | W | A | A | W | A | A | A | W | A | A | W | A | A | A | W | W | W | W | A | A | A | A | W | A | A | W | W | A | W | W | W | W | A | A | W | W | A | A | W | W | W | W | A | W | A | A | W | W | A | A | W | W | W | W | W | A | W | W | W |
| ChrII | 22.5 | W | A | W | A | A | A | A | A | A | A | A | W | A | W | W | W | A | W | A | A | A | W | A | W | W | A | W | W | A | A | A | W | W | W | A | A | W | A | A | A | W | A | A | W | A | A | A | W | W | W | W | A | A | A | A | W | A | A | W | W | A | W | W | W | W | A | A | W | W | A | A | W | W | W | W | A | W | A | A | W | W | A | A | W | W | W | W | W | A | W | W | W |
| ChrII | 22.6 | W | A | W | A | A | A | A | A | A | A | A | W | A | NA | W | W | A | W | A | A | A | W | A | W | W | A | W | W | A | A | A | W | W | W | A | A | W | A | A | A | W | A | A | W | A | A | A | W | W | NA | W | A | A | A | A | W | A | A | W | W | A | W | W | W | NA | A | A | W | W | A | A | W | W | W | W | A | W | A | A | W | W | A | A | W | W | W | NA | W | A | W | W | W |
| ChrII | 22.7 | W | A | W | A | A | A | A | A | A | A | A | W | A | W | W | W | A | W | A | A | A | W | A | W | W | A | W | W | A | A | A | W | W | NA | A | A | NA | A | A | A | W | A | A | W | A | A | A | W | W | NA | W | A | A | A | A | W | A | A | W | W | A | W | W | W | W | NA | A | W | W | A | A | W | W | W | W | A | W | A | A | W | W | A | A | W | W | W | W | W | A | W | W | W |
| ChrII | 22.8 | W | A | W | A | A | A | A | A | A | A | A | W | A | W | W | W | A | W | A | A | A | W | A | W | W | A | W | W | A | A | A | W | W | W | A | A | W | A | A | A | W | A | A | W | A | A | A | W | W | NA | W | A | A | A | A | W | A | A | W | W | A | W | W | W | W | A | A | W | W | A | A | W | W | W | W | A | W | A | A | W | W | A | A | W | W | W | W | W | A | W | W | W |
| ChrII | 22.9 | W | A | W | A | A | A | A | A | A | A | A | W | A | NA | W | W | A | W | A | A | A | W | A | W | W | A | W | W | A | A | A | W | W | W | A | A | W | A | A | A | W | A | A | W | A | A | A | W | W | W | W | A | A | A | A | W | A | A | W | W | A | W | W | W | W | A | A | W | W | A | A | W | W | W | W | A | W | A | A | W | W | A | A | W | W | W | W | W | A | W | W | W |
| ChrII | 23 | W | A | W | A | A | A | A | A | A | A | A | W | A | W | W | W | A | W | A | A | A | W | A | W | W | A | W | W | A | A | A | W | W | W | A | A | W | A | A | A | W | A | A | W | A | A | A | W | W | W | W | A | A | A | A | W | A | A | W | W | A | W | W | W | W | A | A | W | W | A | A | W | W | W | W | A | W | A | A | W | W | A | A | W | W | W | W | W | A | W | W | W |
| ChrII | 23.1 | W | A | W | A | A | A | A | A | A | A | A | W | A | W | W | NA | A | W | A | NA | A | W | A | W | W | A | W | W | A | A | A | W | W | W | A | A | W | A | A | A | W | A | A | W | A | A | A | W | W | NA | W | A | A | A | A | W | A | A | W | NA | A | W | W | W | NA | A | A | W | W | A | A | W | W | W | W | A | W | A | A | W | W | A | A | W | W | W | NA | W | A | W | W | W |
| ChrII | 23.2 | NA | A | NA | A | A | A | A | A | A | A | A | W | A | NA | NA | NA | A | W | A | NA | A | W | A | W | NA | A | W | W | A | A | A | NA | W | W | A | NA | NA | A | A | A | W | A | A | W | A | A | A | W | W | NA | NA | A | A | A | A | W | A | A | W | W | A | W | W | W | NA | A | A | W | W | A | A | W | NA | W | NA | A | W | A | A | W | W | A | A | NA | W | W | NA | W | A | W | W | NA |
| ChrIII | 0 | A | W | W | A | W | W | W | A | A | A | W | A | W | A | W | A | W | A | W | W | A | W | A | W | W | W | W | A | W | W | W | W | W | W | W | W | A | W | A | A | W | A | A | W | A | W | W | A | A | A | W | A | W | W | NA | A | A | A | W | A | W | A | W | A | A | A | W | A | W | W | NA | W | A | A | A | W | A | A | W | W | A | W | W | W | A | A | A | W | W | A | W | W |
| ChrIII | 0.1 | A | W | W | A | W | W | W | A | A | A | W | A | W | A | W | A | W | A | W | W | A | W | A | W | W | W | W | A | W | W | W | W | W | NA | W | W | A | W | A | A | W | A | A | W | A | W | W | A | A | A | W | A | W | W | NA | A | A | A | W | A | W | A | W | A | A | A | W | A | W | W | W | W | A | A | A | W | A | A | W | W | A | W | W | W | A | A | A | W | W | A | W | W |
| ChrIII | 0.2 | A | W | W | A | W | W | NA | A | A | A | W | A | W | A | NA | A | W | A | W | W | A | W | A | W | W | W | W | NA | W | W | W | NA | W | W | W | W | A | W | A | A | W | A | A | W | A | W | W | A | A | A | W | A | W | W | A | A | A | A | W | A | W | A | W | A | A | A | NA | A | W | W | NA | W | A | A | A | W | A | A | W | W | A | W | W | W | A | A | A | W | W | A | W | W |
| ChrIII | 0.3 | A | W | W | A | W | W | W | A | A | A | W | A | W | A | NA | A | W | A | W | W | A | W | A | W | W | W | W | A | W | W | W | W | W | W | W | W | A | W | A | A | W | A | A | W | A | W | W | A | A | A | W | A | W | W | A | A | A | A | W | A | W | A | W | A | A | A | W | A | W | W | NA | W | A | A | A | W | A | A | W | W | A | W | W | W | A | A | A | W | W | A | W | W |
| ChrIII | 0.4 | A | W | W | A | W | W | W | A | A | A | W | A | W | A | W | A | W | A | W | W | A | W | A | W | W | W | W | A | W | W | W | W | W | W | W | W | A | W | A | A | W | A | A | W | A | W | W | A | A | A | W | A | W | W | A | A | A | A | W | A | W | A | W | A | A | A | W | A | W | W | W | W | A | A | A | W | A | A | W | W | A | W | W | W | A | A | A | W | W | A | W | W |
| ChrIII | 0.5 | A | W | W | A | W | W | NA | A | A | A | W | A | W | A | W | A | W | A | W | W | A | NA | A | W | W | W | W | A | W | W | W | W | NA | W | W | W | A | W | A | A | W | A | A | W | A | W | W | A | A | A | W | A | W | W | A | A | A | A | W | A | W | A | W | A | A | NA | A | W | W | W | W | A | A | A | W | A | A | W | W | A | W | W | W | A | A | A | W | W | A | W | W |
| ChrIII | 0.7 | A | W | NA | A | W | W | NA | NA | A | A | W | A | NA | A | NA | A | W | A | NA | W | A | W | A | W | W | NA | W | NA | W | NA | W | NA | NA | W | W | W | A | NA | A | A | W | A | A | W | A | NA | NA | A | A | A | W | A | W | NA | A | A | A | A | W | A | W | A | NA | A | A | A | W | A | W | NA | NA | W | A | A | A | W | A | A | W | W | A | W | W | W | A | A | A | W | W | A | W | W |
| ChrIII | 0.8 | A | W | W | A | W | W | W | A | A | A | W | A | W | A | W | A | W | A | W | W | A | W | A | W | W | W | W | A | W | W | W | W | W | W | W | W | A | W | A | A | W | A | A | W | A | W | W | A | A | A | W | A | W | W | A | A | A | A | W | A | W | A | NA | A | A | A | W | A | W | W | W | W | A | A | A | W | A | A | W | W | A | W | W | W | A | A | A | W | NA | A | W | W |
| ChrIII | 0.9 | A | W | W | A | W | W | W | A | A | A | W | A | W | NA | NA | A | W | A | W | W | A | W | A | W | W | W | W | A | NA | W | W | W | W | W | W | W | A | W | A | A | W | A | A | W | A | W | W | A | A | NA | W | NA | W | W | A | A | A | A | W | A | W | A | W | A | A | A | W | A | W | W | W | W | A | A | A | W | A | A | W | W | NA | NA | W | W | A | A | A | W | W | A | W | W |
| ChrIII | 1.3 | NA | W | W | A | W | W | W | A | A | A | W | NA | W | A | NA | A | W | A | W | W | A | W | A | W | W | W | W | A | W | W | W | W | W | W | W | W | A | W | A | NA | W | A | A | W | A | W | W | A | NA | A | W | A | W | W | A | A | NA | NA | W | A | W | NA | W | A | NA | NA | W | A | W | W | W | W | A | NA | A | W | NA | A | W | W | NA | W | W | W | A | A | A | W | W | A | W | W |
| ChrIII | 1.4 | A | W | W | A | W | W | W | A | A | A | W | A | W | A | NA | A | W | A | NA | W | A | W | A | W | W | W | W | A | W | W | W | W | W | W | W | W | A | W | A | A | W | A | A | W | A | W | W | A | A | A | W | A | W | W | A | A | A | A | W | A | W | A | W | A | A | A | W | A | W | W | NA | W | A | A | A | W | A | A | W | W | A | W | W | W | A | A | A | W | W | A | W | W |
| ChrIII | 1.5 | A | W | W | A | W | W | W | A | A | A | W | NA | W | A | NA | A | W | A | W | W | NA | NA | A | W | W | W | W | A | W | W | W | W | W | W | W | W | A | W | A | A | W | A | A | W | A | W | W | A | A | A | W | A | W | W | A | A | A | A | W | A | W | A | W | A | A | A | W | A | W | W | W | W | A | A | A | W | A | A | W | W | A | W | W | W | A | A | A | W | W | A | NA | W |
| ChrIII | 1.6 | A | W | W | A | W | W | NA | NA | A | A | NA | A | W | A | NA | A | W | A | NA | W | NA | W | A | NA | W | W | W | A | W | NA | W | NA | NA | W | W | W | A | W | A | A | W | A | A | W | A | W | NA | A | A | A | W | A | W | W | NA | A | A | A | W | A | W | A | NA | A | A | NA | W | A | W | W | W | W | A | A | A | NA | A | A | W | NA | A | NA | W | NA | NA | A | NA | NA | W | A | NA | NA |
| ChrIII | 1.7 | A | W | NA | A | W | W | NA | A | A | A | W | A | W | A | NA | A | W | A | NA | W | A | NA | A | W | W | NA | NA | A | W | NA | W | NA | NA | NA | W | W | A | NA | A | A | W | A | A | W | A | NA | W | A | A | A | W | A | W | NA | A | A | A | A | W | A | W | A | NA | A | A | A | NA | A | W | NA | NA | W | A | A | A | W | A | A | W | NA | A | NA | W | NA | A | A | A | NA | NA | A | W | NA |
| ChrIII | 1.8 | A | W | W | A | W | W | NA | A | A | A | W | A | W | A | NA | A | W | A | W | W | NA | W | A | W | W | W | W | A | W | NA | W | W | W | W | W | W | A | W | A | A | W | A | A | W | A | W | W | A | A | A | W | A | W | W | A | A | A | A | W | A | W | A | W | A | A | A | W | A | W | W | W | W | A | A | A | W | A | A | W | W | A | W | W | W | A | A | A | W | W | A | W | W |
| ChrIII | 1.9 | NA | W | W | A | W | W | W | A | A | A | W | A | W | A | W | A | W | A | W | NA | A | W | A | W | W | W | W | A | W | W | W | NA | W | W | W | W | A | W | A | A | W | A | A | W | A | W | W | A | A | A | W | A | W | W | A | A | A | A | W | A | W | A | W | A | A | A | NA | A | W | W | W | W | A | A | A | W | A | A | W | W | A | W | W | W | A | A | A | W | NA | A | W | W |
| ChrIII | 2 | A | W | W | A | W | W | W | A | A | A | W | A | W | A | W | A | W | A | W | W | A | W | A | W | W | W | W | A | W | W | W | W | W | W | W | W | A | W | A | A | W | A | A | W | A | W | W | A | A | A | W | A | W | W | A | A | A | A | W | A | W | A | W | A | A | A | W | A | W | W | W | W | A | A | A | W | A | A | W | W | A | W | W | W | A | A | A | W | W | A | W | W |

| | | | | | | | | | | | | | | | | | | | | | | | | | | | | | | | | | | | | | | | | | | | | | | | | | | | | | | | | | | | | | | | | | | | | | | | | | | | | | | | | | | | | | | | | | | | | | |
|---|---|---|---|---|---|---|---|---|---|---|---|---|---|---|---|---|---|---|---|---|---|---|---|---|---|---|---|---|---|---|---|---|---|---|---|---|---|---|---|---|---|---|---|---|---|---|---|---|---|---|---|---|---|---|---|---|---|---|---|---|---|---|---|---|---|---|---|---|---|---|---|---|---|---|---|---|---|---|---|---|---|---|---|---|---|---|---|---|---|---|---|---|---|
| ChrIII | 2.1 | A | W | W | A | W | W | W | A | A | A | W | A | W | A | W | A | W | A | W | W | A | W | A | W | W | W | W | A | W | W | W | W | W | W | W | W | A | W | A | A | W | A | A | W | A | W | W | A | A | A | W | A | W | W | NA | A | A | A | W | A | W | A | W | A | A | A | W | A | W | W | W | W | A | A | A | W | A | A | W | W | A | W | W | W | A | A | A | W | W | A | W | W |
| ChrIII | 2.2 | A | W | W | A | W | W | W | NA | A | A | W | A | W | A | W | A | W | A | W | W | A | W | NA | W | W | W | W | A | W | W | W | W | W | W | W | W | A | W | A | A | W | A | A | W | A | W | W | A | A | A | W | A | W | W | NA | A | A | A | W | A | W | A | W | A | A | A | W | A | W | W | W | W | A | A | A | W | A | A | W | W | A | W | W | W | A | A | A | W | W | A | W | W |
| ChrIII | 2.3 | A | W | W | A | W | W | W | A | A | A | W | A | W | A | W | A | W | A | W | W | A | W | A | W | W | W | W | A | W | W | W | W | W | W | W | W | A | W | A | A | W | A | A | W | NA | W | W | A | A | NA | W | A | W | W | A | A | A | A | W | A | W | A | W | A | NA | A | W | A | W | W | W | W | A | NA | A | W | A | A | W | W | A | W | W | W | A | A | A | W | W | A | W | W |
| ChrIII | 2.5 | A | W | W | A | W | W | W | NA | A | A | W | A | W | A | NA | A | W | A | W | W | A | W | A | W | W | W | W | A | W | W | W | W | W | W | W | W | A | W | A | A | W | A | A | W | A | W | W | A | A | A | W | A | W | W | NA | A | A | A | W | A | W | A | W | A | A | A | W | A | W | W | W | W | A | A | A | W | A | A | W | W | A | W | W | W | A | A | A | W | W | A | W | W |
| ChrIII | 2.6 | A | W | W | A | W | W | W | A | A | A | W | A | W | A | W | A | W | A | W | W | A | W | A | W | W | W | W | A | W | W | W | W | W | W | W | W | A | W | A | A | W | A | A | W | A | W | W | A | A | A | W | A | W | W | A | A | A | A | W | A | W | A | NA | A | A | A | W | A | W | W | NA | W | A | A | A | W | A | A | W | W | A | W | W | W | A | A | A | W | W | A | W | W |
| ChrIII | 2.7 | A | W | NA | A | W | W | W | A | A | A | W | A | NA | A | NA | A | W | A | W | W | A | W | NA | NA | W | NA | NA | NA | W | W | W | W | NA | W | W | W | A | W | A | A | W | A | A | W | A | W | W | A | A | A | W | A | W | W | A | A | A | A | W | A | W | A | NA | NA | A | A | W | A | W | W | W | W | A | A | A | NA | A | A | NA | W | A | NA | W | W | NA | A | NA | NA | W | A | NA | W |
| ChrIII | 2.8 | A | W | W | A | W | W | W | A | A | A | W | A | W | A | W | A | W | A | W | W | A | W | A | W | W | NA | W | A | W | W | W | W | W | NA | W | W | A | W | A | A | W | A | A | W | A | W | W | A | A | A | W | A | W | W | A | A | A | A | W | A | W | A | NA | A | A | A | W | A | W | W | W | W | A | A | A | W | A | A | W | W | A | W | W | W | A | A | A | W | W | A | NA | W |
| ChrIII | 2.9 | A | W | W | A | W | W | W | A | A | A | W | A | W | A | NA | A | W | A | W | W | A | W | A | W | W | W | W | A | W | W | W | W | W | W | W | W | A | W | A | A | W | A | A | W | A | W | W | A | A | A | W | A | W | W | A | A | A | A | W | A | W | A | W | A | A | A | W | A | W | W | W | W | A | A | A | W | A | A | W | W | A | W | W | W | A | A | A | W | W | A | NA | W |
| ChrIII | 3.1 | A | W | W | A | W | W | W | A | A | A | W | A | W | A | W | A | W | A | W | W | A | W | A | W | W | W | W | A | W | W | W | W | W | W | W | W | A | W | A | A | W | A | A | W | A | W | W | A | A | A | W | A | W | W | A | A | A | A | W | A | W | A | W | A | A | A | W | A | W | W | W | W | A | A | A | W | A | A | W | W | A | W | W | W | A | A | A | W | W | A | W | W |
| ChrIII | 3.2 | A | W | W | A | W | W | W | A | A | A | W | A | W | A | NA | A | W | A | W | W | A | W | A | W | W | W | W | A | W | W | W | W | W | W | W | W | A | W | A | A | W | A | A | W | A | W | W | A | A | A | W | A | W | W | NA | A | A | A | W | A | W | A | W | A | A | A | NA | A | W | W | W | W | A | A | A | W | A | A | W | W | A | W | W | W | A | A | A | W | W | A | W | NA |
| ChrIII | 3.3 | A | W | W | A | W | W | W | A | A | A | W | A | W | A | W | A | W | A | W | W | A | W | A | W | W | W | W | A | W | W | W | W | W | W | W | W | A | W | A | A | W | A | A | W | A | W | W | A | A | A | W | A | W | W | A | A | A | A | W | A | W | A | W | A | A | A | W | A | W | W | W | W | A | A | A | W | A | A | W | W | A | W | W | W | A | A | A | W | W | A | W | W |
| ChrIII | 3.4 | A | W | W | A | W | W | W | A | A | A | W | A | W | A | W | A | W | A | W | W | A | W | A | W | W | W | W | A | W | W | W | W | W | W | W | W | A | W | A | A | W | A | A | W | A | W | W | A | A | A | W | A | W | W | A | A | A | A | W | A | W | A | W | A | A | A | W | A | W | W | W | W | A | A | A | W | A | A | W | W | A | W | W | W | A | A | A | W | W | A | W | W |
| ChrIII | 3.5 | A | W | W | A | W | NA | W | A | A | A | W | A | W | A | NA | A | W | A | W | W | NA | W | A | W | W | W | W | A | W | W | W | W | W | W | W | W | A | W | A | A | W | A | A | W | A | W | W | A | A | A | W | A | W | W | A | A | A | A | W | A | W | A | W | A | A | A | NA | A | W | W | W | W | A | A | A | W | A | A | W | W | A | W | W | W | A | A | A | W | W | A | W | W |
| ChrIII | 3.6 | A | W | W | A | W | W | W | A | A | A | W | A | W | A | W | A | W | A | W | W | A | W | A | W | W | W | W | A | W | W | W | W | W | W | W | W | A | W | A | A | W | A | A | W | A | W | W | A | A | A | W | A | W | W | A | A | A | A | W | A | W | A | W | A | A | A | W | A | W | W | W | W | A | A | A | W | A | A | W | W | A | W | W | W | A | A | A | W | W | A | W | W |
| ChrIII | 3.7 | A | W | W | A | W | W | NA | NA | A | A | W | A | W | NA | W | A | W | A | W | W | A | W | A | W | W | W | W | NA | W | W | W | W | W | W | W | W | A | W | A | A | W | A | A | W | A | W | W | A | A | A | W | A | W | W | A | A | A | A | W | A | W | A | W | A | A | A | W | A | W | W | W | W | A | A | A | W | A | A | W | W | A | W | W | W | A | A | A | W | W | A | W | W |
| ChrIII | 3.8 | A | W | W | A | W | W | W | A | A | A | W | A | W | A | W | A | W | A | W | W | A | W | A | W | W | W | W | A | W | W | W | W | W | W | W | W | A | W | A | A | W | A | A | W | A | W | W | A | A | A | W | A | W | W | A | A | A | A | W | A | W | A | W | A | A | A | W | A | W | W | W | W | A | A | A | W | A | A | W | W | A | W | W | W | A | A | A | W | W | A | W | W |
| ChrIII | 3.9 | A | W | W | A | W | W | W | A | A | A | W | A | W | A | W | A | W | A | W | NA | A | W | A | W | W | W | W | A | W | W | W | W | W | W | W | W | A | W | A | A | W | A | A | W | NA | W | W | A | A | A | W | A | W | W | A | A | A | A | W | A | W | A | W | A | NA | NA | W | A | W | W | W | NA | A | A | A | W | A | A | W | W | A | W | W | W | A | A | A | W | W | A | A | W |
| ChrIII | 4 | A | W | W | A | W | W | W | NA | A | A | W | A | W | A | W | A | W | A | W | W | A | W | A | W | W | NA | W | A | W | W | W | W | W | W | W | W | A | W | A | A | W | A | A | W | A | W | W | A | A | A | W | A | W | W | A | A | A | A | W | A | W | A | W | A | A | A | W | A | W | W | W | W | A | A | A | W | A | A | W | W | A | W | W | W | A | A | A | W | W | A | A | W |
| ChrIII | 4.1 | A | W | NA | A | W | W | W | NA | A | A | W | A | W | A | W | A | W | A | NA | W | NA | W | NA | W | W | W | W | A | W | W | W | W | W | W | W | W | A | W | A | A | W | A | A | W | A | W | W | A | A | NA | W | A | W | W | A | A | A | A | W | A | W | A | W | A | A | A | W | A | W | W | W | W | A | A | A | W | A | A | W | W | A | W | W | W | A | A | A | W | W | A | A | W |
| ChrIII | 4.2 | A | W | NA | A | W | W | W | A | A | A | NA | A | W | A | W | A | W | A | W | W | A | NA | A | NA | W | W | W | A | W | W | W | W | NA | W | W | W | A | W | A | A | W | A | A | W | A | W | NA | A | A | A | W | A | W | W | A | A | A | A | W | A | W | A | W | A | A | A | W | A | W | NA | W | W | A | A | A | NA | A | A | W | W | A | NA | W | W | A | A | A | NA | NA | A | A | NA |
| ChrIII | 4.3 | A | W | W | A | W | W | NA | A | A | A | W | A | W | A | W | A | W | A | W | W | A | W | A | W | W | W | W | A | W | W | W | W | NA | W | W | W | A | W | A | A | W | A | A | W | A | W | NA | A | A | A | W | A | W | W | A | A | A | A | W | A | W | A | W | A | A | A | W | A | W | W | W | W | A | A | A | W | A | A | W | W | A | W | W | W | A | A | A | NA | W | A | A | W |
| ChrIII | 4.4 | A | W | W | A | W | W | W | A | A | A | W | A | W | A | NA | A | W | A | W | W | A | W | A | W | W | W | W | A | W | W | W | W | W | W | W | W | A | W | A | A | W | A | A | W | A | W | W | A | A | A | W | A | W | W | A | A | A | A | W | A | W | A | W | A | A | A | W | A | W | W | W | NA | A | A | A | W | A | A | W | W | A | W | W | W | A | A | A | W | W | A | A | W |
| ChrIII | 4.5 | A | W | W | A | W | W | W | A | A | A | W | A | W | A | W | A | W | A | W | W | A | W | A | W | W | W | W | A | W | W | W | W | W | W | W | W | A | W | A | A | W | A | A | W | A | W | W | A | A | A | W | A | W | W | A | A | A | A | W | A | W | A | W | A | A | A | W | A | W | W | W | W | A | A | A | W | A | A | W | W | A | W | W | W | A | A | A | W | W | A | A | W |
| ChrIII | 4.6 | A | W | W | A | W | W | W | A | A | A | W | A | W | A | W | A | W | A | W | W | A | W | A | W | W | W | NA | A | W | W | W | W | W | W | W | W | A | W | A | A | W | A | A | W | A | W | W | A | A | NA | W | A | W | W | NA | A | A | A | W | A | W | A | W | A | A | A | W | A | W | W | W | W | A | A | A | W | A | A | W | W | A | W | W | W | A | A | NA | W | W | A | A | W |
| ChrIII | 4.7 | NA | W | W | A | W | W | W | NA | A | A | W | A | W | NA | W | NA | W | A | W | W | NA | W | NA | W | W | W | W | A | W | W | W | W | W | W | W | W | A | W | A | NA | W | A | A | W | NA | W | W | A | NA | A | W | A | W | W | NA | A | A | A | W | A | W | NA | W | NA | NA | NA | W | A | W | W | W | W | NA | A | A | W | A | A | W | W | NA | W | W | W | A | A | A | W | W | A | A | W |
| ChrIII | 4.8 | A | W | W | A | W | W | W | A | A | A | W | A | W | A | W | A | W | A | W | W | A | W | A | W | W | W | W | A | W | W | W | W | W | W | W | W | A | W | A | A | W | A | A | W | A | W | W | A | A | A | W | A | W | W | A | A | A | A | W | A | W | A | W | A | A | A | W | A | W | W | W | W | A | A | A | W | A | A | W | W | A | W | W | W | A | A | A | W | W | A | A | W |
| ChrIII | 4.9 | A | W | W | A | W | W | W | A | A | A | W | A | W | A | W | A | W | NA | W | W | NA | W | A | W | W | NA | W | A | W | W | W | W | W | NA | W | W | A | W | A | A | W | A | A | W | A | W | W | A | A | A | W | A | W | W | A | A | A | A | W | A | W | A | W | A | A | A | W | A | W | W | W | W | A | A | A | W | A | A | W | W | A | W | W | W | A | A | A | W | W | A | A | W |
| ChrIII | 5 | A | W | W | A | W | W | W | A | A | A | W | A | W | A | W | A | W | A | W | W | A | W | A | W | W | W | W | A | W | W | W | W | W | W | W | W | A | W | A | A | W | A | A | W | A | W | W | A | A | A | W | A | W | W | A | A | A | A | W | A | W | A | W | A | A | A | W | A | W | W | W | W | A | A | A | W | A | A | W | W | A | W | W | W | A | A | A | W | W | A | A | W |
| ChrIII | 5.1 | A | W | W | A | W | NA | W | A | A | A | W | A | W | A | W | A | W | A | W | W | A | W | A | W | W | W | W | A | W | W | W | W | W | W | W | W | A | W | A | A | W | A | A | W | A | W | W | A | A | A | W | A | W | W | A | A | A | A | W | A | W | A | W | A | A | A | W | A | W | W | W | W | A | A | A | W | A | A | W | W | A | W | W | W | A | A | A | W | W | A | A | W |
| ChrIII | 5.2 | A | W | W | A | W | NA | W | A | A | A | W | A | W | A | W | A | W | A | W | W | A | W | A | W | W | W | W | A | W | W | W | W | W | NA | W | W | A | W | A | A | W | A | A | W | A | W | W | A | A | A | W | A | W | NA | A | A | A | A | W | A | W | A | W | A | A | A | W | A | W | W | W | W | A | A | A | W | A | A | W | W | A | W | W | W | A | A | A | W | W | A | A | W |
| ChrIII | 5.3 | A | W | W | A | W | W | W | A | A | A | W | A | W | A | W | A | W | A | W | W | A | W | A | W | W | W | W | A | W | W | W | W | W | W | W | W | A | W | A | A | W | A | A | W | A | W | W | A | A | A | W | A | W | W | A | A | A | A | W | A | W | A | NA | A | A | A | W | A | W | W | W | W | A | A | A | W | A | A | W | W | A | W | W | W | A | A | A | W | W | A | A | W |
| ChrIII | 5.4 | A | W | W | A | W | W | W | A | A | A | W | A | W | A | W | A | W | A | W | W | A | W | A | W | W | W | W | A | W | W | W | W | W | W | W | W | A | W | A | A | W | A | A | W | A | W | W | A | A | A | W | A | W | W | A | A | A | A | W | A | W | A | W | A | A | A | W | A | W | W | W | W | A | A | A | W | A | A | W | W | A | W | W | W | A | A | A | W | W | A | A | W |
| ChrIII | 5.5 | NA | W | W | A | W | W | W | NA | NA | A | W | NA | W | A | W | A | W | A | W | W | A | W | A | W | W | W | W | NA | W | W | W | W | W | W | W | W | A | W | A | NA | W | A | A | W | NA | W | W | A | A | NA | W | NA | W | W | NA | A | NA | A | W | A | W | A | W | NA | NA | NA | W | A | W | W | W | W | A | A | A | W | A | A | W | W | NA | W | W | W | A | A | A | W | W | A | A | W |
| ChrIII | 5.6 | A | W | W | A | W | W | W | A | A | A | W | A | W | A | W | A | W | A | W | W | A | W | A | W | W | W | W | A | W | W | W | W | W | W | W | W | A | W | A | A | W | A | A | W | A | W | W | A | A | A | W | A | W | W | A | A | A | A | W | A | W | A | W | A | A | A | W | A | W | W | W | W | A | A | A | W | A | A | W | W | A | W | W | W | A | A | A | W | W | A | A | W |
| ChrIII | 5.7 | A | W | W | NA | W | W | NA | A | A | A | W | A | W | A | W | A | W | A | W | NA | A | W | A | W | W | W | W | A | W | W | W | W | W | W | W | W | A | W | A | A | W | A | A | W | A | W | W | A | A | NA | W | NA | W | W | A | A | A | A | W | A | W | A | W | A | NA | A | NA | A | W | W | W | W | A | A | A | W | A | A | W | W | A | W | W | W | A | A | A | W | W | A | A | W |
| ChrIII | 5.8 | A | W | W | A | W | W | NA | A | A | A | W | A | W | A | W | A | W | A | W | W | A | W | A | W | W | W | W | A | W | W | W | NA | W | W | W | W | A | W | A | A | W | A | A | W | A | W | W | A | A | A | W | A | W | W | A | A | A | A | W | A | W | A | W | A | A | A | W | A | W | W | W | W | A | A | A | W | A | A | W | W | A | W | W | W | A | A | A | NA | W | A | A | W |
| ChrIII | 5.9 | A | W | W | A | W | W | W | A | A | A | W | A | W | A | W | A | W | A | W | W | A | W | A | W | W | W | W | A | W | W | W | W | W | W | W | W | A | W | A | A | W | A | A | W | A | W | W | A | A | A | W | A | W | W | A | A | A | A | W | A | W | A | W | A | A | A | W | A | W | W | W | W | A | A | A | W | A | A | W | W | A | W | W | W | A | A | A | W | W | A | A | W |
| ChrIII | 6 | A | W | W | A | W | W | W | A | A | A | W | A | W | A | W | A | W | A | W | W | A | W | A | W | W | W | W | A | W | W | W | W | W | W | W | W | A | W | A | A | W | A | A | W | A | W | W | A | A | A | W | A | W | W | A | A | A | A | W | A | W | A | W | A | A | A | W | A | W | W | W | W | A | A | A | W | A | A | W | W | A | W | W | W | A | A | A | W | W | A | A | W |
| ChrIII | 6.1 | A | W | W | A | W | W | W | A | A | A | W | A | W | A | W | A | W | A | W | W | A | W | A | W | W | W | W | A | W | W | W | W | W | W | W | W | A | W | A | A | W | A | A | W | A | W | W | A | A | A | W | A | W | W | A | A | A | A | W | A | W | A | W | A | A | A | W | A | W | W | W | W | A | A | A | W | A | A | W | W | A | W | W | W | A | A | A | W | W | A | A | W |
| ChrIII | 6.2 | A | W | W | A | W | W | W | NA | A | A | W | A | W | A | W | A | W | A | W | W | NA | W | NA | W | W | W | W | A | W | W | W | W | W | W | W | W | A | NA | A | A | W | A | A | W | NA | W | W | A | A | A | W | NA | W | W | A | A | A | A | W | A | W | A | W | A | A | NA | W | A | W | W | W | W | A | A | A | W | A | A | W | W | NA | W | W | W | A | A | A | W | W | A | A | W |
| ChrIII | 6.3 | A | W | W | A | W | NA | W | A | A | A | W | A | W | A | W | A | W | A | W | W | A | W | A | W | W | W | W | A | W | W | W | W | NA | W | W | W | A | W | A | A | W | A | A | W | A | W | W | A | A | A | W | A | W | W | A | A | A | A | W | A | W | A | W | A | A | A | W | A | W | W | W | W | A | A | A | NA | A | A | W | W | A | W | W | W | A | A | A | W | W | A | A | NA |
| ChrIII | 6.4 | A | W | W | A | W | NA | NA | A | A | A | NA | A | W | NA | NA | A | W | A | NA | W | A | NA | A | W | W | W | W | A | W | W | W | NA | NA | W | W | W | NA | W | A | A | W | A | A | W | A | W | NA | A | A | A | W | A | W | NA | A | A | A | A | W | NA | W | A | W | A | A | A | W | NA | W | NA | NA | NA | NA | A | A | W | A | A | W | NA | A | W | W | NA | NA | A | A | W | W | A | A | NA |
| ChrIII | 6.5 | NA | W | W | A | W | W | W | NA | NA | A | W | NA | W | A | NA | A | W | A | W | W | NA | W | A | W | W | W | W | A | W | W | W | W | W | W | W | W | A | W | A | NA | W | A | A | W | NA | W | W | A | A | A | W | NA | W | W | A | A | A | A | W | A | W | A | W | A | NA | NA | W | A | W | W | W | W | A | A | A | W | A | A | W | W | A | W | W | W | A | A | A | W | W | A | A | W |
| ChrIII | 6.6 | A | W | W | A | W | W | W | A | A | A | W | A | W | A | W | A | W | A | W | W | A | W | A | W | W | W | W | A | W | W | W | W | W | W | W | W | A | W | A | A | W | A | A | W | A | W | W | A | A | A | W | A | W | W | A | A | A | A | W | A | W | A | W | A | A | A | W | A | W | W | NA | W | A | A | A | W | A | A | W | W | A | W | W | W | A | A | A | W | W | A | A | W |
| ChrIII | 6.7 | A | W | W | A | W | W | W | A | A | A | W | A | W | A | W | A | W | A | W | W | A | W | A | W | W | W | W | A | W | W | W | W | W | W | W | W | A | W | A | A | W | A | A | W | A | W | W | A | A | A | W | A | W | W | A | A | A | A | W | A | W | A | W | A | A | A | W | A | W | W | W | W | A | A | A | W | A | A | W | W | A | W | W | W | A | A | A | W | W | A | A | W |
| ChrIII | 6.8 | A | W | W | A | W | W | NA | A | A | A | W | A | W | A | W | A | W | A | W | W | A | W | A | W | W | W | W | A | W | W | W | W | W | W | W | W | A | W | A | A | W | A | A | W | A | W | W | A | A | A | W | A | W | W | NA | A | A | A | W | A | W | A | W | A | A | A | W | A | W | W | W | W | A | A | A | W | A | A | W | W | A | W | W | W | A | A | A | W | W | A | A | W |
| ChrIII | 6.9 | A | W | W | A | W | W | NA | NA | A | A | W | A | W | A | W | A | W | A | W | W | A | W | A | W | W | W | W | A | W | W | W | W | W | W | W | W | A | W | A | A | W | A | A | W | A | W | W | A | A | NA | W | A | W | W | A | A | A | A | W | A | W | A | W | A | A | A | NA | A | W | W | W | W | A | A | A | W | A | A | W | W | A | W | W | W | A | A | A | W | W | A | A | W |
| ChrIII | 7 | A | W | W | A | W | W | W | A | A | A | W | A | W | A | W | A | W | A | W | W | A | W | A | W | W | W | W | A | W | W | W | W | W | W | W | W | A | W | A | A | W | A | A | W | A | W | W | A | A | A | W | A | W | W | A | A | A | A | W | A | W | A | W | A | A | A | W | A | W | W | NA | W | A | A | A | W | A | A | W | W | A | W | W | W | A | A | A | W | W | A | A | W |
| ChrIII | 7.1 | A | W | NA | A | W | W | NA | A | A | A | W | A | W | A | W | A | W | A | W | W | A | NA | A | W | W | W | W | A | W | NA | W | NA | W | W | W | W | A | W | A | A | W | A | A | W | A | W | W | A | A | A | W | A | W | W | A | A | A | A | W | A | W | A | W | A | A | A | W | A | W | NA | W | NA | A | A | A | W | A | A | W | W | A | W | W | NA | A | A | A | W | W | A | A | W |
| ChrIII | 7.2 | A | W | NA | A | W | W | W | A | A | A | W | A | W | A | W | A | W | A | W | W | A | W | A | NA | W | W | W | A | NA | W | W | NA | W | W | W | W | A | W | A | A | W | A | A | W | A | W | NA | A | A | A | W | A | W | NA | A | A | A | A | W | A | W | A | W | A | A | A | NA | A | W | W | W | W | A | A | A | W | A | A | NA | W | A | W | W | W | A | A | A | W | W | A | A | NA |
| ChrIII | 7.3 | A | W | W | A | W | W | W | A | A | A | W | A | W | A | W | A | W | A | W | W | A | W | A | W | W | W | W | A | W | W | W | W | W | W | W | W | A | W | A | A | W | A | A | W | A | W | W | A | A | A | W | A | W | W | A | A | A | A | W | A | W | A | W | A | A | A | W | A | W | W | W | W | A | A | A | W | A | A | W | W | A | W | W | W | A | A | A | W | W | A | A | W |
| ChrIII | 7.4 | A | W | W | A | W | W | NA | NA | A | A | W | A | W | A | W | A | W | A | W | W | NA | W | A | W | W | W | W | A | NA | W | W | W | W | NA | W | W | A | W | A | A | W | A | A | W | NA | W | W | A | A | A | W | A | W | W | A | A | A | A | W | A | W | A | W | A | A | A | W | A | W | W | W | W | A | A | A | W | A | A | W | W | A | W | W | W | A | A | A | W | W | A | A | W |
| ChrIII | 7.5 | A | W | W | A | W | W | W | A | A | A | W | A | W | A | W | A | W | A | W | W | A | W | A | W | W | W | W | A | W | W | W | W | W | W | W | W | A | W | A | A | W | A | A | W | A | W | W | A | A | A | W | A | W | W | A | A | A | A | W | A | W | A | W | A | A | A | W | A | W | W | W | W | A | A | A | W | A | A | W | W | A | W | W | W | A | A | A | W | W | A | A | W |
| ChrIII | 7.6 | A | W | W | A | W | W | W | A | A | A | W | A | W | A | W | A | W | A | W | W | A | W | A | W | W | W | W | A | W | W | W | W | W | W | W | W | A | W | A | A | W | A | A | W | A | W | W | A | A | A | W | A | W | W | A | A | A | A | W | A | W | A | W | A | A | A | W | A | W | NA | W | W | A | A | A | W | A | A | W | W | A | W | W | W | A | A | A | W | W | A | A | NA |
| ChrIII | 7.7 | A | W | W | A | W | W | W | A | A | A | W | A | W | A | W | A | W | A | NA | W | A | W | A | W | W | W | W | A | W | W | W | W | W | W | W | W | A | W | A | A | W | A | A | W | A | W | W | A | A | A | W | A | W | W | A | A | A | A | W | A | W | A | W | A | A | A | W | A | W | W | W | W | A | A | A | W | A | A | W | W | A | W | W | W | A | A | A | W | W | A | A | W |
| ChrIII | 7.8 | A | W | W | A | W | W | W | A | A | A | W | A | W | A | W | A | W | A | NA | W | A | W | A | W | W | W | W | A | W | NA | W | W | W | NA | W | W | A | W | A | A | W | A | A | W | A | W | W | A | A | A | W | A | W | NA | A | A | A | A | W | A | W | A | W | A | A | A | W | A | W | W | W | W | A | A | A | W | A | A | W | W | A | W | W | W | A | A | A | W | W | A | A | W |
| ChrIII | 7.9 | A | W | W | A | W | NA | W | A | A | A | W | A | W | A | W | A | W | A | W | W | A | W | A | W | W | W | W | A | W | W | W | W | W | W | W | W | A | W | A | A | W | A | A | W | A | W | W | A | A | A | W | A | W | W | A | A | A | A | W | A | W | A | W | A | A | A | W | A | W | W | W | W | A | A | A | W | A | A | W | W | A | W | W | W | A | A | A | W | W | A | A | W |
| ChrIII | 8 | A | W | W | A | W | W | W | A | A | A | W | A | W | A | W | A | W | A | W | W | A | W | A | W | W | W | W | A | W | W | W | NA | W | W | W | W | A | W | A | A | W | A | A | W | A | W | W | A | A | A | W | A | W | W | A | A | A | A | W | A | W | A | W | A | A | A | W | A | W | W | W | W | A | A | A | W | A | A | W | W | A | W | W | W | A | A | A | W | W | A | A | NA |
| ChrIII | 8.1 | A | W | W | A | W | W | NA | A | A | A | W | A | W | A | W | A | W | A | W | W | A | W | A | W | W | W | W | A | W | W | W | W | W | W | W | W | A | W | A | A | W | A | A | W | A | W | W | A | A | NA | W | A | W | W | A | A | A | A | W | A | W | A | W | A | A | NA | W | A | W | W | W | W | A | A | A | W | A | A | W | W | A | W | W | W | A | A | A | W | W | A | A | W |
| ChrIII | 8.2 | A | W | W | A | W | W | NA | A | A | A | W | A | W | A | W | A | W | A | W | W | A | W | A | W | W | W | W | A | W | W | W | W | NA | W | W | W | A | W | A | A | W | A | A | W | A | W | W | A | A | A | W | A | W | W | A | A | A | A | W | A | W | A | W | A | A | A | NA | A | W | W | W | W | A | A | A | W | A | A | W | W | A | W | W | W | A | A | A | W | W | A | A | W |
| ChrIII | 8.3 | A | W | W | A | W | W | W | A | A | A | W | A | W | A | W | A | W | A | W | W | A | W | A | W | W | W | W | A | W | W | W | W | W | W | W | W | A | W | A | A | W | A | A | W | A | W | NA | A | A | A | W | A | W | W | A | A | A | A | W | A | W | A | W | A | A | A | NA | A | W | W | NA | W | A | A | A | W | A | A | W | W | A | W | W | W | A | A | A | W | W | A | A | NA |
| ChrIII | 8.4 | A | W | W | A | W | W | W | A | A | A | W | A | W | A | W | A | W | A | W | W | A | W | A | W | W | W | W | A | W | W | W | W | W | W | W | W | A | W | A | A | W | A | A | W | A | W | W | A | A | A | W | A | W | W | A | A | A | A | W | A | W | A | W | A | A | A | W | A | W | W | W | W | A | A | A | W | A | A | W | W | A | W | W | W | A | A | A | W | W | A | A | W |
| ChrIII | 8.5 | A | W | W | A | W | W | W | A | A | A | W | A | W | A | W | A | W | A | W | W | NA | W | A | W | W | W | W | A | W | W | W | W | W | W | W | W | A | W | A | A | W | A | A | W | A | W | W | A | A | A | W | A | W | W | A | A | A | A | W | A | W | A | W | A | A | A | W | A | W | W | W | W | A | A | A | W | A | A | W | W | A | W | W | W | A | A | A | W | W | A | A | W |
| ChrIII | 8.6 | A | W | W | A | W | W | W | A | A | A | W | A | W | A | W | A | W | A | W | W | A | W | A | W | W | W | W | A | W | W | W | W | W | W | W | W | A | W | A | A | W | A | A | W | A | W | W | A | A | A | W | A | W | W | A | A | A | A | W | A | W | A | W | A | A | NA | W | A | W | W | W | W | A | A | A | W | A | A | W | W | A | W | W | W | A | A | A | W | W | A | A | W |
| ChrIII | 8.7 | A | W | W | A | W | W | W | A | A | A | W | A | W | A | W | A | W | A | W | W | A | W | A | W | W | W | W | A | W | W | W | W | W | W | W | W | A | W | A | A | W | A | A | W | A | W | W | A | A | A | W | A | W | W | A | A | A | A | W | A | W | A | W | A | A | A | W | A | W | W | W | W | A | A | A | W | A | A | W | W | A | W | W | W | A | A | A | W | W | A | A | W |
| ChrIII | 8.8 | A | W | W | A | W | W | W | A | A | A | W | A | W | NA | NA | A | W | A | NA | W | A | W | A | W | W | W | W | A | W | NA | W | NA | NA | W | W | W | A | W | A | A | W | A | A | W | A | NA | W | A | A | A | W | A | W | NA | A | A | A | A | W | A | W | A | W | A | A | A | NA | A | W | NA | W | W | NA | A | A | W | A | A | W | NA | A | W | W | NA | A | NA | NA | W | NA | A | A | NA |
| ChrIII | 8.9 | A | W | W | A | W | W | W | A | A | A | W | A | W | A | W | A | W | A | W | W | A | W | A | W | W | W | W | NA | W | W | W | W | W | W | W | W | A | W | A | A | W | A | A | W | A | W | W | A | A | A | W | A | W | W | A | A | A | A | W | A | W | A | W | A | A | A | W | A | W | W | W | W | A | A | A | W | A | A | W | W | A | W | W | W | A | A | A | W | W | A | A | W |
| ChrIII | 9 | A | W | W | A | W | W | W | A | A | A | W | A | W | A | W | A | W | A | W | W | NA | W | A | A | W | W | W | A | W | W | W | W | W | W | W | W | A | W | A | A | W | A | A | W | A | W | W | A | A | A | W | A | W | W | NA | A | A | A | W | A | W | A | W | A | A | A | W | A | W | W | W | W | A | A | A | W | A | A | W | W | A | W | W | W | A | A | A | W | W | A | A | W |
| ChrIII | 9.1 | A | W | W | A | W | W | W | A | A | A | W | A | W | A | W | A | W | A | W | W | NA | W | A | A | W | W | W | A | W | W | W | W | W | NA | W | W | A | W | A | A | W | A | A | W | A | W | W | A | A | A | W | A | W | W | A | A | A | A | W | A | W | A | W | A | A | A | NA | A | W | W | NA | W | A | A | A | W | A | A | W | W | A | W | W | W | A | A | A | W | W | A | A | W |
| ChrIII | 9.2 | A | W | W | A | W | W | W | A | A | A | W | A | W | A | W | A | W | A | W | W | NA | W | A | A | W | W | W | A | W | W | W | W | W | W | W | W | A | W | A | A | W | A | A | W | A | W | W | A | A | A | W | A | W | W | A | A | A | A | W | A | W | A | W | A | A | A | W | A | W | W | W | W | A | A | A | W | A | A | W | W | A | W | W | W | A | A | NA | W | W | A | A | NA |
| ChrIII | 9.3 | A | W | W | A | W | W | W | A | A | A | W | A | W | A | W | A | W | A | W | W | A | W | A | A | W | W | W | A | W | W | W | W | W | NA | W | W | A | W | A | A | W | A | A | W | A | W | W | A | A | A | W | A | W | W | A | A | A | A | W | A | W | A | W | A | A | A | W | A | W | W | W | W | A | A | A | W | A | A | W | W | A | W | W | W | A | A | A | W | W | A | A | W |
| ChrIII | 9.4 | A | W | W | A | W | W | W | A | A | A | W | A | W | A | W | A | W | A | W | W | A | W | A | A | W | W | W | A | W | W | W | W | W | W | W | W | A | W | A | A | W | A | A | W | A | W | W | A | A | A | W | A | W | W | A | A | A | A | W | A | W | A | W | A | A | A | W | A | W | W | W | W | A | A | A | W | A | A | W | W | A | W | W | W | A | A | A | W | W | A | A | W |
| ChrIII | 9.5 | A | W | W | A | W | W | NA | A | A | A | W | A | W | A | W | A | W | A | W | W | A | W | A | A | W | W | W | A | W | W | W | W | W | W | W | W | A | W | A | A | W | A | A | W | A | W | W | A | A | A | W | A | W | W | A | A | A | A | W | A | W | A | W | A | A | A | W | A | W | W | W | W | A | A | A | W | A | A | W | W | A | W | W | W | A | A | A | W | W | A | A | W |
| ChrIII | 9.6 | A | W | W | A | W | W | W | A | A | A | W | A | W | A | W | A | W | A | W | W | A | W | A | A | W | W | W | A | W | W | W | W | W | W | W | W | A | W | A | A | W | A | A | W | A | W | W | A | A | A | W | A | W | W | NA | A | A | A | W | A | W | A | W | A | A | A | W | A | W | W | W | W | A | A | A | W | A | A | W | W | A | W | W | W | A | A | A | W | W | A | A | W |
| ChrIII | 9.8 | NA | W | NA | A | W | W | W | A | NA | A | W | A | W | A | W | A | NA | A | NA | NA | A | NA | A | A | NA | W | NA | A | NA | NA | W | NA | W | W | W | W | A | W | A | A | W | A | A | W | A | W | W | A | A | A | W | A | W | W | A | A | A | A | W | A | W | A | NA | A | A | A | W | A | W | W | NA | NA | A | A | A | W | A | A | W | W | NA | NA | W | NA | A | A | A | W | NA | A | A | NA |
| ChrIII | 10 | A | W | W | A | W | W | W | NA | A | A | W | A | W | A | W | A | W | A | W | W | A | W | A | A | W | NA | NA | A | W | W | W | W | W | W | W | W | A | W | A | A | W | A | A | W | A | W | W | A | A | A | W | A | W | W | A | A | A | A | W | A | W | A | W | A | A | A | W | A | W | W | W | W | A | A | A | W | A | A | W | W | NA | W | W | W | A | A | A | W | W | A | A | NA |
| ChrIII | 10.1 | A | W | W | A | W | W | W | A | A | A | W | A | W | A | W | A | W | A | W | W | A | W | A | A | W | W | W | A | W | W | W | W | W | W | W | W | A | W | A | A | W | A | A | W | A | W | W | A | A | A | W | A | W | W | NA | A | A | A | W | A | W | A | W | A | A | A | W | A | W | W | W | W | A | A | A | W | A | A | W | W | A | W | W | W | A | A | A | W | W | A | A | W |
| ChrIII | 10.2 | A | W | W | A | W | W | W | A | A | A | W | A | W | A | W | A | W | A | W | W | A | W | A | A | W | W | W | A | W | W | W | W | W | W | W | W | A | W | A | A | W | A | A | W | A | W | W | A | A | A | W | A | W | W | A | A | A | A | W | A | W | A | W | A | A | A | W | A | W | W | W | W | A | A | A | W | A | A | W | W | A | W | W | W | A | A | A | W | W | A | A | W |
| ChrIII | 10.3 | A | W | W | A | W | W | W | A | A | A | W | A | W | A | W | A | W | A | W | W | A | W | A | A | NA | W | W | A | W | W | W | W | W | W | W | W | A | W | A | A | W | A | A | W | A | W | W | A | A | A | W | A | W | W | A | A | A | A | W | A | W | A | W | A | A | A | W | A | W | W | W | W | A | A | A | W | A | A | W | W | A | W | W | W | A | A | A | W | W | A | A | W |
| ChrIII | 10.4 | A | W | W | A | W | W | NA | A | A | A | W | A | W | A | W | A | W | A | W | W | A | W | A | A | W | W | W | A | NA | W | W | W | W | W | W | W | A | W | A | A | W | A | A | W | A | W | W | A | A | A | W | A | W | W | A | A | A | A | W | A | W | A | W | A | A | A | W | A | W | W | W | W | A | A | A | W | A | A | W | W | A | W | W | W | A | A | A | W | W | A | A | W |
| ChrIII | 10.5 | A | W | W | A | W | W | W | A | A | A | W | A | W | A | W | A | W | A | W | W | A | W | A | A | W | W | W | A | W | W | W | W | NA | NA | W | W | A | W | A | A | W | A | A | W | A | W | W | A | A | A | W | A | W | NA | A | A | A | A | W | A | W | A | W | A | A | A | W | A | W | W | W | W | NA | A | A | W | A | A | W | W | A | W | W | NA | A | A | A | W | W | A | A | NA |
| ChrIII | 10.6 | A | W | W | NA | W | W | W | A | A | A | W | A | W | A | W | A | W | A | NA | W | A | W | NA | A | W | W | W | NA | W | W | W | W | W | NA | W | W | A | W | A | A | W | A | A | W | A | W | W | A | A | A | W | A | W | W | NA | A | A | A | W | A | W | A | W | A | A | NA | W | A | W | W | W | W | A | NA | A | W | A | A | W | W | A | W | W | W | A | A | A | W | W | A | A | W |
| ChrIII | 10.7 | A | W | W | A | W | W | W | A | A | A | W | A | W | A | W | A | W | A | W | W | A | W | A | A | W | W | W | A | W | W | W | W | W | NA | W | W | A | W | A | A | W | A | A | W | A | W | W | A | A | A | W | A | W | W | A | A | A | A | W | A | W | A | W | A | A | A | W | A | W | W | W | W | A | A | A | W | A | A | W | W | A | W | W | NA | A | A | A | W | NA | A | A | W |
| ChrIII | 10.8 | A | W | W | A | W | W | W | A | A | A | W | A | W | A | W | A | W | A | W | W | A | W | A | A | W | W | W | A | W | W | W | W | NA | W | W | W | NA | W | A | A | W | A | A | W | A | NA | W | A | A | A | W | A | W | NA | A | A | A | A | W | A | W | A | W | A | A | A | W | A | W | W | W | NA | A | A | A | W | A | A | W | NA | A | W | W | NA | A | A | NA | W | NA | A | A | W |
| ChrIII | 10.9 | A | W | W | A | W | W | NA | A | A | A | W | A | W | A | W | A | W | A | W | W | A | W | A | A | W | W | W | A | W | NA | W | W | W | W | W | W | A | W | A | A | W | A | A | W | A | W | W | A | A | A | W | A | W | W | A | A | A | A | W | A | W | A | W | A | A | A | W | A | W | W | W | W | A | A | A | W | A | A | W | NA | A | W | W | W | A | A | A | W | NA | A | A | W |
| ChrIII | 11 | A | W | W | A | W | W | W | A | A | A | W | A | W | A | W | A | W | A | W | W | A | W | A | A | W | W | W | A | W | W | W | W | W | W | W | W | A | W | A | A | W | A | A | W | A | W | W | A | A | A | W | A | W | W | A | A | A | A | W | A | W | A | W | A | A | A | W | A | W | W | W | W | A | A | A | W | A | A | W | W | A | W | W | W | A | A | A | W | NA | A | A | W |
| ChrIII | 11.1 | A | W | W | NA | W | W | NA | A | A | A | NA | A | W | A | W | A | W | A | W | W | A | NA | A | A | W | W | W | A | W | W | W | W | NA | W | W | W | A | W | A | A | W | A | A | W | A | NA | NA | A | A | A | W | A | W | W | NA | A | A | A | W | A | W | A | W | A | A | NA | W | A | W | NA | W | W | A | A | A | W | A | A | W | W | A | W | W | NA | A | A | NA | W | W | A | A | W |
| ChrIII | 11.2 | NA | W | W | A | W | W | W | NA | NA | NA | W | NA | W | A | W | NA | NA | NA | W | W | A | W | A | NA | W | W | W | A | W | W | W | W | W | W | W | W | A | W | A | A | W | NA | A | W | NA | W | W | A | NA | NA | W | A | W | W | NA | A | NA | NA | W | A | W | NA | W | NA | NA | NA | W | NA | W | W | W | W | NA | A | NA | W | NA | A | W | W | A | W | W | W | A | A | NA | W | W | A | NA | W |

| | | | | | | | | | | | | | | | | | | | | | | | | | | | | | | | | | | | | | | | | | | | | | | | | | | | | | | | | | | | | | | | | | | | | | | | | | | | | | | | | | | | | | | | | | | | |
|---|---|---|---|---|---|---|---|---|---|---|---|---|---|---|---|---|---|---|---|---|---|---|---|---|---|---|---|---|---|---|---|---|---|---|---|---|---|---|---|---|---|---|---|---|---|---|---|---|---|---|---|---|---|---|---|---|---|---|---|---|---|---|---|---|---|---|---|---|---|---|---|---|---|---|---|---|---|---|---|---|---|---|---|---|---|---|---|---|---|---|---|---|---|
| ChrIII | 11.3 | A | W | W | A | W | W | W | A | A | A | W | A | NA | A | NA | A | W | A | W | W | A | W | A | A | W | W | W | A | NA | W | W | W | W | W | W | W | A | W | A | A | W | A | A | W | A | W | W | A | A | NA | W | A | W | W | A | A | A | A | W | A | W | A | W | A | NA | A | W | A | W | W | W | NA | A | A | A | W | A | A | W | W | A | W | W | NA | A | A | A | W | NA | A | A | NA |
| ChrIII | 11.4 | A | W | W | A | W | NA | W | A | A | A | NA | A | W | A | W | A | W | A | NA | W | A | NA | A | A | W | W | W | NA | W | W | W | NA | W | NA | W | W | A | W | A | A | W | A | A | W | A | W | W | A | A | A | W | A | W | W | A | A | A | A | W | A | W | A | W | A | A | A | W | A | W | W | W | W | A | A | A | W | A | A | NA | W | A | W | W | NA | A | A | A | W | NA | A | A | W |
| ChrIII | 11.5 | A | W | NA | A | W | W | NA | A | A | A | W | A | W | A | W | A | W | A | W | W | A | NA | A | A | W | W | W | A | W | W | W | NA | W | NA | W | W | A | W | A | A | W | A | A | W | A | W | W | A | A | A | W | A | W | W | A | A | A | A | W | A | W | A | W | A | A | A | W | A | W | NA | W | NA | A | A | A | W | A | A | W | NA | A | W | W | W | A | A | A | W | W | A | A | NA |
| ChrIII | 11.6 | A | W | W | A | W | W | W | A | A | A | W | A | W | A | W | A | W | A | NA | W | A | W | A | A | W | W | W | A | W | NA | W | W | NA | W | W | W | A | W | A | A | W | A | A | W | A | W | W | A | A | A | W | A | W | W | A | A | A | A | W | A | W | A | W | A | A | A | W | A | W | W | W | W | A | A | A | W | A | A | NA | W | A | W | W | NA | A | A | A | W | NA | A | A | W |
| ChrIII | 11.7 | A | W | W | A | W | W | W | A | A | A | W | A | W | A | W | A | W | A | W | W | A | W | A | A | W | W | W | A | W | W | W | W | W | NA | W | W | A | W | A | A | W | A | A | W | A | W | W | A | A | A | W | A | W | W | A | A | A | A | W | A | W | A | W | A | A | A | W | A | W | W | W | W | A | A | A | W | A | A | W | W | A | W | W | W | A | A | A | W | W | A | A | W |
| ChrIII | 11.9 | A | W | W | A | W | W | W | A | A | A | W | A | W | A | W | A | W | A | W | W | A | W | A | A | W | W | W | A | W | W | W | W | W | W | W | W | A | W | A | A | W | A | A | W | A | NA | W | A | A | A | W | A | W | W | A | A | A | A | W | A | W | A | W | A | A | A | NA | A | W | W | W | W | A | A | A | W | A | A | W | W | A | W | W | W | A | A | A | W | W | A | A | W |
| ChrIII | 12 | A | W | W | A | W | NA | W | A | A | A | W | A | W | A | NA | A | W | A | NA | W | A | NA | A | A | W | W | NA | A | NA | NA | W | NA | W | NA | W | W | A | W | A | A | W | A | A | W | A | NA | NA | A | A | A | W | A | W | NA | A | A | A | A | W | A | W | A | W | A | A | A | W | A | W | NA | W | NA | NA | A | A | W | A | NA | W | NA | A | W | W | NA | A | NA | A | W | NA | A | A | NA |
| ChrIII | 12.1 | A | W | W | A | W | W | W | A | A | A | W | A | W | A | W | A | W | A | W | W | A | W | A | A | W | W | W | A | W | W | W | W | W | W | W | W | A | W | A | A | W | A | A | W | A | W | W | A | A | A | W | A | W | W | A | A | A | A | W | A | W | A | W | A | A | A | W | A | W | W | W | W | A | A | A | W | A | A | W | W | A | W | W | W | A | A | NA | W | W | A | A | NA |
| ChrIII | 12.3 | A | W | W | A | W | W | W | A | A | A | W | A | W | A | W | A | W | A | W | W | NA | W | A | A | W | W | W | A | W | W | W | W | W | W | W | W | A | W | A | A | W | A | A | W | A | W | W | A | A | A | W | A | W | W | A | A | A | A | W | A | W | A | W | A | A | A | W | A | W | W | W | W | A | A | A | W | A | A | W | W | A | W | W | W | A | A | A | W | W | A | A | W |
| ChrIII | 12.4 | A | W | W | A | W | W | W | NA | A | A | W | A | W | A | NA | A | W | A | W | W | A | W | A | A | W | W | W | A | W | W | W | W | W | W | W | W | A | W | A | A | W | A | A | W | A | W | W | A | A | A | W | A | W | W | A | A | A | A | W | A | W | A | W | A | A | A | W | A | W | W | W | W | A | A | A | W | A | A | W | W | A | W | W | W | A | A | A | W | W | A | A | W |
| ChrIII | 12.5 | A | W | W | A | W | W | W | A | A | A | W | A | NA | A | W | A | W | A | W | W | A | W | A | A | W | W | W | A | W | W | W | W | W | W | W | W | A | W | A | A | W | A | A | W | A | W | W | A | A | A | W | A | W | W | A | A | A | A | W | A | W | A | W | A | A | A | NA | A | W | W | W | W | A | A | A | W | A | A | W | W | A | W | W | W | A | A | A | W | W | A | A | W |
| ChrIII | 12.6 | A | W | W | A | W | W | W | A | A | A | W | A | W | A | W | A | W | A | W | W | NA | W | A | A | W | W | W | A | W | W | W | W | W | W | W | W | A | W | A | A | W | A | A | W | A | W | W | A | A | A | W | A | W | W | A | A | A | A | W | A | W | A | W | A | A | A | W | A | W | W | W | W | A | A | A | W | A | A | W | W | A | W | W | W | A | A | A | W | W | A | A | W |
| ChrIII | 12.7 | A | W | W | A | W | W | W | A | A | A | W | A | W | A | W | A | W | A | W | W | NA | W | A | A | W | W | W | A | W | W | W | W | W | W | W | W | A | W | A | A | W | A | A | W | A | W | W | A | A | A | W | A | W | W | A | A | A | A | W | A | W | A | W | A | A | A | W | A | W | W | W | W | A | A | A | W | A | A | W | W | A | W | W | W | A | A | A | W | W | A | A | W |
| ChrIII | 12.8 | A | W | NA | A | W | W | NA | A | A | A | W | A | W | A | W | A | W | A | W | W | A | W | A | A | W | W | W | A | W | NA | W | W | W | NA | W | W | A | W | A | A | W | A | A | W | A | NA | W | A | A | A | W | A | W | W | A | A | A | A | W | A | W | A | W | A | A | A | W | A | W | NA | W | W | A | A | A | W | A | A | W | W | A | W | W | NA | A | A | NA | W | W | A | A | W |
| ChrIII | 12.9 | A | W | W | A | W | W | W | A | A | A | W | A | W | A | W | A | W | A | W | W | A | W | NA | A | W | W | W | A | W | W | W | W | W | W | W | W | A | W | A | A | W | A | A | W | A | W | W | A | A | A | W | A | W | W | NA | A | A | A | W | A | W | A | W | A | A | A | W | A | W | W | W | W | A | A | A | W | A | A | W | W | A | W | W | W | A | A | A | W | W | A | A | W |
| ChrIII | 13 | A | W | W | A | W | W | W | A | A | A | W | A | W | A | W | A | W | A | W | W | A | W | A | A | W | W | W | A | W | W | W | W | W | W | W | W | A | W | A | A | W | A | A | W | A | W | W | A | A | A | W | A | W | W | A | A | A | A | W | A | W | A | W | A | A | A | W | A | W | W | W | W | A | A | A | W | A | A | W | W | A | W | W | W | A | A | A | W | W | A | A | W |
| ChrIII | 13.1 | A | W | W | A | W | W | NA | A | A | A | NA | A | W | A | W | A | W | A | NA | W | A | W | A | A | W | W | W | NA | W | W | W | W | W | W | W | W | A | W | A | A | W | A | A | W | A | W | W | A | A | A | W | A | W | W | A | A | A | A | W | A | W | A | W | A | A | A | W | A | W | W | W | W | A | A | A | W | A | A | W | W | A | W | W | W | NA | A | A | W | W | NA | A | W |
| ChrIII | 13.2 | A | W | W | A | W | W | W | A | A | A | W | A | W | A | W | A | W | A | W | W | A | W | A | A | W | W | W | A | W | W | W | W | W | W | W | W | A | W | A | A | W | A | A | W | A | W | W | A | A | A | W | A | W | W | A | A | A | A | W | A | W | A | W | A | A | A | W | A | W | W | W | W | A | A | A | W | A | A | W | W | A | W | W | W | A | A | A | W | W | A | A | W |
| ChrIII | 13.3 | A | W | W | A | W | W | W | A | A | A | W | A | W | A | W | A | W | A | W | W | A | W | A | A | W | W | W | A | W | W | W | W | W | W | W | W | A | W | A | A | W | A | A | W | A | W | W | A | A | A | W | A | W | W | A | A | A | A | W | A | W | A | W | A | A | A | W | A | W | W | W | W | A | A | A | W | A | A | W | W | A | W | W | W | A | A | A | W | W | A | A | W |
| ChrIII | 13.4 | A | W | W | A | W | W | W | A | A | A | W | A | W | A | W | A | W | A | W | W | A | W | A | A | W | W | W | A | W | W | W | W | W | NA | W | W | A | W | A | A | W | A | A | W | A | W | W | A | A | A | W | W | W | W | A | A | A | A | W | A | W | A | W | A | A | A | W | A | W | W | W | W | A | A | A | W | A | A | W | W | A | W | W | W | A | A | A | W | W | A | A | W |
| ChrIII | 13.5 | A | W | W | A | W | W | W | NA | NA | A | W | A | W | A | W | A | W | A | W | W | A | W | A | A | W | W | W | A | W | W | W | W | W | W | W | W | A | W | A | A | W | A | A | W | A | W | W | A | A | NA | W | W | W | W | A | A | A | A | W | A | W | A | W | A | A | A | W | A | W | W | W | W | A | A | A | W | A | A | W | W | A | W | W | W | A | A | A | W | W | A | A | W |
| ChrIII | 13.7 | A | W | W | A | W | NA | NA | A | A | A | W | A | W | A | W | A | W | A | NA | W | A | W | A | A | W | W | W | A | W | W | W | W | NA | W | W | W | A | W | A | A | W | A | A | W | A | W | W | A | A | NA | W | W | W | W | A | A | A | A | W | A | W | A | W | A | A | A | W | A | W | W | W | W | A | A | A | W | A | A | W | NA | A | W | W | NA | A | A | A | W | W | A | A | NA |
| ChrIII | 13.8 | A | W | W | A | W | W | W | A | A | A | W | A | W | A | W | A | W | A | W | W | A | W | A | A | W | W | W | A | W | W | W | W | W | W | W | W | A | W | A | A | W | A | A | W | A | W | W | A | A | A | W | W | W | W | A | A | A | A | W | A | W | A | W | A | A | A | W | A | W | W | W | W | A | A | A | W | A | A | W | W | A | W | W | W | A | A | A | W | W | A | A | W |
| ChrIII | 13.9 | NA | W | W | A | W | W | W | NA | NA | A | NA | A | W | A | W | A | W | A | W | W | A | W | A | NA | W | W | W | A | W | W | W | W | W | W | W | W | A | W | A | A | W | A | A | W | NA | W | W | A | NA | A | W | W | W | W | A | A | A | A | W | NA | W | NA | W | A | A | NA | W | A | W | W | W | W | A | A | A | W | NA | A | W | W | A | W | W | W | A | A | A | W | W | A | A | W |
| ChrIII | 14 | NA | W | W | A | W | W | W | NA | NA | A | W | NA | W | A | W | NA | W | A | W | W | NA | W | NA | A | W | W | W | A | W | W | W | W | W | W | W | W | A | W | A | NA | W | A | A | W | NA | W | NA | A | NA | A | W | NA | W | W | A | A | A | NA | W | NA | W | NA | W | NA | A | NA | W | A | W | W | W | W | A | A | A | W | A | A | W | W | NA | W | W | W | A | A | A | W | W | A | A | W |
| ChrIII | 14.1 | A | W | W | A | W | W | W | A | A | A | W | A | W | A | W | A | W | A | W | W | A | W | A | A | W | W | W | A | W | W | W | A | A | W | W | W | A | W | A | W | W | A | A | W | A | W | A | A | A | A | W | W | W | W | A | A | A | A | W | A | W | A | W | A | A | NA | W | A | W | W | W | W | A | A | A | W | A | A | W | W | A | W | W | W | A | A | A | W | W | A | A | W |
| ChrIII | 14.2 | A | W | W | A | W | W | W | A | A | A | W | A | W | A | W | A | W | A | W | W | A | W | A | A | W | W | W | A | W | W | W | A | A | W | W | W | A | W | A | W | W | A | A | W | A | W | A | A | A | A | W | W | W | W | A | A | A | A | W | A | NA | A | W | A | A | A | W | A | W | W | W | W | A | A | A | W | A | A | W | W | A | W | W | W | A | A | A | W | W | A | A | W |
| ChrIII | 14.3 | A | W | W | A | W | W | NA | A | A | A | W | A | W | A | W | A | W | A | W | W | A | W | A | A | W | W | W | A | W | W | W | A | A | W | W | W | A | W | A | W | W | A | A | W | A | W | A | A | A | A | W | W | W | W | A | A | A | A | W | A | W | A | W | A | A | A | W | A | W | W | W | W | A | A | A | W | A | A | W | W | A | W | W | W | A | A | A | W | W | A | A | W |
| ChrIII | 14.4 | A | W | W | A | W | W | W | NA | A | A | W | NA | W | A | W | NA | W | A | W | W | A | W | A | A | W | W | W | A | W | W | W | A | A | W | W | W | A | W | A | W | W | A | A | W | A | W | A | A | NA | NA | W | W | W | W | A | A | A | A | W | A | W | A | W | A | NA | A | W | A | W | W | W | W | A | A | A | W | A | A | W | W | A | W | W | W | A | A | A | W | W | A | A | W |
| ChrIII | 14.5 | NA | W | W | A | W | W | W | NA | NA | NA | W | NA | W | A | W | A | W | A | W | W | A | W | A | A | W | W | W | A | W | W | W | A | A | W | W | W | A | W | A | W | W | A | A | W | A | W | A | A | NA | A | W | W | W | W | NA | A | NA | NA | W | A | W | A | W | A | NA | NA | W | NA | W | W | W | W | A | A | A | W | A | A | W | W | A | W | W | W | A | A | NA | W | W | A | A | W |
| ChrIII | 14.6 | A | W | W | A | W | W | W | A | A | A | W | A | A | A | W | A | W | A | W | W | A | W | A | A | W | W | W | A | W | W | W | A | A | W | W | W | A | W | A | W | W | A | A | W | A | W | A | A | A | W | W | W | W | W | A | A | A | W | W | A | NA | A | W | A | A | A | NA | A | W | W | W | W | A | A | A | W | A | A | W | W | A | W | W | W | A | A | A | W | W | A | A | W |
| ChrIII | 14.7 | A | W | W | A | W | W | W | A | A | A | W | A | A | A | W | A | W | A | W | W | A | W | A | A | W | W | W | A | W | W | W | A | A | W | W | W | A | W | A | W | W | A | A | W | A | W | A | A | A | W | W | W | W | W | A | A | A | W | W | A | W | A | W | A | A | A | NA | A | W | W | W | W | A | A | A | W | A | A | W | W | A | W | W | W | A | A | A | W | W | A | A | W |
| ChrIII | 14.8 | A | W | W | A | W | W | W | A | A | A | W | A | A | A | W | A | W | A | W | W | A | W | A | A | W | W | W | A | W | W | W | A | A | W | W | W | A | W | A | W | W | A | A | W | A | W | A | A | A | W | W | W | W | W | A | A | A | W | W | A | W | A | W | A | A | A | W | A | W | W | W | W | A | A | A | W | A | A | W | W | A | W | W | W | A | A | A | W | W | A | A | W |
| ChrIII | 14.9 | A | W | W | A | W | W | W | A | A | A | W | A | A | A | W | A | W | A | W | W | A | W | A | A | W | W | W | A | W | W | W | A | A | W | W | W | A | W | A | W | W | A | A | W | A | W | A | A | A | W | W | W | W | W | A | A | A | W | W | A | W | A | W | A | A | A | W | A | W | W | W | W | A | A | A | W | A | A | W | W | A | W | W | W | A | A | A | W | W | A | A | W |
| ChrIII | 15 | A | W | W | A | W | W | W | A | A | A | W | A | A | A | W | A | W | A | W | W | A | W | A | A | W | W | W | A | W | W | W | A | A | W | W | W | A | W | A | W | W | A | A | W | A | W | A | A | A | W | W | W | W | W | A | A | A | W | W | A | W | A | W | A | A | W | W | A | W | W | W | W | A | A | A | W | A | A | W | W | A | W | W | W | A | A | A | W | W | A | A | NA |
| ChrIII | 15.1 | A | W | W | A | W | W | W | A | A | A | W | A | A | A | W | A | W | A | W | W | A | W | A | A | W | W | W | A | W | A | W | A | A | W | W | W | A | W | A | W | W | A | A | W | A | W | A | A | A | W | W | W | W | W | A | A | A | W | W | A | W | A | W | A | A | W | W | A | W | W | W | W | A | A | A | W | A | A | W | W | A | W | W | W | A | A | A | W | W | A | A | W |
| ChrIII | 15.2 | A | W | W | A | W | W | W | NA | A | A | W | A | A | W | NA | A | W | A | W | W | A | W | A | A | W | W | W | A | W | A | W | A | A | W | W | W | A | W | A | W | W | A | A | W | A | W | A | A | A | W | W | W | W | W | A | A | A | W | W | A | W | A | W | A | A | W | W | A | W | W | W | W | A | A | A | W | A | A | W | W | A | W | W | W | NA | A | A | W | W | A | A | W |
| ChrIII | 15.3 | A | W | W | A | W | W | W | A | A | A | W | A | A | W | W | A | W | A | W | W | A | W | A | A | W | W | W | A | W | A | W | A | A | W | W | W | A | W | A | W | W | A | A | W | A | W | A | A | A | W | W | W | W | W | A | A | A | W | W | A | W | A | W | A | A | W | NA | A | W | W | W | W | A | A | A | W | A | A | W | W | A | W | W | W | W | A | A | W | W | A | A | W |
| ChrIII | 15.5 | A | W | W | A | W | W | W | A | A | A | W | A | A | W | W | A | W | A | W | W | A | W | A | A | W | W | W | A | W | A | W | A | A | W | W | W | A | W | A | W | W | A | A | W | A | W | A | A | A | W | W | W | W | W | A | A | A | W | W | A | W | A | W | A | A | W | W | A | A | W | W | W | A | A | A | W | A | W | W | A | A | W | W | W | W | A | A | W | W | A | A | W |
| ChrIII | 15.6 | A | W | W | A | W | W | W | A | A | A | W | A | A | W | W | A | W | A | W | W | A | W | A | A | W | W | W | A | W | A | W | A | A | W | W | W | A | W | A | W | W | A | A | W | A | W | A | A | A | W | W | W | W | W | A | A | A | W | W | A | W | A | W | A | A | W | W | A | A | W | W | W | A | A | A | W | A | W | W | A | A | W | W | W | W | A | A | W | W | A | A | W |
| ChrIII | 15.7 | A | W | W | A | W | W | W | A | A | A | W | A | A | W | W | A | W | A | W | W | A | W | A | A | W | W | W | A | W | A | W | A | A | NA | W | W | A | W | A | W | W | A | A | W | A | W | A | A | A | W | W | W | W | W | A | A | A | W | W | A | W | A | W | A | A | W | NA | A | A | W | W | W | A | A | A | W | A | NA | W | A | A | W | W | W | W | A | A | W | W | A | A | W |
| ChrIII | 15.8 | A | W | W | A | W | W | W | A | A | A | W | A | A | W | W | A | W | A | W | W | A | W | A | A | W | W | W | A | W | A | W | A | A | W | W | W | A | W | A | W | W | A | A | W | A | W | A | A | A | W | W | W | W | W | A | A | A | W | W | A | W | A | W | A | A | W | W | A | A | W | W | W | A | A | A | W | A | W | W | A | A | W | W | W | W | A | A | W | W | A | A | W |
| ChrIII | 15.9 | A | W | W | A | W | W | W | A | A | A | W | A | A | W | W | A | W | A | W | W | A | W | A | A | W | W | W | A | W | A | W | A | A | W | W | W | A | W | A | W | W | A | A | W | A | W | A | A | A | NA | W | W | W | W | A | A | A | W | W | A | W | A | W | A | A | W | W | A | A | W | W | W | A | A | A | W | A | W | W | A | A | W | W | W | W | A | A | W | W | A | A | W |
| ChrIII | 16 | A | W | W | A | W | W | W | A | A | A | W | A | A | NA | W | A | W | A | W | W | A | W | A | A | W | W | W | A | W | A | W | A | A | NA | W | W | A | W | A | W | W | A | A | W | A | W | A | A | A | W | W | W | W | W | A | A | A | W | W | A | W | A | W | A | A | W | W | A | A | W | W | W | A | A | A | W | A | W | W | A | A | W | W | W | W | A | NA | W | W | A | A | W |
| ChrIII | 16.1 | A | W | W | A | W | W | W | A | A | A | W | A | A | W | W | A | W | A | W | W | NA | W | NA | A | W | W | W | A | W | A | W | A | A | W | W | W | A | W | A | W | W | A | A | W | A | W | A | A | A | W | W | W | W | W | A | A | A | W | W | A | W | A | W | A | A | W | W | A | A | W | W | W | A | A | A | W | A | W | W | A | A | W | W | W | W | A | A | W | W | A | A | W |
| ChrIII | 16.2 | A | W | W | A | W | W | W | A | A | A | W | A | A | W | W | A | W | A | W | W | A | W | A | A | NA | W | W | A | W | A | W | A | A | W | W | W | A | W | A | W | W | A | A | W | A | W | A | A | A | W | W | W | W | W | A | A | A | W | W | A | W | A | W | A | A | W | W | A | A | W | W | W | A | A | A | W | A | W | W | A | A | W | W | W | W | A | A | W | W | A | A | W |
| ChrIII | 16.3 | A | W | W | A | W | W | W | A | A | A | W | A | A | W | W | A | W | A | W | W | A | W | A | A | W | W | W | A | W | A | W | A | A | W | W | W | A | W | A | W | W | A | A | W | A | W | A | A | A | W | W | W | W | W | A | A | A | W | W | A | W | A | W | A | A | W | W | A | A | W | W | W | A | A | A | W | A | W | W | A | A | W | W | W | W | A | A | W | W | A | A | W |
| ChrIII | 16.4 | A | W | W | A | W | W | W | A | A | A | W | A | A | W | W | A | W | A | W | W | A | W | A | A | W | W | W | A | W | A | W | A | A | W | W | W | A | W | A | W | W | A | A | W | A | A | A | A | A | W | W | W | W | W | A | A | A | W | W | A | W | A | W | A | A | W | W | A | A | W | W | W | A | A | A | W | A | W | W | A | A | W | W | W | W | A | A | W | W | A | A | W |
| ChrIII | 16.5 | A | W | W | A | W | W | W | A | A | A | W | A | A | W | W | A | W | A | W | W | A | W | A | A | W | W | W | A | W | A | W | A | A | W | W | W | A | W | A | W | W | A | A | W | A | A | A | A | A | W | W | W | W | W | A | A | A | W | W | A | W | A | W | A | A | W | W | A | A | W | W | W | A | W | A | W | W | W | W | A | A | W | W | W | W | A | A | W | W | A | A | W |
| ChrIII | 16.6 | A | W | W | A | W | W | W | A | A | A | W | A | A | W | W | A | NA | A | W | W | A | A | A | A | W | W | W | A | W | A | W | A | A | W | W | W | A | W | A | W | W | A | A | W | A | A | A | A | A | W | W | W | W | W | NA | A | A | W | W | A | W | A | W | A | A | W | W | A | A | W | W | W | A | W | A | W | W | W | W | A | A | NA | NA | W | W | A | A | W | W | A | A | A |
| ChrIII | 16.7 | A | W | W | A | W | W | NA | A | A | A | W | A | A | W | NA | A | A | A | W | W | A | A | A | A | W | W | W | A | W | A | W | A | A | W | W | W | A | W | A | W | W | A | A | W | A | A | A | A | A | W | W | W | W | W | A | A | A | W | W | A | W | A | W | A | A | W | W | A | A | W | W | W | A | W | A | W | W | W | W | A | A | A | A | W | W | A | A | W | W | A | A | A |
| ChrIII | 16.8 | A | W | W | A | W | W | W | NA | A | A | W | A | A | W | W | A | A | A | W | W | A | A | A | A | W | W | W | A | W | A | NA | A | A | W | W | W | A | W | A | W | W | A | A | W | A | A | A | A | A | W | W | W | W | W | A | A | A | W | W | A | W | A | W | A | A | W | W | A | A | W | W | W | A | W | A | W | W | W | W | A | A | A | A | W | W | A | A | W | W | A | A | A |
| ChrIII | 16.9 | A | W | W | A | W | W | W | A | A | A | W | A | A | W | W | A | A | A | W | W | A | A | A | A | W | W | W | A | W | A | A | A | A | W | W | W | A | W | A | W | W | A | A | W | A | A | A | A | A | W | W | W | W | W | A | A | A | W | W | A | W | A | W | A | A | W | W | A | A | W | W | W | A | W | A | W | W | W | W | A | A | A | A | W | W | A | A | W | W | A | A | A |
| ChrIII | 17 | A | W | W | A | W | W | W | A | A | A | W | A | A | W | W | A | A | A | W | W | A | A | A | A | W | W | W | A | W | A | A | A | A | W | W | W | A | W | A | W | W | A | A | W | A | A | A | A | A | W | W | W | W | W | A | A | A | W | W | A | W | A | W | A | A | W | W | A | A | W | W | W | A | W | A | W | W | W | W | A | A | A | A | W | W | A | A | W | W | A | A | A |
| ChrIII | 17.1 | A | W | W | NA | W | W | W | A | A | A | W | A | A | NA | W | A | A | A | NA | W | A | A | A | A | W | W | W | A | W | A | A | A | A | W | W | W | A | W | A | W | W | A | A | W | A | A | A | A | A | W | W | W | W | W | A | A | A | W | W | A | NA | NA | W | A | A | W | W | A | A | NA | W | W | A | W | A | W | W | W | W | A | A | A | A | W | W | A | A | W | W | A | A | A |
| ChrIII | 17.2 | A | W | W | A | W | W | W | A | A | A | W | A | A | W | W | A | A | A | W | W | A | A | A | A | W | W | W | A | W | A | A | A | A | W | W | W | A | W | A | W | W | A | A | W | A | A | A | A | A | W | W | W | W | W | A | A | A | W | W | A | W | A | W | A | A | W | NA | A | A | W | W | W | A | W | A | W | W | W | W | A | A | A | A | W | W | A | A | W | W | A | A | A |
| ChrIII | 17.3 | A | NA | NA | A | W | NA | W | W | A | A | W | A | A | W | W | A | A | A | W | W | W | A | W | A | W | W | W | A | W | A | A | A | NA | W | A | W | NA | W | A | W | W | A | A | W | A | A | A | A | A | NA | W | W | W | W | A | A | A | W | W | A | NA | A | W | A | A | W | W | A | A | W | W | W | A | W | A | W | W | NA | W | A | A | A | A | NA | W | A | NA | W | W | A | A | A |
| ChrIII | 17.4 | A | W | A | A | W | W | W | W | A | A | W | A | A | W | NA | A | A | A | W | W | W | A | W | A | W | W | W | A | W | A | A | A | A | W | A | W | A | W | A | W | W | A | A | W | A | A | A | A | A | NA | W | W | W | W | A | A | A | W | W | A | A | A | W | A | A | W | W | A | A | W | W | W | A | W | A | W | W | W | W | A | A | A | A | W | W | A | A | W | W | A | A | A |
| ChrIII | 17.5 | A | W | A | A | W | W | W | W | A | A | W | A | A | W | W | A | A | A | W | W | W | A | W | A | W | W | W | NA | W | A | A | A | NA | W | A | W | A | W | A | W | W | A | A | W | A | A | A | A | A | W | W | W | W | W | A | A | A | W | W | A | A | A | W | A | A | W | W | A | A | W | W | W | A | W | A | W | W | W | W | A | A | A | A | W | W | A | A | W | W | A | A | A |
| ChrIII | 17.6 | A | NA | A | A | W | NA | W | W | A | A | W | A | A | W | W | A | A | A | W | W | W | A | W | A | W | W | W | A | W | A | A | A | A | W | A | W | A | W | W | W | W | A | A | W | A | A | A | A | A | W | W | W | W | W | A | A | A | W | W | A | A | A | W | A | A | W | W | A | A | W | W | W | A | W | A | W | W | W | W | A | A | A | A | W | W | A | NA | W | W | A | A | A |
| ChrIII | 17.7 | A | A | A | A | W | A | W | W | A | A | W | A | A | W | W | A | A | A | W | W | W | A | W | A | W | W | W | A | W | A | A | A | A | W | A | W | A | W | W | W | W | A | A | W | A | A | A | A | A | W | W | W | W | W | A | A | A | W | W | A | A | A | W | A | A | W | NA | A | A | W | W | W | A | W | A | W | W | W | W | A | A | A | A | W | W | A | A | W | W | A | A | A |
| ChrIII | 17.8 | A | A | A | A | W | A | W | W | NA | A | W | A | A | W | W | A | A | A | W | W | W | A | W | A | NA | W | W | A | W | A | A | A | A | W | A | W | A | W | NA | W | W | A | A | W | NA | A | A | A | NA | W | W | W | W | NA | NA | A | A | W | W | A | A | A | W | A | A | W | W | A | A | NA | W | W | A | W | A | W | W | NA | W | A | A | A | A | W | W | A | A | W | W | A | A | A |
| ChrIII | 17.9 | A | A | A | A | W | A | W | W | A | A | W | A | A | W | W | A | A | A | W | W | W | A | W | A | NA | W | W | A | W | A | A | A | NA | NA | A | W | A | W | W | W | W | A | A | W | A | A | A | A | A | W | W | W | W | W | A | A | A | W | W | A | NA | A | W | A | A | W | NA | A | A | W | W | W | A | W | A | W | W | NA | W | A | A | A | A | W | W | A | A | W | W | A | A | NA |
| ChrIII | 18 | A | A | A | A | W | A | W | W | A | A | W | A | A | W | W | A | A | A | W | W | W | A | W | A | W | W | W | A | W | A | A | A | A | W | A | W | A | W | W | W | W | A | A | W | A | A | A | A | A | W | W | W | W | W | A | A | A | W | W | A | A | A | W | A | A | W | W | A | A | W | W | W | A | W | A | W | W | W | W | A | A | A | A | W | W | A | A | W | W | A | A | A |
| ChrIII | 18.1 | NA | A | A | A | W | A | W | W | NA | A | W | NA | A | W | W | NA | A | NA | W | W | W | A | W | A | W | W | W | NA | W | A | A | A | A | W | A | W | A | W | W | W | W | A | A | W | A | A | A | A | A | W | W | W | W | W | A | A | A | W | W | A | A | A | W | A | NA | W | W | A | A | W | W | W | A | W | A | W | W | W | W | A | A | A | NA | W | W | A | A | W | W | A | A | A |
| ChrIII | 18.2 | A | A | A | A | W | A | W | W | A | A | W | A | A | W | NA | A | A | A | W | W | W | A | W | A | W | W | NA | A | W | A | A | A | A | W | A | W | A | W | W | W | W | A | A | W | A | A | A | A | A | NA | W | W | NA | NA | A | A | A | W | W | A | A | A | W | A | A | W | W | A | A | W | W | W | A | W | A | W | W | W | W | A | A | A | A | W | W | A | A | W | W | A | A | A |
| ChrIII | 18.3 | A | A | A | A | W | A | W | W | A | A | W | A | A | W | W | A | A | A | W | W | W | A | W | A | NA | W | W | A | W | A | A | A | NA | W | A | W | A | W | W | W | W | A | A | W | A | A | A | A | A | W | W | W | W | W | A | A | A | W | W | A | A | A | W | A | A | W | W | A | A | W | W | W | A | W | A | W | W | W | W | A | A | A | A | W | W | A | A | W | W | A | A | A |
| ChrIII | 18.4 | A | A | A | A | W | NA | W | W | A | A | W | A | A | W | W | A | A | A | NA | W | W | A | W | A | NA | W | NA | A | W | A | A | A | NA | NA | A | W | NA | W | NA | W | W | A | A | W | A | A | A | A | A | W | W | W | NA | NA | A | A | A | W | W | A | A | NA | W | A | A | NA | NA | A | NA | NA | W | W | NA | W | A | W | NA | NA | NA | A | A | A | A | NA | W | A | NA | W | NA | A | A | A |
| ChrIII | 18.7 | A | A | A | NA | W | A | W | W | A | A | W | A | A | W | W | A | A | W | W | W | W | A | W | A | W | W | A | A | A | A | A | A | A | W | A | W | A | W | W | W | W | A | A | NA | A | A | A | W | NA | NA | W | W | W | W | A | A | A | W | W | A | A | A | W | A | A | W | W | A | A | W | W | W | A | W | W | A | W | W | W | A | A | A | A | W | W | A | W | W | W | A | A | A |
| ChrIII | 18.8 | A | A | A | W | W | A | W | W | A | A | W | A | A | W | W | A | A | W | W | W | W | A | W | A | A | W | A | A | A | A | A | A | A | NA | A | W | W | W | W | W | W | A | A | A | A | A | A | W | W | W | W | W | W | W | A | A | A | W | W | A | A | A | W | A | A | W | W | A | A | W | W | W | A | W | W | A | W | W | W | A | A | A | A | W | W | A | W | W | W | A | A | A |
| ChrIII | 18.9 | A | A | A | W | W | A | W | W | A | A | W | A | A | W | W | A | A | W | W | W | W | A | W | A | A | W | A | A | A | A | A | A | A | W | A | W | W | W | W | W | W | A | A | A | A | A | A | W | W | W | W | W | W | W | A | A | A | W | W | A | A | A | W | A | A | W | W | A | A | W | W | W | A | W | W | A | W | W | W | A | A | A | A | W | W | A | W | W | W | A | A | A |
| ChrIII | 19 | A | A | A | W | W | A | W | W | A | A | W | A | A | W | W | A | A | W | W | W | W | A | W | A | A | W | A | A | A | A | A | A | A | NA | A | W | W | W | W | W | W | A | A | A | A | A | A | W | W | W | W | W | W | W | A | A | A | W | W | A | A | A | W | A | A | W | W | A | A | W | W | W | A | W | W | A | W | W | W | A | A | A | A | W | W | A | W | W | W | A | A | A |
| ChrIII | 19.1 | A | A | A | W | W | A | W | W | A | A | W | A | A | W | W | A | A | W | W | W | W | A | W | A | A | W | A | A | A | A | A | A | A | W | A | W | W | W | W | W | W | A | A | A | A | A | A | W | W | W | W | W | W | W | A | A | A | W | W | A | A | A | W | A | A | W | W | A | A | W | W | W | A | W | W | A | W | W | W | A | A | A | A | W | W | A | W | W | W | A | A | A |
| ChrIII | 19.4 | A | A | A | W | W | A | W | W | A | A | W | A | A | W | W | A | A | W | W | W | W | A | W | A | A | W | A | A | A | A | A | A | A | W | A | W | W | W | W | W | W | A | A | A | A | A | A | W | W | W | W | W | W | W | A | A | A | W | W | A | A | A | W | A | A | W | W | A | A | W | W | W | A | W | W | A | W | W | W | A | A | A | A | W | W | A | W | W | W | A | A | A |
| ChrIII | 19.5 | A | A | A | W | W | A | W | W | A | A | W | A | A | W | W | A | A | W | W | W | W | A | W | A | A | W | A | A | A | A | A | A | A | W | A | W | W | W | W | W | W | A | A | A | A | A | A | W | W | W | W | W | W | W | A | A | A | W | W | A | A | A | W | A | A | W | W | A | A | W | W | W | A | W | W | A | W | W | W | A | A | A | A | W | W | A | W | W | W | A | A | A |
| ChrIII | 19.6 | A | A | A | W | W | A | W | W | A | A | W | A | A | W | W | A | A | W | W | W | W | A | W | A | A | W | A | A | A | A | A | A | A | W | A | W | W | W | W | W | W | A | A | A | A | A | A | W | W | W | W | W | W | W | A | A | A | W | W | A | A | A | W | A | A | W | W | A | A | W | W | W | A | W | W | A | W | W | W | A | A | A | A | W | W | A | W | W | W | A | A | A |
| ChrIII | 19.7 | A | A | A | W | W | A | W | W | A | A | W | A | A | W | W | A | A | W | W | W | W | A | W | A | A | W | A | A | A | A | A | A | A | W | A | W | W | W | W | W | W | A | A | A | A | A | A | W | W | W | W | W | W | W | A | A | A | W | W | A | A | A | W | A | A | W | W | A | A | W | W | W | A | W | W | A | W | W | W | A | A | A | A | W | W | A | W | W | W | A | A | A |
| ChrIII | 19.8 | A | A | A | W | W | A | W | W | A | A | W | A | A | W | W | NA | A | W | W | W | NA | A | W | A | A | W | A | A | A | A | A | A | A | W | A | W | W | NA | W | W | W | A | A | A | A | A | A | W | W | W | W | W | W | W | A | A | A | W | W | A | A | A | W | A | NA | W | W | A | A | W | W | W | A | W | W | A | W | W | W | A | A | A | NA | W | W | A | W | W | W | A | A | A |
| ChrIII | 19.9 | A | A | A | W | W | A | W | W | A | A | W | A | A | W | W | A | A | W | W | NA | W | A | W | A | A | W | A | A | A | A | A | A | A | W | A | W | W | A | W | W | W | A | A | A | A | A | A | W | W | W | W | W | W | W | A | A | A | W | W | A | A | A | W | A | A | NA | W | A | A | W | W | W | A | W | NA | A | W | W | W | A | A | A | A | W | W | A | W | W | W | A | A | A |
| ChrIII | 20 | A | A | A | W | W | A | W | W | A | A | W | NA | A | W | W | A | A | W | W | W | W | A | W | A | A | W | A | A | A | A | A | A | A | W | A | W | W | A | W | W | W | A | A | A | A | A | A | W | W | W | W | W | W | W | A | A | A | W | W | A | A | A | W | A | A | W | W | A | A | W | W | W | A | W | W | A | W | W | W | A | A | A | A | W | W | A | W | W | W | A | A | A |
| ChrIII | 20.1 | A | A | A | W | W | A | W | W | A | A | W | W | A | W | W | A | A | W | W | W | W | A | W | A | A | W | A | A | A | A | A | A | A | W | A | W | W | A | W | W | W | A | A | A | A | A | A | W | W | W | W | W | W | W | A | A | A | W | W | A | A | A | W | A | A | W | W | A | A | W | W | W | A | W | W | A | W | W | W | A | A | A | A | W | W | A | W | W | W | A | A | A |
| ChrIII | 20.2 | A | A | A | W | W | A | W | W | A | A | W | W | A | W | W | NA | A | W | W | W | W | A | W | A | A | W | A | A | A | A | A | A | A | W | A | W | W | A | W | W | W | A | A | A | A | A | A | W | W | W | W | W | W | W | A | A | A | W | W | A | A | A | W | A | A | W | W | A | A | W | W | W | A | W | W | A | W | W | W | A | A | A | A | W | W | A | W | W | W | A | A | A |
| ChrIII | 20.3 | A | A | A | W | W | A | W | W | A | A | W | W | A | W | W | A | A | W | W | W | W | A | W | A | A | W | A | A | A | A | A | A | A | W | A | W | W | A | W | W | W | A | A | A | A | A | A | W | W | W | W | W | W | W | A | A | A | W | W | A | A | A | W | A | A | W | W | A | A | W | W | W | NA | W | W | A | W | W | W | A | A | A | A | W | W | A | W | W | W | A | A | A |
| ChrIII | 20.5 | A | A | A | W | W | A | W | W | A | A | W | W | A | W | W | A | A | W | W | W | W | A | W | A | A | W | A | NA | A | A | A | A | A | W | A | W | W | A | W | W | W | A | A | A | A | A | A | W | W | W | W | W | W | W | A | A | A | W | W | A | A | A | W | A | A | W | W | A | A | W | W | W | A | W | W | A | W | W | W | A | W | A | A | W | W | A | W | W | W | A | A | A |
| ChrIII | 20.6 | A | A | A | W | W | A | W | W | A | A | W | W | A | W | W | A | A | W | W | W | W | A | W | A | A | W | A | A | A | A | A | A | A | W | A | W | W | A | W | W | W | A | A | A | A | A | A | W | W | NA | W | W | W | W | A | A | A | W | W | A | A | A | W | A | A | W | W | A | A | W | W | W | A | W | W | A | W | W | W | A | W | A | A | W | W | A | W | W | W | A | A | A |
| ChrIII | 20.7 | A | A | A | W | W | A | W | W | A | A | W | W | A | W | W | A | A | W | W | W | W | A | W | A | A | W | A | A | A | A | A | A | A | W | A | W | W | A | W | W | W | A | A | A | A | A | A | W | W | W | W | W | W | W | A | A | A | W | W | A | A | A | W | A | A | W | W | A | A | W | W | W | A | W | W | A | W | W | W | A | W | A | A | W | W | A | W | W | W | A | A | A |
| ChrIII | 20.8 | A | A | A | W | W | A | W | W | A | A | W | W | A | W | W | A | A | W | W | W | W | A | W | A | A | W | A | A | A | A | A | A | A | W | A | W | W | A | W | W | W | A | A | A | A | A | A | W | W | W | W | W | W | W | A | A | A | W | W | A | A | A | W | A | A | W | W | A | A | W | W | W | A | W | W | A | W | W | W | A | W | A | A | W | W | A | W | W | W | A | A | A |
| ChrIII | 20.9 | A | A | A | W | W | A | W | W | A | A | W | W | A | W | W | A | A | W | W | W | W | A | W | A | A | W | A | A | A | A | A | A | NA | W | A | W | W | A | W | W | W | A | A | A | A | A | A | W | W | W | W | W | W | W | NA | A | A | W | W | A | A | A | W | A | A | W | W | A | A | W | W | W | A | W | W | A | W | W | W | A | W | A | A | W | W | A | NA | W | W | A | A | A |

| | | | | | | | | | | | | | | | | | | | | | | | | | | | | | | | | | | | | | | | | | | | | | | | | | | | | | | | | | | | | | | | | | | | | | | | | | | | | | | | | | | | | | | | | | | | | | |
|---|---|---|---|---|---|---|---|---|---|---|---|---|---|---|---|---|---|---|---|---|---|---|---|---|---|---|---|---|---|---|---|---|---|---|---|---|---|---|---|---|---|---|---|---|---|---|---|---|---|---|---|---|---|---|---|---|---|---|---|---|---|---|---|---|---|---|---|---|---|---|---|---|---|---|---|---|---|---|---|---|---|---|---|---|---|---|---|---|---|---|---|---|---|
| ChrIII | 21 | A | A | A | W | W | A | W | W | A | A | W | W | A | W | W | A | A | W | W | NA | W | A | W | A | A | W | A | A | A | A | A | A | A | W | A | W | NA | A | W | W | W | A | A | A | A | A | A | W | W | W | W | W | W | W | A | A | A | W | W | A | A | A | W | A | A | W | W | A | A | W | W | W | A | W | W | A | W | W | W | A | W | A | A | W | W | A | W | W | W | A | A | A |
| ChrIII | 21.2 | A | A | A | W | W | A | W | W | A | A | W | W | A | W | W | A | A | W | W | W | W | A | W | A | A | W | A | A | A | A | A | A | A | W | A | W | W | A | W | W | W | A | A | A | A | A | A | W | W | W | W | W | W | W | A | A | A | W | W | A | A | A | W | A | A | W | W | A | A | W | W | W | A | W | W | A | W | W | W | A | W | A | A | W | W | A | W | W | W | A | A | A |
| ChrIII | 21.3 | A | A | A | W | W | A | W | W | A | A | W | W | A | W | W | A | A | W | W | NA | W | A | W | A | A | W | A | A | A | A | A | A | A | W | A | W | W | A | W | W | W | A | A | A | A | A | A | W | W | W | W | W | W | W | A | A | A | W | W | A | A | A | W | A | A | W | W | A | A | W | W | W | A | W | W | A | W | W | W | A | W | A | A | W | W | A | W | W | W | A | A | A |
| ChrIII | 21.4 | A | A | A | W | W | A | W | W | A | A | W | W | A | W | W | A | NA | W | W | W | W | A | W | A | A | W | A | A | A | A | A | A | A | W | A | W | W | A | W | W | W | A | A | A | A | A | A | W | W | W | W | W | W | W | A | A | A | W | W | A | A | A | W | A | A | W | W | A | A | W | W | W | A | W | W | A | W | W | W | A | W | A | A | W | W | A | W | W | W | A | A | A |
| ChrIII | 21.5 | A | A | A | W | W | A | W | W | NA | NA | W | W | NA | W | W | A | A | W | W | W | W | A | W | A | A | W | A | A | A | A | NA | A | A | W | NA | W | W | A | W | W | W | NA | A | A | NA | A | NA | W | W | W | W | W | W | W | NA | NA | NA | W | W | A | A | NA | W | NA | NA | W | W | NA | NA | W | W | W | A | W | W | A | W | W | W | A | W | NA | A | W | W | A | W | W | W | A | A | A |
| ChrIII | 21.6 | A | A | A | W | W | A | W | NA | A | A | W | W | A | W | W | NA | A | W | W | W | W | A | W | NA | A | W | A | A | NA | A | A | A | A | W | NA | W | W | NA | W | W | W | A | A | NA | A | A | NA | W | W | W | W | W | W | W | NA | A | A | W | W | NA | A | A | W | A | NA | W | W | A | A | W | W | W | NA | W | W | A | W | W | W | A | W | A | A | W | W | A | W | W | W | A | NA | A |
| ChrIV | 0.3 | W | W | W | A | A | W | NA | W | W | W | A | W | A | A | W | A | A | A | A | A | A | W | A | W | A | A | NA | A | W | A | A | A | A | A | A | A | A | A | A | A | A | A | W | W | W | A | W | A | W | W | A | W | A | A | W | W | W | W | W | W | W | W | W | A | W | W | NA | W | NA | W | W | A | W | A | A | W | W | A | W | W | A | A | W | W | A | A | W | A | W | W | A | W |
| ChrIV | 0.4 | W | W | W | A | A | W | A | W | W | W | A | W | A | A | W | A | A | A | A | A | A | W | A | W | A | A | A | A | W | A | A | A | A | A | A | A | A | A | A | A | A | A | W | W | W | A | W | A | W | W | A | W | A | A | W | W | W | W | W | W | W | W | W | A | W | W | A | W | A | W | W | A | W | A | A | W | W | A | W | W | A | A | W | W | A | A | W | A | W | W | A | W |
| ChrIV | 0.5 | W | W | W | NA | A | W | NA | W | W | W | NA | W | A | NA | W | A | A | A | A | A | A | W | NA | W | A | A | NA | A | W | A | NA | A | NA | A | A | NA | A | A | A | NA | A | A | NA | W | W | NA | W | A | W | W | NA | W | NA | NA | W | W | W | W | W | W | W | W | W | A | W | W | A | W | A | W | W | NA | W | A | A | W | W | A | W | W | A | A | W | W | A | NA | W | A | W | W | NA | W |
| ChrIV | 0.6 | W | W | W | A | A | W | A | W | W | W | A | W | A | A | NA | A | A | A | A | A | A | W | A | W | A | A | A | A | W | A | A | A | A | A | A | A | A | A | A | A | A | A | W | W | W | A | W | A | W | W | A | W | A | A | W | W | W | W | W | W | W | W | W | A | W | W | A | W | A | W | W | A | W | A | A | W | W | A | W | W | A | A | W | W | A | A | W | A | W | W | A | W |
| ChrIV | 0.7 | W | W | W | A | A | W | A | W | W | W | A | W | A | A | W | A | A | A | A | A | A | W | A | W | A | A | A | A | W | A | A | A | A | A | A | A | A | A | A | A | A | A | W | W | W | A | W | A | W | W | A | W | A | A | W | W | W | W | W | W | W | W | W | A | W | W | A | W | A | W | W | A | W | A | A | W | W | A | W | W | A | A | W | W | A | A | W | A | W | W | A | W |
| ChrIV | 0.8 | W | NA | W | A | A | W | A | W | W | W | A | W | A | A | W | A | A | A | A | A | A | W | A | NA | A | A | A | A | W | A | A | A | A | A | A | A | A | A | A | A | A | A | W | W | W | A | W | A | W | NA | A | W | A | A | W | W | W | NA | W | W | W | W | W | A | W | W | A | W | A | W | W | A | W | A | A | W | NA | A | W | W | A | A | W | W | A | A | W | A | NA | W | A | NA |
| ChrIV | 0.9 | W | W | W | A | A | NA | A | W | W | NA | A | W | A | A | W | NA | A | A | A | A | A | W | A | W | A | A | NA | A | W | A | A | A | A | A | A | A | A | A | A | A | A | A | W | W | NA | A | W | A | W | W | A | W | A | A | W | W | W | W | NA | W | W | W | NA | A | NA | W | A | NA | A | W | W | A | NA | A | A | W | W | A | NA | W | A | A | W | W | A | A | W | A | W | W | A | W |
| ChrIV | 1 | W | W | W | NA | A | W | A | W | W | W | A | W | A | A | W | A | A | A | A | A | A | W | A | W | A | A | NA | A | W | A | A | A | A | A | A | A | A | A | A | NA | A | A | W | W | W | NA | W | A | W | W | A | W | A | A | W | W | W | W | W | W | W | NA | A | A | W | W | NA | W | A | W | W | A | W | A | A | W | W | A | W | W | A | A | W | W | A | A | W | A | W | W | A | W |
| ChrIV | 1.1 | W | W | W | A | A | W | A | W | W | W | A | W | A | A | W | A | A | A | A | A | A | W | A | W | A | A | A | A | W | A | A | A | A | A | A | A | A | A | A | A | A | A | W | W | W | A | W | A | W | W | A | W | A | A | W | W | W | W | W | W | W | W | A | A | W | W | NA | W | A | W | W | A | W | A | A | W | W | A | W | NA | A | A | W | W | A | A | W | A | W | W | A | W |
| ChrIV | 1.2 | W | W | W | NA | A | W | NA | W | W | W | A | W | A | A | W | A | A | A | A | A | A | W | A | W | A | A | A | A | W | A | A | A | A | A | A | NA | A | A | A | A | A | A | W | W | W | A | W | A | W | W | A | W | A | A | W | W | W | W | W | W | W | W | A | A | W | W | NA | W | A | W | W | A | W | A | A | W | W | A | W | W | A | A | W | W | A | A | W | A | W | W | A | W |
| ChrIV | 2.4 | W | W | W | NA | A | W | A | W | W | W | A | W | A | NA | W | A | A | A | A | A | A | W | A | W | A | A | NA | NA | W | A | A | A | A | A | A | A | A | A | A | NA | A | A | W | W | W | A | W | A | W | NA | A | W | A | NA | W | W | W | W | W | W | W | W | A | A | W | W | A | W | A | W | W | A | W | A | A | W | W | A | W | W | A | A | W | W | A | A | W | A | W | W | A | W |
| ChrIV | 2.5 | W | W | W | A | A | W | A | W | W | W | A | W | A | A | W | A | A | A | A | NA | A | W | A | W | NA | A | A | A | W | A | A | A | NA | A | A | A | A | A | A | A | A | A | W | W | W | NA | W | A | W | W | NA | W | NA | A | W | W | W | W | W | W | W | W | A | NA | W | W | NA | W | NA | W | W | A | W | A | A | W | W | A | W | W | A | A | W | W | A | A | W | NA | W | W | NA | W |
| ChrIV | 3 | W | W | W | A | A | W | A | W | W | W | A | W | A | A | W | A | A | A | A | A | A | W | A | W | A | A | NA | A | W | A | A | A | A | A | A | A | A | A | A | A | A | A | W | W | W | A | W | A | W | W | A | W | A | A | W | W | W | W | W | W | W | W | A | A | W | W | A | W | A | W | W | A | W | A | A | W | W | A | W | W | A | A | W | W | A | A | NA | A | W | W | A | W |
| ChrIV | 3.6 | W | W | W | A | A | W | A | W | W | W | A | W | A | A | W | A | A | A | A | A | A | W | A | W | A | A | NA | A | W | A | A | A | A | A | A | A | A | A | A | A | A | A | W | W | W | A | W | A | W | W | A | W | A | A | W | W | W | W | W | W | W | W | A | A | W | W | A | W | A | W | W | A | W | A | A | W | W | A | W | W | A | A | W | W | A | A | NA | A | W | W | A | W |
| ChrIV | 3.7 | W | W | W | A | A | W | A | W | W | W | A | W | A | A | W | A | A | A | A | A | A | W | A | W | NA | A | A | A | W | A | A | A | NA | A | A | A | A | A | A | A | A | A | W | W | W | NA | W | A | W | W | A | W | NA | NA | NA | W | W | W | W | W | W | W | A | A | W | W | A | W | A | W | W | A | W | A | A | W | W | NA | W | W | A | A | W | W | A | A | W | A | W | W | A | W |
| ChrIV | 3.9 | W | W | W | NA | NA | W | NA | W | W | W | NA | W | A | NA | W | A | NA | NA | A | NA | A | W | NA | W | A | A | NA | A | W | A | A | A | A | A | A | NA | NA | A | A | NA | A | A | W | W | W | NA | W | NA | W | W | NA | W | NA | A | W | W | W | W | W | W | W | W | NA | A | W | W | NA | W | NA | W | W | NA | W | A | A | W | W | NA | W | W | NA | A | W | W | NA | NA | W | NA | W | W | NA | W |
| ChrIV | 4 | W | W | W | A | A | W | A | W | W | W | A | W | A | A | W | A | NA | A | A | A | A | W | A | W | A | A | NA | A | W | A | A | A | NA | A | A | NA | A | A | A | A | A | A | W | W | W | NA | W | A | W | W | A | W | A | A | W | W | W | W | W | W | W | W | A | A | W | W | A | W | NA | W | W | A | W | A | A | W | W | A | W | W | A | A | W | W | A | A | W | A | W | W | A | W |
| ChrIV | 4.1 | W | W | W | NA | A | W | NA | W | W | W | NA | W | A | NA | W | A | NA | A | A | A | A | W | A | W | NA | A | NA | NA | W | A | A | NA | NA | A | A | NA | NA | A | NA | NA | A | NA | W | W | W | A | W | A | W | W | NA | W | NA | NA | W | W | W | W | W | W | W | W | A | A | W | W | NA | W | NA | W | W | A | W | A | A | W | W | NA | W | W | A | NA | W | W | A | A | W | NA | W | W | NA | W |
| ChrIV | 4.3 | W | W | W | A | A | W | NA | W | W | NA | A | W | A | A | W | A | A | A | A | A | A | W | A | W | A | A | A | A | W | A | A | A | A | A | A | A | A | A | A | A | A | A | W | W | W | A | W | A | W | NA | A | W | A | A | NA | W | W | W | NA | W | W | W | NA | A | W | W | A | W | A | W | W | A | W | A | A | W | W | A | W | W | A | A | W | W | A | A | W | A | W | W | A | W |
| ChrIV | 4.4 | W | W | W | A | A | W | A | NA | W | W | A | W | A | A | W | A | A | A | A | A | A | W | A | W | A | A | A | A | W | A | A | A | A | A | A | A | A | A | A | A | A | A | W | W | W | A | W | A | W | NA | A | W | A | A | W | W | W | W | W | W | W | W | A | A | W | W | A | W | A | W | W | A | W | A | A | W | W | A | W | W | A | A | W | W | A | A | W | A | W | W | A | W |
| ChrIV | 4.6 | W | W | W | NA | A | W | A | W | W | W | A | W | A | A | W | A | A | NA | A | A | A | W | NA | W | NA | A | NA | NA | W | A | A | NA | NA | A | A | A | A | A | A | NA | A | NA | W | W | W | NA | W | A | W | W | NA | W | NA | A | W | W | W | W | W | W | W | W | A | NA | W | W | A | W | A | W | W | A | W | A | A | W | W | NA | W | W | A | A | W | W | A | A | W | A | W | W | NA | W |
| ChrIV | 4.7 | W | W | W | NA | A | W | NA | W | W | W | NA | W | A | A | NA | A | A | NA | A | NA | A | W | NA | W | NA | A | NA | NA | W | A | A | NA | A | A | A | NA | NA | A | A | NA | A | NA | W | W | W | NA | W | A | W | NA | NA | W | NA | NA | W | W | W | W | W | W | W | W | A | A | W | NA | A | W | NA | W | W | A | W | A | A | W | W | NA | W | W | A | NA | W | W | A | A | W | NA | W | W | NA | W |
| ChrIV | 4.8 | W | W | W | A | A | W | A | W | W | W | A | W | A | A | W | W | A | A | A | A | A | W | A | W | A | A | A | A | W | A | A | A | A | A | A | A | A | A | A | A | A | A | W | NA | W | A | W | A | W | W | A | W | A | A | W | W | W | W | W | W | W | W | A | A | W | W | A | W | A | W | W | A | W | A | A | W | W | A | W | W | A | A | W | W | A | A | W | NA | W | W | A | W |
| ChrIV | 5.1 | W | W | W | A | A | W | A | W | W | W | A | W | A | A | NA | W | A | A | A | A | A | W | A | W | NA | A | NA | A | W | A | A | A | A | A | A | A | A | A | A | NA | A | NA | W | W | W | A | W | A | W | NA | A | W | A | A | W | W | W | W | W | W | W | W | A | A | W | W | NA | NA | A | W | W | A | W | A | A | W | W | A | W | W | A | A | W | W | A | A | NA | A | W | W | A | W |
| ChrIV | 5.2 | W | W | W | A | A | W | A | W | W | W | NA | W | A | A | W | W | A | A | A | A | A | W | A | W | A | A | A | A | W | A | A | A | A | A | A | A | A | A | A | A | A | A | W | W | W | A | W | A | W | NA | NA | W | A | A | W | W | W | W | W | W | W | W | NA | A | W | W | NA | W | A | W | W | A | W | A | A | W | W | A | W | W | A | A | W | W | A | A | W | A | W | W | A | W |
| ChrIV | 5.4 | W | W | W | NA | A | W | A | W | W | NA | A | W | A | A | W | W | A | A | A | A | NA | W | A | W | A | A | A | A | W | A | A | A | A | A | A | A | NA | A | A | NA | A | A | W | W | W | NA | W | A | W | W | NA | W | A | A | W | W | W | W | W | W | W | W | A | A | W | W | NA | W | A | W | W | A | W | A | A | W | W | A | W | W | A | A | W | W | A | A | W | A | W | W | A | W |
| ChrIV | 5.7 | W | W | W | NA | A | W | A | W | W | W | A | W | NA | A | W | W | A | A | NA | A | A | W | A | W | A | A | NA | NA | W | A | A | A | A | A | NA | NA | A | A | A | A | NA | A | W | W | W | A | W | A | W | W | A | W | A | A | W | W | W | W | W | W | W | W | NA | A | W | W | A | W | NA | W | W | A | W | NA | NA | W | NA | A | W | W | NA | A | W | W | A | NA | W | A | W | W | A | W |
| ChrIV | 5.9 | W | W | W | NA | A | W | A | W | W | W | NA | W | A | A | W | W | NA | NA | A | A | A | W | A | W | NA | A | A | A | W | A | A | A | NA | A | A | NA | NA | A | A | NA | A | NA | W | W | W | NA | W | A | W | W | NA | W | NA | A | W | W | W | W | W | W | W | W | NA | NA | W | W | NA | W | NA | W | W | A | W | A | A | W | W | NA | W | W | A | A | W | W | A | A | W | A | W | W | NA | W |
| ChrIV | 6.5 | W | W | W | A | A | W | NA | W | W | W | A | W | A | A | W | W | A | A | A | A | NA | W | A | W | NA | A | NA | A | W | A | A | A | NA | A | A | NA | A | A | A | NA | A | A | W | W | W | NA | W | NA | W | W | A | W | A | A | W | W | W | W | W | W | W | W | NA | A | W | W | NA | W | A | W | W | A | W | A | A | W | W | A | W | W | A | NA | W | W | A | A | W | A | W | W | A | W |
| ChrIV | 6.7 | W | W | W | NA | A | W | NA | W | W | W | NA | W | A | A | W | W | NA | A | A | A | NA | W | NA | W | NA | A | A | A | W | A | A | NA | A | A | A | NA | NA | A | NA | NA | A | NA | W | W | W | NA | W | A | W | W | NA | W | A | A | W | W | W | W | W | W | W | W | NA | NA | W | W | NA | W | NA | W | NA | A | W | A | A | W | W | NA | W | W | A | A | W | W | A | A | W | A | W | W | NA | W |
| ChrIV | 6.8 | W | W | W | A | A | W | A | W | W | W | A | W | A | A | W | W | A | A | A | A | A | W | A | W | A | A | A | A | W | A | A | A | A | A | A | A | A | A | A | A | A | A | W | W | W | A | W | A | W | W | A | W | A | A | W | W | W | W | W | W | W | W | A | A | W | W | A | W | A | W | W | A | W | A | A | W | W | A | W | W | A | A | W | W | A | A | W | A | W | W | A | W |
| ChrIV | 6.9 | W | NA | W | A | A | W | NA | W | W | W | A | W | A | A | W | NA | A | A | A | A | A | W | A | W | A | A | A | A | W | NA | A | A | NA | NA | A | NA | A | A | A | A | A | A | W | NA | W | NA | W | A | NA | W | A | W | A | A | W | W | W | NA | NA | W | W | W | A | A | W | W | A | W | NA | W | NA | A | W | A | A | W | NA | A | W | W | A | NA | NA | W | A | A | W | A | W | W | A | W |
| ChrIV | 7 | W | W | W | A | A | W | A | W | W | W | A | W | A | A | W | W | A | A | A | A | A | W | A | W | A | A | A | A | W | A | A | A | NA | A | A | A | A | A | A | NA | A | A | W | W | W | A | W | A | W | W | A | W | A | A | W | W | W | W | W | W | W | W | A | A | W | W | NA | W | A | W | W | A | W | A | A | W | W | A | W | W | A | A | W | W | A | A | W | A | W | W | A | W |
| ChrIV | 7.1 | W | W | W | A | A | W | A | W | W | W | A | W | A | A | W | W | A | A | A | A | A | W | A | W | A | A | A | A | W | A | A | A | A | A | A | A | A | A | A | A | A | A | W | W | W | NA | W | A | W | W | A | W | A | A | W | W | W | W | W | W | W | W | A | A | W | NA | A | W | A | W | W | A | W | A | A | W | W | A | W | W | A | A | W | W | A | A | W | A | W | W | A | W |
| ChrIV | 7.6 | W | W | NA | A | A | W | A | W | W | W | A | W | A | A | W | W | A | A | A | A | A | W | A | W | A | A | A | A | W | A | A | A | A | A | A | A | A | A | A | A | A | A | NA | W | W | A | W | A | NA | W | A | W | A | A | W | W | W | NA | NA | W | W | W | A | A | W | NA | A | W | A | W | W | A | W | A | A | W | NA | A | W | W | A | A | W | W | A | A | W | A | W | W | A | NA |
| ChrIV | 8.3 | W | W | W | NA | A | W | A | W | W | W | NA | W | A | A | W | W | NA | A | A | A | A | W | NA | W | NA | A | NA | NA | W | A | A | A | NA | NA | A | A | A | NA | A | NA | NA | NA | W | W | W | NA | W | NA | W | NA | A | W | NA | A | W | W | W | W | W | W | W | W | NA | NA | W | W | NA | W | NA | W | W | NA | W | A | A | W | W | NA | W | W | A | NA | W | W | A | NA | W | NA | W | W | A | W |
| ChrIV | 8.7 | W | W | W | NA | A | W | A | W | W | W | A | W | A | A | W | W | A | A | A | A | A | W | NA | W | A | A | A | A | W | A | A | A | A | A | A | A | NA | A | A | A | A | A | W | W | W | A | W | A | W | W | A | W | A | A | W | W | W | W | W | W | W | W | A | A | W | W | A | W | A | W | W | A | W | A | A | W | W | A | W | W | A | A | W | W | A | A | W | A | W | W | A | W |
| ChrIV | 8.8 | W | W | W | A | A | W | A | W | W | W | A | W | A | A | W | W | A | A | A | A | A | W | A | W | A | A | A | A | W | A | A | A | A | A | A | A | A | A | A | A | A | A | W | W | W | A | W | A | W | W | A | W | A | A | W | W | W | W | W | W | W | W | A | A | W | W | A | W | A | W | W | A | W | A | A | W | W | A | W | W | A | A | W | W | A | A | W | A | W | W | A | W |
| ChrIV | 8.9 | W | W | W | A | A | W | A | W | W | W | A | W | A | A | W | W | A | A | A | A | A | W | A | W | A | A | A | A | W | A | A | A | A | NA | A | A | A | A | A | A | A | A | W | W | W | A | W | A | W | W | A | W | A | A | W | W | W | W | W | W | W | W | A | A | W | W | A | W | A | W | W | A | W | NA | A | W | W | A | W | W | A | A | W | W | A | A | W | A | W | W | A | W |
| ChrIV | 9 | W | W | W | A | A | W | A | W | W | W | A | W | A | A | W | W | A | A | A | A | A | W | A | W | A | A | A | A | W | A | A | A | A | A | A | A | A | A | A | A | A | A | W | W | W | A | W | A | W | W | A | W | A | A | W | W | W | W | W | W | W | W | A | A | W | W | A | W | A | W | W | A | W | A | A | W | W | A | W | W | A | A | W | W | A | A | W | A | W | W | A | W |
| ChrIV | 9.1 | W | W | W | A | A | W | A | W | W | W | A | W | A | A | W | W | A | A | A | A | A | W | A | W | A | A | A | A | W | A | A | A | A | A | A | A | A | A | A | A | A | A | W | W | W | A | W | A | W | W | A | W | A | A | W | W | W | W | NA | W | W | W | A | A | W | W | A | W | A | W | W | A | W | A | A | W | W | A | W | W | A | A | W | W | A | A | W | A | W | W | A | W |
| ChrIV | 9.6 | W | W | W | A | A | W | NA | W | W | W | A | W | A | A | W | W | A | A | A | A | A | W | NA | W | A | A | NA | A | W | A | A | A | NA | A | A | NA | A | A | A | NA | A | A | W | W | W | NA | W | A | W | W | NA | W | NA | NA | W | W | W | W | W | W | W | W | NA | A | W | W | NA | W | A | W | W | A | W | A | A | W | W | NA | W | W | A | NA | W | W | A | NA | W | A | W | W | NA | W |
| ChrIV | 9.7 | W | W | W | A | A | W | A | W | W | W | A | W | A | A | W | W | A | A | A | A | A | W | A | W | A | A | A | A | W | A | A | A | A | A | A | A | A | A | A | A | A | A | W | W | W | NA | W | A | W | NA | A | W | A | A | W | W | W | W | W | W | W | W | A | A | W | W | A | W | A | W | W | A | W | A | A | W | W | A | W | W | A | A | W | W | A | A | W | A | W | W | A | W |
| ChrIV | 9.8 | W | W | W | A | A | W | NA | W | W | W | A | W | A | A | W | W | A | A | A | A | A | W | A | W | A | A | A | A | W | A | A | A | NA | A | A | A | A | A | A | A | A | A | W | W | W | A | W | A | W | W | A | W | A | A | W | W | W | W | W | W | W | W | A | A | W | W | A | W | A | W | W | A | W | A | A | W | W | A | W | W | A | A | W | W | A | A | W | A | W | W | A | W |
| ChrIV | 9.9 | W | W | W | A | A | W | A | W | W | W | A | W | A | A | W | W | A | A | A | A | A | W | A | W | A | A | A | A | W | A | A | A | A | A | A | A | A | A | A | A | A | A | W | W | W | A | W | A | W | W | A | W | A | A | W | W | W | W | W | W | W | W | A | A | W | W | A | W | A | W | W | A | W | A | A | W | W | A | W | W | A | A | W | W | A | A | W | A | W | W | A | W |
| ChrIV | 10 | W | W | W | A | A | W | A | W | W | W | A | W | A | A | W | W | A | A | A | A | A | W | A | W | A | A | A | A | W | A | A | A | A | A | A | A | A | A | A | A | A | A | W | W | W | A | W | A | W | W | A | W | A | A | W | W | W | W | W | W | W | W | A | A | W | W | A | W | A | W | W | A | W | A | A | W | W | A | W | W | A | A | W | W | A | A | W | A | W | W | A | W |
| ChrIV | 10.1 | W | W | W | NA | A | W | A | W | W | W | NA | W | NA | A | W | W | A | A | A | NA | NA | W | A | W | A | A | NA | NA | W | A | A | A | NA | A | A | NA | A | A | A | A | A | A | W | W | W | NA | W | A | W | W | NA | W | NA | NA | W | W | W | W | W | W | W | W | NA | A | W | W | A | W | NA | W | W | NA | W | NA | NA | W | W | NA | W | W | A | NA | W | W | NA | A | W | A | W | W | NA | W |
| ChrIV | 10.3 | W | W | W | A | A | W | A | W | W | W | A | W | A | A | W | W | A | A | A | A | NA | W | A | W | A | A | NA | A | W | A | A | A | A | NA | A | A | A | A | A | A | A | A | W | W | W | NA | W | A | W | W | A | W | A | A | W | W | W | W | W | W | W | W | A | A | W | W | NA | W | A | W | W | A | W | A | A | W | W | NA | W | W | A | A | W | W | A | A | W | A | W | W | A | W |
| ChrIV | 10.4 | W | W | W | A | A | W | A | W | W | W | A | W | A | A | W | W | A | A | A | A | A | W | A | W | A | A | A | A | W | A | A | A | A | A | A | A | A | A | A | A | A | A | W | W | W | A | W | A | W | W | A | W | A | A | W | W | W | W | W | W | W | W | A | A | W | W | A | W | A | W | W | A | W | A | A | W | W | A | W | W | A | A | W | W | A | A | W | A | W | W | A | W |
| ChrIV | 10.5 | W | W | W | A | A | W | A | W | W | W | A | W | A | A | W | NA | A | A | A | A | A | W | A | W | A | A | A | A | W | A | A | A | A | A | A | A | A | A | A | A | A | A | W | W | W | A | W | A | W | W | A | W | A | A | W | W | W | W | W | W | W | W | A | A | W | W | A | W | A | W | W | A | W | A | A | W | W | A | W | W | A | A | W | W | A | A | W | A | W | W | A | W |
| ChrIV | 10.6 | W | W | W | A | A | W | A | W | W | W | A | W | A | A | W | W | A | A | A | A | A | W | A | W | A | A | A | A | W | A | A | A | A | A | A | A | A | A | A | A | A | A | W | W | W | A | W | A | W | NA | A | W | A | A | W | W | W | W | W | W | W | W | A | A | W | W | A | W | A | W | W | A | W | A | A | W | W | A | W | W | A | A | W | W | A | A | W | A | W | W | A | W |
| ChrIV | 10.7 | W | W | W | A | A | W | A | W | W | W | A | W | A | A | W | W | A | A | A | A | A | W | A | W | A | A | A | A | W | A | A | A | NA | A | A | A | A | A | A | A | A | A | W | W | W | A | W | A | W | W | A | W | A | A | W | W | W | W | W | W | W | W | A | A | W | W | A | W | NA | W | W | A | W | A | A | W | W | A | W | W | A | A | W | W | A | A | W | A | W | W | A | W |
| ChrIV | 10.8 | W | W | W | A | A | W | A | W | W | W | A | W | A | A | W | W | A | A | A | A | A | W | A | W | A | A | A | A | W | A | A | A | A | A | A | A | A | A | A | A | A | A | W | W | W | A | W | A | W | W | A | W | A | A | W | W | W | W | W | W | W | W | A | A | W | W | A | W | A | W | W | A | W | A | A | W | W | A | W | W | A | A | W | W | A | A | W | A | W | W | A | W |
| ChrIV | 11.1 | W | W | W | A | A | W | A | W | W | W | A | W | A | A | W | W | A | A | A | A | A | W | A | W | A | A | A | A | W | A | A | A | A | A | A | A | A | A | A | A | A | A | W | W | W | A | W | A | W | W | A | W | A | A | W | W | W | W | W | W | W | W | A | A | W | W | A | W | A | W | W | A | W | A | A | W | W | A | W | W | A | A | W | W | A | A | W | A | W | W | A | W |
| ChrIV | 11.2 | W | W | W | A | A | W | A | W | W | W | A | W | A | A | W | W | A | A | A | A | A | W | A | W | A | A | A | A | W | A | A | A | A | A | A | A | A | A | A | A | A | A | W | W | W | NA | W | A | W | W | A | W | A | A | W | W | W | W | W | W | W | W | A | A | W | W | A | W | A | W | W | A | W | A | A | W | W | A | W | W | A | A | W | W | A | A | W | A | W | W | A | W |
| ChrIV | 11.3 | W | W | W | A | A | W | A | W | W | W | A | W | A | A | W | W | NA | A | A | A | A | W | A | W | NA | A | A | A | W | A | A | A | A | A | A | A | A | A | A | A | A | A | W | W | W | A | W | A | W | W | A | W | A | A | W | W | W | W | W | W | W | W | A | A | W | W | A | W | A | W | W | A | W | A | A | W | W | A | W | W | A | A | W | W | A | A | W | A | W | W | A | W |
| ChrIV | 11.4 | W | W | W | A | A | W | A | W | W | W | A | W | A | A | W | W | A | A | A | A | A | W | A | W | A | A | A | A | W | A | A | A | A | A | A | A | A | A | A | A | A | A | W | W | W | A | W | A | W | W | A | W | A | A | W | W | W | W | W | W | W | W | A | A | W | W | A | W | A | W | W | A | W | A | A | W | W | A | W | W | A | A | W | W | A | A | W | A | W | W | A | W |
| ChrIV | 11.5 | W | W | W | A | A | W | A | NA | W | W | A | W | A | A | W | NA | A | A | A | A | A | W | A | W | A | A | A | A | W | A | A | A | A | A | A | A | A | A | A | A | A | A | W | W | W | A | W | A | NA | W | A | W | A | A | W | W | W | W | W | W | W | W | A | A | W | NA | A | W | A | W | NA | A | W | A | A | W | W | A | NA | W | A | A | W | W | A | A | W | A | W | W | A | W |
| ChrIV | 11.6 | W | NA | NA | A | A | W | A | W | W | W | A | W | A | A | W | NA | A | A | A | NA | NA | W | A | NA | A | A | A | A | NA | A | A | A | A | A | A | A | A | A | A | A | A | A | NA | NA | NA | A | W | A | W | W | A | W | A | A | NA | W | W | NA | W | W | W | W | A | A | W | W | A | W | A | W | W | A | W | A | A | W | W | A | NA | NA | A | A | W | W | NA | A | W | A | W | W | A | W |
| ChrIV | 11.7 | W | W | W | A | A | W | A | W | W | W | A | W | A | A | W | W | A | A | A | A | A | W | A | W | A | A | A | A | W | A | A | A | A | A | A | A | A | A | A | A | A | A | W | W | W | A | W | A | W | W | A | W | A | A | W | W | W | W | W | W | W | W | A | A | W | W | A | W | A | W | W | A | W | A | A | W | W | A | W | W | A | A | W | W | A | A | W | A | W | W | A | W |
| ChrIV | 11.8 | W | W | W | A | A | W | A | W | W | W | A | W | A | A | W | W | A | A | A | A | A | W | A | W | A | A | A | A | W | A | A | A | A | A | A | A | A | A | A | A | A | A | W | W | W | A | W | A | W | W | A | W | A | A | W | W | W | W | W | W | W | W | A | A | W | W | A | W | A | W | W | A | W | A | A | W | W | A | W | W | A | A | W | W | A | A | W | A | W | W | A | W |
| ChrIV | 11.9 | W | W | W | A | A | W | A | W | W | W | A | W | A | A | NA | W | A | A | A | A | A | W | NA | W | A | A | A | A | W | A | A | A | A | A | A | A | A | A | A | A | A | A | W | W | W | A | W | A | W | W | A | W | A | A | W | W | W | W | W | W | W | W | A | A | W | W | A | W | A | W | W | A | W | A | A | W | W | A | W | W | A | A | W | W | A | A | W | A | W | W | A | W |
| ChrIV | 12.1 | W | W | W | A | A | W | A | W | W | W | A | W | A | A | W | NA | A | A | A | A | A | W | A | W | A | A | A | A | W | A | A | A | A | A | A | A | A | A | A | A | A | A | W | W | W | A | W | A | W | W | A | W | A | A | W | W | W | W | W | W | W | W | A | A | W | W | A | W | A | W | W | A | W | A | A | W | W | A | W | W | A | A | W | W | A | A | W | A | W | W | A | W |
| ChrIV | 12.3 | W | W | W | A | A | W | A | W | W | W | A | W | A | A | W | W | A | A | A | A | A | W | A | W | A | A | A | A | W | A | A | A | A | A | A | A | A | A | A | A | A | A | W | W | W | A | W | A | W | W | A | W | A | A | W | W | W | W | W | W | W | W | A | A | W | W | A | W | A | W | W | A | W | A | A | W | W | A | W | W | A | A | W | W | A | A | W | A | W | W | A | W |
| ChrIV | 12.4 | W | W | W | A | A | W | A | W | W | W | A | W | A | A | W | W | A | A | A | A | A | W | A | W | NA | A | A | A | W | A | A | A | A | A | A | A | A | A | A | A | A | A | W | W | W | A | W | A | W | W | A | W | A | A | W | W | W | W | W | W | W | W | A | A | W | W | A | W | A | W | W | A | W | A | A | W | W | A | W | W | A | A | W | W | A | A | W | A | W | W | A | W |
| ChrIV | 12.5 | W | W | W | A | A | W | A | W | W | W | A | W | A | A | W | W | A | A | A | A | A | W | A | W | A | A | A | A | W | A | A | A | A | A | A | A | A | A | A | A | A | A | W | W | W | A | W | A | W | W | A | W | A | A | W | W | W | W | W | W | W | W | A | A | W | W | NA | W | A | W | W | A | W | A | A | W | W | A | W | W | A | A | W | W | A | A | W | A | W | W | A | W |
| ChrIV | 12.6 | W | W | W | A | A | W | A | NA | W | W | A | W | A | A | W | W | A | A | A | NA | NA | W | A | W | A | A | A | A | W | A | A | A | A | A | A | A | A | A | A | A | A | A | W | W | W | A | W | A | W | W | A | W | A | A | W | W | W | W | W | W | W | W | A | A | W | W | A | W | A | W | W | A | W | A | A | W | W | A | W | W | A | A | W | W | A | A | W | A | W | W | A | W |
| ChrIV | 12.7 | W | W | W | A | A | W | A | W | W | W | A | W | A | A | W | W | A | A | A | A | A | W | A | W | A | A | NA | A | W | A | A | A | A | A | A | A | A | A | A | A | A | A | W | NA | W | A | W | A | W | W | A | W | A | A | W | W | W | W | W | W | W | W | A | A | W | NA | A | W | A | W | W | A | W | A | A | W | NA | A | W | W | A | A | W | W | A | A | W | A | W | W | A | W |
| ChrIV | 12.8 | W | W | W | A | A | W | A | W | W | W | A | W | A | A | W | W | A | A | A | A | A | W | A | W | A | A | A | A | W | A | A | A | A | A | A | A | A | A | A | A | A | A | W | W | W | A | W | A | W | W | A | W | A | A | W | W | W | W | W | W | W | W | A | A | W | W | A | W | A | W | W | A | W | A | A | W | W | A | W | W | A | A | W | W | A | A | W | A | W | W | A | W |
| ChrIV | 12.9 | W | W | W | NA | NA | W | A | W | W | W | A | W | A | A | W | W | A | A | A | A | A | W | NA | W | A | A | NA | A | W | A | A | A | NA | A | A | A | A | A | A | NA | A | A | W | W | W | A | W | A | W | W | A | W | A | A | W | W | W | W | W | W | W | W | A | A | W | W | NA | W | NA | W | W | NA | W | A | A | W | W | NA | W | W | A | A | W | W | NA | NA | W | A | W | W | A | W |
| ChrIV | 13 | W | W | W | A | A | W | A | W | W | W | A | W | A | A | W | W | A | A | A | A | A | W | A | W | A | A | NA | A | W | A | A | A | A | A | A | A | A | A | A | A | A | A | W | W | W | NA | W | A | W | W | A | W | A | A | W | W | W | W | W | W | W | W | A | A | W | W | A | W | A | W | W | A | W | A | A | W | W | A | W | W | A | A | W | W | A | A | W | A | W | W | A | W |
| ChrIV | 13.1 | W | W | W | A | A | W | A | W | W | W | A | W | A | A | W | W | A | A | A | A | A | W | A | W | A | A | A | A | W | A | A | A | A | A | A | A | A | A | A | A | A | A | W | W | W | A | W | A | W | W | A | W | A | A | W | W | W | W | W | W | W | W | A | A | W | W | A | W | A | W | W | A | W | A | A | W | W | A | W | W | A | A | W | W | A | A | W | A | W | W | A | W |
| ChrIV | 13.2 | W | W | W | A | A | W | NA | W | W | W | A | W | A | A | W | W | A | A | A | A | A | W | NA | W | A | A | NA | A | W | A | A | A | A | A | A | A | A | A | A | A | A | A | W | W | W | A | W | A | W | W | A | NA | A | A | W | W | W | W | W | W | W | W | A | A | W | W | A | W | A | W | W | A | W | A | A | W | W | A | W | W | A | A | W | W | A | A | W | A | NA | W | A | W |
| ChrIV | 13.3 | W | W | W | A | A | W | A | W | W | W | A | W | A | A | W | W | A | A | A | A | A | W | A | W | A | A | A | A | W | A | A | A | A | A | A | A | A | A | A | A | A | A | W | W | W | A | W | A | W | W | A | A | A | A | W | W | W | W | W | W | W | W | A | A | W | W | A | W | A | W | W | A | W | A | A | W | W | A | W | W | A | A | W | W | A | A | W | A | W | W | A | W |
| ChrIV | 13.4 | W | W | W | A | A | W | A | W | W | W | A | W | A | A | W | NA | A | A | A | A | A | W | A | W | A | A | A | A | W | A | A | A | A | A | A | A | A | A | A | A | A | A | W | W | W | A | W | A | W | W | A | NA | A | A | W | W | W | W | W | W | W | W | A | A | W | W | A | W | A | W | W | A | W | A | A | W | W | A | W | W | A | A | W | W | A | A | W | A | W | W | A | W |
| ChrIV | 13.5 | W | W | W | A | A | W | A | W | W | W | A | W | A | A | W | W | A | A | A | A | A | W | A | W | A | A | A | A | W | A | A | A | A | A | A | A | A | A | A | A | A | A | W | W | W | A | W | A | W | W | A | A | A | A | W | W | W | W | W | W | W | W | A | A | W | W | NA | W | A | W | W | A | W | A | A | W | W | A | W | W | A | A | W | W | A | A | W | A | W | W | A | W |
| ChrIV | 13.6 | W | W | W | A | A | W | A | W | W | W | A | W | A | A | W | W | A | A | A | A | A | W | A | W | A | A | A | A | W | A | A | A | A | A | A | A | A | A | A | NA | A | A | W | W | W | A | W | A | W | W | A | A | A | A | W | W | W | W | W | W | W | W | A | A | W | W | A | W | A | W | W | A | W | A | A | W | W | A | W | W | A | A | W | W | A | A | W | A | W | W | A | W |
| ChrIV | 13.7 | W | W | W | A | A | W | A | W | W | W | A | W | A | A | W | W | A | A | A | A | A | W | A | W | A | A | A | A | W | A | A | A | A | A | A | A | A | A | A | A | A | A | W | W | W | A | W | A | W | W | A | A | A | A | W | W | W | W | W | W | W | W | A | A | W | W | A | W | A | W | W | A | W | A | A | W | W | A | W | W | A | A | W | W | A | A | W | A | W | W | A | W |
| ChrIV | 13.8 | W | W | W | A | A | W | A | W | W | W | A | W | A | A | W | W | A | A | A | A | A | W | A | W | NA | A | A | A | W | A | A | A | A | A | A | A | A | A | A | A | A | A | W | W | W | A | W | A | W | NA | A | A | A | A | W | W | W | W | W | W | W | W | A | A | W | W | A | W | A | W | W | A | W | A | A | W | W | A | W | W | A | A | W | W | A | A | W | A | W | W | A | W |
| ChrIV | 13.9 | W | W | W | A | A | W | A | W | W | W | A | W | A | A | W | W | A | A | A | A | A | W | A | W | A | A | A | A | W | A | A | A | A | A | A | A | A | A | A | A | A | A | W | W | W | A | W | A | W | W | A | A | A | A | W | W | W | W | W | W | W | W | A | A | W | W | A | W | A | W | W | A | W | W | A | W | W | A | W | W | A | A | W | W | A | A | W | A | W | W | A | W |
| ChrIV | 14 | W | W | W | A | A | W | A | W | W | W | A | W | A | A | W | W | A | A | A | A | A | W | A | W | A | A | A | A | W | A | A | A | A | A | A | A | A | A | A | A | A | A | W | W | W | A | W | A | W | W | A | A | A | A | W | W | W | W | W | W | W | W | A | A | W | W | A | W | A | W | W | A | W | NA | A | W | W | A | W | W | A | A | W | W | A | A | W | A | W | W | A | W |
| ChrIV | 14.1 | W | W | W | A | A | W | A | W | W | W | A | W | A | NA | W | NA | A | A | A | A | A | W | A | NA | A | A | A | A | W | A | A | A | A | A | A | A | A | A | A | A | A | A | W | W | NA | A | W | A | W | W | A | A | A | A | W | W | W | W | W | W | W | W | A | A | W | W | A | W | A | W | W | A | W | NA | A | W | NA | A | NA | NA | A | A | W | W | NA | A | W | A | W | W | A | W |

| | | | | | | | | | | | | | | | | | | | | | | | | | | | | | | | | | | | | | | | | | | | | | | | | | | | | | | | | | | | | | | | | | | | | | | | | | | | | | | | | | | | | | | | | | | | | | |
|---|---|---|---|---|---|---|---|---|---|---|---|---|---|---|---|---|---|---|---|---|---|---|---|---|---|---|---|---|---|---|---|---|---|---|---|---|---|---|---|---|---|---|---|---|---|---|---|---|---|---|---|---|---|---|---|---|---|---|---|---|---|---|---|---|---|---|---|---|---|---|---|---|---|---|---|---|---|---|---|---|---|---|---|---|---|---|---|---|---|---|---|---|---|
| ChrIV | 14.2 | W | W | W | A | A | W | A | W | W | W | A | W | A | A | W | W | A | A | A | A | A | W | A | W | A | A | A | A | W | A | A | A | A | A | A | A | A | A | A | A | A | A | W | W | W | A | W | A | W | W | A | A | A | A | W | W | W | W | W | W | W | W | A | A | W | W | A | W | A | W | W | A | W | W | A | W | W | A | W | W | A | A | W | W | A | A | W | A | W | W | A | W |
| ChrIV | 14.3 | W | W | W | A | A | W | A | W | W | W | A | W | A | A | W | W | A | A | A | A | A | W | A | W | A | A | A | A | W | A | A | A | A | A | A | A | A | A | A | A | A | A | W | W | W | A | W | A | W | W | A | A | A | A | W | W | W | W | W | W | W | W | A | A | W | W | A | W | A | W | NA | A | W | W | A | W | W | A | W | W | A | A | W | W | A | A | W | A | W | W | A | W |
| ChrIV | 14.4 | W | W | W | A | A | W | A | W | W | W | A | W | A | A | W | W | NA | A | A | A | A | W | A | W | A | A | NA | A | W | A | A | A | A | A | A | A | A | A | A | A | A | A | W | W | W | A | W | A | W | W | A | A | A | A | W | W | W | W | W | W | W | W | A | A | W | W | A | W | A | W | W | A | W | W | A | W | W | A | W | W | A | A | W | W | A | A | W | A | W | W | A | W |
| ChrIV | 14.5 | W | W | W | A | A | W | A | NA | W | W | A | W | A | A | W | W | A | A | A | A | A | W | A | W | A | A | A | A | W | A | A | A | NA | A | A | A | A | A | A | A | A | A | W | W | W | A | W | A | W | W | A | A | A | A | W | W | W | W | W | W | W | W | A | A | W | W | A | W | A | W | W | A | W | W | A | W | W | A | W | W | A | A | W | W | A | A | W | A | W | W | A | W |
| ChrIV | 14.6 | W | W | W | A | A | W | A | W | W | W | A | W | A | A | W | W | A | A | A | A | A | W | A | W | A | A | A | A | W | A | A | A | A | A | A | A | A | A | A | A | A | A | W | W | W | A | W | A | W | W | A | A | A | A | W | W | W | W | W | W | W | W | A | A | W | W | A | W | A | W | W | A | W | W | A | W | NA | A | W | W | A | A | W | W | A | A | W | A | W | W | A | W |
| ChrIV | 14.7 | W | NA | NA | A | A | W | A | W | W | W | A | W | A | A | W | NA | A | A | A | A | NA | W | A | NA | A | A | NA | A | W | A | NA | A | A | A | A | A | A | A | A | A | A | A | NA | NA | W | NA | NA | A | NA | NA | A | A | A | A | W | W | W | W | W | W | W | W | A | A | W | W | A | W | A | W | NA | A | W | NA | A | W | W | A | NA | W | A | NA | W | W | A | A | W | A | W | W | A | NA |
| ChrIV | 14.8 | W | W | W | A | A | W | A | W | W | W | A | W | A | A | W | W | A | A | A | A | A | W | A | W | A | A | A | A | W | A | A | A | A | A | A | A | A | A | A | A | A | A | W | W | W | A | W | A | W | W | A | A | A | A | W | W | W | W | W | W | W | W | A | A | W | NA | A | W | A | W | W | A | W | W | A | W | W | A | W | W | A | A | W | W | A | A | W | A | W | W | A | W |
| ChrIV | 14.9 | W | W | W | A | A | W | A | W | W | W | A | W | A | A | W | W | A | A | A | NA | A | W | A | W | A | A | A | A | W | A | A | A | A | A | A | A | A | A | A | A | A | A | W | W | W | NA | W | A | W | W | A | A | A | A | W | W | W | W | W | W | W | W | A | A | W | W | NA | W | A | W | W | A | W | W | A | W | W | A | W | W | A | A | W | W | A | A | W | A | W | W | A | W |
| ChrIV | 15 | W | W | W | A | A | W | A | W | W | W | A | W | A | A | W | W | A | A | A | A | A | W | A | W | A | A | A | A | W | A | A | A | A | A | A | A | A | A | A | A | A | A | W | W | W | A | W | A | W | W | A | A | A | A | W | W | W | W | W | W | W | W | A | A | W | W | A | W | A | W | W | A | W | NA | A | W | W | A | W | W | A | A | W | W | A | A | W | A | W | W | A | W |
| ChrIV | 15.1 | W | W | W | A | A | W | A | W | W | W | A | W | A | A | W | W | A | A | A | A | A | W | A | W | A | A | A | A | W | A | A | A | A | A | A | A | A | A | A | A | A | A | W | W | W | A | W | A | W | W | A | A | A | A | W | W | W | W | W | W | W | W | A | A | W | W | A | W | A | W | W | A | W | W | A | W | W | A | W | W | A | A | W | W | A | A | W | A | W | W | A | W |
| ChrIV | 15.2 | W | W | W | A | A | W | A | W | W | W | A | W | A | A | W | W | A | A | A | A | A | W | A | W | A | A | A | A | W | A | A | A | A | A | A | A | A | A | A | A | A | A | W | W | W | A | W | A | W | W | A | A | A | A | W | W | W | W | W | W | W | W | A | A | W | W | A | W | A | W | W | A | W | W | A | W | W | A | W | W | A | A | W | W | A | A | W | A | W | W | A | NA |
| ChrIV | 15.3 | W | W | W | A | A | W | A | W | W | W | A | W | A | A | W | W | A | A | A | A | A | W | A | W | A | A | A | A | W | A | A | A | A | A | A | A | A | A | A | A | A | A | W | W | W | A | W | A | W | W | A | A | A | A | W | W | W | W | W | W | W | W | A | A | W | W | NA | W | A | W | W | A | W | W | A | W | W | A | W | W | A | A | W | W | A | A | W | A | W | W | A | W |
| ChrIV | 15.4 | W | W | W | A | A | W | A | W | W | W | A | W | A | A | W | W | A | A | A | A | A | W | A | W | A | A | A | A | W | A | A | A | A | A | A | A | A | A | A | A | A | A | W | W | W | A | W | A | W | W | A | A | A | A | W | W | W | W | W | W | W | W | A | A | W | W | A | W | A | W | W | A | W | W | A | W | W | A | W | W | A | A | W | W | A | A | W | A | W | W | A | W |
| ChrIV | 15.5 | W | W | W | A | A | W | A | W | W | W | A | W | A | A | W | NA | A | A | A | A | A | W | A | W | A | A | A | A | W | A | A | A | A | A | A | A | A | A | A | A | A | A | W | W | W | A | W | A | W | W | A | A | A | A | W | W | W | W | W | W | W | W | A | A | W | W | A | W | A | W | W | A | W | W | A | W | W | A | W | W | A | A | W | W | A | A | W | A | W | W | A | W |
| ChrIV | 15.6 | W | W | W | A | A | W | A | W | W | W | A | W | A | A | W | W | A | A | A | A | A | W | A | W | A | A | A | A | W | A | A | A | A | A | A | A | A | A | A | A | A | A | W | W | W | A | W | A | W | W | A | A | A | A | W | W | W | W | W | W | W | W | A | A | W | W | A | W | A | W | W | A | W | W | A | W | W | A | W | W | A | A | W | W | A | A | W | A | W | W | A | W |
| ChrIV | 15.7 | W | W | W | A | A | W | A | W | W | W | A | W | A | A | W | W | A | A | A | A | A | W | A | W | A | A | A | A | W | A | A | A | A | A | A | A | A | A | A | A | A | A | W | W | W | A | W | A | W | W | A | A | A | A | NA | W | W | W | W | W | W | W | A | A | W | W | A | W | A | W | W | A | W | W | A | W | W | A | W | W | A | A | W | W | NA | A | W | A | W | W | A | W |
| ChrIV | 15.8 | W | W | W | A | A | W | A | W | W | W | A | W | A | A | W | W | A | A | A | A | A | W | A | W | A | A | A | A | W | A | A | A | A | A | A | A | A | A | A | A | A | A | W | W | W | A | W | A | W | W | A | A | A | A | W | W | W | W | W | W | W | W | A | A | W | W | A | W | A | W | W | A | W | W | A | W | W | A | W | W | A | A | W | W | A | A | W | A | W | W | A | W |
| ChrIV | 15.9 | W | W | W | A | A | W | A | W | W | W | A | W | A | A | W | W | A | A | A | A | A | W | A | W | A | A | A | A | W | A | A | A | NA | A | A | A | A | A | A | A | A | A | W | W | W | A | W | A | W | W | A | A | A | A | W | W | W | W | W | W | W | W | A | A | W | W | A | W | A | W | W | A | W | W | A | W | W | A | W | W | A | A | W | W | A | A | W | A | W | W | A | W |
| ChrIV | 16 | W | W | W | A | A | W | A | W | W | W | A | W | A | A | W | W | A | A | A | NA | A | W | A | W | A | A | A | A | W | A | A | A | A | A | A | A | A | A | A | A | A | A | W | W | W | A | W | A | W | W | A | A | A | A | W | W | W | NA | W | W | W | W | A | A | W | NA | A | W | A | W | NA | A | W | W | A | W | W | A | W | NA | A | A | NA | W | A | A | W | A | W | W | A | NA |
| ChrIV | 16.1 | W | W | W | A | A | W | A | W | W | W | A | W | A | A | W | NA | A | A | A | A | NA | W | A | W | A | A | A | A | W | A | A | A | A | A | A | A | A | A | A | A | A | A | W | W | W | A | W | A | W | W | A | A | A | A | W | W | W | W | W | W | W | W | A | A | W | NA | A | W | A | W | NA | A | W | NA | A | W | NA | A | W | W | A | A | W | W | A | A | W | A | W | W | A | W |
| ChrIV | 16.2 | W | W | W | A | A | W | A | W | W | W | A | W | A | A | W | W | A | A | A | A | A | W | A | W | A | A | A | A | W | A | A | A | A | A | A | A | A | A | A | A | A | A | W | W | W | A | W | A | W | W | A | A | A | A | W | W | W | W | W | W | W | W | A | A | W | W | A | W | A | W | W | A | W | W | A | W | W | A | W | W | A | A | W | W | A | A | W | A | W | W | A | W |
| ChrIV | 16.3 | W | W | W | A | A | W | A | W | W | W | A | W | A | A | W | W | A | A | A | A | A | W | A | W | A | A | NA | A | W | A | A | A | A | A | A | A | A | A | A | A | A | A | W | W | W | A | W | A | W | W | A | A | A | A | W | W | W | W | W | W | W | W | A | A | W | W | A | W | A | W | W | A | W | W | A | W | W | A | W | W | A | A | W | W | A | A | W | A | W | W | A | W |
| ChrIV | 16.4 | W | W | W | A | A | W | A | W | W | W | A | W | A | A | W | W | A | A | A | A | A | W | A | W | A | A | A | NA | W | A | A | A | A | A | A | A | A | A | A | A | A | A | W | W | W | A | W | A | W | W | A | A | A | A | W | W | W | W | W | W | W | W | A | A | W | W | A | W | A | W | W | A | W | W | A | W | W | A | W | W | A | A | W | W | A | A | W | A | W | W | A | W |
| ChrIV | 16.5 | W | W | W | A | A | W | A | W | W | W | A | W | A | A | W | W | A | A | A | A | A | W | A | W | A | A | A | A | W | A | A | A | A | A | A | A | A | A | A | A | A | A | W | W | W | A | W | A | W | W | A | A | A | A | W | W | W | W | W | W | W | W | A | A | W | W | A | W | A | W | W | A | W | W | A | W | NA | A | W | W | A | A | W | W | A | A | W | A | W | W | A | NA |
| ChrIV | 16.6 | W | NA | NA | A | A | W | A | W | W | W | A | W | NA | A | W | NA | A | NA | A | A | A | W | A | NA | A | A | A | NA | NA | A | A | A | NA | NA | A | A | A | A | A | A | A | A | NA | NA | NA | A | NA | A | W | W | A | A | A | A | NA | W | W | NA | W | NA | W | W | A | A | W | W | A | W | A | W | NA | NA | W | NA | A | W | W | A | W | NA | NA | A | W | W | A | A | W | A | W | W | A | W |
| ChrIV | 16.7 | W | W | W | A | A | W | A | W | W | W | A | W | A | A | W | W | A | A | A | A | NA | W | A | W | A | A | NA | A | W | A | A | A | NA | A | A | A | A | A | A | NA | A | A | W | W | W | A | W | A | W | W | A | NA | A | NA | W | W | W | W | W | W | W | W | NA | A | W | W | NA | W | NA | W | W | A | W | W | A | W | W | A | W | W | A | NA | W | W | A | A | W | A | W | W | A | W |
| ChrIV | 16.8 | W | W | W | A | A | W | A | W | W | W | A | W | A | A | W | W | A | A | A | A | A | W | A | W | A | A | A | A | W | A | A | A | A | A | A | A | A | A | A | A | A | A | W | W | W | A | W | A | W | W | A | A | A | A | W | W | W | W | W | W | W | W | A | A | W | W | A | W | A | W | W | A | W | W | A | W | NA | A | W | W | A | A | W | W | A | NA | W | A | W | W | A | W |
| ChrIV | 16.9 | W | W | W | A | A | W | A | W | W | W | A | W | A | A | W | W | A | A | A | A | A | W | A | W | A | A | A | A | W | A | A | A | A | A | A | A | A | A | A | A | A | A | W | W | W | A | W | A | W | W | A | A | A | A | W | W | W | W | W | W | W | W | A | A | W | W | A | W | A | W | W | A | W | W | A | W | W | A | W | W | A | A | W | W | A | A | W | A | W | W | A | W |
| ChrIV | 17 | W | W | W | A | A | W | A | W | W | W | A | W | NA | A | NA | NA | A | A | A | A | A | W | A | W | A | A | A | A | W | A | A | A | A | NA | A | A | A | A | A | A | A | A | W | W | W | A | W | A | W | W | A | A | A | A | W | W | W | W | W | W | W | W | A | A | W | W | A | W | A | W | W | A | W | W | A | W | W | A | W | W | A | A | W | W | A | A | W | A | W | W | A | W |
| ChrIV | 17.1 | W | W | W | NA | A | W | NA | W | W | W | A | W | A | A | W | W | A | A | A | NA | A | W | NA | W | NA | NA | NA | A | W | NA | A | NA | NA | A | A | NA | A | A | A | NA | A | A | W | W | W | A | W | A | W | NA | NA | NA | A | NA | W | W | W | W | W | W | W | W | NA | NA | W | W | NA | W | NA | W | W | A | W | W | A | W | W | NA | W | W | NA | NA | W | NA | A | A | NA | NA | W | W | NA | W |
| ChrIV | 17.2 | W | W | W | A | A | W | A | W | W | W | A | W | A | A | W | W | A | A | A | A | A | W | A | W | A | A | A | A | W | A | A | A | A | A | A | A | A | A | A | A | A | A | W | W | W | A | W | A | W | W | A | A | A | A | W | W | W | W | W | W | W | W | A | A | W | W | A | W | A | W | W | NA | W | W | A | W | NA | A | W | W | A | A | W | W | A | A | W | A | W | W | A | W |
| ChrIV | 17.3 | W | NA | W | A | A | W | A | W | W | W | A | W | A | A | W | NA | A | A | NA | A | A | W | A | NA | A | NA | A | A | W | A | A | A | A | A | A | A | A | A | A | A | A | A | NA | NA | NA | A | W | A | W | W | A | A | A | A | NA | W | W | NA | W | W | W | W | A | A | W | NA | A | W | A | W | NA | A | W | NA | A | W | NA | A | W | NA | A | A | W | W | A | A | W | NA | W | NA | A | W |
| ChrIV | 17.4 | W | W | NA | A | A | W | A | NA | W | W | A | W | NA | A | W | NA | A | A | NA | A | A | W | A | NA | A | A | A | A | W | A | A | A | A | A | NA | A | A | A | A | A | NA | A | W | NA | W | A | W | A | NA | W | A | A | A | A | NA | W | W | NA | W | NA | W | W | A | A | W | NA | A | W | A | W | NA | A | W | NA | NA | W | NA | A | NA | NA | A | A | NA | W | A | NA | W | A | W | NA | A | NA |
| ChrIV | 17.5 | W | NA | NA | A | A | W | A | W | W | W | A | W | A | A | W | W | A | A | A | A | A | W | A | NA | A | A | A | NA | W | A | A | A | A | A | A | A | A | A | A | A | A | A | W | NA | NA | A | NA | A | W | W | A | A | A | A | NA | W | W | NA | W | NA | W | W | A | A | W | W | A | W | A | W | W | A | W | NA | A | W | NA | A | NA | NA | A | A | NA | W | A | A | W | NA | W | NA | A | NA |
| ChrIV | 17.6 | W | W | W | A | A | W | A | NA | W | W | A | W | A | A | W | W | A | A | A | A | A | W | A | NA | A | A | A | A | W | A | A | A | A | NA | A | A | A | A | A | A | A | A | NA | NA | NA | A | NA | A | NA | W | A | A | A | A | W | W | W | NA | W | W | W | W | A | A | W | NA | A | W | A | W | NA | A | W | W | A | W | NA | A | W | NA | A | A | W | W | A | A | W | A | W | NA | A | W |
| ChrIV | 17.7 | W | W | W | A | A | W | A | W | W | W | A | W | A | A | W | NA | A | A | A | A | A | W | A | W | A | A | A | A | W | A | A | A | A | A | A | A | A | A | A | A | A | A | W | W | W | A | W | A | W | W | A | A | A | A | NA | W | W | W | W | NA | W | W | A | A | W | W | A | W | A | W | W | A | W | W | A | W | W | A | W | W | A | A | W | W | A | A | W | A | W | W | A | W |
| ChrIV | 17.8 | W | NA | NA | A | A | W | A | W | W | W | A | W | A | A | W | NA | A | A | A | A | A | W | A | NA | A | NA | A | A | W | A | A | A | A | A | A | A | A | A | A | A | A | A | NA | W | W | A | NA | A | NA | W | A | A | A | A | NA | W | W | NA | W | NA | W | W | A | A | W | NA | A | W | A | W | W | A | W | NA | A | W | NA | A | NA | NA | A | A | NA | W | A | A | W | A | W | W | A | NA |
| ChrIV | 17.9 | W | NA | NA | A | A | W | A | NA | W | W | A | W | A | A | W | W | A | NA | A | A | NA | W | A | NA | A | A | A | A | W | A | A | A | A | A | A | A | A | A | A | A | NA | A | NA | NA | W | A | NA | A | NA | W | A | A | A | A | W | W | W | W | W | W | W | W | A | A | W | W | A | W | A | W | NA | A | W | NA | A | W | NA | A | NA | NA | A | A | W | W | A | A | W | A | W | W | A | NA |
| ChrIV | 18 | W | NA | NA | A | A | W | A | NA | W | W | A | W | A | A | W | NA | A | A | A | NA | A | W | A | NA | A | A | A | A | W | A | A | A | A | A | A | A | A | A | A | A | A | A | W | W | NA | A | W | A | W | W | A | A | A | A | NA | W | W | NA | W | NA | W | W | A | A | W | W | A | W | A | W | W | A | W | NA | A | W | NA | A | W | NA | A | A | NA | W | A | A | W | A | W | W | A | NA |
| ChrIV | 18.2 | W | W | W | A | A | W | A | W | W | W | A | W | A | A | W | NA | A | A | A | A | NA | W | A | W | A | A | A | A | W | A | A | A | A | A | A | A | A | A | A | A | A | A | W | W | NA | A | W | A | NA | W | A | A | A | A | W | W | W | W | W | W | W | W | A | A | W | W | A | W | A | W | W | A | W | NA | A | W | W | A | W | NA | A | A | W | W | A | A | W | A | W | W | A | W |
| ChrIV | 18.3 | W | W | W | A | A | W | A | W | W | W | A | W | A | A | W | W | A | A | A | A | A | W | A | W | A | A | A | A | W | A | A | A | A | A | A | A | A | A | A | A | A | A | W | W | W | A | W | A | W | W | A | A | A | A | W | W | W | W | W | W | W | W | A | A | W | W | A | W | A | W | W | A | W | W | A | W | W | A | W | W | A | A | W | W | A | A | W | A | W | W | A | W |
| ChrIV | 18.4 | W | W | W | A | A | W | A | W | W | W | A | W | A | A | NA | NA | A | A | A | A | A | W | A | W | A | A | A | A | W | A | A | A | A | A | A | A | A | A | A | A | A | A | W | W | W | A | W | A | W | W | A | A | A | A | W | W | W | W | W | W | W | W | A | A | W | W | A | W | A | W | NA | A | W | W | A | W | W | A | W | W | A | A | W | W | A | A | W | A | W | W | A | W |
| ChrIV | 18.5 | W | NA | W | A | A | W | A | W | W | W | A | W | A | A | W | W | A | A | A | NA | A | W | A | W | A | A | A | A | W | A | A | A | A | A | A | A | A | A | A | A | A | A | NA | NA | NA | A | W | A | W | W | A | A | A | A | W | W | W | NA | W | NA | W | W | A | A | W | NA | A | W | A | W | NA | A | W | NA | A | W | NA | A | NA | NA | A | A | W | W | A | NA | W | A | W | W | A | W |
| ChrIV | 18.6 | W | W | W | A | A | W | A | NA | W | W | A | W | A | A | W | W | A | A | A | A | NA | W | A | W | A | A | A | A | W | A | A | A | A | A | A | A | A | A | A | A | A | A | W | W | W | A | W | A | W | W | A | A | A | A | W | W | W | W | W | W | W | W | A | A | W | W | A | W | A | W | W | A | W | NA | A | W | W | A | W | W | A | A | W | W | A | A | W | A | W | W | A | NA |
| ChrIV | 18.8 | W | W | W | A | A | W | A | W | W | W | A | W | A | A | W | NA | A | A | A | A | A | W | A | W | A | A | A | A | W | A | A | A | A | A | A | A | A | A | A | A | A | A | W | W | W | A | W | A | W | W | A | A | A | A | W | W | W | W | W | W | W | W | A | A | W | NA | A | W | A | W | W | A | W | W | A | W | W | A | W | W | A | A | W | W | A | A | W | A | W | W | A | W |
| ChrIV | 18.9 | W | W | W | A | A | W | A | W | W | W | A | W | A | A | W | W | A | A | A | NA | A | W | A | W | A | A | A | A | W | A | A | A | A | A | A | A | A | A | A | A | A | A | W | W | W | A | W | A | W | W | A | A | A | A | W | W | W | W | W | W | W | W | A | A | W | W | A | W | A | W | W | A | W | W | A | W | W | A | W | W | A | A | W | W | A | A | W | A | W | W | A | W |
| ChrIV | 19.1 | W | W | W | A | A | W | A | W | W | W | A | W | A | A | W | NA | A | A | A | A | A | W | A | W | A | A | A | A | W | A | A | A | A | A | A | A | A | A | A | A | A | A | W | W | W | A | W | A | W | W | A | A | A | A | NA | W | W | W | W | W | W | W | A | A | W | W | A | W | A | W | W | A | W | W | A | W | W | A | W | W | A | A | W | W | A | A | W | A | W | W | A | W |
| ChrIV | 19.2 | W | W | W | A | A | W | A | W | W | W | A | W | A | A | W | W | A | A | A | A | A | W | A | W | A | A | A | A | W | A | A | A | A | A | A | A | A | A | A | A | A | A | W | W | W | A | W | A | W | W | A | A | A | A | W | W | W | W | W | W | W | W | A | A | W | W | A | W | A | W | W | A | W | W | A | W | W | A | W | W | A | A | W | W | A | A | W | A | W | W | A | W |
| ChrIV | 19.3 | W | W | W | A | A | W | A | W | W | W | A | W | A | A | W | W | A | A | A | A | A | W | A | W | A | A | A | A | W | A | A | A | A | A | A | A | A | A | A | A | A | A | W | W | W | A | W | A | W | W | A | A | A | A | W | W | W | W | W | W | W | W | A | A | W | W | A | W | A | W | W | A | W | W | A | W | W | A | W | W | A | A | W | W | A | A | W | A | W | W | A | W |
| ChrIV | 19.4 | W | W | W | A | A | W | A | W | W | W | A | W | A | A | W | W | A | A | A | A | A | W | A | W | A | A | A | A | W | A | A | A | A | A | A | A | A | A | A | A | A | A | W | W | W | A | W | A | W | W | A | A | A | A | W | W | W | W | W | W | W | W | A | A | W | W | NA | W | A | W | W | A | W | W | A | W | W | A | W | W | A | A | W | W | A | A | W | A | W | W | A | W |
| ChrIV | 19.5 | W | W | W | A | A | W | A | W | W | W | A | W | A | A | W | NA | A | A | A | A | A | W | A | W | A | A | A | A | W | A | A | A | A | A | A | A | A | A | A | A | A | A | W | W | W | A | W | A | W | W | A | A | A | A | W | W | W | W | W | W | W | W | A | A | W | W | A | W | A | W | W | A | W | W | A | W | W | A | W | W | A | A | W | W | A | A | W | A | W | W | A | W |
| ChrIV | 19.6 | W | W | W | A | A | W | A | W | W | W | A | W | A | A | W | W | A | A | A | A | A | W | A | W | A | A | NA | A | W | A | A | A | A | A | A | A | A | A | A | A | A | A | W | W | W | A | W | A | W | W | A | A | A | A | W | W | W | W | W | W | W | W | A | A | W | W | A | W | A | W | W | A | W | W | A | W | W | A | W | W | A | A | W | W | A | A | W | A | W | W | A | W |
| ChrIV | 19.7 | W | W | W | A | A | W | A | W | W | W | A | W | A | A | W | W | A | A | A | A | A | W | A | W | A | A | A | A | W | A | A | A | A | A | A | A | A | A | A | A | A | A | W | W | W | A | W | A | W | W | A | A | A | A | W | W | W | W | W | W | W | W | A | A | W | W | A | W | A | W | W | A | W | W | A | W | W | A | W | W | A | A | W | W | A | A | W | A | W | W | A | W |
| ChrIV | 19.8 | W | W | W | A | A | W | A | W | W | W | A | W | A | A | W | W | A | A | A | A | A | W | A | W | A | A | A | A | W | A | A | A | NA | A | A | A | A | A | A | A | A | A | W | W | W | NA | W | A | W | W | A | A | A | A | W | W | W | W | W | W | W | W | A | A | W | W | A | W | A | W | W | A | W | W | A | W | W | NA | W | W | A | A | W | W | A | A | W | A | W | W | A | W |
| ChrIV | 19.9 | W | W | W | A | A | W | A | W | W | W | A | W | A | A | W | W | A | A | A | A | A | W | A | W | A | A | NA | A | W | A | A | A | A | NA | A | A | A | A | A | A | A | A | W | W | W | NA | W | A | W | W | A | A | A | A | W | W | W | W | W | W | W | W | A | A | W | W | A | W | A | W | W | A | W | W | A | W | W | A | W | W | A | A | W | W | A | NA | W | A | W | W | A | W |
| ChrIV | 20 | W | W | W | A | A | W | A | W | W | W | A | W | A | A | W | W | A | A | A | A | A | W | A | W | A | A | A | A | W | A | A | A | A | A | A | A | A | A | A | A | A | A | W | W | W | A | W | A | W | W | A | A | A | A | W | W | W | W | W | W | W | W | A | A | W | W | A | W | A | W | W | A | W | W | A | W | W | A | W | W | A | A | W | W | A | A | W | A | W | W | A | W |
| ChrIV | 20.1 | W | W | W | A | A | W | A | W | W | W | A | W | A | A | NA | W | A | A | A | A | A | W | A | W | A | A | A | A | W | A | A | A | A | A | A | A | A | A | A | A | A | A | W | W | W | A | W | A | W | W | A | A | A | A | W | W | W | W | W | W | W | W | A | A | W | W | A | W | A | W | W | A | W | W | A | W | W | A | W | W | A | A | W | W | A | A | W | A | W | W | A | W |
| ChrIV | 20.2 | W | W | W | A | A | W | A | W | W | W | A | W | A | A | W | W | A | A | A | A | A | W | A | W | A | A | A | A | W | A | A | A | A | A | A | A | A | A | A | A | A | A | W | W | W | A | W | A | W | W | A | A | A | A | W | W | W | W | W | W | W | W | A | A | W | W | A | W | A | W | W | A | W | W | A | W | W | A | W | W | A | A | W | W | A | A | W | A | W | W | A | W |
| ChrIV | 20.3 | W | W | W | A | A | W | A | W | W | W | A | W | A | A | W | W | A | A | A | A | A | W | A | W | A | A | NA | A | W | A | A | A | A | A | A | A | A | A | A | A | A | A | W | W | W | NA | W | A | W | W | A | A | A | A | W | W | W | W | W | W | W | W | A | A | W | W | A | W | A | W | W | A | W | W | A | W | W | A | W | W | A | A | W | W | A | A | W | A | W | W | A | W |
| ChrIV | 20.4 | W | W | W | A | A | W | A | W | W | W | A | W | A | A | W | W | A | A | A | A | A | W | A | W | A | A | NA | A | W | A | A | A | A | A | A | A | A | A | A | A | A | A | W | W | W | NA | W | A | W | W | NA | A | A | A | W | W | W | W | W | W | W | W | A | A | W | W | NA | W | A | W | W | A | W | W | A | W | W | A | W | W | A | A | W | W | A | A | W | A | W | W | A | W |
| ChrIV | 20.5 | W | W | W | A | A | W | A | W | W | W | A | W | A | A | W | W | A | A | A | A | A | W | A | W | A | A | W | A | W | A | A | A | A | A | A | A | A | A | A | A | A | A | W | W | W | A | W | A | W | W | A | A | A | A | W | W | W | W | W | W | W | W | A | A | W | W | NA | W | A | W | W | A | W | W | A | W | W | A | W | W | A | A | W | W | A | A | W | A | W | W | A | W |
| ChrIV | 20.6 | W | W | W | A | A | W | A | W | W | W | A | W | A | A | W | W | A | A | A | A | A | W | NA | W | A | A | W | A | W | A | A | A | A | A | A | A | A | A | A | A | A | A | W | W | W | A | W | A | W | W | A | A | A | A | W | W | W | W | W | W | W | W | A | A | W | W | A | W | A | W | W | A | W | W | A | W | W | A | W | W | A | A | W | W | A | A | W | A | W | W | A | W |
| ChrIV | 20.7 | W | W | W | A | NA | W | NA | NA | W | W | A | W | A | A | W | NA | A | NA | A | A | A | W | A | W | A | NA | W | A | W | A | NA | A | A | A | NA | A | A | NA | A | A | NA | A | W | NA | NA | A | W | A | NA | W | A | A | A | A | NA | W | W | NA | W | W | W | W | A | A | W | NA | A | W | A | W | NA | A | W | W | A | W | W | A | W | W | A | A | NA | W | A | A | W | NA | W | NA | A | W |
| ChrIV | 20.8 | W | W | W | A | A | W | A | W | W | W | A | W | A | A | W | W | A | A | A | A | A | W | A | W | A | A | W | A | W | A | A | A | A | A | A | A | A | A | A | A | A | A | W | W | W | A | W | A | W | W | A | A | A | A | W | W | W | W | W | W | W | W | A | A | W | A | A | W | A | W | W | A | W | W | A | W | W | A | W | W | A | A | W | W | A | A | W | A | W | W | A | W |
| ChrIV | 20.9 | W | W | W | A | A | W | A | W | W | W | A | W | A | A | W | W | A | A | A | A | A | W | A | W | A | A | W | A | W | A | A | A | A | A | A | A | A | A | A | A | A | A | W | W | W | A | W | A | W | W | A | A | A | A | W | W | W | W | W | W | W | W | A | A | W | A | A | W | A | W | W | A | W | W | A | W | W | A | W | W | A | A | W | W | A | A | W | A | W | W | A | W |
| ChrIV | 21 | W | W | W | A | A | W | A | W | W | W | A | W | A | A | W | W | A | A | A | A | A | W | A | W | A | A | W | A | W | A | A | A | A | A | A | A | A | A | A | A | A | A | W | W | W | A | W | A | W | W | A | A | A | A | W | W | W | W | W | W | W | W | A | A | W | A | A | W | A | W | W | A | W | W | A | W | W | A | W | W | A | A | W | W | A | A | W | A | W | W | A | W |
| ChrIV | 21.1 | W | W | W | A | A | W | A | W | W | W | A | W | A | A | W | W | A | A | A | A | A | W | A | W | A | A | W | A | W | A | A | A | A | A | A | A | A | A | A | A | A | A | W | W | W | A | W | A | W | W | A | A | A | A | W | W | W | W | W | W | W | W | A | A | W | A | A | W | A | W | W | A | W | W | A | W | W | A | W | W | A | A | W | W | A | A | W | A | W | W | A | W |
| ChrIV | 21.2 | W | W | W | A | A | W | A | W | W | W | A | W | A | A | W | W | A | A | A | A | A | W | A | W | A | A | W | A | W | A | A | A | A | A | A | A | A | A | A | A | A | A | W | W | W | A | W | A | W | W | A | A | A | A | W | W | W | W | W | W | W | W | A | A | W | A | A | W | A | W | W | A | W | W | A | W | W | A | W | W | A | A | W | W | A | A | W | A | W | W | A | W |
| ChrIV | 21.3 | W | W | W | A | A | W | A | W | W | W | A | W | A | A | NA | W | A | A | A | A | A | W | A | W | A | A | W | A | W | A | A | A | A | A | A | A | A | A | A | A | A | A | W | W | W | A | W | A | W | W | A | A | A | A | W | W | W | W | W | W | W | W | A | A | W | A | A | W | A | W | W | A | W | W | A | W | W | A | W | W | A | A | W | W | A | A | W | A | W | W | A | W |
| ChrIV | 21.4 | W | W | W | A | A | W | A | W | W | W | A | W | A | A | W | W | A | A | A | A | A | W | A | W | A | A | W | A | W | A | A | A | A | A | A | A | A | A | A | A | A | A | W | W | W | A | W | A | W | W | A | A | A | A | NA | W | W | W | W | W | W | W | A | A | W | A | A | W | A | W | W | A | W | W | A | W | W | A | W | W | A | A | W | W | A | A | W | A | W | W | A | W |
| ChrIV | 21.5 | W | NA | W | A | A | W | A | W | W | W | A | W | A | A | NA | W | NA | A | A | A | A | W | A | W | A | A | W | A | W | A | A | A | A | A | A | A | A | A | A | A | A | A | W | W | W | A | W | A | W | W | A | A | A | A | W | W | W | W | W | W | W | W | A | A | W | A | A | W | A | W | W | A | W | W | A | W | W | A | W | NA | A | A | W | W | A | A | W | A | W | W | A | W |
| ChrIV | 21.6 | W | W | W | A | A | W | A | W | W | W | A | W | A | A | W | W | A | A | A | A | A | W | A | W | A | A | W | A | W | A | A | A | A | A | A | A | A | A | A | A | A | A | W | W | W | A | W | A | W | W | A | A | A | A | W | W | W | W | W | W | W | W | A | A | W | A | A | W | A | W | W | A | W | W | A | W | W | A | W | W | A | A | W | W | A | A | W | A | W | W | A | W |
| ChrIV | 21.7 | W | W | W | A | A | W | A | W | W | W | NA | W | A | A | W | W | A | A | A | A | A | W | A | W | NA | A | W | A | W | A | A | NA | NA | A | A | NA | A | A | A | A | A | A | W | W | W | NA | W | A | W | W | NA | NA | NA | NA | W | W | W | W | W | W | W | W | NA | A | W | A | A | W | NA | W | W | A | W | W | A | W | W | A | W | W | A | NA | W | W | A | A | W | A | W | W | A | W |
| ChrIV | 21.8 | W | W | W | A | A | W | A | W | W | W | A | W | A | NA | W | W | NA | A | A | A | A | W | A | W | A | A | W | A | W | A | A | NA | NA | A | A | NA | A | A | A | A | A | A | W | W | W | A | W | A | W | W | A | A | A | A | W | W | W | W | W | W | W | W | NA | A | W | NA | NA | W | NA | W | W | A | W | W | A | W | W | A | W | W | NA | NA | W | W | NA | A | W | A | W | W | A | W |
| ChrIV | 21.9 | W | W | W | A | A | W | A | W | W | W | A | W | A | NA | W | W | A | A | A | A | NA | W | A | W | A | A | W | A | W | A | A | A | A | A | A | A | A | A | A | A | A | A | W | W | W | A | W | A | W | W | A | A | A | A | W | W | W | W | W | W | W | W | A | A | W | A | A | W | A | W | W | A | W | W | A | W | W | A | W | W | A | A | W | W | A | A | W | A | W | W | A | W |
| ChrIV | 22 | W | W | W | NA | A | W | A | W | W | W | NA | W | A | A | W | W | A | A | NA | A | A | W | A | W | NA | A | W | A | W | A | A | NA | NA | A | A | NA | A | A | A | A | NA | NA | W | W | W | NA | W | A | W | W | NA | NA | NA | NA | W | W | W | W | W | W | W | W | NA | A | W | NA | A | W | NA | W | W | A | W | W | A | W | W | NA | W | W | A | NA | W | W | NA | A | W | NA | W | W | A | W |
| ChrIV | 22.1 | W | W | W | A | A | W | A | W | W | W | A | W | A | A | W | W | NA | A | A | A | A | W | A | W | A | A | W | A | W | A | A | A | A | A | A | A | A | A | A | A | A | A | W | W | W | A | W | A | W | NA | A | A | A | A | W | W | W | W | W | W | W | W | A | A | W | A | A | W | A | W | W | A | W | W | A | W | W | A | W | W | A | A | W | W | A | A | W | A | W | W | A | W |
| ChrIV | 22.2 | W | W | W | A | A | W | A | W | W | W | A | W | A | A | W | W | A | A | A | A | A | W | A | W | A | A | W | A | W | A | A | A | A | A | A | A | A | A | A | A | A | A | W | W | W | A | W | A | W | W | A | A | A | A | W | W | W | W | W | W | W | W | A | A | W | A | A | W | A | W | W | A | W | W | A | W | W | A | W | W | A | A | W | W | A | A | W | A | W | W | A | W |
| ChrIV | 22.3 | W | W | W | A | A | W | A | W | W | W | A | W | A | A | W | NA | A | A | A | A | A | W | A | W | A | A | W | A | W | A | A | A | NA | A | A | A | A | A | A | A | A | A | W | W | W | A | W | A | W | NA | A | A | A | A | W | W | W | W | W | W | W | W | A | A | W | A | A | W | A | W | W | A | W | W | A | W | W | A | W | W | A | A | W | W | A | A | W | A | W | W | A | W |
| ChrIV | 22.4 | W | W | W | A | A | W | A | W | W | W | A | W | A | A | W | W | A | A | A | A | A | W | A | W | A | A | W | A | W | A | A | A | A | A | A | A | A | A | A | A | A | A | W | W | W | A | W | A | W | W | A | A | A | A | W | W | W | W | W | W | W | W | A | A | W | A | A | W | A | W | W | A | W | W | A | W | W | A | W | W | A | A | W | W | A | A | W | A | W | W | A | W |
| ChrIV | 22.5 | W | W | W | A | A | W | A | W | W | W | A | W | A | A | NA | W | A | A | A | A | A | W | A | W | A | A | W | A | W | A | A | A | A | A | A | A | A | A | A | A | A | A | W | W | W | A | W | A | W | W | A | A | A | A | W | W | W | W | W | W | W | W | A | A | W | A | A | W | A | W | W | A | W | W | A | W | W | A | W | W | A | A | W | W | A | A | W | A | W | W | A | W |
| ChrIV | 22.6 | W | W | W | A | A | W | A | W | W | W | A | W | A | A | W | W | A | A | A | A | A | W | A | W | A | A | W | A | W | A | A | A | A | A | A | A | A | A | A | A | A | A | W | W | W | A | W | A | W | W | A | A | A | A | W | W | W | W | W | W | W | W | A | A | W | A | NA | W | A | W | W | A | W | W | A | W | W | A | W | W | A | A | W | W | A | A | NA | A | W | W | A | W |
| ChrIV | 22.7 | W | W | W | A | A | W | A | W | W | W | A | W | A | A | W | W | A | A | A | A | NA | W | A | W | A | A | W | A | W | A | A | A | A | A | A | A | A | A | A | A | A | A | W | W | W | A | W | A | W | NA | A | A | A | A | W | W | W | W | W | W | W | W | A | A | W | NA | A | W | A | W | W | A | W | W | A | W | W | A | W | W | A | A | W | W | A | A | A | A | W | W | A | W |
| ChrIV | 22.8 | W | W | W | A | A | W | A | W | W | W | A | W | A | A | W | W | A | A | A | A | A | W | A | W | A | A | W | A | W | A | A | A | A | A | A | A | A | A | A | A | A | A | W | W | W | A | W | A | W | W | A | A | A | A | W | W | W | W | W | W | W | W | A | A | W | A | A | W | A | W | W | A | W | W | A | W | W | A | W | W | A | A | W | W | A | A | A | A | W | W | A | W |
| ChrIV | 22.9 | W | W | W | A | A | W | A | W | W | W | A | W | A | A | W | W | A | A | A | A | A | W | A | W | A | W | W | A | W | A | A | A | A | A | A | A | A | A | A | A | A | A | W | W | W | A | W | A | W | W | A | A | A | A | W | W | W | W | W | W | W | W | A | A | W | A | NA | W | A | W | W | A | W | W | A | W | W | A | W | W | A | A | W | W | A | A | NA | A | W | W | A | W |
| ChrIV | 23 | W | W | W | A | A | W | A | W | W | W | A | W | A | A | W | W | A | A | A | A | NA | W | NA | W | A | W | W | A | W | A | A | A | NA | A | A | NA | A | A | A | A | A | A | W | W | W | A | W | A | W | W | A | A | A | A | W | W | W | W | W | W | W | W | A | A | W | A | A | W | A | W | W | A | W | W | A | W | W | NA | W | A | A | A | W | W | A | A | A | A | W | W | A | W |
| ChrIV | 23.1 | W | W | W | A | A | W | A | W | W | W | A | W | A | NA | NA | W | A | A | A | A | A | W | A | W | A | W | W | A | W | A | A | A | A | A | A | W | A | A | A | A | A | A | W | W | W | A | W | A | W | W | A | A | A | A | W | W | W | W | W | W | W | W | A | A | W | A | A | W | A | W | W | A | W | W | A | W | W | A | W | A | A | A | W | W | A | A | A | A | W | W | A | W |
| ChrIV | 23.2 | W | W | W | A | A | W | A | W | W | W | A | W | A | A | W | W | A | A | A | A | A | W | A | W | A | W | W | A | W | A | A | A | A | A | A | W | A | A | A | A | A | A | W | W | W | A | W | A | W | W | A | A | A | A | W | W | W | W | W | W | W | W | A | A | W | A | A | W | A | W | W | A | W | W | A | W | W | A | W | A | A | A | W | W | A | A | A | A | W | W | A | W |

| | | | | | | | | | | | | | | | | | | | | | | | | | | | | | | | | | | | | | | | | | | | | | | | | | | | | | | | | | | | | | | | | | | | | | | | | | | | | | | | | | | | | | | | | | | | | | |
|---|---|---|---|---|---|---|---|---|---|---|---|---|---|---|---|---|---|---|---|---|---|---|---|---|---|---|---|---|---|---|---|---|---|---|---|---|---|---|---|---|---|---|---|---|---|---|---|---|---|---|---|---|---|---|---|---|---|---|---|---|---|---|---|---|---|---|---|---|---|---|---|---|---|---|---|---|---|---|---|---|---|---|---|---|---|---|---|---|---|---|---|---|---|
| ChrIV | 23.3 | W | W | W | A | A | W | A | W | W | W | A | W | A | A | W | W | A | A | A | A | A | W | A | W | A | W | W | A | W | A | A | A | A | A | A | W | A | A | A | A | A | A | W | W | W | NA | W | A | W | W | A | A | A | A | W | W | W | W | W | W | W | W | A | A | W | A | A | W | A | W | W | A | W | W | A | W | W | A | W | A | A | A | W | W | A | A | NA | A | W | W | A | W |
| ChrIV | 23.4 | W | W | W | A | A | W | A | W | W | W | A | W | A | A | W | W | A | A | A | A | A | W | A | W | A | W | W | A | W | A | A | A | A | A | A | W | A | A | A | A | A | A | W | W | W | A | W | A | W | W | A | A | A | A | W | W | W | W | W | W | W | W | A | A | W | A | A | W | A | W | W | A | W | W | A | W | W | A | W | A | A | A | W | W | A | A | A | A | W | W | A | A |
| ChrIV | 23.5 | W | W | W | A | A | W | A | W | W | W | A | NA | A | A | W | W | A | A | A | A | A | W | A | W | NA | W | W | A | W | A | A | A | A | A | A | W | A | A | A | A | A | A | W | W | W | A | W | A | W | W | A | A | A | A | W | W | W | W | W | W | W | W | A | A | W | A | A | W | A | W | W | A | W | W | A | W | W | A | W | A | A | A | W | W | A | A | NA | A | W | W | A | A |
| ChrIV | 23.6 | W | W | W | A | A | W | A | W | W | W | A | W | A | A | W | W | A | A | A | A | A | W | NA | W | W | W | W | A | W | A | A | A | A | A | A | W | A | A | A | A | A | A | W | W | W | A | W | A | W | W | A | A | A | A | W | W | W | W | W | W | W | W | A | A | W | A | A | W | A | W | W | A | W | W | A | W | W | A | W | A | A | A | W | W | A | A | A | A | W | W | A | A |
| ChrIV | 23.7 | W | W | W | A | A | W | A | W | W | W | A | W | A | A | W | W | A | A | A | NA | A | W | A | W | W | W | W | A | W | A | A | NA | A | A | A | W | A | A | A | A | A | A | W | W | W | A | W | A | W | W | A | A | A | A | W | W | W | W | W | W | W | W | A | A | W | A | A | W | A | W | W | A | W | NA | A | W | W | A | A | A | A | A | W | W | A | A | A | A | W | W | A | A |
| ChrIV | 23.8 | W | W | W | A | A | W | A | W | W | W | A | W | A | A | W | W | A | A | A | A | A | W | A | W | W | W | W | A | W | A | A | W | A | A | A | W | A | A | A | A | A | A | W | W | W | A | W | A | W | W | A | A | A | A | W | W | W | W | W | W | W | W | A | A | W | A | A | W | A | W | W | A | W | A | A | W | W | A | A | A | A | A | W | W | A | A | A | A | W | W | A | A |
| ChrIV | 23.9 | W | W | W | A | A | W | A | W | W | W | A | W | A | A | W | W | A | A | A | A | A | W | A | W | W | W | W | A | W | A | A | W | A | A | A | W | A | A | A | A | A | A | W | W | W | A | W | A | W | W | A | A | A | A | W | W | W | W | W | W | W | W | A | A | W | A | A | W | A | W | W | A | W | A | A | W | W | A | A | A | A | A | W | W | A | A | A | A | W | W | A | A |
| ChrIV | 24 | W | W | W | A | A | W | A | W | W | W | A | W | A | A | W | W | A | A | A | A | A | W | NA | W | W | W | W | A | W | A | A | W | NA | A | A | W | A | A | A | A | A | A | W | W | W | A | W | A | W | W | A | A | A | A | W | W | W | W | W | W | W | W | A | A | W | A | A | W | A | W | W | A | W | A | A | W | W | A | A | A | A | A | W | W | A | A | A | A | W | W | A | A |
| ChrIV | 24.1 | W | W | W | A | A | W | A | W | W | W | A | W | A | A | W | W | A | A | A | A | A | W | A | W | W | W | W | A | W | A | A | W | A | A | A | W | A | A | A | A | A | A | W | W | W | A | W | A | W | W | A | A | A | A | W | W | W | W | W | W | W | W | A | A | W | A | A | W | A | W | W | A | W | A | A | W | W | A | A | A | A | A | W | W | A | A | A | A | W | W | A | A |
| ChrIV | 24.2 | W | W | W | A | A | W | A | W | W | W | A | W | A | A | W | W | A | A | A | A | A | W | A | W | W | W | NA | A | W | A | A | W | A | A | A | W | A | A | A | A | A | A | W | W | W | A | W | A | W | NA | A | A | A | A | W | W | W | W | W | W | W | W | A | A | W | A | A | W | A | W | W | A | W | A | A | W | W | A | A | A | A | A | W | W | A | A | A | A | W | W | A | A |
| ChrIV | 24.3 | W | W | W | A | A | W | A | NA | W | W | A | W | A | A | W | W | A | A | A | A | A | W | A | NA | W | W | W | NA | W | A | A | W | A | A | A | W | A | A | A | A | A | A | W | NA | NA | A | W | A | W | NA | A | A | A | A | NA | W | W | NA | W | NA | W | W | A | NA | W | A | A | W | A | W | W | NA | W | NA | A | W | NA | A | A | A | A | A | NA | W | A | A | A | A | W | NA | A | NA |
| ChrIV | 24.4 | W | W | W | A | A | W | A | A | W | W | A | W | A | A | W | W | A | A | A | A | A | W | A | W | W | W | W | A | W | A | A | W | A | A | A | W | A | A | A | A | A | A | W | W | W | A | W | A | NA | NA | A | A | A | A | W | W | W | W | NA | W | W | W | A | A | W | A | A | W | A | W | NA | A | W | A | A | W | W | A | A | A | A | A | NA | W | A | A | A | A | A | W | A | A |
| ChrIV | 24.5 | W | W | W | NA | A | W | A | A | W | W | A | W | A | A | W | W | A | A | A | A | A | W | A | W | W | W | W | A | W | A | A | W | NA | A | A | W | A | A | A | A | A | A | W | W | W | NA | W | A | W | W | NA | A | A | A | W | W | W | W | A | W | W | NA | NA | A | W | A | NA | W | A | W | W | A | NA | A | A | W | W | NA | A | A | A | A | W | W | A | A | A | A | A | W | A | A |
| ChrIV | 24.6 | W | NA | W | A | A | W | A | A | W | W | A | W | A | A | W | W | A | A | A | A | A | W | A | W | W | W | NA | A | W | A | A | W | A | A | A | W | A | A | A | A | A | A | W | W | W | A | W | A | W | W | A | A | A | A | W | W | W | W | A | W | W | W | A | A | W | A | A | W | A | W | W | A | W | A | A | W | W | A | A | A | A | A | W | W | A | A | A | A | A | W | A | A |
| ChrIV | 24.7 | W | W | W | A | A | W | A | A | W | W | A | W | A | A | W | W | A | A | A | A | A | W | A | W | W | W | W | A | W | A | A | W | A | A | A | W | A | A | A | A | A | A | W | W | W | A | W | A | W | W | A | A | A | A | W | W | W | W | A | W | W | W | A | A | W | A | A | W | A | W | NA | A | W | A | A | W | W | A | A | A | A | A | W | W | A | A | A | A | A | W | A | A |
| ChrIV | 24.8 | W | W | W | A | A | W | A | A | W | W | A | W | A | A | W | W | NA | A | A | A | A | W | A | W | W | W | W | A | W | A | A | W | A | A | A | W | A | A | A | A | A | A | W | W | W | A | W | A | W | NA | A | A | A | A | W | W | W | W | A | W | W | W | A | A | W | A | A | W | A | W | W | A | W | A | A | W | W | A | A | A | A | A | W | W | A | A | NA | A | A | W | A | A |
| ChrIV | 24.9 | W | W | W | A | A | W | A | A | W | W | A | W | A | A | W | NA | W | A | A | A | W | W | A | W | W | W | W | A | W | A | A | W | A | A | A | W | A | A | A | A | A | A | W | W | W | A | W | A | W | W | A | A | A | A | W | W | W | W | A | W | W | W | A | A | W | A | A | W | A | W | A | A | W | A | A | W | W | A | A | A | A | A | W | W | A | A | A | A | A | W | A | A |
| ChrIV | 25 | W | W | W | A | A | W | A | A | W | W | A | W | A | A | W | W | W | A | A | A | W | W | A | W | W | W | W | A | W | A | A | W | A | A | A | W | A | A | A | A | A | A | W | W | W | A | W | A | W | W | A | A | A | A | W | W | W | W | A | W | W | W | A | A | W | A | A | W | A | W | A | A | W | A | A | W | W | A | A | A | A | A | W | W | A | A | A | A | A | W | A | A |
| ChrIV | 25.1 | W | W | W | A | A | W | A | A | W | W | A | W | A | A | W | W | W | A | A | A | W | W | A | W | W | W | W | A | W | A | A | W | A | A | A | W | A | A | A | A | A | A | W | W | W | A | W | A | W | W | A | A | A | A | W | W | W | W | A | W | W | W | A | A | W | A | NA | W | A | W | A | A | W | A | A | W | W | A | A | A | A | A | W | W | A | A | A | A | A | W | A | A |
| ChrIV | 25.2 | W | W | W | A | A | W | A | A | W | W | A | W | A | A | W | W | W | A | A | A | W | W | A | W | W | W | W | A | W | A | A | W | A | A | A | W | A | A | A | A | A | A | W | W | W | A | W | A | W | W | A | A | A | A | W | W | W | W | A | W | W | W | A | A | W | A | A | W | A | W | A | A | W | A | A | W | W | A | A | A | A | A | W | W | A | A | A | A | A | W | A | A |
| ChrIV | 25.3 | W | W | W | A | A | W | A | A | W | W | A | NA | A | A | NA | W | W | A | A | A | W | W | A | W | W | W | W | A | W | A | A | W | A | A | A | W | A | A | A | A | A | A | W | W | W | A | W | A | W | W | A | A | A | A | W | W | W | W | A | W | W | W | A | A | W | A | A | W | A | W | A | A | W | A | A | W | NA | A | A | A | A | A | W | W | A | A | A | A | A | W | A | A |
| ChrIV | 25.4 | W | W | W | A | A | W | A | A | W | W | A | NA | A | A | W | W | W | A | A | A | NA | W | A | W | W | W | W | NA | W | A | A | W | A | A | A | W | A | A | A | A | A | A | W | W | W | A | W | A | W | W | A | A | A | A | NA | W | W | W | A | W | W | W | A | A | W | A | A | W | A | W | A | A | W | A | A | W | W | A | A | A | A | A | W | W | A | A | A | W | A | W | A | A |
| ChrIV | 25.5 | W | W | W | A | A | W | A | A | W | W | A | NA | A | A | W | W | W | A | A | A | NA | W | A | W | W | NA | W | W | W | A | A | W | A | NA | A | W | A | A | A | A | A | A | NA | W | W | A | W | A | W | W | A | A | A | A | W | W | W | W | A | W | W | W | A | A | W | NA | A | W | A | W | A | A | W | A | A | W | W | A | A | A | A | A | NA | W | A | A | A | W | A | NA | A | A |
| ChrIV | 25.7 | W | W | W | NA | A | W | NA | NA | W | W | A | NA | A | A | W | W | W | NA | A | A | W | W | NA | W | W | W | W | NA | W | A | A | W | A | A | A | W | NA | A | A | A | A | A | W | W | W | NA | W | NA | W | W | NA | NA | A | A | W | W | W | W | NA | W | NA | W | NA | A | W | NA | A | W | A | W | NA | A | W | A | A | W | W | A | NA | NA | A | NA | W | W | A | A | NA | W | NA | W | A | NA |
| ChrIV | 25.8 | W | W | W | A | A | W | A | A | W | W | A | W | A | A | W | W | W | A | A | A | W | W | A | W | W | W | W | NA | W | A | A | W | A | A | A | W | A | A | A | A | A | A | W | W | W | A | W | A | W | W | A | NA | A | A | W | W | W | W | NA | W | NA | W | A | A | W | A | NA | W | A | W | A | A | W | A | A | W | W | A | A | A | A | A | W | W | A | A | NA | W | A | W | A | A |
| ChrIV | 26 | W | W | W | NA | A | W | A | A | W | W | A | W | A | A | W | W | W | A | A | A | NA | W | A | W | W | W | W | W | W | A | A | W | A | A | A | W | A | A | W | A | A | A | W | W | W | A | W | A | W | W | A | A | NA | A | W | W | W | W | A | W | W | W | A | A | W | A | A | W | A | W | A | A | W | A | A | W | W | A | A | A | A | A | W | W | A | A | A | W | A | W | A | A |
| ChrIV | 26.1 | W | W | W | A | A | W | A | A | W | W | A | W | A | A | W | W | W | A | A | A | NA | W | NA | W | W | W | W | W | W | A | A | W | A | A | A | W | A | A | W | A | A | A | W | W | W | A | W | A | W | W | A | A | W | A | W | W | W | W | A | W | W | W | A | A | W | A | A | W | A | W | A | A | W | A | A | W | W | A | A | A | A | A | W | W | A | A | A | W | A | W | A | A |
| ChrIV | 26.2 | W | W | W | A | A | W | A | A | W | W | A | W | A | A | W | W | W | A | A | A | W | W | A | W | W | W | W | W | W | A | A | W | A | A | A | W | A | A | W | A | A | A | W | W | W | A | W | A | W | W | A | A | W | A | W | W | W | W | A | W | W | W | A | A | W | A | A | W | A | W | A | A | W | A | A | W | W | A | A | A | A | A | W | W | A | A | A | W | A | W | A | A |
| ChrIV | 26.3 | W | W | W | A | A | W | A | A | W | W | A | W | A | NA | W | W | W | A | A | A | W | W | A | W | W | W | W | W | W | A | A | W | A | A | A | W | A | A | W | A | A | A | W | W | W | A | W | A | W | W | A | A | W | A | W | W | W | W | A | W | W | W | A | A | W | A | A | W | A | W | A | A | W | A | A | W | W | A | A | A | A | A | W | W | A | A | A | W | A | W | A | A |
| ChrIV | 26.4 | W | W | W | A | A | W | A | A | W | W | A | W | A | A | W | W | W | A | A | A | W | W | A | W | W | W | W | W | W | A | A | W | A | A | A | NA | A | A | W | A | A | A | W | W | W | A | W | A | W | W | A | A | W | A | W | W | W | W | A | W | W | W | A | A | W | A | A | W | A | W | A | A | W | A | A | W | NA | A | A | A | A | A | W | W | A | A | NA | W | A | W | A | A |
| ChrIV | 26.5 | W | W | W | A | A | W | A | A | W | W | A | W | A | A | W | W | W | A | A | A | W | W | A | W | W | W | W | W | W | A | A | W | A | A | W | W | A | A | W | A | W | A | W | W | W | A | W | A | W | W | A | A | W | A | W | W | W | W | A | W | W | W | A | A | W | A | A | W | A | W | A | A | W | A | A | W | W | A | A | A | A | A | W | W | A | A | A | W | A | W | A | A |
| ChrIV | 26.6 | W | W | W | A | A | W | A | A | W | W | A | W | A | A | W | W | W | A | A | A | W | W | A | W | W | W | W | W | W | A | A | W | A | A | W | W | A | A | W | A | W | A | W | W | W | A | W | A | A | W | A | A | W | A | W | W | W | W | A | W | W | W | A | A | W | A | A | W | A | W | A | A | W | A | A | W | W | A | A | A | A | A | W | W | A | A | A | W | A | W | A | A |
| ChrIV | 26.7 | W | W | W | A | A | W | A | A | W | W | A | W | A | A | W | W | W | A | A | A | W | W | A | W | W | W | W | W | W | A | A | W | A | A | W | W | A | A | W | A | W | A | W | W | W | A | W | A | A | NA | A | A | W | A | W | W | W | W | A | W | W | W | A | A | W | A | A | W | A | W | A | A | W | A | A | W | W | A | A | A | A | A | W | W | A | A | A | W | A | W | A | A |
| ChrIV | 26.8 | W | W | W | A | A | W | A | A | W | W | A | W | A | A | W | W | W | A | A | A | W | W | A | W | W | W | W | W | W | A | A | W | A | A | W | W | A | A | W | A | W | A | W | W | W | A | W | A | A | NA | A | A | W | A | W | W | W | W | A | W | W | W | A | A | W | A | A | W | A | W | A | A | W | A | A | W | W | A | A | A | A | A | W | W | A | A | A | W | A | W | A | A |
| ChrIV | 26.9 | W | W | W | A | A | W | A | A | W | W | A | W | A | A | W | W | W | A | A | A | W | W | A | W | W | W | W | W | W | A | A | W | A | A | W | W | A | A | W | A | W | A | W | W | W | A | W | A | A | W | A | A | W | A | W | W | W | W | A | W | W | W | A | A | W | A | A | W | A | W | A | A | W | A | A | W | W | A | A | A | A | A | W | W | A | A | A | W | A | W | A | A |
| ChrIV | 27 | W | W | W | A | A | W | A | A | W | W | A | NA | A | A | W | W | W | A | A | A | W | W | A | W | W | W | W | W | W | A | A | W | A | A | W | W | A | A | W | A | W | A | W | W | W | A | W | NA | A | W | A | A | W | A | W | W | W | W | A | W | W | W | A | A | W | A | A | W | A | W | NA | A | W | A | A | W | W | A | A | A | W | A | W | NA | A | A | A | W | A | W | A | A |
| ChrIV | 27.1 | W | W | W | A | A | W | A | A | W | W | A | W | A | A | W | W | W | A | A | A | W | W | A | W | W | W | W | W | W | A | A | W | A | A | W | NA | A | A | W | A | W | A | W | W | W | A | W | W | NA | W | A | A | W | A | W | W | W | W | A | W | NA | W | A | A | W | A | A | W | A | W | NA | A | W | A | A | W | W | A | A | A | W | A | W | A | A | A | A | W | A | W | A | A |
| ChrIV | 27.2 | W | W | W | A | A | NA | A | A | W | W | NA | W | A | A | NA | W | W | A | A | A | W | W | A | W | W | W | NA | W | W | A | A | W | A | A | W | W | A | A | W | NA | W | A | W | W | W | NA | W | W | A | W | A | A | W | A | W | W | NA | W | A | W | W | NA | A | A | NA | A | NA | W | A | W | A | A | NA | A | A | W | W | NA | A | A | W | A | W | NA | A | A | NA | W | A | W | NA | A |
| ChrIV | 27.3 | W | W | W | NA | A | W | A | A | W | W | A | W | A | A | W | W | W | A | A | A | W | W | A | W | W | W | W | W | W | A | A | W | NA | A | W | W | NA | A | W | A | W | NA | W | W | W | NA | W | W | A | W | NA | NA | W | A | W | W | W | W | A | W | W | W | A | A | W | A | A | W | A | W | A | A | NA | A | A | W | W | NA | A | A | W | A | W | A | A | A | A | W | A | W | NA | A |
| ChrIV | 27.4 | W | W | W | NA | A | W | A | A | W | W | NA | W | A | A | W | W | W | A | NA | NA | W | W | A | W | W | W | W | W | W | A | W | W | A | A | W | W | NA | A | W | A | W | NA | W | W | W | A | W | W | A | W | NA | A | W | A | W | W | W | W | NA | W | W | W | A | A | W | A | A | W | NA | W | A | A | W | A | A | W | W | W | A | NA | W | W | W | NA | A | A | NA | W | NA | W | A | A |
| ChrIV | 27.5 | W | W | W | A | A | W | NA | A | W | W | A | W | A | A | W | W | W | A | A | A | W | W | A | W | W | W | W | W | W | A | W | W | A | A | W | W | A | A | W | A | W | A | W | W | W | A | W | W | A | W | A | A | W | A | W | W | W | W | A | W | W | W | A | A | W | A | A | W | A | W | A | A | W | A | A | W | W | W | A | A | W | W | W | A | A | A | A | W | A | W | A | A |
| ChrIV | 27.6 | W | W | W | A | A | W | A | A | W | W | A | W | A | NA | W | W | W | A | A | A | W | W | A | W | W | W | W | W | W | A | W | W | A | NA | W | W | A | A | W | A | W | A | W | W | W | A | W | W | A | W | A | A | W | A | W | W | W | W | A | W | W | W | A | A | W | A | A | W | A | W | A | A | W | A | A | W | W | W | A | A | W | W | W | A | A | A | A | W | A | W | A | A |
| ChrIV | 27.7 | W | W | W | A | A | W | NA | A | W | W | A | W | A | A | W | W | W | A | A | A | W | W | A | W | W | W | W | W | W | A | W | W | A | A | W | W | A | A | W | A | W | A | W | W | W | A | W | W | A | NA | A | A | W | A | W | W | W | W | A | W | W | W | A | A | W | A | A | W | A | W | A | A | W | A | A | W | W | W | A | A | W | W | W | A | A | A | A | W | A | W | A | A |
| ChrIV | 27.8 | W | W | W | A | A | W | A | A | W | W | A | W | A | A | W | W | W | A | A | A | W | W | A | W | W | W | W | W | W | A | W | W | A | A | W | W | A | A | W | A | W | A | W | W | W | A | W | W | A | W | A | A | W | A | W | W | W | W | A | W | W | W | A | A | W | A | A | W | A | W | A | A | W | A | A | W | W | W | A | A | W | W | W | A | A | A | A | W | A | W | A | A |
| ChrIV | 27.9 | W | W | W | A | W | W | A | A | W | W | A | W | A | A | W | W | W | A | A | A | W | W | A | W | W | W | W | W | W | A | W | W | A | A | W | W | A | A | W | A | W | A | W | W | W | A | W | W | A | W | A | A | W | A | W | W | W | W | A | W | W | W | A | A | W | A | A | W | A | W | NA | A | W | A | A | W | W | W | A | A | W | W | W | A | A | A | A | W | A | W | A | A |
| ChrIV | 28 | W | W | W | A | W | W | A | A | W | W | A | W | A | A | W | W | W | A | A | A | W | W | A | W | W | W | W | W | W | A | W | W | A | A | W | W | A | A | W | A | W | A | W | W | W | A | W | W | A | W | A | A | W | A | W | W | W | W | A | W | W | W | A | A | W | A | A | W | A | W | A | A | W | A | A | W | W | W | A | A | W | W | W | A | A | A | A | W | A | W | A | A |
| ChrIV | 28.1 | W | W | W | A | W | W | A | A | W | W | A | NA | A | A | W | W | W | A | A | A | W | W | A | W | W | W | W | W | W | A | W | W | A | A | W | W | A | A | W | A | W | A | W | W | W | A | W | W | A | W | A | A | W | A | W | W | W | W | A | W | W | W | A | A | W | A | A | W | A | W | A | A | W | A | A | W | W | W | A | A | W | W | W | A | A | A | A | W | A | W | A | A |
| ChrIV | 28.2 | W | W | W | A | W | W | A | A | W | W | A | W | A | A | W | W | W | A | NA | A | W | W | A | W | W | W | W | W | W | A | W | W | A | A | W | W | A | A | W | A | W | A | W | W | W | NA | W | W | A | W | A | A | W | A | W | W | W | W | A | W | W | W | A | A | W | A | A | W | A | W | NA | A | W | A | A | W | W | W | A | NA | W | W | W | A | A | A | NA | W | A | W | A | NA |
| ChrIV | 28.3 | W | W | W | A | W | W | A | A | W | W | A | W | A | A | W | W | W | A | A | A | W | W | A | W | W | W | W | W | W | A | W | W | A | A | W | W | A | A | W | A | W | A | W | W | W | A | W | W | A | W | A | A | W | A | W | W | W | W | A | W | W | W | A | A | W | A | A | W | A | W | A | A | W | A | A | W | W | W | A | A | W | W | W | A | A | A | A | W | A | W | A | A |
| ChrIV | 28.4 | W | W | W | A | W | W | A | A | W | W | A | W | A | A | W | W | W | A | A | A | W | W | A | W | W | W | W | W | W | A | W | W | A | A | W | W | A | A | W | A | W | A | W | W | W | A | W | W | A | W | A | A | W | A | W | W | W | W | A | W | W | W | A | A | W | A | A | W | A | W | A | A | W | A | A | W | W | W | A | A | W | W | W | A | A | A | A | W | A | W | A | A |
| ChrIV | 28.5 | W | W | W | A | W | W | A | A | W | W | A | W | A | A | W | W | W | A | A | A | W | W | A | W | W | W | W | W | W | A | W | W | NA | A | W | W | A | A | W | NA | W | A | W | W | W | A | W | W | A | W | A | A | W | A | NA | W | W | W | A | NA | W | W | A | A | W | A | A | W | A | W | A | A | NA | A | A | W | W | W | A | A | W | W | W | A | A | A | A | W | A | W | A | A |
| ChrIV | 28.6 | W | W | W | A | W | W | A | A | W | W | A | W | A | A | W | W | W | A | A | A | W | W | A | W | W | W | W | W | W | A | W | W | W | A | W | W | A | A | W | W | W | A | W | W | W | A | W | W | A | W | A | A | W | A | W | W | W | W | A | W | W | W | A | A | W | A | A | W | A | W | A | A | A | A | A | W | W | W | A | A | W | W | W | A | A | A | A | W | A | W | A | A |
| ChrIV | 28.7 | W | W | W | A | W | W | A | A | W | W | A | W | A | A | W | W | W | A | A | A | W | W | A | W | W | W | W | W | W | A | W | W | W | A | W | W | A | A | W | W | W | A | W | W | W | A | W | W | A | W | A | A | W | A | W | W | W | W | A | W | W | W | A | A | W | A | A | W | A | W | A | A | A | A | A | W | W | W | A | A | W | W | W | A | A | A | A | W | A | W | A | A |
| ChrIV | 28.8 | W | W | W | A | W | W | A | A | W | W | A | W | A | A | W | W | W | A | A | A | W | W | A | W | W | W | W | W | W | A | W | W | W | A | W | W | A | A | W | W | W | A | W | W | W | A | W | W | A | W | A | A | W | A | W | W | W | W | A | W | W | W | A | A | W | A | A | W | A | W | A | A | A | A | A | W | W | W | A | A | W | W | W | A | A | A | A | W | A | W | A | A |
| ChrIV | 28.9 | W | W | W | A | W | W | A | A | W | W | A | W | A | A | W | W | W | A | A | A | W | W | A | W | W | W | W | W | W | A | W | W | W | A | W | W | A | A | W | W | W | A | W | W | W | A | W | W | A | W | A | A | W | A | W | W | W | W | A | W | W | W | A | A | W | A | A | W | A | W | A | A | A | A | A | W | W | W | A | A | W | W | W | A | A | A | NA | W | A | W | A | A |
| ChrIV | 29 | W | W | W | A | W | W | A | A | W | W | A | W | A | A | W | W | W | A | A | A | W | W | A | W | W | W | W | W | W | A | W | W | W | A | W | W | A | A | W | W | W | A | W | W | W | A | W | W | A | W | A | A | W | A | NA | W | W | W | A | W | W | W | A | A | W | A | A | W | A | W | A | A | A | A | A | W | W | W | A | A | W | W | W | A | A | A | NA | W | A | W | A | A |
| ChrIV | 29.1 | W | W | W | NA | W | W | A | A | W | W | A | W | A | A | W | W | W | A | A | NA | W | W | NA | W | W | W | W | W | W | A | W | W | W | A | W | W | A | A | W | W | W | A | W | W | W | A | W | W | A | W | A | A | W | A | W | W | W | W | A | W | W | W | A | A | W | A | A | W | NA | W | A | A | A | A | A | W | W | W | A | A | W | W | W | A | NA | A | A | W | A | W | A | A |
| ChrIV | 29.2 | W | W | W | NA | W | W | A | A | W | W | A | W | A | W | W | W | W | A | A | A | W | W | A | W | W | W | W | W | W | A | W | W | W | A | W | W | A | A | W | W | W | A | W | W | W | A | W | W | A | W | A | A | W | A | W | W | W | W | A | W | W | W | A | A | W | A | NA | W | NA | W | A | A | A | A | A | W | W | W | A | A | W | W | W | A | W | A | A | W | A | W | A | A |
| ChrIV | 29.3 | W | W | W | W | W | W | A | A | W | W | A | W | A | W | W | W | W | A | A | A | W | W | A | W | W | W | W | W | W | A | W | W | W | A | W | W | A | A | W | W | W | A | W | W | W | A | W | W | A | W | A | A | W | A | W | W | W | W | A | W | W | W | A | A | W | A | A | W | W | W | A | A | A | A | A | W | W | W | A | A | W | W | W | A | W | A | A | W | A | W | A | A |
| ChrIV | 29.5 | W | W | W | W | W | W | A | A | W | W | A | W | A | W | W | W | W | A | A | A | W | W | NA | W | W | W | W | W | W | A | W | W | W | A | W | W | A | A | W | W | W | A | W | W | W | A | W | W | A | W | A | A | W | A | W | W | W | W | A | W | W | W | A | A | W | A | A | W | W | W | A | A | A | A | A | W | W | W | A | A | W | W | W | A | W | A | NA | W | A | W | A | A |
| ChrIV | 29.6 | W | W | W | W | W | W | NA | A | W | W | A | W | A | W | W | W | W | A | A | A | W | W | A | W | W | W | W | W | W | A | W | W | W | A | W | W | A | A | W | W | W | A | W | W | W | A | W | W | A | W | A | A | W | A | W | W | W | W | A | W | W | W | A | A | W | A | A | W | W | W | A | A | A | A | A | W | W | W | A | A | W | W | W | A | W | A | A | W | A | W | A | A |
| ChrIV | 29.7 | W | W | W | W | W | W | A | A | W | W | A | W | A | W | W | W | W | A | A | A | W | W | A | W | W | W | W | W | W | A | W | W | NA | A | W | W | A | A | W | W | W | A | W | W | W | A | W | W | A | W | A | A | W | A | W | W | W | W | A | W | NA | W | A | A | W | A | A | W | W | W | A | A | A | A | A | W | W | W | A | A | W | W | W | A | W | A | A | W | A | W | A | A |
| ChrIV | 29.8 | W | W | W | W | W | W | A | A | W | W | A | W | A | W | W | W | W | A | A | A | W | W | NA | W | W | W | W | W | W | A | W | W | W | A | W | W | A | A | W | W | W | A | W | W | W | A | W | W | A | W | A | NA | W | NA | W | W | W | W | A | W | W | W | A | A | W | A | A | W | W | W | A | A | A | A | A | W | W | W | A | A | W | W | W | A | W | A | A | W | A | W | A | A |
| ChrIV | 29.9 | W | W | W | W | W | W | A | A | W | W | A | W | A | W | W | W | W | A | A | A | W | W | A | W | W | W | W | W | W | A | W | W | W | A | W | W | A | A | W | W | W | A | W | W | W | A | W | W | A | W | A | A | W | A | W | W | W | W | A | W | W | W | A | A | W | A | A | W | W | W | A | A | A | A | A | W | W | W | A | A | W | W | W | A | W | A | A | W | A | W | A | A |
| ChrIV | 30 | W | W | W | W | W | W | A | A | W | W | A | W | A | W | W | W | W | A | A | A | W | W | A | W | W | W | W | W | W | A | W | W | W | A | W | W | A | A | W | W | W | A | W | W | W | A | W | W | A | W | A | A | W | A | W | W | W | W | A | W | W | W | A | A | W | A | A | W | W | W | A | A | A | A | A | W | W | W | A | A | W | W | W | A | W | A | A | W | A | W | A | A |
| ChrIV | 30.1 | W | W | W | W | W | W | A | A | W | W | A | W | A | W | W | W | W | A | A | A | W | W | A | W | W | W | W | W | W | A | W | W | W | A | W | W | A | A | W | W | W | A | W | W | W | A | W | W | A | W | A | A | W | A | W | W | W | W | A | W | W | W | A | A | W | A | A | W | W | W | A | A | A | A | A | W | W | W | A | A | W | W | W | A | W | A | A | W | A | W | A | A |
| ChrIV | 30.2 | W | W | W | W | W | W | NA | A | W | W | A | W | A | W | W | W | W | A | A | A | W | W | NA | W | W | W | W | W | W | A | W | W | W | A | W | W | A | A | W | W | W | A | W | W | W | A | W | W | A | W | A | NA | W | NA | W | W | W | W | A | W | W | W | A | A | W | A | NA | W | W | W | A | A | A | A | A | W | W | W | A | A | W | W | W | A | W | A | NA | W | A | W | A | A |
| ChrIV | 30.3 | W | W | W | W | W | W | A | A | W | W | A | W | A | W | W | W | W | A | A | A | W | W | A | W | W | W | W | W | W | A | W | W | W | A | W | W | A | A | W | W | W | A | W | W | W | A | W | W | A | W | A | A | W | A | W | W | W | W | A | W | W | W | A | A | W | A | A | W | W | W | A | A | A | A | A | W | W | W | A | A | W | W | W | A | W | A | A | W | A | W | A | A |
| ChrIV | 30.4 | W | W | W | W | W | W | NA | A | W | W | A | W | A | W | W | W | W | A | A | NA | W | W | A | W | W | W | W | W | W | A | W | W | W | NA | W | W | A | A | W | W | W | A | W | W | W | A | W | W | A | W | A | A | W | A | W | W | W | W | A | W | W | W | A | A | W | NA | A | W | W | W | NA | A | A | A | A | W | W | W | A | A | W | W | W | A | NA | A | NA | W | A | W | A | A |
| ChrV | 0 | W | A | A | W | A | W | A | W | A | A | A | W | A | A | A | A | W | A | A | A | A | A | A | W | W | A | A | A | W | A | A | A | W | W | A | A | A | W | A | W | W | W | A | W | W | A | W | A | W | A | A | A | W | A | W | A | W | W | W | A | A | A | A | W | A | A | A | A | W | A | W | A | W | A | W | A | A | W | W | W | A | A | W | A | A | W | W | W | A | W | W | A |
| ChrV | 0.1 | W | A | A | W | A | W | A | W | A | A | A | W | A | A | NA | A | W | A | A | A | A | A | A | W | W | A | A | A | W | A | A | A | W | W | A | A | A | W | A | W | W | W | A | W | W | A | W | A | W | A | A | A | W | A | W | A | W | W | W | A | A | A | A | W | A | A | A | A | W | A | NA | A | W | A | W | A | A | W | W | W | A | A | W | A | A | W | NA | W | A | W | W | A |
| ChrV | 0.2 | W | A | A | W | A | W | A | W | A | A | A | W | A | A | NA | A | W | A | NA | A | A | A | A | W | W | A | A | A | W | A | A | A | W | W | A | A | A | W | A | W | W | W | A | W | W | A | W | A | W | A | A | NA | W | A | W | A | W | W | W | NA | A | NA | A | W | NA | A | A | A | W | NA | W | A | W | A | W | A | A | W | W | W | A | A | W | A | A | W | W | W | A | W | W | A |
| ChrV | 0.3 | W | A | A | W | A | W | A | W | A | A | A | W | A | A | A | A | W | A | A | A | A | A | A | W | W | A | A | A | W | A | A | A | W | W | A | A | A | W | A | W | W | W | A | W | W | A | W | A | W | A | A | A | W | A | W | A | W | W | W | A | A | A | A | W | A | A | A | A | W | A | W | A | W | A | W | A | A | W | W | W | A | A | W | A | A | W | W | W | A | W | W | A |
| ChrV | 0.4 | W | A | A | W | A | W | A | W | A | A | A | W | A | NA | A | A | W | A | A | A | A | A | A | W | W | A | A | A | W | A | A | A | W | W | A | A | A | W | A | W | W | W | A | W | W | A | W | A | W | A | A | A | W | A | W | A | W | W | W | A | A | A | A | W | A | A | A | A | W | A | W | A | W | A | W | A | A | W | W | W | A | A | W | A | A | W | W | W | A | W | W | A |
| ChrV | 0.5 | W | A | A | W | A | W | A | W | A | A | A | W | A | A | A | A | W | A | A | A | A | A | A | W | W | A | A | A | W | A | A | A | W | W | A | A | A | W | A | W | W | W | A | W | W | A | W | A | W | A | A | A | W | A | W | A | W | W | W | A | A | A | A | W | A | A | A | A | W | A | W | A | W | A | W | A | A | W | W | W | A | A | W | A | A | W | W | W | A | W | W | A |
| ChrV | 0.7 | W | A | A | W | A | W | A | W | A | A | A | W | A | A | A | A | W | A | A | A | A | A | A | W | W | W | A | A | W | A | A | A | W | W | A | A | A | W | A | W | W | W | A | W | W | A | W | A | W | A | A | A | W | A | W | A | W | W | W | A | A | W | A | W | A | A | A | A | W | A | W | A | W | A | W | A | A | W | W | W | A | A | W | A | A | W | W | W | A | W | W | A |
| ChrV | 0.8 | W | A | A | W | A | W | A | W | A | A | A | W | A | A | A | A | W | A | A | A | A | A | A | W | W | W | A | A | W | A | A | A | W | W | A | A | A | W | A | W | W | W | A | W | W | A | W | A | W | A | A | A | W | A | W | A | W | W | W | A | A | W | A | W | A | A | A | A | W | A | W | A | W | A | W | A | A | W | W | W | A | A | W | A | A | W | W | W | A | W | W | A |
| ChrV | 1 | W | A | NA | NA | NA | NA | A | NA | NA | A | NA | NA | A | NA | A | NA | W | A | A | A | A | A | A | W | W | W | A | A | W | A | NA | NA | W | W | A | A | A | W | A | W | W | W | A | W | NA | A | NA | A | W | A | NA | NA | W | A | NA | NA | W | W | W | NA | NA | W | NA | W | A | NA | A | A | W | A | W | NA | NA | A | W | A | A | NA | W | W | A | A | W | A | NA | W | W | NA | A | W | W | A |
| ChrV | 1.1 | W | A | A | W | NA | W | A | W | A | A | A | NA | A | NA | A | A | W | A | A | NA | A | A | A | W | W | W | NA | A | W | A | A | A | W | W | A | A | A | W | A | NA | W | W | A | W | NA | A | W | A | W | NA | NA | A | W | A | W | A | W | W | W | A | NA | W | A | W | A | A | A | A | W | A | NA | A | NA | A | W | A | A | W | W | W | A | NA | W | A | A | W | W | NA | A | W | W | A |
| ChrV | 1.2 | W | A | A | W | A | W | A | W | A | A | A | W | A | A | A | A | W | A | A | NA | A | A | A | W | W | W | A | A | W | A | A | A | W | W | A | NA | A | W | A | W | W | W | A | W | W | A | W | A | W | A | A | NA | W | A | W | A | W | W | W | NA | A | W | NA | W | NA | A | A | A | W | A | W | A | W | A | W | A | A | W | W | W | A | A | W | A | A | W | W | W | NA | W | W | A |
| ChrV | 1.3 | W | A | A | NA | A | NA | A | NA | A | A | A | NA | A | A | A | A | W | A | A | A | A | A | A | W | W | W | A | A | W | A | A | A | W | W | A | A | A | W | A | W | W | NA | A | W | NA | A | NA | A | W | A | A | NA | W | A | NA | A | W | W | NA | A | A | W | A | W | A | A | A | A | W | A | NA | A | NA | A | W | A | A | NA | W | W | A | A | W | A | A | NA | W | NA | A | W | W | A |
| ChrV | 1.4 | W | A | A | W | A | W | A | W | A | A | A | NA | A | A | A | A | W | A | A | A | A | A | A | W | W | W | A | A | W | A | A | A | W | W | A | A | A | W | A | W | W | W | A | W | W | A | W | A | W | A | A | A | W | A | W | W | W | W | A | A | A | W | A | W | A | A | NA | A | W | A | NA | A | W | A | W | A | A | W | W | W | A | A | W | A | A | W | W | W | A | W | W | A |
| ChrV | 1.5 | W | A | A | W | A | W | A | W | A | A | A | W | A | A | NA | A | W | A | A | A | A | A | NA | W | W | W | A | A | W | A | A | A | W | W | A | A | A | W | A | NA | W | W | NA | W | W | A | W | A | W | A | A | A | W | A | W | W | W | W | A | A | A | W | NA | W | A | A | A | A | W | A | W | A | W | A | W | A | A | W | W | W | A | A | W | A | A | W | W | W | A | W | W | A |
| ChrV | 1.6 | W | A | A | NA | A | W | A | W | A | A | A | W | A | A | W | A | W | A | A | A | A | A | A | W | W | W | NA | A | W | A | A | A | W | W | A | A | A | W | A | A | W | W | W | W | W | A | W | A | W | A | A | A | W | A | W | W | W | W | A | A | A | W | A | W | A | A | A | A | W | A | NA | A | W | A | W | A | A | W | W | W | A | A | W | A | A | W | W | W | A | W | W | A |
| ChrV | 1.7 | W | NA | A | W | A | NA | A | W | NA | NA | A | NA | A | A | W | A | W | NA | A | A | A | A | A | W | W | NA | A | A | W | A | A | NA | W | W | A | A | A | W | A | A | W | W | NA | W | NA | A | NA | A | W | A | A | A | W | A | NA | W | W | W | A | A | NA | W | NA | W | A | A | A | A | W | A | W | A | W | A | W | A | NA | NA | W | W | A | A | W | A | A | W | W | NA | NA | W | W | A |
| ChrV | 1.8 | W | A | A | W | A | W | A | W | A | A | A | W | A | A | W | A | W | A | A | A | A | A | A | W | W | W | A | A | W | A | A | A | NA | W | A | A | A | W | A | A | W | W | W | W | W | A | W | A | W | A | A | A | W | A | W | W | W | W | A | A | A | W | A | W | A | A | A | A | W | A | W | A | W | A | W | A | A | W | W | W | A | A | W | A | A | W | W | W | A | W | W | NA |
| ChrV | 2 | W | A | A | W | A | W | A | W | A | A | A | W | A | A | W | A | W | A | A | A | A | A | A | W | W | W | A | A | NA | A | A | A | W | W | A | A | A | W | A | A | W | W | W | W | W | A | W | A | W | A | A | A | W | A | W | W | W | W | A | A | A | W | A | W | NA | A | A | A | W | A | W | A | W | A | W | A | A | W | W | W | A | A | W | NA | A | W | W | W | A | W | W | A |
| ChrV | 2.1 | W | A | A | W | A | W | A | W | A | A | A | W | A | A | W | A | W | A | A | A | NA | A | A | W | W | W | A | A | W | A | A | A | W | W | A | A | A | W | A | A | W | W | W | W | W | A | W | A | W | A | A | NA | W | A | W | W | W | W | A | A | A | W | NA | W | A | A | A | A | W | A | W | A | W | A | W | A | A | NA | W | W | A | A | W | A | A | W | W | W | A | W | W | A |

| | | | | | | | | | | | | | | | | | | | | | | | | | | | | | | | | | | | | | | | | | | | | | | | | | | | | | | | | | | | | | | | | | | | | | | | | | | | | | | | | | | | | | | | | | | | | | | |
|---|---|---|---|---|---|---|---|---|---|---|---|---|---|---|---|---|---|---|---|---|---|---|---|---|---|---|---|---|---|---|---|---|---|---|---|---|---|---|---|---|---|---|---|---|---|---|---|---|---|---|---|---|---|---|---|---|---|---|---|---|---|---|---|---|---|---|---|---|---|---|---|---|---|---|---|---|---|---|---|---|---|---|---|---|---|---|---|---|---|---|---|---|---|
| ChrV | 2.2 | W | A | A | W | A | NA | A | NA | A | A | A | W | A | A | W | A | W | A | A | A | A | A | A | W | W | NA | A | A | W | A | A | A | W | W | A | A | A | W | A | A | W | W | W | W | W | A | W | A | W | A | A | A | W | A | W | W | W | W | A | A | A | W | A | W | A | A | A | A | W | A | NA | A | NA | A | W | A | A | NA | W | W | A | A | W | A | A | NA | W | W | A | W | W | A |
| ChrV | 2.3 | W | A | A | W | A | W | A | W | A | A | A | W | A | A | W | A | W | A | A | A | A | A | A | W | W | W | A | A | W | A | A | A | W | W | A | A | A | W | A | A | W | W | W | W | NA | A | W | A | W | NA | A | A | W | NA | W | W | W | W | A | A | A | W | NA | W | NA | A | A | A | W | A | NA | A | NA | A | W | A | A | NA | W | W | A | A | W | A | A | NA | W | W | A | W | W | A |
| ChrV | 2.4 | W | A | A | W | A | W | A | W | A | A | A | W | A | A | W | A | W | A | A | A | NA | A | A | W | W | W | A | NA | W | A | A | A | W | W | A | A | A | W | A | A | W | W | W | W | W | NA | W | A | W | NA | A | A | W | NA | W | W | W | W | A | A | A | W | A | W | NA | A | A | A | W | A | W | A | W | NA | W | A | A | NA | W | W | A | A | W | A | A | W | W | W | A | W | W | A |
| ChrV | 2.6 | W | A | A | W | A | W | A | W | A | A | A | W | A | A | W | A | W | A | A | A | A | A | A | W | W | W | A | A | W | A | A | A | W | W | A | A | A | W | A | A | W | W | W | W | W | A | W | A | W | A | A | A | W | A | W | W | W | W | A | A | A | W | A | W | A | A | A | A | W | A | W | A | W | A | W | A | A | NA | W | W | A | A | W | A | A | W | W | W | A | W | W | A |
| ChrV | 2.7 | W | A | A | W | A | NA | A | W | A | A | A | A | A | A | W | A | W | A | A | A | A | A | A | W | W | W | A | A | W | W | A | A | W | W | A | A | A | W | A | A | W | W | NA | W | W | A | W | A | W | A | A | A | W | A | W | W | W | W | A | A | A | W | A | W | A | NA | A | A | W | A | NA | A | NA | A | W | A | A | NA | W | W | A | A | W | A | A | W | W | NA | A | W | W | A |
| ChrV | 2.8 | W | A | A | W | A | W | A | W | A | A | A | A | A | A | W | A | W | A | A | A | A | A | NA | W | W | W | A | NA | W | W | A | A | W | W | A | A | A | W | A | A | W | W | W | W | W | A | W | A | W | A | A | A | W | A | W | W | W | W | A | A | A | W | NA | W | A | A | A | A | W | A | W | A | W | A | W | A | A | NA | W | W | A | A | W | A | A | W | W | W | A | W | W | NA |
| ChrV | 2.9 | W | A | A | W | A | W | A | W | A | A | A | A | A | A | W | A | W | A | A | A | A | A | A | W | W | W | A | A | W | W | A | A | W | W | A | A | A | W | A | A | W | W | NA | W | NA | A | W | A | W | A | A | A | W | A | W | NA | W | W | A | A | A | W | NA | W | A | A | A | A | W | A | W | A | NA | A | W | A | A | NA | W | W | A | A | W | A | A | W | W | W | A | W | W | A |
| ChrV | 3 | W | A | A | NA | A | W | A | NA | A | A | A | A | A | A | W | NA | W | A | A | A | A | A | A | W | W | W | A | A | NA | W | A | A | W | NA | A | A | A | W | A | A | W | NA | NA | W | W | A | NA | A | W | NA | A | A | W | A | NA | NA | W | W | A | A | A | W | NA | W | A | NA | A | A | W | A | NA | A | NA | A | W | A | A | W | W | W | A | A | W | A | NA | NA | W | NA | A | NA | W | A |
| ChrV | 3.1 | W | A | A | NA | A | W | A | NA | A | A | A | A | A | A | W | A | W | A | A | NA | A | A | A | W | W | W | A | W | W | W | A | A | NA | NA | NA | A | A | W | A | A | W | NA | NA | NA | NA | A | NA | A | W | NA | A | A | W | A | NA | NA | W | W | A | A | NA | W | NA | W | A | A | A | A | W | A | NA | A | NA | A | W | A | A | W | W | W | A | A | W | A | A | NA | W | NA | A | NA | W | A |
| ChrV | 3.2 | W | A | A | W | A | W | A | W | A | A | A | A | A | A | W | A | W | A | A | A | A | A | A | W | W | W | A | W | W | W | A | A | W | W | A | A | A | W | A | A | W | W | W | W | W | A | W | A | W | A | A | A | W | A | W | W | W | W | A | A | A | W | NA | W | A | A | A | A | W | A | W | A | W | A | W | A | A | NA | W | W | A | A | W | A | A | W | W | W | A | W | W | A |
| ChrV | 3.3 | W | A | A | W | A | W | A | NA | A | A | A | A | A | A | W | A | W | A | A | A | A | A | A | W | W | W | A | W | W | W | A | A | W | W | A | A | A | W | A | A | W | W | W | W | NA | A | W | A | W | A | A | A | W | A | W | W | W | W | A | A | A | W | NA | W | A | A | A | A | W | A | W | A | W | A | W | A | A | NA | W | W | A | A | W | A | A | W | W | W | A | W | W | A |
| ChrV | 3.6 | W | A | NA | W | A | W | A | W | NA | A | A | NA | NA | NA | W | A | W | A | A | A | A | A | A | W | W | W | A | W | NA | W | A | A | W | W | A | A | A | W | A | A | W | NA | W | W | NA | A | NA | A | W | NA | A | A | W | A | NA | NA | W | W | A | A | NA | W | W | W | A | A | A | A | W | A | NA | NA | NA | A | W | A | A | NA | W | W | A | NA | W | A | A | W | W | W | A | W | W | NA |
| ChrV | 3.7 | NA | A | A | W | A | W | NA | W | A | A | A | A | A | NA | NA | A | W | A | A | A | A | A | A | W | W | W | A | W | W | W | A | A | W | W | A | A | A | W | A | A | W | NA | NA | W | NA | A | W | A | W | A | NA | A | W | A | NA | NA | W | W | A | A | NA | W | W | W | A | NA | A | A | W | A | NA | A | NA | A | W | A | A | NA | W | W | A | A | W | A | A | W | W | NA | A | W | W | A |
| ChrV | 3.8 | W | A | A | W | A | W | A | W | A | A | A | A | A | A | W | A | W | A | A | A | A | A | A | W | W | W | A | W | NA | W | A | A | W | W | A | A | A | NA | A | A | W | W | W | W | W | A | W | A | W | NA | A | NA | W | A | NA | W | W | W | NA | A | A | W | W | W | NA | A | A | W | W | A | W | A | W | A | W | NA | A | NA | W | W | A | A | W | NA | A | W | W | W | A | W | W | NA |
| ChrV | 3.9 | W | A | A | W | A | W | A | W | A | A | A | A | A | A | W | A | W | A | A | NA | A | A | A | W | W | W | A | W | W | W | A | A | W | W | A | A | A | A | A | A | W | W | W | W | W | A | W | A | W | A | A | A | W | A | W | NA | W | W | A | A | A | W | W | W | A | A | A | W | W | A | NA | A | NA | A | W | A | A | NA | W | W | A | A | W | A | A | W | NA | W | A | W | W | A |
| ChrV | 4 | W | A | A | W | A | W | A | NA | A | A | A | A | A | A | W | A | W | A | A | A | A | A | A | W | W | NA | A | W | W | W | A | A | W | W | A | A | A | A | A | A | W | W | W | W | W | A | W | A | W | A | A | A | W | A | W | W | W | W | A | A | A | W | W | W | A | A | A | W | W | A | W | A | W | A | W | A | A | NA | W | W | A | A | W | A | A | W | A | W | A | W | W | A |
| ChrV | 4.1 | W | A | A | W | A | W | A | W | A | A | A | A | A | A | W | A | W | A | A | A | A | A | A | W | NA | W | A | W | W | W | A | A | W | W | A | A | A | A | A | A | W | W | W | W | NA | A | W | A | W | A | A | A | W | A | W | W | W | W | A | A | A | W | W | W | A | A | A | W | W | A | W | A | W | A | W | A | A | W | W | W | A | A | W | A | A | W | A | W | A | W | W | A |
| ChrV | 4.2 | W | A | A | W | A | W | A | NA | A | A | A | A | A | A | W | A | W | A | A | NA | A | A | A | W | W | NA | A | W | W | W | A | A | W | W | A | A | A | A | A | A | W | W | W | W | W | A | W | A | W | A | A | A | W | A | NA | W | W | W | A | A | A | W | W | W | A | A | A | W | W | A | W | A | NA | A | W | A | A | NA | W | W | A | A | W | A | A | W | A | W | A | W | W | A |
| ChrV | 4.3 | W | A | A | W | A | W | NA | W | A | A | A | A | A | A | W | A | W | A | A | A | A | A | A | W | W | W | A | W | W | W | A | A | W | NA | A | A | A | A | A | A | W | NA | W | W | W | A | W | A | W | A | A | A | W | A | NA | NA | W | W | A | A | NA | W | W | W | A | NA | A | W | W | A | W | A | NA | A | W | A | A | W | W | W | A | A | W | A | A | W | NA | NA | A | W | W | A |
| ChrV | 4.4 | W | A | A | NA | A | W | A | W | A | A | A | A | A | A | W | A | W | A | A | NA | A | A | A | W | W | W | A | W | W | W | A | A | W | W | A | A | A | A | A | A | W | W | W | W | W | A | W | A | W | A | A | A | W | A | W | W | W | W | A | A | A | W | W | W | A | A | A | W | W | A | W | A | W | A | W | A | A | NA | W | W | A | A | W | A | A | W | A | W | A | W | W | A |
| ChrV | 4.5 | W | A | A | W | A | W | A | W | A | A | A | A | A | A | W | A | W | A | A | NA | A | A | A | W | W | W | A | W | W | W | A | A | NA | W | A | A | A | A | A | A | W | W | W | W | W | A | W | A | W | A | A | A | W | A | W | W | W | W | A | A | A | W | W | W | A | A | A | W | W | A | W | A | W | A | W | A | A | NA | W | W | A | A | W | A | A | W | A | W | A | W | W | A |
| ChrV | 4.6 | W | NA | A | W | A | W | A | W | A | A | A | NA | A | NA | W | NA | W | A | A | W | A | NA | A | W | W | W | NA | W | W | W | A | A | W | W | A | A | A | A | A | A | W | W | W | A | W | A | W | A | W | NA | NA | NA | W | A | W | W | W | W | A | A | NA | W | W | W | A | A | A | W | W | W | W | NA | W | A | W | A | A | NA | W | W | A | A | W | A | NA | W | A | W | A | W | W | A |
| ChrV | 4.8 | W | A | A | NA | A | W | A | NA | A | A | A | A | A | A | NA | A | W | A | W | W | A | A | A | W | W | W | A | W | NA | W | A | A | W | W | A | A | A | A | A | A | W | W | W | A | W | A | W | A | W | A | A | A | W | A | W | W | W | W | A | A | A | W | W | W | A | A | A | W | W | W | W | A | W | A | W | A | A | NA | W | W | A | A | W | A | A | W | A | W | A | W | W | A |
| ChrV | 4.9 | NA | A | A | NA | A | W | A | NA | A | A | A | A | A | A | NA | A | W | A | W | W | A | A | A | W | W | W | A | W | W | W | A | A | W | NA | A | A | A | A | NA | A | W | W | W | A | W | NA | NA | A | W | A | A | A | W | A | W | W | W | W | A | A | A | W | W | W | NA | A | A | W | W | W | W | A | W | A | W | A | A | NA | W | W | A | A | W | A | A | W | NA | W | A | W | W | A |
| ChrV | 5 | W | A | A | W | A | W | A | W | A | A | A | A | A | A | W | A | W | A | W | W | NA | A | A | W | W | W | NA | W | W | W | A | NA | W | W | A | NA | A | NA | A | A | W | W | W | A | W | NA | W | NA | W | A | A | NA | W | A | W | W | W | W | NA | A | A | W | W | W | NA | A | A | W | W | W | W | A | W | NA | W | A | A | NA | W | W | A | A | W | NA | A | W | A | W | A | W | W | NA |
| ChrV | 5.2 | W | A | A | W | A | W | A | W | A | A | A | NA | NA | A | W | NA | W | A | W | W | NA | A | NA | W | W | W | A | W | W | W | NA | A | W | W | NA | A | A | NA | NA | A | W | W | W | A | W | NA | W | A | W | A | A | A | W | NA | W | W | W | W | NA | NA | A | W | W | W | A | NA | A | W | W | W | W | NA | W | A | W | NA | A | NA | W | W | NA | A | W | A | A | W | A | W | NA | W | W | NA |
| ChrV | 5.3 | W | A | A | W | A | W | A | NA | A | A | A | A | A | A | W | A | W | A | W | W | A | A | A | W | W | W | A | W | W | W | A | A | W | W | A | A | A | A | A | A | W | W | W | A | W | A | W | A | W | A | A | A | W | A | W | W | W | W | A | A | A | W | W | W | A | A | A | W | W | W | W | A | NA | A | W | A | A | NA | W | W | A | A | W | A | A | W | A | W | A | W | W | A |
| ChrV | 5.4 | W | A | A | NA | A | W | NA | W | A | A | A | NA | NA | A | W | NA | W | A | W | NA | A | NA | A | W | W | W | A | W | NA | W | A | A | NA | W | A | A | A | A | A | A | W | W | NA | A | W | A | NA | A | W | A | A | A | W | A | W | NA | W | W | A | A | A | W | W | W | A | A | A | W | W | W | NA | A | NA | A | W | A | A | NA | W | W | A | A | W | A | A | W | A | NA | A | W | W | A |
| ChrV | 5.5 | W | A | A | W | A | W | NA | W | A | A | A | A | A | NA | W | A | W | A | W | W | A | A | A | W | W | W | A | W | W | W | A | A | W | W | A | A | A | A | A | A | W | W | W | A | W | A | W | NA | A | A | W | A | W | W | W | W | A | A | A | W | W | W | A | A | A | W | W | W | NA | A | W | A | W | A | A | NA | W | W | A | A | W | A | A | W | A | W | A | W | W | A |
| ChrV | 5.6 | W | A | A | W | A | W | A | W | A | A | A | A | A | A | NA | A | W | A | W | W | A | A | A | W | W | W | A | W | W | W | A | A | W | W | A | A | A | A | A | A | W | W | W | A | W | A | W | A | W | NA | A | A | W | A | W | W | W | W | A | A | A | W | W | W | A | A | A | W | W | W | NA | A | W | A | W | A | A | NA | W | W | A | A | W | A | A | W | A | W | A | W | W | A |
| ChrV | 5.7 | W | A | A | NA | A | W | A | W | A | A | A | A | A | A | NA | A | W | A | W | NA | A | A | A | W | W | W | A | W | W | W | A | A | W | W | A | A | A | A | A | A | W | W | W | A | W | A | W | A | W | A | A | A | W | A | W | NA | W | W | A | A | A | W | W | W | A | A | A | W | W | W | W | A | NA | A | W | A | A | NA | W | W | A | A | W | A | A | W | A | W | A | W | W | A |
| ChrV | 5.8 | W | A | A | W | A | W | A | W | A | A | A | A | A | A | NA | A | W | A | W | W | A | A | A | W | W | W | A | W | W | W | A | A | W | W | A | A | A | A | A | A | W | W | W | A | W | A | W | A | W | A | A | A | W | A | W | W | W | W | A | A | A | W | W | W | A | A | A | W | W | W | NA | A | NA | A | W | A | A | NA | W | W | A | A | W | A | A | W | A | W | A | W | W | A |
| ChrV | 5.9 | W | A | A | W | A | W | A | W | A | A | A | A | A | A | NA | A | W | A | W | W | A | A | A | W | W | W | A | W | W | W | A | A | W | W | A | A | A | A | A | A | W | W | W | A | W | A | W | A | W | A | A | A | W | A | W | W | W | W | A | A | A | W | W | W | A | A | A | W | W | W | W | A | W | A | W | A | A | NA | W | W | A | A | W | A | A | W | A | W | A | W | W | A |
| ChrV | 6 | W | A | A | W | A | W | A | W | A | A | A | A | A | A | W | A | W | A | W | W | A | A | A | W | W | W | A | W | W | W | A | A | W | W | A | A | A | A | A | A | W | W | W | A | W | A | W | A | W | A | A | A | W | A | W | W | W | W | A | A | A | W | W | W | A | A | A | W | W | W | W | A | W | A | W | A | A | NA | W | W | A | A | W | NA | A | W | A | W | A | W | W | A |
| ChrV | 6.1 | W | A | A | W | A | W | A | W | A | A | A | A | A | A | W | A | W | A | W | W | A | A | A | W | W | W | A | W | W | W | A | A | W | W | A | A | A | A | A | A | W | W | W | A | W | A | W | A | W | NA | A | A | W | A | W | W | W | W | A | A | A | W | W | W | A | A | A | W | W | W | W | A | W | A | W | A | A | NA | W | W | A | A | W | W | A | W | A | W | A | W | W | A |
| ChrV | 6.2 | W | A | A | W | A | W | A | W | A | A | A | A | A | A | W | A | W | A | W | W | A | A | A | W | W | W | A | W | W | W | A | A | W | W | A | A | A | A | A | A | W | W | W | A | W | A | W | A | W | A | A | A | W | A | W | W | W | W | A | A | A | W | W | W | A | A | A | W | W | W | W | A | W | A | W | A | A | NA | W | W | A | A | W | W | A | W | A | W | A | W | W | A |
| ChrV | 6.3 | W | A | A | W | A | W | A | W | A | A | A | A | A | A | W | A | W | A | W | W | A | A | A | W | W | W | A | W | W | W | A | A | W | W | A | A | A | A | A | A | W | W | W | A | W | A | W | A | W | A | A | A | W | A | W | W | W | W | A | A | A | W | W | W | A | A | A | W | W | W | W | A | W | A | W | A | A | NA | W | W | A | A | W | W | A | W | A | W | A | W | W | A |
| ChrV | 6.4 | W | A | A | W | A | W | A | W | A | A | A | A | A | A | NA | A | W | A | W | W | A | A | A | W | W | W | A | W | W | W | A | A | W | W | A | A | A | A | A | A | W | W | W | A | W | A | W | A | W | A | A | A | W | A | W | W | W | W | A | A | A | W | W | W | A | A | A | W | W | W | NA | A | W | A | W | A | A | NA | W | W | A | A | W | W | A | W | A | W | A | W | W | A |
| ChrV | 6.5 | W | A | A | W | A | W | A | W | A | A | A | A | A | A | W | A | W | A | W | W | A | A | NA | W | W | W | A | W | W | W | A | A | W | W | A | A | A | A | A | A | W | W | W | A | W | A | W | A | W | A | A | A | W | A | W | W | W | W | A | A | A | W | W | W | A | A | A | W | W | W | W | A | W | A | W | A | A | NA | W | W | A | A | W | W | A | W | A | W | A | W | W | A |
| ChrV | 6.6 | W | A | A | W | A | W | A | W | A | A | A | A | A | A | NA | A | W | A | W | W | A | A | A | W | W | W | A | W | W | W | A | A | W | W | A | A | A | A | A | A | W | W | W | A | W | A | W | A | W | A | A | A | W | A | W | W | W | W | A | A | A | W | W | W | A | A | A | W | W | W | W | A | W | A | W | A | A | W | W | W | A | A | W | W | A | W | A | W | A | W | W | A |
| ChrV | 6.7 | W | A | A | W | A | W | A | W | A | A | A | A | A | A | W | A | W | A | W | W | A | A | A | W | W | W | A | W | W | W | A | A | W | W | A | A | A | A | A | A | W | W | W | A | W | A | W | A | W | A | A | A | W | A | W | W | W | W | A | A | A | W | W | W | A | A | A | W | W | W | W | A | W | A | W | A | A | W | W | W | A | A | W | W | A | W | A | W | A | W | W | A |
| ChrV | 6.8 | W | A | A | W | A | W | A | W | A | A | A | A | A | A | W | A | W | A | W | W | A | A | A | W | W | W | A | W | W | W | A | A | W | W | A | A | A | A | A | A | W | W | W | A | W | A | W | A | W | A | A | A | W | A | W | W | W | W | A | A | A | W | W | W | A | A | NA | W | W | W | W | A | W | A | W | A | A | W | W | W | A | A | W | W | A | W | A | W | A | W | W | A |
| ChrV | 6.9 | W | A | A | W | A | W | A | W | A | A | A | A | A | A | W | A | W | A | W | W | NA | A | A | W | W | W | A | W | W | W | A | A | W | W | A | A | A | A | A | A | W | W | W | A | W | NA | W | A | W | A | A | NA | W | A | W | W | W | W | A | A | A | W | W | W | A | NA | W | W | W | W | W | A | W | A | W | A | A | W | W | W | A | A | W | W | A | W | A | W | A | W | W | NA |
| ChrV | 7 | W | A | A | W | A | W | A | W | A | A | A | A | A | A | NA | A | W | A | W | W | A | A | A | W | W | W | A | W | W | W | A | A | W | W | A | A | A | A | A | A | W | W | W | A | W | A | W | A | W | A | A | A | W | A | W | W | W | W | A | A | A | W | W | W | A | A | W | W | W | W | W | A | W | A | W | A | A | W | W | W | A | A | W | W | A | W | A | W | A | W | W | A |
| ChrV | 7.1 | W | A | A | W | A | W | A | W | A | A | A | A | A | A | W | A | W | A | W | W | A | A | A | W | W | W | NA | W | W | W | A | A | W | W | A | A | A | A | A | A | W | W | W | A | W | A | W | A | W | A | A | NA | W | A | W | W | W | W | A | A | A | W | W | W | A | A | W | W | W | W | W | A | W | A | W | A | NA | W | W | W | A | A | W | W | A | W | A | W | A | W | W | A |
| ChrV | 7.2 | W | A | A | W | A | W | A | W | A | A | A | A | A | A | W | A | W | A | W | W | A | A | A | W | W | W | A | W | W | W | A | A | W | W | A | A | A | A | A | A | W | W | W | A | W | A | W | A | W | A | A | A | W | A | W | W | W | W | A | A | A | W | W | W | A | A | W | W | W | W | W | A | W | A | W | A | A | W | W | W | A | A | W | W | A | W | A | W | A | W | W | A |
| ChrV | 7.3 | W | A | A | W | A | W | A | NA | A | A | A | A | A | NA | NA | A | W | A | W | NA | A | A | A | W | W | NA | A | W | W | W | A | A | W | W | A | A | A | A | A | A | W | W | W | A | W | A | W | A | W | A | A | A | W | A | W | W | W | W | A | A | A | W | W | W | A | A | W | W | W | W | W | A | NA | A | W | A | A | W | W | W | A | A | W | W | A | W | A | W | A | W | W | A |
| ChrV | 7.5 | W | A | A | W | A | W | A | W | A | A | A | A | A | A | W | A | W | A | W | W | NA | A | A | W | W | W | A | W | W | W | A | A | W | W | A | A | A | A | A | A | W | W | W | A | W | A | W | A | W | A | A | NA | W | A | W | W | W | W | A | A | A | W | W | W | A | NA | W | W | W | W | W | A | W | A | W | A | A | W | W | W | A | A | W | W | A | W | A | W | A | W | W | A |
| ChrV | 7.6 | W | A | A | W | A | W | A | W | A | A | A | A | A | A | NA | A | W | A | W | W | A | A | A | W | W | W | A | W | W | W | A | A | W | W | A | A | A | A | A | A | W | W | W | A | W | A | W | A | W | A | A | A | W | A | W | W | NA | W | A | A | A | W | W | W | A | A | W | W | W | W | W | A | W | A | W | A | A | W | W | W | A | A | W | W | A | W | A | W | A | W | W | A |
| ChrV | 7.8 | W | A | A | W | A | W | A | W | A | NA | A | A | A | A | W | A | W | NA | W | W | A | A | NA | W | W | W | NA | W | W | W | A | A | W | W | A | A | A | A | A | A | W | W | W | A | W | A | W | A | W | NA | W | A | NA | A | W | A | W | W | W | W | A | A | A | W | W | W | A | A | W | W | W | W | W | A | W | A | W | A | A | W | W | W | A | A | W | W | A | W | NA | W | A | W | W | NA |
| ChrV | 7.9 | W | A | A | W | A | W | A | NA | A | A | A | A | A | A | W | A | W | A | W | W | A | A | A | W | W | W | A | W | W | W | A | A | W | W | A | A | A | A | A | A | W | W | W | A | W | A | W | A | W | A | A | A | W | A | W | W | W | W | A | A | A | W | W | W | A | A | W | W | W | W | W | A | W | A | W | A | A | W | W | W | A | A | W | W | A | W | A | W | A | W | W | A |
| ChrV | 8 | W | A | A | W | A | W | A | W | A | A | A | A | A | A | W | A | W | A | W | W | A | A | A | W | W | NA | A | W | W | W | A | A | W | W | A | A | A | A | A | A | W | W | W | A | W | A | W | A | W | A | A | A | W | A | W | W | W | W | A | A | A | W | W | W | A | A | W | W | W | W | W | A | W | A | W | A | A | W | W | W | A | A | W | W | A | W | A | W | A | W | W | A |
| ChrV | 8.1 | W | A | A | W | A | W | A | W | A | A | A | A | A | A | W | A | W | A | W | W | A | A | A | W | W | W | NA | W | W | W | A | A | W | W | A | A | A | A | A | A | W | W | W | A | W | A | W | A | W | A | A | A | W | A | W | W | W | W | A | A | A | W | W | W | A | A | W | W | W | W | W | A | W | A | W | A | A | W | W | W | A | A | W | W | A | W | A | W | A | W | W | A |
| ChrV | 8.2 | W | A | A | NA | A | W | A | NA | A | A | A | A | A | A | NA | A | W | A | W | NA | A | A | A | W | W | W | A | W | W | W | A | A | W | W | A | A | A | A | A | A | W | W | W | A | W | A | W | A | W | A | A | A | W | A | W | W | NA | W | A | A | A | W | W | W | A | A | W | W | W | W | NA | A | NA | A | W | A | A | W | W | W | A | A | W | W | A | W | A | W | A | W | W | A |
| ChrV | 8.3 | W | A | A | W | A | W | A | W | A | A | A | A | A | A | W | A | W | A | W | W | A | A | A | W | W | NA | A | W | W | W | A | A | W | W | A | A | A | A | A | A | W | W | W | A | W | A | W | A | W | NA | A | A | W | A | W | W | W | W | A | A | NA | W | W | W | A | NA | W | W | W | W | W | A | W | A | W | A | A | W | W | W | A | A | W | W | A | W | A | W | A | W | W | A |
| ChrV | 8.4 | NA | A | A | W | A | W | A | W | A | A | A | A | A | A | W | A | W | A | W | NA | A | A | A | W | W | W | NA | W | W | W | A | A | W | W | A | A | A | A | A | A | W | W | NA | A | W | A | W | A | W | NA | NA | A | W | A | NA | NA | NA | W | A | A | NA | W | W | W | A | A | W | W | W | W | W | A | W | A | W | A | A | W | W | W | A | A | W | W | A | W | A | NA | A | W | W | A |
| ChrV | 8.8 | W | A | A | W | A | W | A | W | A | A | A | A | A | A | W | A | W | A | W | W | NA | A | A | W | W | W | A | W | W | W | A | A | W | W | A | A | A | A | A | A | W | W | W | A | W | NA | W | NA | W | A | A | NA | W | A | W | W | W | W | A | NA | A | W | W | W | NA | A | W | W | W | W | W | A | W | A | W | NA | NA | W | W | W | NA | A | W | W | A | W | A | W | A | W | W | NA |
| ChrV | 9 | W | A | A | W | A | W | A | W | A | A | A | A | A | A | NA | A | W | A | W | W | A | A | A | W | W | W | A | W | W | W | A | A | W | W | A | A | A | A | A | A | W | W | W | A | W | A | W | A | W | A | A | A | W | A | W | W | W | W | A | A | A | W | W | W | A | A | W | W | W | W | W | A | W | A | W | A | A | W | W | W | A | A | W | W | A | W | A | W | A | W | W | A |
| ChrV | 9.1 | W | A | A | W | A | W | A | W | A | A | A | A | A | A | W | A | W | A | W | W | A | A | A | W | W | W | A | W | W | W | A | A | W | W | A | A | A | A | A | A | W | W | W | A | W | A | W | A | W | A | A | A | W | A | W | W | W | W | A | A | A | W | W | W | A | A | W | W | W | W | W | A | W | A | W | A | A | W | W | W | A | A | W | W | A | W | A | W | A | W | W | A |
| ChrV | 9.2 | W | A | A | W | A | W | A | W | A | A | A | A | A | A | W | A | W | A | W | W | A | A | A | W | W | W | A | W | W | W | A | A | W | W | A | A | A | A | A | A | W | W | W | A | W | A | W | A | W | A | A | A | W | A | W | W | W | W | A | A | A | W | W | W | A | A | W | W | W | W | W | A | W | A | W | A | A | W | W | W | A | A | W | W | A | W | A | W | A | W | W | A |
| ChrV | 9.3 | W | A | A | W | A | W | A | W | A | A | A | A | A | A | NA | A | W | A | W | W | A | A | A | W | W | W | A | W | W | W | A | A | W | W | A | A | A | A | A | A | W | W | W | A | W | A | W | A | W | A | A | A | W | A | W | W | W | W | A | A | A | W | W | W | A | A | W | W | W | W | NA | A | W | A | W | A | A | W | W | W | A | A | W | W | A | W | A | W | A | W | W | A |
| ChrV | 9.4 | NA | A | A | NA | A | W | A | W | A | A | A | A | A | A | NA | A | W | A | W | W | NA | A | A | W | NA | NA | A | W | NA | W | A | A | W | W | A | A | A | A | NA | A | NA | W | W | A | W | A | W | A | W | A | A | A | W | A | NA | W | NA | W | A | A | A | NA | W | W | A | A | W | W | W | W | NA | A | NA | A | W | A | A | W | W | NA | A | A | W | W | A | W | A | W | A | W | W | A |
| ChrV | 9.6 | W | A | A | W | A | W | A | W | A | A | A | A | A | A | NA | A | W | NA | W | W | A | A | NA | W | W | W | A | W | W | W | A | A | W | W | A | A | A | A | A | A | W | W | W | A | W | A | W | NA | W | A | A | A | W | A | W | W | W | W | NA | NA | A | W | W | W | NA | A | W | W | W | W | W | A | W | A | W | A | A | W | W | W | A | A | W | W | A | W | A | W | A | W | W | NA |
| ChrV | 9.7 | W | A | A | W | A | W | A | NA | A | A | A | A | A | A | W | A | W | A | W | W | A | A | A | W | W | W | A | W | W | W | A | A | W | W | A | A | A | A | A | A | W | W | W | A | W | A | W | A | W | NA | A | A | W | A | W | W | W | W | A | A | A | W | W | W | A | A | W | W | W | W | W | A | W | A | W | A | A | W | W | W | A | A | W | W | A | W | NA | W | A | W | W | A |
| ChrV | 9.8 | W | A | A | W | A | W | A | W | A | A | A | A | A | A | NA | A | W | A | W | W | NA | A | A | W | W | W | A | W | W | W | A | A | W | W | A | A | A | A | A | A | W | W | W | A | W | A | W | A | W | A | A | A | W | A | W | W | W | W | A | A | A | W | W | W | A | A | W | W | W | W | W | A | W | A | W | A | A | W | W | W | A | A | W | W | A | W | A | W | A | W | W | A |
| ChrV | 9.9 | W | A | A | W | A | W | A | W | A | A | A | A | A | A | W | A | W | A | W | W | A | A | A | W | W | W | A | W | W | W | A | A | W | W | A | A | A | A | A | A | W | W | W | A | W | A | W | A | W | A | A | A | W | A | W | W | W | W | A | A | A | W | W | W | A | A | W | W | W | W | W | A | W | A | W | A | A | W | W | W | A | A | W | W | A | W | A | W | A | W | W | A |
| ChrV | 10.4 | W | A | A | NA | A | W | A | W | A | A | A | A | A | A | NA | A | W | A | W | W | A | A | A | W | W | W | A | W | W | W | A | A | W | W | A | A | A | A | A | A | W | W | W | A | NA | A | W | A | W | A | A | A | W | A | W | W | W | W | A | A | A | W | W | W | A | A | W | W | W | W | W | A | W | A | W | A | A | W | W | W | A | A | W | W | A | W | A | W | A | W | W | A |
| ChrV | 10.5 | W | A | A | W | A | W | A | W | A | A | A | A | A | A | W | A | W | A | W | W | A | A | A | W | W | W | A | W | W | W | A | A | W | W | A | A | A | A | A | A | W | W | W | A | W | A | W | A | W | A | A | A | W | A | W | W | W | W | A | A | A | W | W | W | A | A | W | W | W | W | W | A | W | A | W | A | A | W | W | W | A | A | W | W | A | W | A | W | A | W | W | A |
| ChrV | 10.6 | W | A | A | NA | A | W | A | W | A | A | A | A | A | A | NA | A | W | A | W | NA | A | A | A | W | W | W | A | W | NA | W | A | A | W | W | A | A | A | A | A | A | W | NA | W | A | NA | A | W | A | W | NA | A | A | W | A | W | NA | NA | W | A | A | A | W | W | W | A | A | W | W | W | W | NA | A | NA | A | W | A | A | NA | W | W | A | A | W | W | A | W | A | NA | A | W | W | A |
| ChrV | 10.7 | W | A | A | W | A | W | A | W | A | A | A | A | A | A | W | A | W | A | W | W | A | A | A | W | W | W | A | W | W | W | A | A | W | W | A | A | A | A | A | A | W | W | W | A | W | A | W | A | W | NA | A | A | W | A | W | W | W | W | A | A | A | W | W | W | A | A | W | W | W | W | W | A | W | A | W | A | A | W | W | W | A | A | W | W | A | W | A | W | A | W | W | A |
| ChrV | 10.8 | W | A | A | W | A | W | A | W | A | A | A | A | A | A | W | A | W | A | W | NA | A | A | A | W | W | W | A | W | W | W | A | A | W | W | A | A | A | A | A | A | W | W | W | A | W | A | W | A | W | A | A | A | W | A | NA | W | NA | W | A | A | A | W | W | W | A | A | W | W | W | W | NA | A | W | A | W | A | A | W | W | W | A | A | W | W | A | W | A | W | A | W | W | A |
| ChrV | 10.9 | W | A | A | W | A | W | A | W | A | A | A | A | A | A | NA | A | W | A | W | W | A | A | A | W | W | W | A | W | W | W | A | A | W | W | A | A | A | A | A | A | W | W | W | A | W | A | W | A | W | A | A | A | W | A | W | W | W | W | A | A | A | W | W | W | A | A | W | W | W | W | W | A | W | A | W | A | A | W | W | W | A | A | W | W | A | W | A | W | A | W | W | A |
| ChrV | 11 | W | A | A | W | A | W | NA | W | A | A | A | A | A | A | W | A | W | A | W | W | A | A | A | W | W | W | A | W | W | W | A | A | W | W | A | A | A | A | A | A | W | W | W | A | W | A | W | A | W | A | A | A | W | A | W | W | NA | W | A | A | A | W | W | W | A | A | W | W | W | W | W | A | W | A | W | A | A | W | W | W | A | A | W | W | A | W | A | W | A | W | W | A |
| ChrV | 11.1 | W | A | A | W | A | W | A | W | A | A | A | A | A | A | W | A | W | A | W | W | A | A | A | W | W | W | A | W | W | W | A | A | W | W | A | A | A | A | A | A | W | W | W | A | W | A | W | A | W | A | A | A | W | A | W | W | W | W | A | A | A | W | W | W | A | A | W | W | W | W | W | A | W | A | W | A | A | W | W | W | A | A | W | W | A | W | A | W | A | W | W | A |
| ChrV | 11.2 | W | A | A | W | A | W | A | W | A | A | A | A | A | A | W | A | W | A | W | W | A | A | A | W | W | W | NA | W | NA | W | A | A | W | W | A | A | A | A | A | A | W | W | W | A | W | A | W | A | W | A | A | A | W | A | W | W | W | W | A | A | A | W | W | W | A | A | W | W | W | W | NA | A | NA | A | W | A | A | W | W | W | A | A | W | W | A | W | A | W | A | W | W | A |
| ChrV | 11.3 | W | A | A | W | A | W | A | NA | A | A | A | A | A | A | W | A | W | A | W | NA | A | A | A | W | W | W | A | W | W | W | A | A | W | W | A | A | A | A | A | A | W | W | W | A | W | A | W | A | W | A | A | A | W | A | W | W | W | W | A | A | A | W | W | W | A | A | W | W | W | W | W | A | W | A | W | A | A | W | W | W | A | A | W | W | A | W | A | W | A | W | W | A |
| ChrV | 11.4 | W | A | A | W | A | W | A | NA | A | A | A | A | A | A | NA | A | W | A | W | W | A | A | A | W | W | W | A | W | W | W | A | A | W | W | A | A | A | A | A | A | W | W | W | A | W | A | W | A | W | A | A | A | W | A | W | NA | W | W | A | A | A | W | W | W | A | A | W | W | W | W | W | A | NA | A | W | A | A | W | W | W | A | A | W | W | A | W | A | W | A | W | W | A |
| ChrV | 11.5 | W | A | A | W | A | W | NA | W | A | A | A | A | A | A | W | NA | W | A | W | W | A | A | A | W | W | W | A | W | W | W | A | A | W | W | A | A | A | A | A | A | W | W | W | A | W | A | W | A | W | A | A | A | W | A | W | W | W | W | A | A | NA | W | W | W | A | A | W | W | W | W | W | A | W | A | W | A | A | W | W | W | A | A | W | W | A | W | A | W | A | W | W | A |
| ChrV | 11.6 | W | A | A | W | A | W | A | W | A | A | A | A | A | A | NA | A | W | A | W | W | NA | A | A | W | W | W | A | W | W | W | A | A | W | W | A | A | A | A | A | A | W | W | W | A | W | NA | W | A | W | NA | A | A | W | A | W | W | W | W | NA | NA | A | W | W | W | A | A | W | W | W | W | W | A | W | NA | W | A | A | W | W | W | A | A | W | W | A | W | A | W | A | W | W | NA |
| ChrV | 11.7 | W | A | A | W | A | W | A | W | A | A | A | A | A | A | NA | A | W | A | W | W | A | A | A | W | W | W | A | W | W | W | A | A | W | W | A | A | A | A | A | A | W | W | W | A | W | A | W | A | W | A | A | A | W | A | NA | W | NA | W | A | A | A | W | W | W | A | A | W | W | W | W | W | A | NA | A | W | A | A | W | W | W | A | A | W | W | A | W | A | W | A | W | W | A |
| ChrV | 11.8 | NA | A | A | W | A | W | A | W | A | A | A | A | A | A | NA | A | W | A | W | W | A | A | A | W | W | W | A | W | W | W | A | A | W | W | A | A | A | A | A | A | W | NA | W | A | NA | A | W | A | W | A | A | A | W | A | W | W | W | W | A | A | A | W | W | W | A | A | W | W | W | W | NA | A | NA | A | W | A | A | W | W | W | A | A | W | W | A | W | A | W | A | W | W | A |
| ChrV | 11.9 | NA | A | A | W | A | W | A | NA | A | A | A | NA | A | NA | NA | A | W | A | W | NA | A | A | A | W | W | W | A | W | W | W | A | A | W | W | A | A | A | A | A | A | W | W | W | A | NA | A | W | A | NA | A | W | A | W | NA | NA | W | A | A | NA | W | W | W | A | A | W | W | W | W | NA | A | NA | A | W | A | A | NA | W | W | A | A | W | W | NA | W | A | NA | A | W | W | A |
| ChrV | 12 | W | A | A | W | A | W | A | W | A | A | A | A | A | A | NA | A | W | A | W | NA | A | A | A | W | W | W | A | W | W | W | A | A | W | W | A | A | A | A | A | A | W | W | W | A | W | A | W | NA | A | A | W | A | W | W | W | W | A | A | A | W | W | W | A | A | W | W | W | W | W | A | W | A | W | A | A | W | W | W | A | A | W | W | NA | W | A | NA | A | W | W | A |
| ChrV | 12.1 | W | A | A | W | A | W | A | NA | A | A | A | A | A | A | W | A | W | A | W | W | A | A | A | W | W | W | A | W | W | W | A | A | W | W | A | A | A | A | A | A | W | W | W | A | W | A | W | A | W | A | A | A | W | A | W | NA | W | W | A | A | A | W | W | W | A | A | W | W | W | W | NA | A | W | A | W | A | A | W | W | W | A | A | W | W | A | W | A | W | A | W | W | A |
| ChrV | 12.2 | W | A | A | W | A | W | A | W | A | A | A | A | A | A | NA | A | W | A | W | W | A | A | A | W | W | W | NA | W | W | W | A | A | W | W | A | A | A | A | A | A | W | W | W | A | W | A | W | A | W | A | A | A | W | A | W | W | W | W | A | A | A | W | W | W | A | A | W | W | W | W | NA | A | W | A | W | A | A | W | W | W | A | A | W | W | A | W | A | W | A | W | W | A |
| ChrV | 12.3 | W | A | A | W | A | W | A | W | A | A | A | A | A | A | W | A | W | A | W | W | A | A | A | W | W | W | A | W | W | W | A | A | W | W | A | A | A | A | A | A | W | W | W | A | W | A | W | A | W | A | A | A | W | A | NA | W | W | W | A | A | A | W | W | W | A | A | W | W | W | W | NA | A | W | A | W | A | A | W | W | W | A | A | W | W | A | W | A | W | A | W | W | A |
| ChrV | 12.4 | W | A | A | W | A | W | A | NA | A | A | A | A | A | A | W | A | W | A | W | W | A | A | A | W | W | NA | A | W | W | W | A | A | W | W | A | A | A | A | A | A | W | W | W | A | W | A | W | A | W | A | A | A | W | A | W | W | W | W | A | A | A | W | W | W | A | A | W | W | W | W | NA | A | W | A | W | A | A | W | W | W | A | A | W | W | NA | W | A | W | A | W | W | A |
| ChrV | 12.6 | W | A | A | W | A | W | A | W | A | A | A | A | A | A | W | A | W | A | W | W | A | A | A | W | W | W | A | W | W | W | A | A | W | W | A | A | NA | A | NA | A | W | W | W | NA | W | NA | W | NA | W | A | A | NA | W | NA | W | W | W | W | A | A | A | W | W | W | NA | A | W | W | W | W | W | A | W | NA | W | NA | NA | W | W | W | NA | A | W | W | A | W | A | W | A | W | W | A |

| | | | | | | | | | | | | | | | | | | | | | | | | | | | | | | | | | | | | | | | | | | | | | | | | | | | | | | | | | | | | | | | | | | | | | | | | | | | | | | | | | | | | | | | | | | | | | |
|---|---|---|---|---|---|---|---|---|---|---|---|---|---|---|---|---|---|---|---|---|---|---|---|---|---|---|---|---|---|---|---|---|---|---|---|---|---|---|---|---|---|---|---|---|---|---|---|---|---|---|---|---|---|---|---|---|---|---|---|---|---|---|---|---|---|---|---|---|---|---|---|---|---|---|---|---|---|---|---|---|---|---|---|---|---|---|---|---|---|---|---|---|---|
| ChrV | 12.7 | W | A | A | W | A | W | A | W | A | A | A | A | A | A | W | A | W | A | W | W | NA | A | NA | W | W | W | NA | W | W | W | A | A | W | W | A | A | A | A | NA | A | W | W | W | A | W | A | W | NA | W | A | A | NA | W | A | W | W | W | W | A | A | A | W | W | W | NA | A | W | W | W | W | W | A | W | A | W | NA | NA | W | W | W | A | A | W | W | A | W | A | W | NA | W | W | A |
| ChrV | 12.8 | W | A | A | W | A | W | NA | W | A | A | A | A | A | A | W | A | W | A | W | W | NA | A | NA | W | W | W | NA | W | W | W | A | A | W | W | A | A | A | A | A | A | W | W | W | A | W | A | W | A | W | A | A | A | W | A | W | W | W | W | A | A | A | W | W | W | NA | A | W | W | W | W | W | A | W | A | W | A | A | W | W | W | A | A | W | W | A | W | A | W | A | W | W | NA |
| ChrV | 12.9 | W | A | A | W | A | W | A | W | A | A | A | A | A | A | W | A | W | A | W | W | A | A | A | W | W | W | A | W | W | W | A | A | W | W | A | A | A | A | A | A | W | W | W | A | W | A | W | A | W | A | A | A | W | A | W | W | W | W | A | A | A | W | W | W | A | A | W | W | W | W | W | A | W | A | W | A | A | W | W | W | A | A | W | W | A | W | A | W | A | W | W | A |
| ChrV | 13 | W | A | A | W | A | W | A | W | A | A | A | A | A | A | W | A | W | A | W | W | A | A | A | W | W | W | A | W | W | W | A | A | W | W | A | A | A | A | A | A | W | W | W | A | W | A | W | A | W | A | A | A | W | A | NA | W | NA | W | A | A | A | W | W | W | A | NA | W | W | W | W | W | A | W | A | W | A | A | W | W | W | A | A | W | W | A | W | A | W | A | W | W | A |
| ChrV | 13.1 | W | A | A | W | A | W | A | W | A | A | A | A | A | A | W | A | W | A | W | W | A | A | A | W | W | W | A | W | W | W | A | A | W | W | A | A | A | A | A | A | W | W | W | A | W | A | W | A | W | NA | A | A | W | A | W | W | NA | W | A | A | A | W | W | W | A | A | W | W | W | W | NA | A | NA | A | W | A | A | W | W | W | A | A | W | W | A | W | A | W | A | W | W | A |
| ChrV | 13.2 | W | A | A | W | A | W | A | W | A | A | A | A | A | A | W | A | W | A | W | W | A | A | A | W | W | W | A | W | W | W | A | A | W | W | A | A | A | A | A | A | W | W | W | A | W | A | W | A | W | A | A | A | W | A | W | W | W | W | A | A | A | W | W | W | A | A | W | W | W | W | NA | A | W | A | W | A | A | W | W | W | A | A | W | W | A | W | NA | W | A | W | W | A |
| ChrV | 13.3 | NA | A | A | W | A | W | A | NA | A | A | A | A | A | A | NA | A | W | A | W | W | NA | A | NA | W | NA | W | A | W | W | W | A | A | W | W | A | A | A | A | A | A | W | NA | NA | A | NA | A | W | A | NA | A | A | A | W | A | W | NA | NA | W | NA | A | A | W | W | W | A | A | W | W | W | W | NA | A | NA | A | W | A | A | W | W | W | A | A | W | W | A | W | NA | W | NA | W | W | NA |
| ChrV | 13.4 | W | A | A | NA | A | W | A | W | A | A | A | A | A | A | NA | A | W | A | W | W | NA | A | A | W | W | W | A | W | W | W | A | A | NA | W | A | A | A | A | A | A | W | NA | W | A | W | A | W | A | W | A | A | A | W | A | W | NA | NA | W | A | A | A | W | W | W | A | A | W | W | W | W | W | NA | NA | A | W | A | A | W | W | W | A | A | W | W | A | W | A | W | A | W | W | A |
| ChrV | 13.5 | W | A | A | W | A | W | A | W | A | A | A | A | A | A | W | A | W | A | W | W | A | A | A | W | W | W | A | W | W | W | A | A | W | W | A | A | A | A | A | A | W | W | W | A | W | A | W | A | W | NA | A | A | W | A | W | W | W | W | A | A | A | W | W | W | A | A | W | W | W | W | W | A | W | A | W | A | A | W | W | W | A | A | W | W | A | W | A | W | A | W | W | A |
| ChrV | 13.6 | W | A | A | W | A | W | A | W | A | A | A | A | A | A | W | A | W | A | W | W | A | A | A | W | W | W | A | W | W | W | A | A | W | W | A | A | A | A | A | A | W | W | W | A | W | NA | W | A | W | A | A | A | W | A | W | W | W | W | A | A | A | W | W | W | A | A | W | W | W | W | W | A | W | A | W | A | A | W | W | W | A | A | W | W | A | W | A | W | A | W | W | A |
| ChrV | 13.7 | NA | A | A | W | A | W | A | W | A | A | A | A | A | A | NA | A | W | A | W | NA | A | A | A | W | W | W | A | W | W | W | A | A | W | W | A | A | A | A | A | A | W | NA | W | A | W | A | NA | A | W | A | A | A | W | A | W | NA | NA | W | A | A | A | W | W | W | A | A | W | W | W | W | NA | A | NA | A | W | A | A | W | W | W | A | A | W | W | A | W | A | W | A | W | W | A |
| ChrV | 13.8 | W | A | A | NA | A | W | A | NA | A | A | A | NA | A | A | NA | A | W | A | W | NA | A | NA | A | W | W | NA | A | W | W | W | A | A | W | W | A | A | A | A | A | A | W | NA | NA | A | NA | NA | NA | A | W | A | NA | A | W | A | NA | NA | NA | W | NA | A | A | W | W | W | NA | A | W | W | W | W | NA | NA | NA | A | W | NA | A | W | W | W | A | A | W | W | NA | W | NA | W | NA | W | W | NA |
| ChrV | 13.9 | W | A | A | W | A | W | A | W | A | A | A | A | A | A | W | A | W | A | W | W | NA | A | A | W | W | W | A | W | W | W | A | A | NA | W | A | A | A | A | A | A | W | W | W | A | W | A | W | A | W | A | A | A | W | A | W | W | W | W | A | A | A | W | W | W | A | A | W | W | W | W | NA | A | W | A | W | A | A | W | W | W | A | A | W | W | A | W | A | W | NA | W | W | NA |
| ChrV | 14 | W | A | A | W | A | W | A | W | A | A | A | A | A | A | W | A | W | A | W | W | NA | A | NA | W | W | W | NA | W | W | W | A | A | W | W | A | A | A | A | A | A | W | W | W | A | W | A | W | NA | W | A | A | A | W | A | W | W | W | W | NA | A | A | W | W | W | A | A | W | W | W | W | W | A | W | A | W | A | NA | NA | W | W | A | A | W | W | A | W | NA | W | NA | W | W | A |
| ChrV | 14.1 | W | A | A | NA | A | W | A | NA | A | A | A | A | A | A | NA | A | W | A | W | NA | A | A | A | W | W | NA | NA | W | NA | W | A | A | W | W | A | A | A | A | A | A | W | W | W | A | W | A | W | A | W | A | A | A | W | A | W | W | W | W | A | A | NA | W | W | W | A | NA | W | W | W | W | NA | A | NA | A | W | A | A | W | W | W | A | A | W | W | NA | W | A | W | A | W | W | A |
| ChrV | 14.2 | W | A | A | W | A | W | A | W | A | A | A | A | A | A | W | A | W | A | W | W | NA | A | A | W | W | W | A | W | W | W | A | A | W | W | A | A | A | A | A | A | W | W | W | A | W | A | W | A | W | A | A | A | W | A | W | W | W | W | A | A | A | W | W | W | A | A | W | W | W | W | W | A | W | A | W | A | A | W | W | W | A | A | W | W | A | W | A | W | A | W | W | A |
| ChrV | 14.3 | W | A | A | W | A | W | A | W | A | A | A | A | A | A | W | A | W | A | W | W | A | A | A | W | W | W | A | W | W | W | A | A | W | W | A | A | A | A | A | A | W | W | W | A | W | A | W | A | W | NA | A | A | W | A | W | W | W | W | A | A | A | W | W | W | A | A | W | W | W | W | W | A | W | A | W | A | A | W | W | W | A | A | W | W | A | W | A | W | A | W | W | A |
| ChrV | 14.4 | NA | A | A | W | NA | W | A | NA | A | A | A | A | A | A | NA | A | W | A | W | NA | A | A | A | W | W | NA | A | W | NA | W | A | A | W | W | A | A | A | A | A | A | W | W | W | A | NA | A | W | A | W | A | A | A | W | A | W | NA | NA | W | A | A | NA | W | W | W | A | A | W | W | W | W | NA | A | W | A | W | A | A | W | W | W | A | A | W | W | A | W | A | NA | A | W | W | A |
| ChrV | 14.5 | W | A | A | W | A | W | A | NA | A | A | A | A | A | A | W | A | W | A | W | W | A | A | A | W | W | W | A | W | W | W | A | A | W | W | A | A | A | A | A | A | W | W | NA | A | W | A | W | A | W | A | A | A | W | A | W | W | W | W | A | A | A | W | W | W | A | A | W | W | W | W | W | A | W | A | W | A | A | W | W | W | A | A | W | W | A | W | A | W | A | W | W | A |
| ChrV | 14.6 | W | A | A | W | A | W | NA | W | A | A | A | A | A | A | W | A | W | A | W | NA | A | A | A | W | W | W | NA | W | W | W | A | A | W | W | A | A | A | A | A | A | W | W | W | A | W | A | W | A | W | A | A | A | W | A | W | W | W | W | A | A | A | W | W | W | A | A | W | W | W | W | W | A | W | A | W | A | A | W | W | W | A | A | W | W | A | W | A | W | A | W | W | A |
| ChrV | 14.7 | W | A | A | W | A | W | A | W | A | A | A | A | A | A | W | A | W | A | W | W | A | A | A | W | W | W | A | W | W | W | A | A | W | W | A | A | A | A | A | A | W | W | W | A | W | A | W | A | W | A | A | A | W | A | W | W | W | W | A | A | A | W | W | W | A | A | W | W | W | W | W | A | W | A | W | A | A | W | W | W | A | A | W | W | A | W | A | W | A | W | W | A |
| ChrV | 14.8 | W | A | A | W | A | W | A | W | A | A | A | A | A | A | W | A | W | A | W | W | A | A | A | W | W | W | A | W | W | W | A | A | W | NA | A | A | A | A | A | A | W | W | W | A | W | A | W | A | W | A | A | A | W | A | W | W | NA | W | A | A | A | W | W | W | A | A | W | W | W | W | W | A | W | A | W | A | A | W | W | W | A | A | W | W | A | W | A | W | A | W | W | A |
| ChrV | 14.9 | W | A | A | W | A | W | A | W | A | A | A | A | A | A | W | A | W | A | W | W | A | A | A | W | W | W | A | W | W | W | A | A | W | W | A | A | A | A | A | A | W | W | W | A | W | A | W | A | W | A | A | A | W | A | W | W | W | W | A | A | A | W | W | W | A | A | W | W | W | W | W | A | W | A | W | A | A | W | W | W | A | A | W | W | A | W | A | W | A | W | W | A |
| ChrV | 15 | W | A | A | W | A | W | A | W | A | A | A | A | NA | NA | W | NA | W | A | W | W | NA | A | A | W | W | W | A | W | W | W | A | A | W | W | A | A | A | A | A | A | W | W | W | A | W | NA | W | NA | W | NA | A | A | W | A | W | W | W | W | NA | A | A | W | W | W | NA | A | W | W | W | W | W | A | W | NA | W | NA | NA | W | W | W | A | A | W | W | A | W | NA | W | A | W | W | A |
| ChrV | 15.1 | W | A | A | W | A | W | A | W | A | A | A | A | A | A | W | A | W | A | W | W | A | A | A | W | W | W | A | W | W | W | A | A | W | W | A | A | A | A | A | A | W | W | W | A | W | A | W | A | W | A | A | A | W | A | W | W | W | W | A | A | A | W | W | W | A | A | W | W | W | W | W | A | W | A | W | A | A | W | W | W | A | A | W | W | A | W | A | W | A | W | W | A |
| ChrV | 15.2 | W | A | A | W | A | W | A | W | A | A | A | A | A | A | W | A | W | A | W | W | A | A | A | W | W | W | A | W | W | W | A | A | W | W | A | A | A | A | A | A | W | W | W | A | W | A | W | A | W | A | A | A | W | A | W | W | W | W | A | A | A | W | W | W | A | A | W | W | W | W | W | A | W | A | W | A | A | W | W | W | A | A | W | W | A | W | NA | W | A | W | W | A |
| ChrV | 15.3 | W | A | A | W | A | W | A | W | A | A | A | A | A | A | W | A | W | A | W | W | A | A | A | W | W | W | A | W | W | W | A | A | W | W | A | A | A | A | A | A | W | W | W | A | W | A | W | A | W | NA | A | A | W | A | W | W | W | W | A | A | A | W | NA | W | A | A | W | W | W | W | W | A | W | A | W | A | A | W | W | W | A | A | W | W | A | W | A | W | A | W | W | A |
| ChrV | 15.4 | W | A | A | W | A | W | A | W | A | A | A | A | A | A | W | A | W | A | W | W | NA | A | NA | W | W | W | A | W | W | W | A | A | W | W | A | A | A | A | A | A | W | W | W | A | W | A | W | A | W | A | A | A | W | A | W | W | W | W | A | A | A | W | W | W | A | A | W | W | W | W | W | A | W | A | W | A | A | W | W | W | A | A | W | W | A | W | A | W | A | W | W | A |
| ChrV | 15.5 | W | A | A | W | A | W | A | W | A | A | A | A | A | A | W | A | W | A | W | W | A | A | A | W | W | W | A | W | W | W | A | A | W | W | A | A | A | A | A | A | W | W | W | A | W | A | W | A | W | A | A | A | W | A | W | W | W | W | A | A | A | W | W | W | A | A | W | W | W | W | W | A | W | A | W | A | A | W | W | W | A | A | W | W | A | W | A | W | A | W | W | A |
| ChrV | 15.6 | W | A | A | W | A | W | A | W | A | A | A | A | A | A | W | A | W | A | W | W | A | A | A | W | W | W | A | W | W | W | A | A | W | NA | A | A | A | A | A | A | W | W | W | A | W | A | W | A | W | A | A | A | W | A | W | W | W | W | A | A | A | W | W | W | A | A | W | W | W | W | NA | A | W | A | W | A | A | W | W | W | A | A | W | W | A | W | A | W | A | W | W | A |
| ChrV | 15.7 | W | A | A | W | A | W | A | W | A | A | A | A | A | A | W | A | W | A | W | W | A | A | A | W | W | W | A | W | W | W | A | A | W | W | A | A | A | A | A | A | W | W | W | A | W | A | W | A | W | A | A | A | W | A | W | W | W | W | A | A | A | W | W | W | A | A | W | W | W | W | W | A | W | A | W | A | A | W | W | W | A | A | W | W | A | W | A | W | A | W | W | A |
| ChrV | 15.8 | W | A | A | W | A | W | A | W | A | A | A | A | A | A | W | A | W | A | W | W | A | A | A | W | W | W | A | W | W | W | A | A | W | W | A | A | A | A | A | A | W | W | W | A | W | A | W | A | W | A | A | A | W | A | W | W | W | W | A | A | A | W | W | W | A | A | W | W | W | W | W | A | W | A | W | A | A | W | W | W | A | A | W | W | A | W | A | W | A | W | W | A |
| ChrV | 15.9 | W | A | A | W | A | W | A | W | A | A | A | A | A | A | NA | A | W | A | W | NA | A | A | A | W | W | NA | A | W | NA | W | A | A | W | NA | A | A | A | A | A | A | W | W | W | A | W | A | W | A | W | NA | A | A | W | A | W | W | W | W | A | A | A | W | W | W | A | A | W | W | W | W | NA | A | NA | A | W | A | A | W | W | W | A | A | W | W | A | W | A | W | A | W | W | A |
| ChrV | 16 | W | A | A | NA | A | W | A | NA | A | A | A | A | A | A | W | A | W | A | W | W | A | A | A | W | W | NA | A | W | W | W | A | A | W | W | A | A | A | A | A | A | W | W | W | A | W | A | W | A | W | A | A | A | W | A | NA | NA | NA | W | A | A | A | W | W | W | A | A | W | W | W | W | NA | A | W | A | W | A | A | W | W | W | A | A | W | W | A | W | A | W | A | W | W | A |
| ChrV | 16.1 | W | A | A | W | A | W | A | W | A | A | A | A | A | A | NA | A | W | A | W | W | A | A | A | W | W | W | A | W | W | W | A | A | W | W | A | A | A | A | A | A | W | W | W | A | W | A | W | A | W | A | A | A | W | A | W | W | W | W | A | A | A | W | W | W | A | A | W | W | W | W | NA | A | W | A | W | A | A | W | W | W | A | A | W | W | A | W | A | W | A | W | W | A |
| ChrV | 16.2 | W | A | A | W | A | W | A | W | A | A | A | A | A | A | NA | A | W | A | W | W | A | A | A | W | W | NA | A | W | W | W | A | A | W | NA | A | A | A | A | A | A | W | W | W | A | W | A | W | A | W | A | A | A | W | A | W | W | W | W | A | A | A | W | W | W | A | A | W | W | W | W | W | A | W | A | W | A | A | W | W | W | A | A | W | W | A | W | A | W | A | W | W | A |
| ChrV | 16.3 | W | A | A | W | A | W | A | W | A | A | A | A | A | A | W | A | W | A | W | W | A | A | A | W | W | W | A | W | W | W | A | A | W | W | A | A | A | A | A | A | W | W | W | A | W | A | W | A | W | A | A | A | W | A | W | W | W | W | A | A | A | W | W | W | A | A | W | W | W | NA | NA | A | W | A | W | A | A | W | W | W | A | A | W | W | A | W | A | W | A | W | W | A |
| ChrV | 16.4 | W | A | A | W | A | W | A | W | A | A | A | A | A | A | W | A | W | A | W | W | A | A | A | W | W | W | A | W | W | W | A | A | W | W | A | A | A | A | A | A | W | W | W | A | W | A | W | A | W | A | A | A | W | A | W | W | W | W | A | A | A | W | W | W | A | A | W | W | W | W | NA | A | W | A | W | A | A | W | W | W | A | A | W | W | A | W | A | W | A | W | W | A |
| ChrV | 16.5 | W | A | A | NA | A | W | NA | W | A | A | A | A | A | A | W | A | W | A | W | NA | A | A | A | W | W | W | A | W | W | W | A | A | W | W | A | A | A | A | A | A | W | W | NA | A | NA | A | W | A | W | A | A | A | W | A | NA | NA | W | W | A | A | A | W | W | W | A | A | W | W | W | NA | W | A | NA | A | W | A | A | W | W | W | A | A | W | W | NA | W | A | W | A | W | W | A |
| ChrV | 16.6 | W | A | A | W | A | W | A | W | A | A | A | A | A | A | W | A | W | A | W | W | A | A | A | W | W | W | A | W | W | W | A | A | W | W | A | A | A | A | A | A | W | W | W | A | W | A | W | A | W | A | A | A | W | A | W | W | W | W | A | A | A | W | W | W | A | A | W | W | W | W | NA | A | NA | A | W | A | A | W | W | W | A | A | W | W | A | W | A | W | A | W | W | A |
| ChrV | 16.7 | W | A | A | W | A | W | A | NA | A | A | A | A | A | A | W | A | W | A | W | NA | A | A | A | W | W | W | A | W | W | W | A | A | W | NA | A | A | A | A | A | A | W | W | W | A | W | A | W | A | W | A | A | A | W | A | W | NA | W | W | A | A | A | W | W | W | A | A | W | W | W | NA | W | A | W | A | W | A | A | W | W | W | A | A | W | W | A | W | A | W | A | W | W | A |
| ChrV | 16.8 | W | A | A | W | A | W | A | W | A | A | A | A | A | A | W | A | W | A | W | W | A | A | A | W | W | NA | A | W | W | W | A | A | W | NA | A | A | A | A | A | A | W | W | W | A | W | A | W | A | W | NA | A | A | W | A | W | NA | W | W | A | A | A | W | W | W | A | A | W | W | W | W | NA | A | W | A | W | A | A | W | W | W | A | A | W | W | A | W | A | W | A | W | W | A |
| ChrV | 16.9 | W | A | A | W | A | W | A | W | A | A | A | A | A | A | NA | A | W | A | W | W | NA | A | A | W | W | W | A | W | W | W | A | A | W | W | A | A | A | A | A | A | W | W | W | A | W | A | W | A | W | A | A | A | W | A | W | W | W | W | A | A | A | W | W | W | A | NA | W | W | W | W | W | A | W | A | W | A | A | W | W | W | A | A | W | W | A | W | A | W | A | W | W | A |
| ChrV | 17 | W | A | A | W | A | W | A | W | A | A | A | A | A | A | W | A | W | A | W | W | A | A | A | W | W | W | A | W | W | W | A | A | W | W | A | A | A | A | A | A | W | W | W | A | W | A | W | NA | W | A | A | A | W | A | W | W | W | W | A | A | A | W | W | W | A | A | W | W | W | W | W | A | W | A | W | A | A | W | NA | W | A | A | W | W | A | W | A | W | A | W | W | A |
| ChrV | 17.2 | W | A | A | W | A | W | A | W | A | A | A | A | A | A | W | A | W | A | W | W | A | A | A | W | W | W | A | W | W | W | A | A | W | W | A | A | A | A | A | A | W | W | W | A | W | A | W | A | W | A | A | A | W | A | NA | W | W | W | A | A | A | W | W | W | A | A | W | W | W | W | W | A | W | A | W | A | A | W | W | W | A | A | W | W | A | W | A | W | A | W | W | A |
| ChrV | 17.3 | W | A | A | W | A | W | A | W | A | A | A | A | A | A | NA | A | W | A | W | NA | A | A | A | W | W | W | NA | W | NA | W | A | A | W | W | A | A | A | A | A | A | W | W | W | A | NA | A | W | A | W | A | A | A | W | A | W | W | NA | W | A | A | A | W | W | W | A | NA | W | W | NA | W | NA | A | NA | A | W | A | A | W | W | W | A | A | W | W | A | W | A | W | A | W | W | A |
| ChrV | 17.4 | W | A | A | W | A | W | A | W | A | A | A | A | A | A | W | A | W | A | W | W | A | A | A | W | W | W | A | W | W | W | A | A | W | W | A | A | A | A | A | A | W | W | W | A | W | A | W | A | W | A | A | A | W | A | W | W | W | W | A | A | A | W | W | W | A | A | W | W | W | W | W | A | W | A | W | A | A | W | W | W | A | A | W | W | A | W | A | W | A | W | W | A |
| ChrV | 17.5 | W | A | A | W | A | W | A | W | A | A | A | A | A | A | NA | A | W | A | W | W | A | A | A | W | W | W | A | W | W | W | A | A | W | W | A | A | A | A | A | A | W | W | W | A | W | A | W | A | W | A | A | A | W | A | NA | W | W | W | A | A | A | W | W | W | A | A | W | W | W | W | W | A | W | A | W | A | A | W | W | W | A | A | W | W | A | W | A | W | A | W | W | A |
| ChrV | 17.6 | W | A | A | W | A | W | A | W | A | A | A | A | A | A | W | A | W | A | W | W | A | A | A | W | W | W | A | W | W | W | A | A | W | W | A | A | A | A | A | A | W | W | W | A | W | A | W | A | W | A | A | A | W | A | W | W | W | W | A | A | A | W | W | W | A | A | W | W | W | NA | NA | A | W | A | W | A | A | W | W | W | A | A | W | W | A | W | A | W | A | W | W | A |
| ChrV | 17.7 | W | A | A | W | A | W | A | W | A | A | A | A | A | A | W | A | W | A | W | W | A | A | A | W | W | W | A | W | W | W | A | A | W | NA | A | A | A | A | A | A | W | W | W | A | W | A | W | A | W | A | A | A | W | A | W | W | W | W | A | A | A | W | W | W | A | A | W | W | W | W | NA | A | NA | A | W | A | A | W | W | W | A | A | W | W | A | W | A | W | A | W | W | A |
| ChrV | 17.8 | W | A | A | W | A | W | A | W | A | A | A | A | NA | A | W | A | W | A | W | W | NA | A | A | W | W | W | A | W | W | W | NA | A | W | W | A | A | A | A | A | A | W | W | W | A | W | NA | W | A | W | A | A | A | W | A | W | W | W | W | A | A | NA | W | W | W | A | A | W | W | W | W | W | A | W | A | W | A | A | W | W | W | A | NA | W | W | A | W | NA | W | A | W | W | NA |
| ChrV | 17.9 | W | A | A | W | A | W | A | W | A | A | A | A | A | A | W | A | W | A | W | W | A | A | A | W | W | W | A | W | W | W | A | A | W | W | A | A | A | A | A | A | W | W | W | A | W | A | W | A | W | A | A | A | W | A | W | W | W | W | A | A | A | W | W | W | A | A | W | W | W | W | W | A | W | A | W | A | A | W | W | W | A | A | W | W | A | W | A | W | A | W | W | A |
| ChrV | 18 | W | A | A | W | A | W | A | NA | A | A | A | A | A | A | NA | A | W | A | W | NA | A | A | A | W | W | W | A | W | W | W | A | A | W | NA | A | A | A | A | A | A | W | W | W | A | W | A | W | A | W | A | A | A | W | A | W | W | NA | W | A | A | A | W | W | W | A | A | W | W | W | NA | NA | A | W | A | W | A | A | W | W | W | A | A | W | W | A | W | A | W | A | W | W | A |
| ChrV | 18.1 | W | A | A | W | A | W | A | W | A | A | A | A | A | A | W | A | W | A | W | W | A | A | A | W | W | W | A | W | W | W | A | A | W | W | A | A | A | A | A | A | W | W | W | A | W | A | W | A | W | A | A | A | W | A | W | W | W | W | A | A | A | W | W | W | A | A | W | W | W | W | W | A | W | A | W | A | NA | W | W | W | A | A | W | W | A | W | A | W | A | W | W | A |
| ChrV | 18.2 | W | A | A | W | A | W | A | W | A | A | A | A | A | A | NA | A | W | A | W | NA | NA | A | A | W | W | NA | A | W | W | W | A | A | W | NA | A | A | A | A | A | A | W | W | W | A | NA | A | W | A | W | A | A | A | W | A | W | W | NA | W | A | A | A | W | W | W | A | A | W | W | W | NA | W | A | W | A | W | A | W | W | NA | W | A | A | W | W | A | W | A | W | A | W | W | A |
| ChrV | 18.3 | W | A | A | W | A | W | A | W | A | A | A | A | A | A | W | A | W | A | W | NA | A | A | A | W | W | W | A | W | W | W | A | A | W | NA | A | A | A | A | A | A | W | W | W | A | W | A | W | A | W | A | A | A | W | A | NA | W | W | W | A | A | A | W | W | W | A | A | W | W | W | W | NA | A | W | A | W | A | W | W | W | W | A | A | W | W | A | W | A | W | A | W | W | A |
| ChrV | 18.4 | W | NA | A | W | A | W | A | NA | A | A | A | A | A | A | W | A | W | A | W | NA | A | A | A | W | W | W | A | W | W | W | A | A | W | W | A | A | A | A | A | A | W | W | W | A | W | A | W | A | W | A | NA | A | W | A | W | W | NA | W | A | A | A | W | W | W | A | A | W | W | W | W | NA | A | W | A | W | A | W | W | W | W | A | A | W | W | A | W | A | W | A | W | W | A |
| ChrV | 18.5 | W | A | A | W | A | W | A | W | A | A | A | A | A | A | W | A | W | A | W | W | A | A | A | W | W | W | A | NA | W | W | A | A | W | W | A | A | A | A | A | A | W | W | W | A | W | A | W | A | W | A | A | A | W | A | W | W | W | W | A | A | A | W | W | W | A | A | W | W | W | W | W | A | W | A | W | A | W | W | W | W | A | A | W | W | A | W | A | W | A | W | W | A |
| ChrV | 18.6 | W | A | A | W | A | W | A | W | A | A | A | A | A | A | W | A | W | A | W | W | A | A | A | W | W | W | A | W | W | W | A | A | W | NA | A | A | A | A | A | A | W | W | W | A | W | A | W | A | W | A | A | A | W | A | W | W | W | W | A | A | A | W | W | W | A | A | W | W | W | W | W | A | W | A | W | A | W | W | W | W | A | A | W | W | A | W | A | W | A | W | W | A |
| ChrV | 18.7 | W | A | A | W | A | W | A | W | A | A | A | A | A | A | W | A | W | A | W | W | A | A | A | W | W | W | NA | W | W | W | A | A | W | W | A | A | A | A | A | A | NA | W | W | A | W | A | W | A | W | A | A | A | W | A | W | W | W | W | A | A | A | W | W | W | A | A | W | W | W | W | W | A | W | A | W | A | W | W | W | W | A | A | W | W | A | W | A | W | A | W | W | A |
| ChrV | 18.8 | W | A | A | W | A | W | A | W | A | A | A | A | A | A | NA | A | W | A | W | W | A | A | A | W | W | NA | W | W | NA | W | A | A | W | NA | A | A | A | A | A | A | W | W | NA | A | W | A | W | A | W | A | A | A | W | A | W | NA | W | W | A | A | A | W | NA | W | A | A | W | W | NA | NA | NA | NA | NA | A | W | A | W | W | W | W | A | A | W | W | A | W | A | NA | A | W | W | A |
| ChrV | 18.9 | W | A | A | W | A | W | A | W | A | A | A | A | A | A | W | A | W | A | W | W | A | A | A | W | W | W | W | W | W | W | A | A | W | NA | A | A | A | A | A | A | W | W | W | A | W | A | W | A | W | A | A | A | W | A | W | W | W | W | A | A | A | W | W | W | A | A | W | W | W | W | A | W | W | A | W | A | W | W | W | W | A | A | W | W | A | W | A | W | A | W | W | A |
| ChrV | 19 | W | A | A | W | A | W | A | NA | A | A | A | A | A | A | W | A | W | A | W | W | A | A | A | W | W | W | W | W | W | W | A | A | W | W | A | A | A | A | A | A | W | W | W | A | W | A | W | A | W | A | A | A | W | A | NA | W | W | W | A | A | A | W | W | W | A | A | W | W | W | W | A | W | W | A | W | A | W | W | W | W | A | A | W | W | A | W | A | W | A | W | W | A |
| ChrV | 19.1 | W | A | A | W | A | W | A | W | A | A | A | A | A | A | NA | A | W | A | W | W | A | A | A | W | W | W | NA | W | W | W | A | A | W | W | A | A | A | A | A | A | W | W | W | A | W | A | W | A | W | A | A | A | W | A | A | W | W | W | A | A | A | W | W | W | A | A | W | W | W | W | A | W | W | A | W | A | W | W | W | W | A | A | W | W | A | W | A | W | A | W | W | A |
| ChrV | 19.2 | W | A | A | W | A | W | A | NA | A | A | A | A | A | A | NA | A | W | A | W | NA | A | A | A | W | W | W | W | W | W | W | A | A | W | NA | A | A | A | A | A | A | W | W | W | A | W | A | W | A | W | A | A | A | W | A | A | W | W | W | A | A | A | W | W | W | A | A | W | W | W | W | NA | W | W | A | W | A | W | W | W | W | A | A | NA | W | A | W | A | W | A | W | W | A |
| ChrV | 19.3 | W | A | A | W | A | W | A | A | A | A | A | A | A | A | NA | A | W | A | W | W | A | A | A | W | W | NA | W | W | W | W | A | A | W | NA | A | A | A | A | A | A | W | W | W | A | W | A | W | A | W | A | A | A | W | A | A | NA | NA | W | A | A | A | W | W | NA | A | A | W | W | W | NA | A | W | NA | A | W | A | W | W | W | W | A | NA | A | W | A | W | A | W | A | W | W | A |
| ChrV | 19.4 | W | A | A | W | A | W | A | A | A | A | A | A | A | A | NA | A | W | A | W | W | A | A | A | W | W | W | W | W | W | W | A | A | W | W | A | A | A | A | A | A | W | W | W | A | W | A | W | A | W | A | A | A | W | A | A | W | W | W | A | A | A | W | W | W | A | A | W | W | W | W | A | W | W | A | W | A | W | W | W | W | A | A | A | W | A | W | A | W | A | W | W | A |
| ChrV | 19.5 | W | A | A | W | A | W | A | A | NA | A | A | A | A | A | W | A | W | A | W | W | A | A | A | W | W | W | NA | W | W | W | A | A | W | W | A | A | A | A | A | A | W | W | W | A | W | A | W | A | W | A | A | A | W | A | A | W | NA | W | A | A | A | W | W | W | A | A | W | W | W | W | A | W | W | A | W | A | W | W | NA | W | A | A | A | W | A | W | A | W | A | W | W | A |
| ChrV | 19.6 | W | A | A | W | A | W | A | A | W | A | A | A | A | A | W | A | W | A | W | W | A | A | A | W | W | W | W | W | W | W | A | A | W | W | A | A | A | A | A | A | W | W | W | A | W | A | W | A | W | A | A | NA | W | A | A | W | W | W | A | A | A | W | W | W | A | A | W | W | W | W | A | W | W | A | W | A | W | W | NA | W | A | A | A | W | A | W | A | W | A | W | W | A |
| ChrV | 19.7 | W | A | A | W | A | W | A | A | W | A | A | A | A | NA | W | A | W | A | W | W | A | A | A | W | W | W | W | W | W | W | A | A | W | W | A | A | A | A | A | A | W | W | W | A | W | A | W | A | W | A | A | A | W | A | A | W | W | W | A | A | A | W | W | NA | A | A | W | W | W | W | A | W | NA | A | W | A | W | W | NA | W | A | A | A | W | A | W | A | W | A | W | W | A |
| ChrV | 19.9 | W | A | A | W | A | W | A | A | W | A | A | A | A | A | W | A | NA | A | W | W | A | A | A | W | W | NA | W | W | W | W | A | A | W | W | A | A | A | A | A | A | W | W | W | A | W | A | W | A | W | A | A | A | W | A | A | W | W | W | A | A | A | W | W | W | A | A | NA | W | W | W | A | W | W | A | W | A | W | W | W | W | A | A | A | W | A | W | A | W | A | W | W | A |
| ChrV | 20 | W | A | NA | W | A | W | A | A | W | A | A | A | A | A | W | A | W | A | W | W | A | A | A | W | W | W | W | W | W | W | A | A | W | W | A | A | A | A | A | A | W | W | W | A | W | A | W | A | W | A | A | NA | W | A | A | W | W | W | A | NA | A | W | W | W | NA | A | W | W | W | W | A | W | W | A | W | A | W | W | W | W | A | A | A | W | A | W | NA | W | A | W | W | A |
| ChrV | 20.1 | W | A | A | W | A | W | A | A | W | A | A | A | A | A | W | A | W | A | W | W | A | A | A | W | W | W | NA | W | W | W | A | A | W | W | A | A | A | A | A | A | W | W | W | A | W | A | W | A | W | A | A | A | W | A | A | W | W | W | A | A | A | W | W | W | A | A | W | W | W | W | A | W | W | A | W | A | W | W | W | W | A | A | A | W | A | W | A | W | A | W | W | A |
| ChrV | 20.2 | W | A | A | W | A | W | A | A | W | A | A | A | A | A | W | A | W | A | W | W | A | A | A | W | W | W | NA | W | W | W | A | A | W | W | A | A | A | A | A | A | W | W | W | A | W | A | W | A | W | A | A | NA | W | A | A | W | W | W | A | A | A | W | W | W | A | A | W | W | W | W | A | W | NA | A | W | A | W | W | W | W | A | A | A | W | A | W | A | W | A | W | W | A |
| ChrV | 20.3 | W | A | A | W | A | W | A | A | W | A | A | A | A | A | W | A | W | A | W | NA | NA | A | A | W | W | W | W | W | W | W | A | A | W | W | A | A | A | A | A | A | W | A | W | A | W | A | W | A | W | A | A | W | W | A | A | W | W | W | A | A | A | W | W | W | A | A | W | W | W | W | A | W | W | A | W | A | W | W | W | W | A | A | A | W | A | W | A | W | A | W | W | A |
| ChrV | 20.4 | NA | A | A | NA | A | W | A | A | NA | A | A | A | A | A | NA | A | W | A | W | W | A | A | A | W | W | NA | NA | NA | NA | W | A | A | W | W | A | A | A | A | A | A | NA | A | W | A | W | A | NA | A | W | A | A | W | W | A | A | NA | NA | W | A | A | A | W | NA | NA | A | NA | W | W | NA | NA | A | NA | NA | A | W | A | W | NA | NA | W | A | A | A | W | A | W | A | W | A | W | W | A |
| ChrV | 20.5 | W | W | A | W | A | W | A | A | NA | A | W | A | A | A | W | A | W | A | W | W | A | A | A | W | W | W | NA | W | W | W | A | A | W | W | W | W | A | A | A | A | W | A | W | A | W | A | W | A | W | A | A | W | W | A | A | W | W | W | A | A | A | W | W | W | A | A | W | W | W | W | A | W | W | A | W | A | W | W | W | W | A | A | A | W | A | W | A | W | A | W | W | A |
| ChrV | 20.6 | W | W | A | NA | A | W | A | NA | W | A | W | A | A | A | W | A | W | A | W | W | A | A | A | W | W | W | W | W | W | W | A | A | W | W | W | W | A | A | A | A | W | A | W | A | W | A | W | A | W | A | A | W | W | A | A | W | NA | W | A | A | A | W | W | W | A | A | W | W | W | W | A | W | W | A | W | A | W | A | W | W | A | A | NA | W | A | W | A | W | A | W | W | A |
| ChrV | 20.7 | W | W | A | W | A | W | A | A | W | A | W | A | A | A | W | A | W | A | W | W | NA | A | A | W | W | W | W | W | W | W | A | A | W | W | W | W | A | A | A | A | W | A | W | A | W | A | W | A | W | NA | A | W | W | A | A | W | W | W | A | A | A | W | W | W | A | A | W | W | W | W | A | W | W | A | W | A | W | A | W | W | A | A | A | W | A | W | A | W | A | W | W | A |
| ChrV | 20.8 | W | W | A | W | A | W | A | A | W | A | W | NA | A | A | W | A | W | A | W | W | A | A | A | W | W | W | W | W | W | W | A | A | W | W | W | W | NA | A | A | A | W | A | W | A | W | A | W | A | W | A | A | W | W | A | A | W | W | W | NA | A | A | W | W | W | NA | A | W | W | W | W | A | W | W | A | W | A | W | A | W | W | A | A | A | W | A | W | NA | W | A | W | W | NA |
| ChrV | 20.9 | W | W | A | W | A | W | A | A | W | A | W | A | A | A | W | A | W | A | W | W | A | A | A | W | W | W | W | W | W | W | A | A | W | W | W | W | A | A | A | A | W | A | W | A | W | A | W | A | W | A | A | W | W | A | A | W | W | W | A | A | A | W | W | W | A | A | W | W | W | W | A | W | W | A | W | A | W | A | W | W | A | A | A | W | A | W | A | W | A | W | W | A |
| ChrV | 21 | W | NA | A | W | A | W | A | A | W | A | W | A | A | A | NA | A | W | A | W | NA | NA | A | A | W | W | NA | NA | W | NA | W | NA | A | W | W | W | W | A | A | A | A | NA | A | W | A | W | A | W | A | W | A | A | W | W | A | NA | NA | NA | W | A | A | A | W | NA | NA | A | A | W | W | NA | NA | A | W | W | A | W | A | W | A | NA | W | A | A | A | NA | A | W | A | W | A | NA | W | A |
| ChrV | 21.1 | NA | NA | A | W | NA | W | A | NA | W | A | W | A | A | A | NA | NA | W | A | W | NA | NA | A | A | W | W | NA | NA | W | W | W | W | A | W | NA | W | W | A | A | A | A | NA | A | W | NA | NA | A | W | A | W | NA | A | W | W | A | A | NA | NA | W | A | A | A | W | NA | NA | A | NA | W | W | NA | W | A | NA | NA | A | W | A | W | A | W | W | A | NA | A | W | A | W | A | W | A | NA | W | A |
| ChrV | 21.2 | W | W | A | W | A | W | NA | A | W | A | W | A | A | A | W | A | W | A | W | W | A | A | A | W | W | W | W | W | W | W | W | A | W | NA | W | W | A | A | A | A | W | A | W | A | W | A | W | A | W | A | A | W | W | A | A | W | W | W | A | A | A | W | W | W | NA | A | W | W | W | W | A | W | W | A | W | A | W | A | W | W | A | A | A | W | A | W | A | W | A | W | W | A |
| ChrV | 21.3 | W | W | A | W | A | W | A | A | W | A | W | A | A | A | W | A | W | A | W | NA | A | A | A | W | W | W | W | W | W | W | W | A | W | W | W | W | A | A | A | A | W | A | W | A | W | A | W | A | W | A | A | W | W | A | A | W | W | W | A | A | A | W | W | W | W | A | W | W | W | W | A | W | W | A | W | A | W | A | W | W | A | A | A | W | A | W | A | W | A | W | W | A |
| ChrV | 21.4 | W | W | A | W | A | W | A | A | W | A | W | A | A | A | W | A | W | A | W | W | A | A | A | W | W | W | W | W | W | W | W | A | W | W | W | W | A | A | A | A | W | A | W | A | W | A | W | A | W | A | A | W | W | A | A | W | W | W | A | A | A | W | W | W | W | A | W | W | W | W | NA | W | W | A | W | A | W | A | W | W | A | A | A | W | A | W | A | W | A | W | W | A |
| ChrV | 21.5 | W | W | A | W | A | W | A | A | W | A | W | A | A | A | W | A | W | A | W | W | A | A | A | W | W | W | W | W | W | W | W | A | W | W | W | W | A | A | A | A | W | A | W | A | W | A | W | A | W | A | A | W | W | W | A | W | NA | W | A | A | A | W | W | W | W | A | W | W | W | NA | A | W | W | W | NA | A | W | A | W | W | A | A | A | W | A | W | A | W | A | W | W | A |
| ChrV | 21.6 | W | W | A | W | A | W | A | A | W | A | W | A | A | A | W | A | W | A | W | W | A | A | A | W | W | W | W | W | W | W | W | A | W | W | W | W | A | A | A | A | W | A | W | A | W | A | W | A | W | A | A | W | W | W | A | W | A | W | A | A | A | W | W | W | W | A | W | W | W | W | A | W | W | W | A | A | W | A | W | W | A | A | A | W | A | W | A | W | A | W | W | A |

| | | | | | | | | | | | | | | | | | | | | | | | | | | | | | | | | | | | | | | | | | | | | | | | | | | | | | | | | | | | | | | | | | | | | | | | | | | | | | | | | | | | | | | | | | | | | |
|---|---|---|---|---|---|---|---|---|---|---|---|---|---|---|---|---|---|---|---|---|---|---|---|---|---|---|---|---|---|---|---|---|---|---|---|---|---|---|---|---|---|---|---|---|---|---|---|---|---|---|---|---|---|---|---|---|---|---|---|---|---|---|---|---|---|---|---|---|---|---|---|---|---|---|---|---|---|---|---|---|---|---|---|---|---|---|---|---|---|---|---|---|---|
| ChrV | 21.8 | W | W | A | W | A | W | A | NA | W | A | W | NA | A | A | W | A | W | A | W | NA | A | W | A | W | W | W | W | W | W | W | W | A | W | W | W | W | A | NA | A | A | W | NA | W | A | W | A | W | A | W | A | A | W | W | W | NA | W | NA | W | A | A | A | W | W | W | W | A | W | W | W | W | A | W | W | W | NA | A | W | NA | W | W | A | A | NA | W | A | W | NA | W | A | W | W | A |
| ChrV | 21.9 | W | W | A | W | A | W | A | A | W | A | W | A | A | A | W | A | W | A | W | W | A | W | A | W | W | W | NA | W | W | W | W | A | W | W | W | W | A | A | A | A | W | A | W | A | W | A | W | A | W | A | A | W | W | W | A | W | A | W | A | A | A | W | W | W | W | A | W | W | NA | W | A | W | NA | W | A | A | W | A | NA | W | A | A | A | W | A | W | A | W | A | NA | W | A |
| ChrV | 22 | W | NA | A | W | W | W | A | A | W | NA | W | A | A | A | W | NA | W | A | W | NA | NA | W | A | NA | W | NA | NA | NA | W | W | NA | A | W | W | W | W | A | A | A | A | NA | NA | NA | A | W | A | NA | A | W | A | NA | W | W | NA | A | W | NA | W | A | A | A | W | NA | W | W | A | W | W | NA | W | A | W | W | W | A | A | W | A | NA | W | A | A | A | W | NA | W | A | W | A | W | W | A |
| ChrV | 22.1 | W | W | A | W | W | W | A | A | W | A | W | A | A | A | W | A | W | A | W | W | A | W | A | A | W | W | W | W | W | W | W | A | W | W | W | W | A | A | A | A | W | A | W | A | W | A | W | A | W | A | A | W | W | W | A | W | A | W | A | A | A | W | W | W | W | A | W | W | W | W | A | W | W | W | A | A | W | A | W | W | A | A | A | W | A | W | A | W | A | W | W | A |
| ChrV | 22.2 | W | W | A | W | W | W | A | A | W | A | W | A | A | A | W | A | W | A | W | W | A | W | A | A | W | NA | NA | W | NA | W | W | A | W | W | W | W | A | A | A | A | W | A | W | A | W | A | W | A | W | A | A | W | W | W | A | W | A | W | A | A | A | W | W | W | W | A | W | W | W | W | A | W | NA | W | A | A | W | A | W | W | A | A | A | NA | A | W | A | W | A | W | W | A |
| ChrV | 22.3 | W | W | A | W | W | W | A | A | W | A | W | A | A | A | W | A | W | A | W | W | A | W | A | A | W | W | NA | NA | W | W | W | A | W | W | W | W | A | A | A | A | W | A | W | A | W | A | W | A | W | A | A | W | W | W | A | W | A | W | A | A | A | W | W | W | W | A | W | NA | W | NA | A | W | NA | W | A | A | W | A | W | W | A | A | A | NA | A | W | A | W | A | W | W | A |
| ChrV | 22.4 | W | W | A | W | W | W | A | A | W | A | W | A | A | A | W | A | W | A | W | W | A | W | A | A | W | W | W | W | W | W | W | A | W | W | W | W | A | A | A | A | A | A | W | A | W | A | W | A | W | A | A | W | W | W | A | W | A | W | A | A | A | W | W | W | W | A | W | W | W | W | A | W | W | W | A | A | W | A | W | W | A | A | A | W | A | W | A | W | A | W | W | A |
| ChrV | 22.5 | W | W | A | W | W | W | A | A | W | A | W | A | A | A | NA | A | W | A | W | W | A | W | A | A | W | W | W | W | W | W | W | A | W | W | W | W | A | A | A | A | A | A | W | A | A | A | W | A | W | A | A | W | W | W | A | W | A | W | A | A | A | W | W | W | W | A | W | NA | W | W | A | W | W | W | A | A | W | A | W | W | A | A | A | W | A | W | A | W | A | W | W | A |
| ChrV | 22.6 | W | NA | NA | W | W | NA | NA | A | W | A | NA | A | NA | A | W | A | W | A | W | NA | A | NA | A | A | NA | W | W | W | W | W | W | A | W | W | NA | NA | A | A | A | A | A | A | W | A | A | NA | W | A | W | A | A | W | W | W | A | W | A | W | A | NA | A | NA | W | W | NA | A | NA | W | W | NA | A | W | W | W | NA | A | W | A | W | W | A | A | A | W | A | W | NA | W | A | W | W | A |
| ChrV | 22.7 | W | W | A | W | W | W | A | A | W | A | W | A | A | A | W | A | W | A | W | W | A | W | A | A | W | W | W | NA | W | W | W | A | W | NA | W | W | A | A | A | A | A | A | W | A | A | A | W | A | W | A | A | W | W | W | A | W | A | W | A | A | A | W | W | W | W | A | W | W | W | NA | A | W | W | W | A | A | W | A | W | W | A | A | A | W | A | W | A | W | A | W | W | A |
| ChrV | 22.8 | W | W | A | W | W | W | A | A | W | A | W | A | A | A | W | A | W | A | W | W | A | W | A | A | W | W | W | W | W | W | W | A | W | W | W | W | A | A | A | A | A | A | W | A | A | A | W | A | W | A | A | W | W | W | A | W | A | W | A | A | A | W | W | W | W | A | W | W | W | W | A | W | W | W | A | A | W | A | W | W | A | A | A | W | A | W | A | W | A | W | W | A |
| ChrV | 22.9 | W | W | A | W | W | W | A | A | W | A | W | A | A | A | NA | A | W | A | W | W | NA | W | A | A | W | W | W | W | W | W | W | A | W | W | W | W | A | A | A | A | A | A | W | A | A | A | W | A | W | A | A | W | W | W | A | W | A | W | A | A | A | W | W | W | W | A | W | W | W | W | A | W | W | W | A | A | W | A | W | W | A | A | A | W | A | W | A | W | A | W | W | A |
| ChrV | 23 | W | W | A | W | W | W | A | A | W | A | W | A | A | A | W | A | W | A | W | NA | A | W | A | A | W | NA | W | NA | W | W | W | A | W | W | W | W | A | A | A | A | A | A | W | A | A | A | W | A | W | A | A | W | W | W | A | W | A | W | A | A | A | W | W | NA | W | A | W | NA | NA | NA | A | W | NA | W | A | A | W | A | W | W | A | A | A | W | A | W | A | W | A | W | W | A |
| ChrV | 23.1 | W | W | NA | W | W | W | A | A | W | NA | W | A | A | A | NA | A | W | A | W | W | A | NA | A | A | W | W | W | W | W | W | W | A | W | W | W | W | A | A | A | A | A | A | W | A | A | A | W | A | W | A | A | W | W | W | A | W | A | W | A | A | A | W | W | W | W | NA | W | W | W | NA | NA | W | NA | W | A | A | W | A | W | W | A | A | NA | NA | A | W | A | W | A | W | W | A |
| ChrV | 23.2 | W | W | A | W | W | W | A | A | W | A | W | A | A | A | W | A | W | A | W | W | A | W | A | A | W | W | W | W | W | W | W | A | W | W | W | W | A | A | A | A | A | A | W | A | A | A | W | A | W | NA | A | W | W | W | A | W | A | W | A | A | A | W | W | W | W | A | W | W | W | W | A | W | W | W | A | A | W | A | W | W | A | A | A | W | A | W | A | W | A | NA | W | A |
| ChrV | 23.3 | W | W | A | W | W | W | A | A | W | A | W | A | A | A | NA | A | W | A | W | W | A | NA | A | A | W | W | NA | W | W | W | W | A | W | W | W | W | A | A | A | A | A | A | W | A | A | A | W | A | W | A | A | W | W | W | A | W | A | W | A | A | A | W | W | W | W | A | W | NA | W | W | A | W | W | W | A | A | W | A | W | W | A | A | A | W | A | W | A | W | A | W | W | A |
| ChrV | 23.4 | W | W | A | W | W | W | A | A | W | A | W | A | A | A | W | A | W | A | W | W | A | NA | NA | A | W | W | W | W | W | W | W | A | W | W | W | W | A | A | A | A | A | A | W | A | A | A | W | A | W | A | A | W | W | W | A | W | A | W | A | A | A | NA | W | W | NA | A | W | W | W | W | A | W | W | W | A | A | W | A | W | W | A | A | A | W | A | W | A | W | A | W | W | A |
| ChrV | 23.5 | W | W | A | W | W | W | A | A | W | A | W | A | A | A | W | A | W | A | W | W | A | W | A | A | W | W | W | W | W | W | W | A | W | W | W | W | A | A | A | A | A | A | W | A | A | A | W | A | W | A | A | W | W | W | A | W | A | W | A | A | A | W | W | W | W | A | W | W | W | NA | A | W | W | W | A | A | W | A | W | W | A | A | A | W | A | W | A | W | A | W | W | A |
| ChrV | 23.6 | W | NA | A | W | W | W | A | A | W | A | W | A | A | A | W | A | W | A | W | W | A | NA | A | A | W | W | NA | W | W | W | W | A | W | W | W | W | A | A | A | A | A | A | NA | A | A | A | W | A | W | A | A | W | W | NA | A | W | A | W | A | A | A | W | W | W | W | A | W | W | W | NA | A | W | W | W | A | A | W | A | W | W | A | A | A | W | A | W | A | W | A | NA | W | A |
| ChrV | 23.7 | W | W | A | W | W | W | A | A | W | A | W | A | A | A | W | A | W | A | W | W | A | W | A | A | W | W | W | W | W | W | W | A | W | NA | W | W | A | A | A | A | A | A | W | A | A | A | W | A | W | A | A | W | W | W | A | W | A | W | A | A | A | W | W | W | W | A | W | W | W | W | A | W | W | W | A | A | W | A | W | W | A | A | A | W | A | W | A | W | A | W | W | A |
| ChrV | 23.8 | W | W | A | W | W | W | A | A | W | A | W | A | A | A | NA | A | W | A | W | W | A | W | A | A | W | W | W | W | W | W | W | A | W | W | W | W | A | A | A | A | A | A | W | A | A | A | W | A | W | A | A | W | W | W | A | W | A | W | A | A | A | W | W | W | W | A | W | W | W | W | A | W | W | W | A | A | W | A | W | W | A | A | A | W | A | W | A | W | A | W | W | A |
| ChrX | 0 | W | W | W | W | W | W | W | W | W | W | W | W | W | W | W | W | W | W | W | W | W | W | W | W | W | W | W | W | W | W | W | W | W | NA | W | W | W | W | W | W | W | W | W | W | W | NA | W | W | W | W | W | W | W | W | W | W | W | W | W | W | W | W | W | W | W | W | W | W | W | W | W | W | W | W | W | W | W | W | W | W | W | W | W | W | W | W | W | W | W | W | W | W |
| ChrX | 0.1 | W | W | W | W | W | W | W | W | W | W | W | W | W | W | W | W | W | W | W | W | NA | W | W | W | W | W | NA | W | W | W | W | W | W | W | W | W | W | W | W | W | W | W | W | W | W | W | W | W | W | W | W | W | W | W | W | W | W | W | W | W | W | W | W | W | NA | NA | W | W | W | W | W | W | W | W | W | W | W | W | W | W | W | W | W | W | W | W | W | W | W | W | W | W |
| ChrX | 0.2 | W | W | W | W | W | W | W | W | W | W | W | W | W | W | W | W | W | W | W | W | W | W | NA | W | W | W | W | W | W | W | W | W | W | W | W | W | W | W | W | W | W | W | W | W | W | W | W | W | W | W | W | W | W | W | W | W | W | W | W | W | W | W | W | W | W | W | W | W | W | W | W | W | W | W | W | W | W | W | W | W | W | W | W | W | W | W | W | W | W | W | W | W |
| ChrX | 0.3 | W | W | NA | W | W | W | NA | W | W | W | W | W | W | W | W | W | W | W | W | W | NA | W | W | W | W | NA | W | W | W | W | W | W | W | W | W | W | W | W | W | W | W | W | W | W | W | W | W | W | W | W | W | W | W | W | W | W | W | W | W | W | W | W | W | W | W | W | W | NA | W | W | NA | W | W | W | W | W | NA | W | W | W | W | W | W | W | W | W | W | W | W | W | W | W |
| ChrX | 0.4 | W | W | W | W | W | W | W | W | W | W | W | W | W | W | W | W | W | W | W | W | W | W | W | W | W | W | W | W | W | W | W | W | W | W | W | W | W | W | W | W | W | W | W | W | W | W | W | NA | W | W | W | W | W | W | W | W | W | W | W | W | W | W | W | W | W | W | W | W | W | W | W | W | W | W | W | W | W | W | W | W | W | W | W | W | W | W | W | W | W | W | W | W |
| ChrX | 0.5 | W | W | W | W | W | W | W | W | W | W | W | W | W | W | W | W | W | W | W | W | W | W | W | W | W | W | W | W | W | W | W | W | W | W | W | W | W | W | W | W | W | W | W | W | W | W | W | W | W | W | W | W | W | W | W | W | W | W | W | W | W | W | W | W | W | W | W | W | W | W | W | W | W | W | W | W | W | W | W | W | W | W | W | W | W | W | W | W | W | W | W | W |
| ChrX | 0.7 | W | W | W | W | W | W | W | W | W | W | W | W | W | W | W | W | W | W | W | W | W | W | W | W | W | W | W | W | W | W | W | W | W | W | W | W | W | W | W | W | W | W | W | W | W | W | W | W | W | W | W | NA | W | W | W | W | W | W | W | W | W | W | W | W | W | W | W | W | W | W | W | W | W | W | W | W | W | W | W | W | W | W | W | W | W | W | W | W | W | W | W | W |
| ChrX | 0.8 | W | W | W | W | W | W | W | W | W | W | W | W | W | W | W | W | W | W | W | W | W | W | W | W | W | W | W | W | W | W | W | W | W | W | W | W | W | W | W | W | W | W | W | W | W | W | W | W | W | W | W | W | W | W | W | W | W | W | W | W | W | W | W | W | W | W | W | W | W | W | W | W | W | W | W | W | W | W | W | W | W | W | W | W | W | W | W | W | W | W | W | W |
| ChrX | 0.9 | W | W | W | W | W | W | W | W | W | W | W | W | W | W | W | W | W | W | W | W | W | W | W | W | W | W | NA | W | W | W | W | W | W | W | W | W | W | W | W | W | W | W | W | W | W | W | W | W | W | W | W | W | W | W | W | W | W | W | W | W | W | W | W | W | W | W | NA | W | W | W | W | W | W | W | W | W | W | W | W | W | W | W | W | W | W | W | W | W | W | W | W | W |
| ChrX | 1 | W | W | W | W | W | W | W | W | W | W | W | W | W | W | W | W | W | W | W | W | W | W | W | W | W | W | W | W | W | W | W | W | W | W | W | W | W | W | W | W | W | W | W | W | W | W | W | W | W | W | W | W | W | W | W | W | W | W | W | W | W | W | W | W | W | W | W | W | W | W | W | W | W | W | W | W | W | W | W | W | W | W | W | W | W | W | W | W | W | W | W | W |
| ChrX | 1.1 | W | W | W | W | W | W | W | W | W | W | W | W | W | W | W | W | NA | W | W | W | W | W | W | W | W | W | W | W | W | W | W | W | W | W | W | W | W | W | W | W | W | W | W | W | W | W | W | W | W | NA | W | W | W | W | W | W | W | W | W | W | W | W | W | W | W | W | W | W | W | W | W | W | W | W | W | W | W | W | W | W | W | W | W | W | W | W | W | W | W | W | W | W |
| ChrX | 1.2 | W | W | W | W | W | W | W | W | W | W | W | W | W | W | W | W | W | W | W | W | W | W | W | W | W | W | W | W | W | W | W | W | W | W | W | W | W | W | W | W | W | W | W | W | W | W | W | W | W | W | W | W | W | W | W | W | W | W | W | W | W | W | W | W | W | W | NA | W | W | W | W | W | W | W | W | W | W | W | W | W | W | W | W | W | W | W | W | W | W | W | W | W |
| ChrX | 1.3 | W | W | W | W | W | W | W | W | W | W | W | W | W | W | W | W | W | W | W | W | W | W | W | W | W | W | W | W | W | W | W | W | W | W | W | W | W | W | W | W | W | W | W | W | W | W | W | W | W | W | W | W | W | W | W | W | W | W | W | W | W | W | W | W | W | W | W | W | W | W | W | W | W | W | W | W | W | W | W | W | W | W | W | W | W | W | W | W | W | W | W | W |
| ChrX | 1.4 | W | W | W | W | W | W | W | W | W | W | W | W | W | W | W | W | W | W | W | W | W | W | W | W | W | W | W | W | W | W | W | W | W | W | W | W | W | W | W | W | W | W | W | W | W | W | W | W | W | W | W | W | W | W | W | W | W | W | W | W | W | W | W | W | W | W | W | W | W | W | W | W | W | W | W | W | W | W | W | W | W | W | W | W | W | W | W | W | W | W | W | W |
| ChrX | 1.5 | W | W | NA | W | W | W | W | W | W | W | W | W | W | W | W | W | W | W | W | NA | NA | W | W | W | W | W | W | W | W | W | W | NA | W | W | W | W | W | W | W | W | W | W | W | W | W | W | W | W | W | W | W | W | W | W | W | W | W | W | W | W | W | W | W | W | W | W | NA | NA | W | W | NA | W | W | W | W | W | W | NA | W | W | W | W | W | W | W | W | W | W | W | W | W | W |
| ChrX | 1.6 | W | W | W | W | W | W | W | W | W | W | W | W | W | W | W | W | W | W | W | W | W | W | W | W | W | W | W | W | W | W | W | W | W | W | W | W | W | W | W | W | W | W | W | W | W | W | W | W | W | W | W | W | W | W | W | W | W | W | W | W | W | W | W | W | W | W | W | W | W | W | W | W | W | W | W | W | W | W | W | W | W | W | W | W | W | W | W | W | W | W | W | W |
| ChrX | 1.7 | W | W | W | W | W | W | W | W | W | W | W | W | W | W | W | W | W | W | W | W | W | W | W | W | W | W | W | W | W | W | W | W | W | W | W | W | W | W | W | W | W | W | W | W | W | W | W | NA | W | W | W | W | W | W | NA | W | W | W | W | W | W | W | W | W | W | W | W | W | W | W | W | W | W | W | W | W | W | W | W | W | W | W | W | W | W | W | W | W | W | W | W | W |
| ChrX | 1.8 | W | W | W | W | W | W | W | W | W | W | W | W | W | W | W | W | W | W | W | W | W | W | W | W | W | W | W | W | W | W | W | W | W | W | W | W | W | W | W | W | W | W | W | W | W | NA | W | W | W | W | W | W | W | W | W | W | W | W | W | W | W | W | W | W | W | NA | NA | W | W | W | W | W | W | W | W | W | W | W | W | W | W | NA | W | W | W | W | W | W | W | W | W | W |
| ChrX | 1.9 | W | W | W | W | W | W | W | W | W | W | W | W | W | W | W | W | W | W | W | W | W | W | W | W | W | W | W | W | W | W | W | W | W | W | W | W | W | W | W | W | W | W | W | W | W | W | W | W | W | W | W | W | W | W | W | W | W | W | W | W | W | W | W | W | W | W | W | W | W | W | W | W | W | W | W | W | W | W | W | W | W | W | W | W | W | W | W | W | W | W | W | W |
| ChrX | 2 | W | W | W | W | W | W | W | W | W | W | W | W | W | W | W | W | W | W | W | W | W | W | W | W | W | W | W | W | W | W | W | W | W | W | W | W | W | W | W | W | W | W | W | W | W | W | W | W | W | W | W | W | W | W | W | W | W | W | W | W | W | W | W | W | W | W | W | W | W | W | W | W | W | W | W | W | W | W | W | W | W | W | W | W | W | W | W | W | W | W | W | W |
| ChrX | 2.1 | W | NA | W | W | NA | W | W | W | W | W | W | W | W | W | W | W | W | W | W | W | W | W | W | W | NA | W | NA | W | W | W | W | NA | W | W | W | W | W | NA | W | W | NA | W | W | NA | W | W | W | W | W | W | NA | W | W | W | NA | NA | W | W | W | W | W | W | W | W | NA | NA | NA | W | W | W | W | W | W | W | W | W | W | NA | NA | W | W | W | W | W | W | W | W | W | W | W | W | W |
| ChrX | 2.2 | W | NA | W | W | W | W | W | W | W | W | W | NA | W | W | W | W | W | NA | W | W | W | W | W | W | NA | NA | W | W | NA | W | W | W | NA | W | W | NA | W | W | W | W | W | NA | W | NA | W | NA | W | W | W | W | NA | NA | W | W | NA | NA | W | W | W | W | W | W | NA | W | NA | W | W | W | W | W | W | W | W | W | W | W | W | W | NA | W | NA | NA | W | NA | W | W | W | W | W | W | W | W |
| ChrX | 2.3 | W | W | W | W | W | NA | W | W | W | W | W | W | W | W | W | W | W | W | W | W | W | W | W | W | W | W | W | W | W | W | W | W | W | W | W | W | W | W | W | W | W | W | W | W | W | W | W | W | W | W | W | W | W | W | W | W | W | W | W | W | W | W | W | W | W | W | W | W | W | W | W | W | W | W | W | W | W | W | W | W | W | W | W | W | W | W | W | W | W | W | W | W |
| ChrX | 2.4 | W | W | W | W | W | W | W | W | W | W | W | W | W | W | W | W | W | W | W | W | W | W | W | W | W | W | NA | W | W | W | W | W | W | W | W | W | W | W | W | W | W | W | W | W | W | W | W | W | W | W | W | W | W | W | W | W | W | W | W | W | W | W | W | W | NA | W | W | W | W | W | W | W | W | W | W | W | W | W | W | W | W | W | W | W | W | W | W | W | W | W | W | W |
| ChrX | 2.5 | W | W | W | W | W | W | W | W | W | W | W | W | W | W | W | W | W | W | W | W | W | W | W | W | W | W | W | W | W | W | W | W | W | W | W | W | W | W | W | W | W | W | W | W | W | W | W | W | W | W | W | W | W | W | W | W | W | W | W | W | W | W | W | W | W | W | W | W | W | W | W | W | W | W | W | W | W | W | W | W | W | W | W | W | W | W | W | W | W | W | W | W |
| ChrX | 2.6 | W | W | W | W | W | W | W | W | W | W | W | W | W | W | W | W | W | W | W | W | W | W | W | W | W | W | W | W | W | W | W | W | W | W | W | W | W | W | W | W | W | W | W | W | W | W | W | W | W | W | W | W | W | W | W | W | W | W | W | W | W | W | W | W | W | W | W | W | W | W | W | W | W | W | W | W | W | W | W | W | W | W | W | W | W | W | W | W | W | NA | W | W |
| ChrX | 2.7 | W | W | W | NA | W | W | W | W | W | W | W | W | W | NA | NA | W | NA | W | NA | W | W | W | W | W | W | W | W | NA | W | W | W | NA | W | W | W | W | NA | W | W | W | W | W | W | NA | W | W | W | W | W | W | W | W | NA | W | W | W | W | W | W | NA | W | W | NA | W | NA | NA | W | W | W | W | W | W | W | W | W | W | W | W | NA | W | W | W | W | W | W | W | W | W | W | W | NA | W |
| ChrX | 2.8 | W | W | W | W | W | W | W | W | W | W | W | W | W | W | W | W | W | W | W | W | W | W | W | W | W | W | W | W | W | W | W | W | W | W | W | W | W | W | W | W | W | W | W | W | W | W | W | W | W | W | W | W | W | W | W | W | W | W | W | W | W | W | W | W | W | W | W | W | W | W | W | W | W | W | W | W | W | W | W | W | W | W | W | W | W | W | W | W | W | W | W | W |
| ChrX | 2.9 | W | W | W | W | W | W | W | W | W | W | W | W | W | W | W | W | W | W | W | W | W | W | W | W | W | W | W | W | W | W | W | W | W | W | W | W | W | W | W | W | W | W | W | W | W | W | W | W | W | W | W | W | W | W | W | W | W | W | W | W | W | W | W | W | W | W | W | W | W | W | W | W | W | W | W | W | W | W | W | W | W | W | W | W | W | W | W | W | W | W | W | W |
| ChrX | 3 | W | W | W | W | W | W | W | W | W | W | W | W | W | NA | W | W | NA | W | W | W | W | W | W | W | W | W | W | W | NA | W | W | W | W | W | W | W | W | W | W | W | W | W | W | W | W | W | W | W | W | W | W | W | W | W | NA | W | W | W | W | W | W | W | W | W | NA | W | W | W | W | W | W | W | W | W | W | W | W | NA | W | W | W | W | W | W | W | W | W | W | W | W | W | W |
| ChrX | 3.1 | W | W | W | W | W | W | W | W | W | W | W | W | W | W | W | W | W | W | W | W | W | W | W | W | W | W | W | W | W | W | W | W | W | W | W | W | W | W | W | W | W | W | W | W | W | W | W | W | W | W | W | W | W | W | W | W | W | W | W | W | W | W | W | W | W | W | W | W | W | W | W | W | W | W | W | W | W | W | W | W | W | W | W | W | W | W | W | W | W | W | W | W |
| ChrX | 3.2 | W | W | W | W | W | W | W | W | W | W | W | W | W | W | W | W | NA | W | W | W | W | W | W | W | W | NA | W | W | W | W | W | W | W | W | W | W | W | W | W | W | W | W | W | W | W | W | W | W | W | W | W | W | W | W | W | W | W | W | W | W | W | W | W | W | W | W | W | W | W | W | W | W | W | W | W | W | W | W | W | W | W | W | W | W | W | W | W | W | W | W | W | W |
| ChrX | 3.3 | W | W | W | W | W | W | W | W | W | W | W | W | W | W | W | W | W | W | W | W | W | W | W | W | W | W | W | W | W | W | W | W | W | W | W | W | W | W | W | W | W | W | W | W | W | W | W | W | W | W | W | W | W | W | W | W | W | W | W | W | W | W | W | W | W | W | W | W | W | W | W | W | W | W | W | W | W | W | W | W | W | W | W | W | W | W | W | W | W | W | W | W |
| ChrX | 3.4 | W | W | W | W | W | W | W | W | W | W | W | W | W | W | W | W | W | W | W | W | W | W | W | W | W | W | W | W | W | W | W | W | W | W | W | W | W | W | W | W | W | W | W | W | W | W | W | W | W | W | W | W | W | W | W | W | W | W | W | W | W | W | W | W | W | W | W | W | W | W | W | W | W | W | W | W | W | W | W | W | W | W | W | W | W | W | W | W | W | W | W | W |
| ChrX | 3.5 | W | W | W | W | W | W | W | W | W | W | W | W | W | W | W | W | W | W | W | W | W | W | W | W | W | W | W | W | W | W | W | W | W | W | W | W | W | W | W | W | W | W | W | W | W | W | W | W | W | W | W | W | W | W | W | W | W | W | W | W | W | W | W | W | W | W | W | W | W | W | W | W | W | W | W | W | W | W | W | W | W | W | W | W | W | W | W | W | W | W | W | W |
| ChrX | 3.6 | W | W | W | W | W | W | W | W | W | W | W | W | W | W | W | W | W | W | W | W | W | W | W | W | W | W | NA | W | W | W | W | W | W | W | W | W | W | W | W | W | W | W | W | W | W | W | W | NA | W | W | W | W | W | W | W | W | W | W | W | W | W | W | W | W | NA | W | W | W | W | W | W | W | W | W | W | W | W | W | W | W | W | W | W | W | W | W | W | W | W | W | W | W |
| ChrX | 3.7 | W | W | W | W | W | W | W | W | W | W | W | W | W | W | W | W | W | W | W | W | W | W | W | W | W | W | W | W | W | W | W | W | W | W | W | W | W | W | W | W | W | W | W | W | W | W | W | W | W | W | W | W | W | W | W | W | W | W | W | W | W | W | W | W | W | W | W | W | W | W | W | W | W | W | W | W | W | W | W | W | W | W | W | W | W | W | W | W | W | W | W | W |
| ChrX | 3.8 | W | W | W | W | W | W | W | W | W | W | W | W | W | W | W | W | W | W | W | W | W | W | W | W | W | W | W | W | W | W | W | W | W | W | W | W | W | W | W | W | W | W | W | W | W | W | W | W | W | W | W | W | W | W | NA | W | W | W | W | NA | W | W | W | W | W | W | W | W | W | W | W | W | W | W | W | W | W | W | W | W | W | W | W | W | W | W | W | W | W | W | W | W |
| ChrX | 3.9 | W | W | W | W | W | W | W | W | W | W | W | W | W | W | W | W | W | W | W | W | W | W | W | W | W | W | W | W | W | W | W | W | W | W | W | W | W | W | W | W | W | W | W | W | W | W | W | W | W | W | W | W | W | W | W | W | W | W | W | W | W | W | W | W | W | W | W | W | W | W | W | W | W | W | W | W | W | W | W | W | W | W | W | W | W | W | W | W | W | W | W | W |
| ChrX | 4 | W | W | W | W | W | W | W | W | W | W | W | W | W | W | W | W | W | NA | W | W | W | W | W | W | W | W | W | W | W | W | W | W | W | W | W | W | W | W | W | W | W | W | W | W | W | NA | W | W | W | W | W | NA | W | W | W | NA | W | W | W | W | W | W | W | W | W | W | W | W | W | W | W | W | W | W | W | W | W | NA | W | W | W | W | W | W | W | NA | W | W | W | W | W | W |
| ChrX | 4.1 | W | W | W | W | W | W | W | W | W | W | W | W | W | W | W | W | W | W | W | W | W | W | W | W | W | W | W | W | W | W | W | W | W | W | W | W | W | W | W | W | W | W | W | W | W | W | W | W | W | W | W | W | W | W | W | W | W | W | W | W | W | W | W | W | W | W | W | W | W | W | W | W | W | W | W | W | W | W | W | W | W | W | W | W | W | W | W | W | W | W | W | W |
| ChrX | 4.2 | W | W | W | W | W | W | W | W | W | W | W | W | W | NA | W | W | W | W | W | W | W | W | W | W | W | W | W | W | W | W | W | W | W | NA | W | W | W | W | W | W | W | W | W | W | W | W | W | W | W | W | W | W | W | W | W | W | W | W | W | W | W | W | W | W | NA | W | W | W | W | W | W | W | W | W | W | W | NA | W | W | W | W | NA | W | W | W | W | W | W | W | W | W | W |
| ChrX | 4.3 | W | W | W | W | W | W | W | W | W | W | W | W | W | NA | NA | W | W | W | W | W | W | W | W | W | W | W | W | W | W | W | W | W | W | W | W | W | W | W | W | W | W | W | W | W | W | W | W | W | W | W | W | W | W | W | W | W | W | W | W | W | W | W | W | W | W | NA | W | W | W | W | W | W | W | W | W | W | W | W | W | W | W | W | W | W | W | W | W | W | W | W | W | W |
| ChrX | 4.4 | W | W | W | W | W | W | W | W | W | W | W | W | W | NA | W | W | W | W | W | W | W | W | W | W | W | W | W | W | W | W | W | W | W | W | W | W | W | W | W | W | W | W | W | W | W | NA | W | W | W | W | W | W | W | W | W | W | W | W | W | W | W | W | W | W | W | W | W | W | W | W | W | W | W | W | W | W | W | W | W | W | W | W | W | W | W | W | W | W | W | W | W | W |
| ChrX | 4.5 | W | W | W | W | W | W | W | W | W | W | W | W | W | W | W | W | W | W | W | W | W | W | W | W | W | W | W | W | W | W | W | W | W | W | W | W | W | W | W | W | W | W | W | W | W | W | W | W | W | W | W | W | W | W | W | W | W | W | W | W | W | W | W | W | W | W | W | W | W | W | W | W | W | W | W | W | W | W | W | W | W | W | W | W | W | W | W | W | W | W | W | W |
| ChrX | 4.6 | W | W | W | W | W | W | W | W | W | W | W | W | W | W | W | W | W | W | W | W | W | W | W | W | W | W | W | W | W | W | W | W | W | W | W | W | W | W | W | W | W | W | W | W | W | W | W | W | W | W | W | W | W | W | W | W | W | W | W | W | W | W | W | W | W | W | W | W | W | W | W | W | W | W | W | W | W | W | W | W | W | W | W | W | W | W | W | W | W | W | W | W |
| ChrX | 4.7 | W | W | W | W | W | W | W | W | W | W | W | W | W | W | W | W | W | W | W | W | W | W | W | W | W | W | W | W | W | W | W | W | W | W | W | W | W | W | W | W | W | W | W | W | W | W | W | W | W | W | W | W | W | W | W | W | W | W | W | W | W | W | W | W | W | W | W | W | W | W | W | W | W | W | W | W | W | W | W | W | W | W | W | W | W | W | W | W | W | W | W | W |
| ChrX | 4.8 | W | W | W | W | W | W | W | W | W | W | W | W | W | W | W | W | W | W | W | W | W | W | W | W | W | W | W | W | W | W | W | W | W | W | W | W | W | W | W | W | W | W | W | W | W | W | W | W | W | W | W | W | W | W | W | W | W | W | W | W | W | W | W | W | W | W | W | W | W | W | W | W | W | W | W | W | W | W | W | W | W | W | W | W | W | W | W | W | W | W | W | W |
| ChrX | 4.9 | W | W | W | W | W | W | W | W | W | W | W | W | W | W | W | W | W | W | W | W | W | W | W | W | W | W | W | W | W | W | W | W | W | W | W | W | W | W | W | W | W | W | W | W | W | NA | W | W | W | W | W | W | W | W | W | W | W | W | W | W | W | W | W | W | W | W | W | W | W | W | W | W | W | W | W | W | W | W | W | W | W | W | W | W | W | W | W | W | W | W | W | W |
| ChrX | 5 | W | W | W | W | W | W | W | W | W | W | W | W | W | W | W | W | NA | W | W | W | W | W | NA | W | W | NA | W | W | W | W | W | W | W | W | W | W | W | W | W | W | W | W | W | W | W | W | W | W | W | W | W | W | NA | W | W | W | W | W | W | W | W | W | W | W | W | W | W | W | W | W | W | W | NA | W | W | W | W | W | W | W | W | W | W | W | W | W | W | W | W | W | W | W |
| ChrX | 5.1 | W | W | W | W | W | W | W | W | W | W | W | W | W | W | W | W | W | W | W | W | W | W | NA | W | W | W | W | W | W | W | W | W | W | W | W | W | W | W | W | W | W | W | W | W | W | W | W | W | W | W | W | W | W | W | W | W | W | W | W | W | W | W | W | W | W | W | W | W | W | W | W | W | W | W | W | W | W | W | W | W | W | W | W | W | W | W | W | W | W | W | W | W |
| ChrX | 5.2 | W | W | W | W | W | W | W | W | W | W | W | W | W | W | W | W | W | W | W | W | W | W | W | W | W | W | W | W | W | W | W | W | W | W | W | W | W | W | W | W | W | W | W | W | W | W | W | W | W | W | W | W | W | W | W | W | W | NA | W | W | W | W | W | W | W | W | W | W | W | W | W | W | W | W | NA | W | W | W | W | W | W | W | W | W | W | W | W | W | W | W | W | W |
| ChrX | 5.3 | W | W | W | W | W | W | W | NA | W | W | W | W | W | W | W | W | W | W | W | W | W | W | W | W | W | W | W | W | W | W | W | W | W | W | W | W | W | W | W | W | W | W | W | W | W | W | W | W | W | W | W | W | W | W | W | W | W | W | W | W | W | W | W | W | W | W | W | W | W | W | W | W | W | W | W | W | W | W | W | W | W | W | W | W | W | W | W | W | W | W | W | W |
| ChrX | 5.4 | W | W | W | NA | W | W | W | W | W | W | W | W | W | W | W | W | W | W | W | W | W | W | W | W | W | W | W | W | W | W | W | W | W | W | W | W | W | W | W | W | W | W | W | W | W | W | W | W | W | W | W | W | W | W | W | W | W | W | W | W | W | W | W | W | W | W | W | W | W | W | W | W | NA | W | W | W | W | W | W | W | W | W | W | W | W | W | W | W | W | W | W | W |
| ChrX | 5.5 | W | W | W | W | W | W | W | W | W | W | W | W | W | W | W | W | W | W | W | W | W | W | W | W | W | W | W | W | W | W | W | W | W | W | W | W | W | W | W | W | W | W | W | W | W | W | W | W | W | W | W | W | W | W | W | W | W | W | W | W | W | W | W | W | W | W | W | W | W | W | W | W | W | W | W | W | W | W | W | W | W | W | W | W | W | W | W | W | W | W | W | W |
| ChrX | 5.6 | W | W | W | W | W | W | W | W | W | W | W | W | W | W | W | W | W | W | W | W | W | W | W | W | W | W | W | W | W | W | W | W | W | W | W | W | W | W | W | W | W | W | W | W | W | W | W | W | W | W | W | W | W | W | W | W | W | W | W | W | W | W | W | W | W | W | W | W | W | W | W | W | W | W | W | W | W | W | W | W | W | W | W | W | W | W | W | W | W | W | W | W |
| ChrX | 5.7 | W | W | W | W | W | W | W | W | W | W | W | W | W | W | NA | W | W | W | W | W | W | W | W | W | W | W | W | W | W | W | W | W | W | W | W | W | W | W | W | W | W | W | W | W | W | W | W | W | W | W | W | W | W | W | W | W | W | W | W | W | W | W | W | W | W | W | W | W | W | W | W | W | W | W | W | W | W | W | W | W | W | W | W | W | W | W | W | W | W | W | W | W |
| ChrX | 5.8 | W | W | W | W | W | W | W | W | W | W | W | W | W | NA | W | W | W | W | W | W | W | W | W | W | W | W | W | W | W | W | W | W | W | W | W | W | W | W | W | W | W | W | W | W | W | W | W | W | W | W | W | W | W | W | W | W | W | W | W | W | W | W | W | W | W | W | W | W | W | W | W | W | W | W | W | W | W | W | W | W | W | W | W | W | W | W | W | W | W | W | W | W |
| ChrX | 5.9 | W | W | W | W | W | W | W | W | W | W | W | W | W | W | W | W | W | W | W | W | W | W | W | W | W | W | W | W | W | W | W | W | W | W | W | W | W | W | W | W | W | W | W | W | W | W | W | W | W | W | W | W | W | W | W | W | W | W | NA | W | W | W | W | NA | W | NA | W | W | W | W | W | W | W | W | W | W | W | W | W | W | W | W | W | W | W | W | W | W | W | W | W | W |
| ChrX | 6 | W | W | W | W | W | W | W | W | W | W | W | W | W | W | W | W | W | W | W | W | W | W | W | W | W | W | W | W | W | W | W | W | W | W | W | W | W | W | W | W | W | W | W | W | W | W | W | W | W | W | W | W | W | W | W | W | W | W | W | W | W | W | W | W | W | W | W | W | W | W | W | W | W | W | W | W | W | W | W | W | W | W | W | W | W | W | W | W | W | W | W | W |
| ChrX | 6.1 | W | W | W | W | W | W | W | W | W | W | W | W | W | W | W | W | W | W | W | W | W | W | W | W | W | W | W | W | W | W | W | W | W | W | W | W | W | W | W | W | W | W | W | W | W | W | W | W | W | W | W | W | W | W | W | W | W | W | W | W | W | W | W | W | W | W | W | W | W | W | W | W | W | W | W | W | W | W | W | W | W | W | W | W | W | W | W | W | W | W | W | W |
| ChrX | 6.2 | W | W | W | W | W | W | NA | W | W | W | W | W | W | W | W | W | W | W | W | W | W | W | W | W | W | W | W | W | W | W | W | W | W | W | W | W | W | W | W | W | W | W | W | W | W | W | W | W | W | NA | W | W | W | W | W | W | W | W | W | W | W | W | W | W | W | W | W | W | W | W | W | W | W | NA | W | W | W | W | W | W | W | W | W | W | W | W | W | W | W | W | W | W |
| ChrX | 6.3 | W | W | W | W | W | W | W | W | W | W | W | W | W | W | W | W | W | W | W | W | W | W | W | W | W | W | W | W | W | W | W | W | W | W | W | W | W | W | W | W | W | W | W | W | W | W | W | W | W | W | W | W | W | W | W | W | W | W | W | W | W | W | W | W | W | W | W | W | W | W | W | W | W | W | W | W | W | W | W | W | W | W | W | W | W | W | W | W | W | W | W | W |
| ChrX | 6.4 | W | W | W | W | W | W | W | W | W | W | W | W | W | W | W | W | W | W | W | W | W | W | W | W | W | W | W | W | W | W | W | W | W | W | W | W | W | W | W | W | W | W | W | W | W | W | W | W | W | W | W | W | W | W | W | W | W | W | W | W | W | W | W | W | W | W | W | W | W | W | W | W | W | W | W | W | W | W | W | W | W | W | W | W | W | W | W | W | W | W | W | W |
| ChrX | 6.6 | W | W | W | W | W | W | W | W | W | W | W | W | W | W | W | W | W | W | W | W | W | W | W | W | W | W | W | W | W | W | W | W | W | NA | W | W | W | W | W | W | W | W | W | W | W | W | W | W | W | W | W | W | W | W | W | W | W | W | W | W | W | W | W | W | W | W | W | W | W | W | W | W | W | W | W | W | W | W | W | W | W | W | W | W | W | W | W | W | W | W | W | W |
| ChrX | 6.7 | W | W | W | W | W | W | W | W | W | W | W | W | W | W | W | W | W | W | W | W | W | W | W | W | W | W | W | W | W | W | W | W | W | W | W | W | W | W | W | W | W | W | W | W | W | W | W | NA | W | W | W | W | W | W | W | W | W | W | W | W | W | W | W | W | W | W | W | W | W | W | W | W | W | W | W | W | W | W | W | W | W | W | W | W | W | W | W | W | W | W | W | W |
| ChrX | 6.8 | W | W | W | W | W | W | W | W | W | W | W | W | W | W | W | W | W | W | W | W | W | W | W | W | W | W | W | W | W | W | W | W | W | W | W | W | W | W | W | W | W | W | W | W | W | W | W | W | W | W | W | W | W | W | W | W | W | W | W | W | W | W | W | W | W | W | W | W | W | W | W | W | W | W | W | W | W | W | W | W | W | W | W | W | W | W | W | W | W | W | W | W |

| | | | | | | | | | | | | | | | | | | | | | | | | | | | | | | | | | | | | | | | | | | | | | | | | | | | | | | | | | | | | | | | | | | | | | | | | | | | | | | | | | | | | | | | | | | | | | |
|---|---|---|---|---|---|---|---|---|---|---|---|---|---|---|---|---|---|---|---|---|---|---|---|---|---|---|---|---|---|---|---|---|---|---|---|---|---|---|---|---|---|---|---|---|---|---|---|---|---|---|---|---|---|---|---|---|---|---|---|---|---|---|---|---|---|---|---|---|---|---|---|---|---|---|---|---|---|---|---|---|---|---|---|---|---|---|---|---|---|---|---|---|---|
| ChrX | 6.9 | W | W | W | W | W | W | W | W | W | W | W | W | W | W | W | W | W | W | W | W | W | W | W | W | W | W | W | W | W | W | W | W | W | W | W | W | W | W | W | W | W | W | W | W | W | W | W | W | W | W | W | W | W | W | W | W | W | W | W | W | W | W | W | W | W | W | W | W | W | W | W | W | W | W | W | W | W | W | W | W | W | W | W | W | W | W | W | W | W | W | W | W |
| ChrX | 7 | W | W | W | W | W | W | W | W | W | W | W | W | W | W | W | W | W | W | W | W | W | W | W | W | W | W | W | W | W | W | W | W | W | W | W | W | W | W | W | W | W | W | W | W | W | W | W | W | W | W | W | W | W | W | W | W | W | W | W | W | W | W | W | W | W | W | W | W | W | NA | W | W | W | W | W | W | W | W | W | W | W | W | W | W | W | W | W | W | W | W | W | W |
| ChrX | 7.1 | W | W | W | W | W | W | W | W | W | W | W | W | W | W | W | W | W | W | W | W | W | W | W | W | W | W | W | W | W | W | W | W | W | W | W | W | W | W | W | W | W | W | W | W | W | NA | W | W | W | W | W | W | W | W | W | W | W | W | W | W | W | W | W | W | W | W | W | W | W | W | W | W | W | W | W | W | W | W | W | W | W | W | W | W | W | W | W | W | W | W | W | W |
| ChrX | 7.2 | W | W | W | W | W | W | W | W | W | W | W | W | W | W | W | W | W | W | W | W | W | W | W | W | W | W | W | W | W | W | W | W | W | W | W | W | W | W | W | W | W | W | W | W | W | W | W | W | W | W | W | W | W | W | W | W | W | W | NA | W | W | W | W | W | W | W | W | W | W | W | W | W | W | W | W | W | W | W | W | W | W | W | W | W | W | W | W | W | W | W | W | W |
| ChrX | 7.3 | W | W | W | W | W | W | W | W | W | W | W | W | W | W | W | W | W | W | W | W | W | W | W | W | W | W | W | W | W | W | W | W | W | W | W | W | W | W | W | W | W | W | W | W | W | W | W | W | W | W | W | W | W | W | W | W | W | W | W | W | W | W | W | W | W | W | W | W | W | W | W | W | W | W | W | W | W | W | W | W | W | W | W | W | W | W | W | W | W | W | W | W |
| ChrX | 7.4 | W | W | W | W | W | W | W | W | W | W | W | W | W | W | NA | W | W | W | W | W | W | W | W | W | W | W | W | W | W | W | W | W | W | W | W | W | W | W | W | W | W | NA | W | W | W | W | W | W | W | W | NA | W | W | W | W | W | W | W | W | W | W | W | W | W | W | W | W | W | W | W | W | W | W | W | W | W | W | W | W | W | W | W | W | W | W | W | W | W | W | W | W | W |
| ChrX | 7.5 | W | W | W | W | W | W | W | W | W | W | W | W | W | W | W | W | W | W | W | W | W | W | W | W | W | W | W | W | W | W | W | W | W | W | W | W | W | W | W | W | W | W | W | W | W | W | W | W | W | W | W | W | W | W | W | W | W | W | W | W | W | W | W | W | W | W | W | W | W | W | W | W | W | W | W | W | W | W | W | W | W | W | W | W | W | W | W | W | W | W | W | W |
| ChrX | 7.6 | W | W | W | NA | W | W | W | W | W | W | W | W | W | W | W | W | W | W | W | W | W | W | W | W | W | W | W | W | W | W | W | W | W | NA | W | W | W | W | W | W | W | W | W | W | W | W | W | W | W | W | W | W | W | W | W | W | W | W | W | W | W | W | W | W | W | W | W | W | W | W | W | W | W | W | W | W | W | W | W | W | W | W | W | W | W | W | W | W | W | W | W | W |
| ChrX | 7.7 | W | W | W | W | W | W | W | W | W | W | W | W | W | W | W | W | W | W | W | W | W | NA | W | W | W | W | W | W | W | W | W | W | W | W | W | W | W | W | W | W | W | W | W | W | W | W | W | W | W | W | W | W | W | W | W | W | W | W | W | W | W | W | W | W | W | W | W | W | W | W | W | W | W | W | W | W | W | W | W | W | W | W | W | W | W | W | W | W | W | W | W | W |
| ChrX | 8.1 | W | W | W | W | W | W | NA | W | W | W | W | W | W | NA | NA | W | W | NA | W | W | W | W | W | W | W | W | W | W | W | W | W | W | W | W | W | W | W | W | W | W | W | W | W | W | W | W | W | NA | W | W | W | W | W | W | W | W | W | W | W | W | W | W | W | W | W | W | W | W | W | W | W | W | W | W | W | W | NA | W | W | W | W | W | W | W | W | W | W | W | W | W | W | W |
| ChrX | 8.2 | W | W | W | W | W | W | W | W | W | W | W | W | W | W | W | W | W | W | W | W | W | W | W | W | W | W | W | W | W | W | W | W | W | W | W | W | W | W | W | W | W | W | W | W | W | W | W | W | W | W | W | W | W | W | W | W | W | W | W | W | W | W | W | W | W | W | W | W | W | W | W | W | W | W | W | W | W | W | W | W | W | W | W | W | W | W | W | W | W | W | W | W |
| ChrX | 8.3 | W | W | W | W | W | W | W | W | W | W | W | W | W | W | W | W | W | W | W | W | W | W | W | W | W | W | W | W | W | W | W | W | W | W | W | W | W | W | W | W | W | W | W | W | W | W | W | W | W | W | W | W | W | W | W | W | W | W | W | W | W | W | W | W | W | W | W | W | W | W | W | W | W | W | W | W | W | W | W | W | W | W | W | W | W | W | W | W | W | W | W | W |
| ChrX | 8.4 | W | W | W | W | W | W | W | W | W | W | W | W | W | W | W | W | W | W | W | W | W | W | W | W | W | W | W | W | W | W | W | W | W | W | W | W | W | W | W | W | W | W | W | W | W | W | W | W | W | W | W | W | W | W | W | W | W | W | W | W | W | W | W | W | W | W | W | W | W | W | W | W | W | W | W | W | W | W | W | W | W | W | W | W | W | W | W | W | W | W | W | W |
| ChrX | 8.5 | W | W | W | W | W | W | W | W | W | W | W | W | W | W | W | W | W | W | W | W | W | W | W | W | W | W | W | W | W | W | W | W | W | W | W | W | W | W | W | W | W | W | W | W | W | W | W | W | W | W | W | W | W | W | W | W | W | W | W | W | W | W | W | W | W | W | W | W | W | W | W | W | W | W | W | W | W | W | W | W | W | W | W | W | W | W | W | W | W | W | W | W |
| ChrX | 8.6 | W | W | W | W | W | W | W | W | W | W | W | W | W | W | W | W | W | W | W | W | W | W | W | W | W | W | W | W | W | W | W | W | W | W | W | W | W | W | W | W | W | W | W | W | W | W | W | W | W | W | W | W | W | W | W | W | W | W | W | W | W | W | W | W | W | W | W | W | W | W | W | W | W | W | W | W | W | W | W | W | W | W | W | W | W | W | W | W | W | W | W | W |
| ChrX | 8.7 | W | W | W | W | W | W | W | W | W | W | W | W | W | W | NA | W | W | W | W | W | W | W | W | W | W | W | W | W | W | W | W | W | W | W | W | NA | W | W | W | W | W | W | W | W | W | W | W | W | W | W | W | W | W | W | W | W | W | W | W | W | W | W | W | W | W | W | W | W | W | W | W | W | W | W | W | W | W | W | W | W | W | W | W | W | W | W | W | W | W | W | W | W |
| ChrX | 8.8 | W | W | W | W | W | W | W | W | W | W | W | W | W | W | W | W | W | W | W | W | W | W | W | W | W | W | W | W | W | W | W | W | NA | W | W | W | W | W | W | W | W | W | W | W | W | W | W | W | W | W | W | W | W | W | W | W | W | W | W | W | W | W | W | W | W | W | W | W | W | W | W | W | W | W | W | W | W | W | W | W | W | W | W | W | W | W | W | W | W | W | W | W |
| ChrX | 8.9 | W | W | W | W | W | W | W | W | W | W | W | W | W | W | W | W | W | W | W | W | W | W | W | W | W | W | W | W | W | W | W | W | W | W | W | W | W | W | W | W | W | W | W | NA | W | W | W | W | W | W | W | W | W | W | W | W | W | W | W | W | W | W | W | W | W | W | W | W | W | W | W | W | W | W | W | W | W | W | W | W | W | W | W | W | W | W | W | W | W | W | W | W |
| ChrX | 9 | W | W | W | W | W | W | W | W | W | W | NA | W | W | NA | W | W | NA | W | W | W | W | W | W | W | W | W | NA | W | W | W | W | W | W | W | W | W | W | W | W | W | W | W | W | W | W | W | W | W | W | W | W | W | W | W | W | W | W | NA | W | W | W | NA | W | NA | W | NA | W | W | W | W | W | W | NA | W | W | W | W | W | W | W | W | W | W | W | W | NA | W | NA | W | W | W |
| ChrX | 9.1 | W | W | W | W | W | W | W | W | W | W | W | W | W | W | W | W | W | W | W | W | W | W | W | W | W | W | W | W | W | W | W | W | W | W | W | W | W | W | W | W | W | W | W | W | W | W | W | W | W | W | W | W | W | W | W | W | W | W | W | W | W | W | W | W | W | W | W | W | W | W | W | W | W | W | W | W | W | W | W | W | W | W | W | W | W | W | W | W | W | W | W | W |
| ChrX | 9.2 | W | W | W | W | W | W | W | W | W | W | W | W | W | W | W | W | W | W | W | W | W | W | W | W | W | W | W | W | W | W | W | W | W | W | W | W | W | W | W | W | W | W | W | W | W | W | W | W | W | W | W | W | W | W | W | W | W | W | W | W | W | W | W | W | W | W | W | W | W | W | W | W | W | W | W | W | W | W | W | W | W | W | W | W | W | W | W | W | W | W | W | W |
| ChrX | 9.3 | W | W | W | W | W | W | W | W | W | W | W | W | W | W | W | W | W | W | W | W | W | W | W | W | W | W | W | W | W | W | W | W | W | NA | W | W | W | W | W | W | W | W | W | W | W | W | W | W | W | W | W | W | W | W | W | W | W | W | W | W | W | W | W | W | W | W | W | W | W | W | W | W | W | W | W | W | W | W | W | W | W | W | W | W | W | W | W | W | W | W | W | W |
| ChrX | 9.4 | NA | W | W | NA | W | W | NA | W | W | W | W | W | W | W | W | W | W | W | W | W | W | W | W | W | W | W | W | W | W | W | W | W | W | W | W | W | W | W | W | W | W | W | W | W | NA | W | W | NA | W | W | W | W | W | W | W | W | W | W | NA | W | W | NA | W | NA | W | W | NA | W | W | W | W | W | W | W | W | W | NA | W | W | W | W | W | W | W | W | NA | NA | NA | NA | W | W | W |
| ChrX | 9.5 | W | W | W | W | W | W | W | W | W | W | W | W | W | W | W | W | W | W | W | W | W | W | W | W | W | W | W | W | W | W | W | W | W | W | W | W | W | W | W | W | W | W | W | W | W | W | W | W | W | W | W | W | W | W | W | W | W | W | W | W | W | W | W | W | W | W | W | W | W | W | W | W | W | W | W | W | W | W | W | W | W | W | W | W | W | W | W | W | W | W | W | W |
| ChrX | 9.6 | W | W | W | W | W | W | NA | W | W | W | W | W | W | W | W | W | W | W | W | NA | W | W | W | W | W | W | W | W | W | W | W | W | W | W | W | W | W | W | W | W | W | W | W | W | W | W | W | W | W | W | W | W | W | W | W | W | W | W | W | W | W | W | W | W | W | W | W | W | W | W | W | W | W | W | W | W | W | W | W | W | W | W | W | W | W | W | W | W | W | W | W | W |
| ChrX | 9.7 | W | W | W | W | W | W | W | W | W | W | W | W | W | W | W | W | W | W | W | W | W | W | W | W | W | W | W | W | W | W | W | W | W | W | W | W | W | W | W | W | W | W | W | W | W | W | W | W | W | W | W | W | W | W | W | W | W | W | W | W | W | W | W | W | W | W | W | W | W | W | W | W | W | W | W | W | W | W | W | W | W | W | W | W | W | W | W | W | W | W | W | W |
| ChrX | 9.8 | W | W | W | W | W | W | W | W | W | W | W | W | W | W | W | W | W | W | W | W | W | W | W | W | W | W | W | W | W | W | W | W | W | W | W | W | W | W | W | W | W | W | W | W | W | W | W | W | W | W | W | W | W | W | W | W | W | W | W | W | W | W | W | W | W | W | W | W | W | W | W | W | W | W | W | W | W | W | W | W | W | W | W | W | W | W | W | W | W | W | W | W |
| ChrX | 9.9 | W | W | W | W | W | W | W | NA | W | W | W | W | W | NA | W | W | W | W | W | W | W | W | W | W | W | W | W | W | W | W | W | W | W | W | W | W | W | W | W | W | W | W | W | W | W | W | W | W | W | W | W | W | W | W | W | W | W | W | W | NA | W | W | W | W | W | W | W | W | W | W | W | NA | W | W | W | W | W | W | W | W | W | W | W | W | W | W | W | W | W | W | W | W |
| ChrX | 10 | W | W | W | W | W | W | W | W | W | W | W | W | W | W | NA | W | W | W | W | W | W | W | NA | W | W | W | W | W | W | W | W | W | W | W | W | NA | W | W | W | W | W | W | W | W | W | W | W | W | W | W | W | W | W | W | W | W | W | NA | W | W | W | W | W | NA | W | W | W | W | W | W | W | W | W | W | W | W | W | W | W | W | W | W | NA | W | W | W | W | W | W | W | NA | W |
| ChrX | 10.1 | W | W | W | W | W | NA | W | W | W | W | NA | NA | W | W | W | W | W | NA | W | W | W | W | W | W | W | W | W | W | W | W | W | W | W | W | W | W | NA | W | W | NA | W | W | W | W | NA | W | W | W | W | W | W | NA | NA | W | W | W | W | NA | NA | NA | W | W | W | NA | NA | W | W | W | W | NA | W | W | NA | NA | W | W | W | W | W | NA | W | W | W | NA | W | NA | W | W | NA | NA | W | W |
| ChrX | 10.2 | W | W | W | W | W | W | W | W | W | W | W | W | W | W | W | W | W | W | W | NA | W | W | W | W | W | W | W | W | W | W | W | W | W | W | W | W | W | W | W | W | W | W | W | W | W | W | W | W | W | W | W | W | W | W | W | W | W | W | W | W | W | W | W | W | W | W | W | W | W | W | W | W | W | W | W | W | W | W | W | W | W | W | W | W | W | W | W | W | W | W | W | W |
| ChrX | 10.3 | W | W | W | W | W | W | W | W | W | W | W | W | W | W | W | W | W | W | W | W | W | W | W | W | W | W | W | W | W | W | W | W | W | W | W | W | W | W | W | W | W | W | W | W | W | W | W | W | W | W | W | W | W | W | W | W | W | W | W | W | W | W | W | W | W | W | W | W | W | W | NA | W | W | W | W | W | W | W | W | W | W | W | W | W | W | W | W | W | W | W | W | W |
| ChrX | 10.4 | W | W | W | W | W | W | W | W | W | W | W | W | W | W | W | W | W | W | W | W | W | W | W | W | W | W | W | W | W | W | W | W | W | W | W | W | W | W | W | W | W | W | W | W | W | W | W | W | W | W | W | W | W | W | W | W | W | W | W | W | W | W | W | W | W | W | W | W | W | W | W | W | W | W | W | W | W | W | W | W | W | W | W | W | W | W | W | W | W | W | W | W |
| ChrX | 10.5 | W | W | W | W | W | W | W | W | W | W | W | W | W | W | W | W | W | W | W | W | W | W | W | W | W | W | W | W | W | W | W | W | W | W | W | W | W | W | W | W | W | W | W | W | W | W | W | W | W | W | W | W | W | W | W | W | W | W | W | W | W | W | W | W | W | W | W | W | W | W | W | W | W | W | W | W | W | W | W | W | W | W | W | W | W | W | W | W | W | W | W | W |
| ChrX | 10.6 | W | W | W | W | W | W | W | W | W | W | W | W | W | W | W | W | W | W | W | W | W | W | W | W | W | W | W | W | W | W | W | W | W | W | W | W | W | W | W | W | W | W | W | W | W | NA | W | W | W | W | W | W | W | W | W | W | W | W | W | W | W | W | W | W | W | W | W | W | W | W | W | W | W | W | W | W | W | W | W | W | W | W | W | W | W | W | W | W | W | W | W | W |
| ChrX | 10.8 | W | W | W | NA | W | W | W | W | W | W | NA | W | NA | NA | W | W | W | W | W | W | W | NA | NA | W | W | NA | W | NA | W | W | W | W | W | W | W | W | W | W | W | W | W | W | W | W | W | NA | W | NA | W | NA | W | W | W | W | W | W | W | W | NA | W | W | W | W | W | W | W | W | W | W | NA | W | W | NA | W | W | W | NA | W | W | W | NA | W | W | NA | W | W | NA | W | W | W | W | W |
| ChrX | 10.9 | W | W | W | W | W | W | W | W | W | W | W | W | W | W | NA | W | W | W | W | W | W | W | W | W | W | W | W | W | W | W | W | W | W | W | W | W | W | W | W | W | W | W | W | W | W | W | W | W | W | W | W | W | W | W | W | W | W | W | W | W | W | W | W | W | W | W | W | W | W | W | W | W | W | W | W | W | W | W | W | W | W | W | W | W | W | W | W | W | W | W | W | W |
| ChrX | 11 | W | W | W | W | W | W | W | W | W | W | W | W | W | NA | W | W | W | W | W | W | W | W | NA | W | W | W | W | W | W | W | W | W | W | W | W | W | W | W | W | W | W | W | W | W | W | W | W | W | W | NA | W | W | W | W | NA | W | W | W | W | W | W | W | W | W | W | W | W | W | W | W | W | W | W | W | W | W | W | W | W | W | W | W | W | W | W | W | W | W | W | W | W | W |
| ChrX | 11.1 | W | W | W | NA | W | W | W | W | W | W | W | W | W | NA | NA | W | W | W | W | W | W | W | W | W | W | NA | W | W | W | W | W | W | W | W | W | W | W | W | W | W | W | W | W | W | NA | W | W | W | W | W | W | NA | W | W | NA | W | W | W | W | W | W | W | W | W | W | W | W | W | W | W | W | W | W | NA | W | W | W | W | W | W | W | W | W | W | W | W | W | W | W | W | W | W |
| ChrX | 11.2 | W | W | W | W | W | W | W | W | W | W | W | W | W | W | W | W | W | W | W | W | W | W | W | W | W | W | W | W | W | W | W | W | W | W | W | W | W | W | W | W | W | W | W | W | W | W | W | W | W | W | W | W | W | W | W | W | W | W | W | W | W | W | W | W | W | W | W | W | W | W | W | W | W | W | W | W | W | W | W | W | W | W | W | W | W | W | W | W | W | W | W | W |
| ChrX | 12 | W | W | W | W | W | W | W | W | W | W | W | W | W | W | W | W | W | W | W | W | W | W | W | W | W | W | W | W | W | W | W | W | W | W | W | W | W | W | W | W | W | W | W | W | W | W | W | W | W | W | W | W | W | W | W | W | W | W | W | W | W | W | W | W | W | W | W | W | W | W | W | W | W | W | W | W | W | W | W | W | W | W | W | W | W | W | W | W | W | W | W | W |
| ChrX | 12.1 | W | W | W | W | W | W | W | W | W | W | W | W | W | W | W | W | W | W | W | W | W | W | W | W | W | W | W | W | W | W | W | W | W | W | W | W | W | W | W | W | W | W | W | W | W | W | W | W | W | W | W | W | W | W | W | W | W | W | W | W | W | W | W | W | W | W | W | W | W | W | W | W | W | W | W | W | W | W | W | W | W | W | W | W | W | W | W | W | W | W | W | W |
| ChrX | 12.2 | W | W | W | W | W | W | W | W | W | W | W | W | W | W | W | W | W | W | W | W | W | W | W | W | W | W | W | W | W | W | W | W | W | W | W | W | W | W | W | W | W | W | W | W | W | W | W | W | W | W | W | W | W | W | W | W | W | W | W | W | W | W | W | W | W | W | W | W | W | W | W | W | W | W | W | W | W | W | W | W | W | W | W | W | W | W | W | W | W | W | W | W |
| ChrX | 12.3 | W | W | W | W | W | W | W | W | W | W | W | W | W | W | W | W | W | W | W | W | W | W | W | W | W | W | W | W | W | W | W | W | W | W | W | W | W | W | W | W | W | W | W | W | W | W | W | W | W | W | W | W | W | W | W | W | W | W | W | W | W | W | W | W | W | W | W | W | W | W | W | W | W | W | W | W | W | W | W | W | W | W | W | W | W | W | W | W | W | W | W | W |
| ChrX | 12.4 | W | W | W | W | W | W | NA | W | W | W | W | W | W | W | NA | W | W | W | W | W | W | W | W | W | W | W | W | W | W | W | W | W | W | W | W | W | W | W | W | W | W | W | W | W | W | W | W | W | W | W | W | W | W | W | W | W | W | NA | W | W | W | W | W | W | W | W | W | W | W | W | W | W | W | W | W | W | W | W | W | W | W | W | W | W | W | W | W | W | W | W | W | W |
| ChrX | 12.6 | W | W | W | W | W | W | W | W | W | W | W | W | W | NA | W | W | W | W | W | W | W | W | W | W | W | W | W | W | W | W | W | W | W | W | W | W | W | W | W | W | W | W | W | W | W | W | W | W | W | W | W | W | W | W | W | W | W | W | W | W | W | W | W | W | W | W | W | W | W | W | W | W | W | W | W | W | W | W | W | W | W | W | W | W | W | W | W | W | W | W | W | W |
| ChrX | 12.7 | W | W | W | W | W | W | W | W | W | W | W | W | W | W | W | W | W | W | W | W | W | W | W | W | W | W | W | W | W | W | W | W | W | W | W | W | W | W | W | W | W | W | W | W | W | W | W | W | W | W | W | W | W | W | W | W | W | W | W | W | W | W | W | W | W | W | W | W | W | W | W | W | W | W | W | W | W | W | W | W | W | W | W | W | W | W | W | W | W | W | W | W |
| ChrX | 12.8 | W | W | W | W | W | W | W | W | W | W | W | W | W | NA | W | W | W | W | W | W | W | W | W | W | W | W | W | W | W | W | W | W | W | NA | W | W | W | W | W | W | W | W | W | W | W | W | W | W | W | W | W | W | W | W | W | W | W | W | W | W | W | W | W | W | W | W | W | W | W | W | W | W | W | W | W | W | W | W | W | W | W | W | W | W | W | W | W | W | W | W | W | W |
| ChrX | 12.9 | W | W | W | W | W | W | NA | W | W | W | W | W | W | NA | W | W | W | W | W | W | W | W | W | W | W | W | W | W | W | W | W | W | W | W | W | W | W | W | W | W | W | W | W | W | W | W | W | W | W | W | W | W | W | W | W | W | W | W | W | W | W | W | W | W | W | W | W | W | W | W | W | W | W | W | W | W | W | W | W | W | W | W | W | W | W | NA | W | W | W | W | W | W |
| ChrX | 13 | W | W | W | W | W | W | W | W | W | W | W | W | W | NA | NA | W | W | W | W | W | W | W | W | W | W | W | W | W | W | W | W | W | W | W | W | W | W | W | W | W | W | W | W | W | NA | NA | W | NA | W | NA | W | W | W | W | W | W | W | NA | W | W | W | W | W | W | W | W | W | W | W | W | W | W | NA | W | W | W | W | W | W | W | W | W | NA | NA | W | W | W | W | W | W | W | W |
| ChrX | 13.1 | W | W | W | W | W | W | W | W | W | W | W | W | W | W | W | W | W | W | W | W | W | W | NA | W | W | W | W | W | W | W | W | W | W | W | W | NA | W | W | W | W | W | W | W | W | W | W | W | W | W | W | W | W | W | W | NA | W | W | W | W | W | W | W | W | W | NA | W | W | W | W | W | W | W | W | W | W | W | W | W | W | W | W | NA | W | W | W | W | W | W | W | W | W | W |
| ChrX | 13.2 | W | W | W | W | W | W | W | W | W | W | W | W | W | NA | W | W | W | W | W | W | W | W | W | W | W | W | W | W | W | W | W | W | W | W | W | W | W | W | W | W | W | W | W | W | W | W | W | W | W | W | W | W | W | W | W | W | W | W | W | W | W | W | W | W | W | W | W | W | W | W | W | W | W | W | W | W | W | W | W | W | W | W | W | W | W | W | W | W | W | W | W | W |
| ChrX | 13.3 | W | W | W | W | W | W | W | W | W | W | W | W | W | W | W | W | W | W | W | W | W | W | W | W | W | W | W | W | W | W | W | W | W | W | W | W | W | W | W | W | W | W | W | W | W | W | W | W | W | W | W | W | W | W | W | W | W | W | W | W | W | W | W | W | W | W | W | W | W | W | W | W | W | W | W | W | W | W | W | W | W | W | W | W | W | W | W | W | W | W | W | W |
| ChrX | 13.4 | W | W | W | W | W | W | W | W | W | W | W | W | W | W | W | W | W | NA | W | W | W | W | NA | W | W | W | W | W | W | W | W | W | W | NA | W | W | W | W | W | W | W | W | W | W | W | NA | W | W | W | W | W | W | W | W | W | W | W | W | NA | NA | W | W | W | W | W | W | W | W | W | W | W | W | W | W | W | W | W | NA | W | W | W | W | W | W | W | W | W | W | W | W | W | W |
| ChrX | 13.5 | W | W | W | NA | W | W | W | W | W | W | W | W | NA | NA | NA | W | W | W | W | W | W | NA | W | W | W | W | W | W | W | NA | W | W | W | W | W | NA | NA | W | W | NA | W | W | W | W | W | NA | W | W | W | W | W | NA | W | W | W | NA | W | NA | W | W | W | W | W | NA | W | W | W | W | W | W | W | NA | W | NA | NA | W | W | NA | W | W | W | W | W | W | W | W | W | W | W | W | NA | W |
| ChrX | 13.6 | W | W | W | W | W | W | W | W | W | W | W | W | W | NA | W | W | W | W | W | W | W | W | W | W | NA | W | W | W | W | W | W | W | W | W | W | W | W | W | W | W | W | W | W | W | W | W | W | NA | W | W | W | W | W | W | NA | W | W | NA | W | W | W | W | W | W | W | NA | W | W | W | W | W | NA | W | W | W | W | NA | W | W | W | W | W | W | W | W | W | W | W | W | W | W | W |
| ChrX | 13.7 | NA | W | W | NA | W | NA | NA | W | NA | W | NA | NA | W | W | NA | W | W | W | W | W | W | NA | NA | W | W | W | W | W | W | NA | W | W | W | W | W | NA | NA | W | W | NA | W | W | W | W | W | NA | W | W | W | W | W | NA | NA | W | W | NA | W | W | W | W | NA | W | W | NA | NA | W | W | W | W | NA | W | W | NA | W | W | W | W | NA | W | W | W | NA | W | NA | W | W | W | W | W | W | NA | W |
| ChrX | 14.2 | W | W | W | W | W | W | W | W | W | W | W | W | W | W | W | W | W | W | W | W | W | W | W | W | W | W | W | W | W | W | W | W | W | W | W | W | W | W | W | W | W | W | W | W | W | W | W | W | W | W | W | W | W | W | W | W | W | W | W | W | W | W | W | W | W | W | W | W | W | W | W | W | W | W | W | W | W | W | W | W | W | W | W | W | W | W | W | W | W | W | W | W |
| ChrX | 14.3 | W | W | W | W | W | W | W | W | W | W | W | W | W | W | W | W | W | W | W | W | W | W | W | W | W | W | W | W | W | W | W | W | W | W | W | W | W | W | W | W | W | W | W | W | W | W | W | W | W | W | W | W | W | W | W | W | W | W | W | W | W | W | W | W | W | W | W | W | W | W | W | W | W | W | W | W | W | W | W | W | W | W | W | W | W | W | W | W | W | W | W | W |
| ChrX | 14.4 | W | W | W | W | W | W | W | W | W | W | W | W | W | W | W | W | W | W | W | W | W | W | W | W | W | W | W | W | W | W | W | W | W | W | W | W | W | W | W | W | W | W | W | W | W | W | W | W | W | W | W | W | W | W | W | W | W | W | W | W | W | W | W | W | W | W | W | W | W | W | W | W | W | W | W | W | W | W | W | W | W | W | W | W | W | W | W | W | W | W | W | W |
| ChrX | 14.5 | W | W | W | W | W | W | W | W | W | W | W | W | W | W | NA | W | W | W | W | W | W | W | W | W | W | W | W | W | W | W | W | W | W | W | W | W | W | W | W | W | W | W | W | W | W | W | W | W | W | W | W | W | W | W | W | W | W | W | W | W | W | W | W | W | W | W | W | W | W | W | W | W | W | W | W | W | W | W | W | W | W | W | W | W | W | W | W | W | W | W | W | W |
| ChrX | 14.6 | W | W | W | W | W | W | W | W | W | W | W | W | W | W | W | W | W | W | W | W | W | W | W | W | W | W | W | W | W | W | W | W | W | W | W | W | W | W | W | W | W | W | W | W | W | W | W | W | W | W | W | W | W | W | W | W | W | W | W | W | W | W | W | W | W | W | W | W | W | W | W | W | W | W | W | W | W | W | W | W | W | W | W | W | W | W | W | W | W | W | W | W |
| ChrX | 14.7 | W | W | W | W | W | W | W | W | W | W | W | W | W | W | W | W | W | W | W | W | W | W | W | W | W | W | W | W | W | W | W | W | W | W | W | W | W | W | W | W | W | W | W | W | W | W | W | W | W | W | W | W | W | W | W | W | W | W | W | W | W | W | W | W | W | W | W | W | W | W | W | W | W | W | W | W | W | W | W | W | W | W | W | W | W | W | W | W | W | W | W | W |
| ChrX | 14.8 | W | W | W | W | W | W | W | W | W | W | W | W | W | W | W | W | W | W | W | W | W | W | W | W | W | W | W | W | W | W | W | W | W | W | W | W | W | W | W | W | W | W | W | W | W | W | W | W | W | W | W | W | W | W | W | W | W | W | W | W | W | W | W | W | W | W | W | W | W | W | W | W | W | W | W | W | W | W | W | W | W | W | W | W | W | W | W | W | W | W | W | W |
| ChrX | 14.9 | W | W | W | W | W | W | W | W | W | W | W | W | W | W | W | W | W | W | W | W | W | W | W | W | W | W | W | W | W | W | W | W | W | NA | W | W | W | W | W | W | W | W | W | W | W | NA | W | W | W | W | W | W | W | W | W | W | W | W | W | W | W | W | W | W | W | W | W | W | W | W | W | W | W | W | W | W | W | W | W | W | W | W | W | W | W | W | W | W | W | W | W | W |
| ChrX | 15 | W | W | W | W | W | W | W | W | W | W | W | W | W | W | W | W | W | W | W | W | W | W | W | W | W | W | W | W | W | W | W | W | W | W | W | W | W | W | W | W | W | W | W | W | W | W | W | W | W | W | W | W | W | W | W | W | W | W | W | W | W | W | W | W | W | W | W | W | W | W | W | W | W | W | W | W | W | W | W | W | W | W | W | W | W | W | W | W | W | W | W | W |
| ChrX | 15.1 | W | W | W | W | W | W | W | W | W | W | W | W | W | W | W | W | W | W | W | W | W | W | W | W | W | W | W | W | W | W | W | W | W | W | W | W | W | W | W | W | W | W | W | W | W | W | W | W | W | W | W | W | W | W | W | W | W | W | W | W | W | W | W | W | W | W | W | W | W | W | W | W | W | W | W | W | W | W | W | W | W | W | W | W | W | W | W | W | W | W | W | W |
| ChrX | 15.2 | W | W | W | W | W | W | W | W | W | W | W | W | W | W | W | W | W | W | W | W | W | W | W | W | W | W | W | W | W | W | W | W | W | W | W | W | W | W | W | W | W | W | W | W | W | W | W | W | W | W | W | W | W | W | W | W | W | W | W | W | W | W | W | W | W | W | W | W | W | W | W | W | W | W | W | W | W | W | W | W | W | W | W | W | W | W | W | W | W | W | W | W |
| ChrX | 15.3 | W | W | W | W | W | W | W | W | W | W | W | W | W | W | W | W | W | W | W | W | W | W | W | W | W | W | W | W | W | W | W | W | W | W | W | W | W | W | W | W | W | W | W | W | W | W | W | W | W | W | W | W | W | W | W | W | W | W | W | W | W | W | W | W | W | W | W | W | W | W | W | W | W | W | W | W | W | W | W | W | W | W | W | W | W | W | W | W | W | W | W | W |
| ChrX | 15.4 | W | W | W | W | W | W | W | W | W | W | W | W | W | NA | NA | W | W | W | W | W | W | W | W | W | W | W | W | W | W | W | W | W | W | W | W | W | W | W | W | W | W | W | W | W | W | W | W | W | W | W | W | W | W | W | W | W | W | W | W | W | W | W | W | W | W | W | W | W | W | W | W | W | W | W | W | W | W | W | W | W | W | W | W | W | W | W | W | W | W | W | W | W |
