## Supplementary material for "Analysis of meiosis in *Pristionchus pacificus* reveals plasticity in homolog pairing and synapsis in the nematode lineage": Table S7 Crossover frequencies

| CHR | MB | cM | map |
|---|---|---|---|
| ChrI | 0.1 | 0 | XO |
| ChrI | 0.8 | 1.098083146 | XO |
| ChrI | 1.3 | 3.819874666 | XO |
| ChrI | 1.5 | 6.541666307 | XO |
| ChrI | 2.5 | 7.628751058 | XO |
| ChrI | 2.8 | 11.2740264 | XO |
| ChrI | 2.9 | 13.07227996 | XO |
| ChrI | 3.4 | 15.24758018 | XO |
| ChrI | 3.9 | 16.33468598 | XO |
| ChrI | 4.1 | 18.5099711 | XO |
| ChrI | 5 | 21.77548897 | XO |
| ChrI | 8.3 | 25.04101776 | XO |
| ChrI | 9 | 26.12812331 | XO |
| ChrI | 9.3 | 27.21525134 | XO |
| ChrI | 13.3 | 28.30237938 | XO |
| ChrI | 21.7 | 29.38950741 | XO |
| ChrI | 25 | 30.47663545 | XO |
| ChrI | 30.4 | 31.56376348 | XO |
| ChrI | 31.8 | 32.65089152 | XO |
| ChrI | 32.5 | 33.73801955 | XO |
| ChrI | 32.9 | 34.8255418 | XO |
| ChrI | 33 | 35.91305293 | XO |
| ChrI | 34.6 | 38.08834925 | XO |
| ChrI | 34.7 | 40.26959714 | XO |
| ChrI | 35 | 42.45082271 | XO |
| ChrI | 35.3 | 44.63099567 | XO |
| ChrI | 35.5 | 47.90145704 | XO |
| ChrI | 35.8 | 48.98921988 | XO |
| ChrI | 36.1 | 50.07700526 | XO |
| ChrI | 36.3 | 51.16412217 | XO |
| ChrI | 37.7 | 53.33942957 | XO |
| ChrI | 37.9 | 54.42667783 | XO |
| ChrI | 38.1 | 55.5139372 | XO |
| ChrI | 38.6 | 56.60008638 | XO |
| ChrII | 0.1 | 0 | XO |
| ChrII | 1.5 | 1.098234194 | XO |
| ChrII | 2.7 | 2.185483783 | XO |
| ChrII | 3.5 | 4.361323522 | XO |
| ChrII | 3.7 | 5.449352638 | XO |
| ChrII | 4.2 | 8.715297185 | XO |
| ChrII | 5.2 | 10.89084828 | XO |
| ChrII | 6.4 | 11.97823079 | XO |

| | | | |
|---|---|---|---|
| ChrII | 6.7 | 13.06549017 | XO |
| ChrII | 7.1 | 14.15273842 | XO |
| ChrII | 9.3 | 16.32804582 | XO |
| ChrII | 14.6 | 17.41514026 | XO |
| ChrII | 16.6 | 20.68068022 | XO |
| ChrII | 17 | 22.85594679 | XO |
| ChrII | 17.4 | 25.76337259 | XO |
| ChrII | 17.5 | 27.20936809 | XO |
| ChrII | 17.8 | 28.29688315 | XO |
| ChrII | 18.2 | 29.9283389 | XO |
| ChrII | 18.4 | 31.56011395 | XO |
| ChrII | 18.7 | 32.64756602 | XO |
| ChrII | 19.1 | 33.7348029 | XO |
| ChrII | 19.3 | 37.00413162 | XO |
| ChrII | 19.4 | 38.09322235 | XO |
| ChrII | 20.5 | 44.65588202 | XO |
| ChrII | 20.7 | 46.83543897 | XO |
| ChrII | 21.4 | 49.01532012 | XO |
| ChrII | 22.1 | 50.10172427 | XO |
| ChrIII | 0.4 | 0 | XO |
| ChrIII | 4.5 | 1.086149174 | XO |
| ChrIII | 9.4 | 2.173277209 | XO |
| ChrIII | 13.8 | 3.260371113 | XO |
| ChrIII | 14.1 | 7.620976483 | XO |
| ChrIII | 14.8 | 10.88816526 | XO |
| ChrIII | 15 | 11.97535724 | XO |
| ChrIII | 15.1 | 13.06260548 | XO |
| ChrIII | 15.3 | 15.23870628 | XO |
| ChrIII | 15.5 | 18.50506385 | XO |
| ChrIII | 16.4 | 19.59215827 | XO |
| ChrIII | 16.5 | 21.76742008 | XO |
| ChrIII | 16.7 | 27.22523422 | XO |
| ChrIII | 16.9 | 28.31363291 | XO |
| ChrIII | 17.4 | 34.87804031 | XO |
| ChrIII | 18 | 38.14856919 | XO |
| ChrIII | 18.7 | 48.49905368 | XO |
| ChrIII | 18.9 | 51.34779356 | XO |
| ChrIII | 20 | 53.52301471 | XO |
| ChrIII | 20.1 | 53.52306604 | XO |
| ChrIII | 20.7 | 54.60922047 | XO |
| ChrIV | 0.4 | 0 | XO |
| ChrIV | 1.1 | 1.086411918 | XO |
| ChrIV | 6.8 | 2.173802695 | XO |

| | | | |
|---|---|---|---|
| ChrIV | 13.3 | 3.260930727 | XO |
| ChrIV | 13.9 | 4.348058761 | XO |
| ChrIV | 20.5 | 5.435318136 | XO |
| ChrIV | 20.8 | 6.522577511 | XO |
| ChrIV | 22.8 | 7.609683547 | XO |
| ChrIV | 22.9 | 8.697331862 | XO |
| ChrIV | 23.2 | 10.87319342 | XO |
| ChrIV | 23.4 | 11.96031033 | XO |
| ChrIV | 23.6 | 13.04781876 | XO |
| ChrIV | 23.8 | 16.31374508 | XO |
| ChrIV | 24.7 | 20.23909827 | XO |
| ChrIV | 25 | 22.84589881 | XO |
| ChrIV | 25.4 | 24.29236043 | XO |
| ChrIV | 26.2 | 27.20027346 | XO |
| ChrIV | 26.5 | 29.3755623 | XO |
| ChrIV | 26.6 | 30.46267387 | XO |
| ChrIV | 27 | 32.07551415 | XO |
| ChrIV | 27.8 | 37.00460484 | XO |
| ChrIV | 28 | 38.09167013 | XO |
| ChrIV | 28.6 | 41.3571875 | XO |
| ChrIV | 29.3 | 45.71511541 | XO |
| ChrV | 0 | 0 | XO |
| ChrV | 0.7 | 2.174326521 | XO |
| ChrV | 1.4 | 4.352059125 | XO |
| ChrV | 2.6 | 7.622935097 | XO |
| ChrV | 2.8 | 11.43779111 | XO |
| ChrV | 3.2 | 11.43780111 | XO |
| ChrV | 4.1 | 15.25269079 | XO |
| ChrV | 3.9 | 15.25270079 | XO |
| ChrV | 4.6 | 18.5239176 | XO |
| ChrV | 5.9 | 19.61655817 | XO |
| ChrV | 6.7 | 20.70435365 | XO |
| ChrV | 7.2 | 21.79148168 | XO |
| ChrV | 18.5 | 22.87899016 | XO |
| ChrV | 18.9 | 26.14534297 | XO |
| ChrV | 19.1 | 27.23350367 | XO |
| ChrV | 19.4 | 29.40961664 | XO |
| ChrV | 20.1 | 30.49738552 | XO |
| ChrV | 20.3 | 32.67512791 | XO |
| ChrV | 20.5 | 37.03788144 | XO |
| ChrV | 20.9 | 38.12659702 | XO |
| ChrV | 21.2 | 40.30181589 | XO |
| ChrV | 21.3 | 40.30187074 | XO |

| | | | |
|---|---|---|---|
| ChrV | 21.6 | 44.66076308 | XO |
| ChrV | 21.9 | 45.74917588 | XO |
| ChrV | 22.1 | 47.92580411 | XO |
| ChrV | 22.8 | 50.10013064 | XO |
| ChrI | 0.8 | 0 | XX |
| ChrI | 1.5 | 2.174337562 | XX |
| ChrI | 2 | 3.261443361 | XX |
| ChrI | 2.5 | 5.436745371 | XX |
| ChrI | 2.7 | 6.5242565 | XX |
| ChrI | 2.9 | 7.611738112 | XX |
| ChrI | 3.7 | 13.06886721 | XX |
| ChrI | 3.9 | 14.15663616 | XX |
| ChrI | 5 | 15.24375307 | XX |
| ChrI | 5.2 | 17.41906047 | XX |
| ChrI | 5.8 | 18.50617738 | XX |
| ChrI | 6 | 19.5938339 | XX |
| ChrI | 6.3 | 20.68147928 | XX |
| ChrI | 6.7 | 22.85784569 | XX |
| ChrI | 7.7 | 25.03368001 | XX |
| ChrI | 8.3 | 26.12079692 | XX |
| ChrI | 9.3 | 27.20792495 | XX |
| ChrI | 13.3 | 28.29505299 | XX |
| ChrI | 21.5 | 29.38218102 | XX |
| ChrI | 27 | 30.46944065 | XX |
| ChrI | 27.5 | 31.55894752 | XX |
| ChrI | 29.2 | 32.64830627 | XX |
| ChrI | 30.7 | 34.82360825 | XX |
| ChrI | 31.8 | 35.91069189 | XX |
| ChrI | 32.8 | 38.82068139 | XX |
| ChrI | 33 | 40.26989442 | XX |
| ChrI | 33.3 | 41.35701878 | XX |
| ChrI | 33.5 | 42.44414681 | XX |
| ChrI | 33.8 | 43.53127485 | XX |
| ChrI | 34.2 | 44.61839177 | XX |
| ChrI | 34.6 | 46.79368814 | XX |
| ChrI | 34.8 | 48.97004958 | XX |
| ChrI | 35 | 50.05819697 | XX |
| ChrI | 35.8 | 52.23509718 | XX |
| ChrI | 36.1 | 54.4120418 | XX |
| ChrI | 37.7 | 55.49817984 | XX |
| ChrII | 0.1 | 0 | XX |
| ChrII | 1.9 | 4.140024627 | XX |
| ChrII | 2.7 | 5.489809836 | XX |

| | | | |
|---|---|---|---|
| ChrII | 4.9 | 7.667128426 | XX |
| ChrII | 5.2 | 10.38950243 | XX |
| ChrII | 6.4 | 13.10932908 | XX |
| ChrII | 7.1 | 14.19644034 | XX |
| ChrII | 9.3 | 15.28353427 | XX |
| ChrII | 16.6 | 19.64242847 | XX |
| ChrII | 17 | 21.81769106 | XX |
| ChrII | 17.5 | 23.99298745 | XX |
| ChrII | 19.1 | 25.08005832 | XX |
| ChrII | 19.4 | 30.54007827 | XX |
| ChrII | 19.6 | 36.00581552 | XX |
| ChrII | 20.5 | 40.37037932 | XX |
| ChrII | 20.7 | 43.6357205 | XX |
| ChrII | 21.4 | 43.63585212 | XX |
| ChrII | 22.1 | 46.90139091 | XX |
| ChrII | 22.3 | 49.0757062 | XX |
| ChrIII | 0.4 | 0 | XX |
| ChrIII | 0.8 | 1.086412081 | XX |
| ChrIII | 1.4 | 2.174197329 | XX |
| ChrIII | 2 | 3.261708578 | XX |
| ChrIII | 2.3 | 5.438081494 | XX |
| ChrIII | 2.5 | 6.526525501 | XX |
| ChrIII | 3.1 | 7.613916567 | XX |
| ChrIII | 3.3 | 8.701022112 | XX |
| ChrIII | 3.6 | 11.96656207 | XX |
| ChrIII | 4 | 14.14237966 | XX |
| ChrIII | 4.3 | 15.2305544 | XX |
| ChrIII | 4.5 | 16.31819741 | XX |
| ChrIII | 5 | 18.49349379 | XX |
| ChrIII | 5.4 | 20.66877916 | XX |
| ChrIII | 5.9 | 22.84406453 | XX |
| ChrIII | 6.7 | 25.01936092 | XX |
| ChrIII | 7.3 | 26.10647783 | XX |
| ChrIII | 14.1 | 27.19373724 | XX |
| ChrIII | 14.3 | 28.2812512 | XX |
| ChrIII | 14.7 | 30.45709066 | XX |
| ChrIII | 14.8 | 31.54447321 | XX |
| ChrIII | 15.5 | 32.63159012 | XX |
| ChrIII | 16.3 | 34.80689753 | XX |
| ChrIII | 16.9 | 35.89401444 | XX |
| ChrIII | 18 | 36.98111995 | XX |
| ChrIII | 18.2 | 40.25037082 | XX |
| ChrIII | 18.9 | 43.51959543 | XX |

| | | | |
|---|---|---|---|
| ChrIII | 20 | 47.87902564 | XX |
| ChrIII | 20.1 | 48.96666285 | XX |
| ChrIII | 20.8 | 50.05281202 | XX |
| ChrIV | 0.4 | 0 | XX |
| ChrIV | 0.7 | 4.357949839 | XX |
| ChrIV | 1.2 | 5.446248589 | XX |
| ChrIV | 2.4 | 7.62476613 | XX |
| ChrIV | 2.5 | 8.874192091 | XX |
| ChrIV | 3 | 15.16423111 | XX |
| ChrIV | 3.6 | 16.428725 | XX |
| ChrIV | 4 | 17.51814208 | XX |
| ChrIV | 4.4 | 19.15103446 | XX |
| ChrIV | 4.6 | 20.78984799 | XX |
| ChrIV | 4.8 | 22.96872965 | XX |
| ChrIV | 5.7 | 25.15921679 | XX |
| ChrIV | 6.8 | 28.4371642 | XX |
| ChrIV | 9.9 | 29.52426961 | XX |
| ChrIV | 13.3 | 30.61139765 | XX |
| ChrIV | 16.9 | 31.69852568 | XX |
| ChrIV | 20.2 | 32.78565372 | XX |
| ChrIV | 22.8 | 33.87278175 | XX |
| ChrIV | 23.2 | 34.95990979 | XX |
| ChrIV | 23.8 | 36.04703782 | XX |
| ChrIV | 24.7 | 37.1342972 | XX |
| ChrIV | 25 | 38.22155103 | XX |
| ChrIV | 25.8 | 39.85406716 | XX |
| ChrIV | 26.2 | 41.48657218 | XX |
| ChrIV | 26.5 | 43.66187409 | XX |
| ChrIV | 27.8 | 44.74801214 | XX |
| ChrV | 0 | 0 | XX |
| ChrV | 0.7 | 1.086138036 | XX |
| ChrV | 1.4 | 3.26332312 | XX |
| ChrV | 2.6 | 5.805095402 | XX |
| ChrV | 2.8 | 8.346795316 | XX |
| ChrV | 3.2 | 9.615033234 | XX |
| ChrV | 4.1 | 10.88322239 | XX |
| ChrV | 4.4 | 14.56374201 | XX |
| ChrV | 4.9 | 18.68329164 | XX |
| ChrV | 6.7 | 20.70748875 | XX |
| ChrV | 7.2 | 21.79460722 | XX |
| ChrV | 14.7 | 22.88173525 | XX |
| ChrV | 14.9 | 23.96885217 | XX |
| ChrV | 15.5 | 26.14415957 | XX |

| | | | |
|---|---|---|---|
| ChrV | 16.4 | 27.231531 | XX |
| ChrV | 17.2 | 29.40737042 | XX |
| ChrV | 17.4 | 30.49475296 | XX |
| ChrV | 18.6 | 31.58213189 | XX |
| ChrV | 18.7 | 33.72608052 | XX |
| ChrV | 18.9 | 38.11949319 | XX |
| ChrV | 20.1 | 39.20807967 | XX |
| ChrV | 20.4 | 40.47678547 | XX |
| ChrV | 20.5 | 41.74544403 | XX |
| ChrV | 20.9 | 46.84934687 | XX |
| ChrV | 22.4 | 49.02461238 | XX |
| ChrV | 23.5 | 50.1227012 | XX |
| ChrX | 0.5 | 0 | XX |
| ChrX | 1.6 | 8.784172407 | XX |
| ChrX | 2.3 | 10.95928115 | XX |
| ChrX | 2.5 | 10.95933247 | XX |
| ChrX | 2.8 | 20.86971182 | XX |
| ChrX | 4.5 | 36.58519792 | XX |
| ChrX | 5.5 | 38.76032907 | XX |
| ChrX | 6.3 | 40.93562546 | XX |
| ChrX | 7.3 | 42.02273126 | XX |
| ChrX | 7.5 | 44.19803866 | XX |
| ChrX | 8.2 | 45.28512147 | XX |
| ChrX | 9.1 | 49.64307132 | XX |
