## Supplementary material for "Analysis of meiosis in *Pristionchus pacificus* reveals plasticity in homolog pairing and synapsis in the nematode lineage": Table S8 Crossovers per chromosome

| | | XX | | | XO | |
|---|---|---|---|---|---|---|
| | crossovers | **count** | map length (cM) | crossovers | **count** | map length (cM) |
| **ChrI** | 2 | 3 | 55.5 | 2 | 1 | 56.6 |
| | 1 | 45 | | 1 | 50 | |
| | 0 | 44 | | 0 | 41 | |
| **ChrII** | 1 | 45 | 49.1 | 1 | 46 | 50.1 |
| | 0 | 47 | | 0 | 46 | |
| **ChrIII** | 1 | 46 | 50.1 | 1 | 50 | 54.6 |
| | 0 | 46 | | 0 | 42 | |
| **ChrIV** | 2 | 0 | 44.74 | 2 | 1 | 45.7 |
| | 1 | 41 | | 1 | 40 | |
| | 0 | 51 | | 0 | 51 | |
| **ChrV** | 2 | 1 | 50.1 | 2 | 0 | 50.1 |
| | 1 | 44 | | 1 | 46 | |
| | 0 | 47 | | 0 | 46 | |
| **ChrX** | 1 | 45 | 49.64 | | | |
| | 0 | 47 | | | | |
